# Cell surface markers identify astrocyte subpopulations in the adult hippocampus with a heterogeneous response to aging

**DOI:** 10.1101/2025.11.07.685575

**Authors:** Lucía Casares-Crespo, Marife Cano-Jaimez, Raquel Anta, Alba Ripollés-Boix, Fabián Robledo-Yagüe, Sheila Zúñiga-Trejos, Helena Mira

## Abstract

Astrocyte diversity is currently expanding both between and within specific brain regions. Here, we assessed the spatial distribution and transcriptomic profile of two hippocampal astrocyte subpopulations, defined by combinatorial expression of the cell surface astrocyte markers ACSA-1 or GLAST/SLC1A3, and ACSA-2 or ATP1B2. Fluorescence activated cell sorting and genome-wide transcriptomics by bulk RNAseq uncovered distinct transcriptional signatures of the two astrocyte subsets and highlighted heterogeneous responses during aging. The most abundant ATP1B2/GLAST double-positive astrocytes corresponded to mature glial cells with increased protein glycosylation and stable gene expression patterns. Signatures related to mitochondrial respiration and cholesterol metabolism were induced during aging in ATP1B2 single-positive astrocytes, while cell adhesion genes from the γ-protocadherin cluster were repressed in double-positive astrocytes. Heterochronic co-culture assays with primary neurons show the loss of synaptogenic function of old ATP1B2/GLAST astrocytes. Our results complement previous studies demonstrating the presence of morphological and molecular astrocyte heterogeneity within the hippocampus, and uncover differences among astrocyte subsets in their transcriptomic response to aging.

## Introduction

Astrocytes play many essential roles in brain homeostasis. They provide trophic and metabolic support to neurons, regulate ionic balance, actively modulate synapses, participate in myelination and contribute to the blood-brain barrier. Recent studies reveal unexpected high levels of astrocyte heterogeneity, challenging the historical view of astrocytes as a uniform cell population. Boosted by the advent of single-cell RNA sequencing (sc-RNAseq) and spatial transcriptomics, astrocyte diversity continues to emerge both between and within specific brain areas (Chai et al., 2017; Morel et al., 2017; Boisvert et al., 2018; Lanjakornsiripan et al., 2018; Batiuk et al. 2020; Bayraktar et al., 2020; Borggrewe et al. 2021; Kantzer et al. 2021; Karpf et al., 2022, Su et al., 2022; De Ceglia et al. 2023; Viana et al. 2023; Zhou et al., 2025; Bocchi et al., 2025). To what extent astrocytes clustered according to their individual transcriptome are actually specialized cell subtypes with a distinct function and developmental trajectory is currently unknown (Hennes et al., 2025; Williamson et al., 2025). Exciting studies are shedding light on the origin of astrocyte diversity (Hochstim et al., 2008; Tsai et al., 2012; Molofsky et al., 2014; Liu et al Sparc Development 2022, Clavreul et al., 2022, Markey et al., 2023; Xie et al., 2025; Zhou et al., 2025); however, most transcriptomic data still lack developmental perspective, functional validation, or both. Thus, additional efforts are needed to support astrocyte subtype classifications and to uncover how astrocyte diversity translates into functional specialization.

Astrocyte cell membrane proteins have been instrumental in the identification and isolation of central nervous system glial subpopulations (Lin et al., 2017; Morel et al., 2019; Borggrewe et al. 2021; Kantzer et al. 2021). In a function-based approach, differential expression of glutamate transporter-1 (GLT-1), involved in clearing excess glutamate from the synapse, allowed to uncover transcriptionally distinct cortical astrocytes with different electrophysiological properties (Morel et al., 2019). Expression of glutamate-aspartate transporter (GLAST) and another cell membrane protein, the astrocyte marker ATPase, Na+/K+ transporting, beta 2 polypeptide ATP1B2, also distinguished astrocyte subpopulations with specific transcriptional profiles in the forebrain, spinal cord, hindbrain and cerebellum (Borggrewe et al. 2021; Kantzer et al. 2021). An extracellular ATP1B2 epitope is recognized by the widely employed astrocyte cell surface antigen-2 (ACSA-2) (Kantzer et al., 2017), allowing for the isolation of astrocytes from freshly dissected tissue. One report showed that hindbrain and spinal cord GLAST-positive and GLAST-negative ATP1B2/ACSA-2-positive astrocytes respond differently to neuroinflammatory insults (Borggrewe et al. 2021), linking classifications based on the astrocyte cell surface proteome to gene expression profiles and functional responses. Age-related astrocyte changes stemming from GLAST and ATP1B2 cell classifications have not been addressed so far.

During aging, astrocytes undergo transcriptomic alterations that reveal a decline in their normal physiological functions (Boisvert et al., 2018; Clarke et al., 2018; Pan et al., 2020). These changes might contribute to cognitive impairment and vulnerability to neurodegenerative diseases (Labarta-Bajo and Allen, 2025). The hippocampus is one of the brain regions most susceptible to age-related degeneration (Bettio et al., 2017; Hernández-Frausto and Vivar, 2024), but astrocyte heterogeneity and astrocyte functional changes during normal aging in the hippocampal formation remain underexplored (Clarke et al., 2018; Habib et al., 2020; Wu et al., 2025). To better understand astrocyte diversity and alterations in the young and old hippocampus, in this study we characterized ATP1B2 and GLAST expression at the mRNA and protein level in different hippocampal subfields during aging. We FACS-isolated two astrocyte subpopulations on the basis of ATP1B2/ACSA-2 and GLAST/ACSA-1 cell surface protein expression and analysed their transcriptome by bulk RNAseq. The results uncover subpopulation-specific transcriptomic signatures and a differential impact of aging on the gene expression landscape of the astrocyte subpopulations. Repression of structural synapse formation genes in ATP1B2 and GLAST double-positive astrocytes reveals a negative impact on excitatory synaptogenesis. This age-related loss of function was confirmed in heterochronic hippocampal astrocyte-neuron co-culture assays.

## Methods

### Animals

All experimental procedures were approved by the Institutional Animal Care and Use Committee of Instituto de Biomedicina de Valencia and by the Bioethical Committee of CSIC and CIPF (protocol 2023-VSC-PEA-0212 and 2024/VSC/PEA/0094). Mice were maintained in the animal facility on a 12h/12h light/ dark cycle under constant temperature (23 °C) with food and water provided *ad libitum*. Wild-type C57BL/6JRccHsd mice were purchased from Inotiv (https://www.inotivco.com/model/c57bl-6jrcchsd) and wild-type RjOrl:SWISS mice from Janvier Labs (https://janvier-labs.com/en/fiche_produit/swiss_mouse/).

Male C57Bl6/JRccHsd mice from in-house breeding were used throughout unless otherwise stated. Data for RNAseq analysis was collected from astrocytes isolated from 2 and 18-months-old mice by FACS (in quadruplicate and triplicate, respectively). Mice aged to 2, 18 and 24 months-old were used for MACS and ISH (*in situ* hybridization) experiments (three replicates of each age). For immunohistochemistry assays, wild-type RjOrl:SWISS male mice aged 2 and 18 months-old were used (three replicates of each age). Neurons were isolated by MACS from embryonic day 18 (E18) RjOrl:SWISS mice. Finally, for lectin flow cytometry, 5 months-old RjOrl:SWISS male mice were used.

### Astrocyte isolation for FACS

Cervical dislocation was used as euthanasia method FACS procedures. Brains were removed and hippocampi were dissected in cold Hank’s balanced salt solution (HBSS, Sigma-Aldrich, 55021C), under a binocular microscope. The myelin present in hippocampi was discarded in order to clean them as much as possible before the dissociation step. Hippocampal cell suspension was obtained using the Neural Tissue Dissociation Kit (Trypsin, Miltenyi, 130-093-231) in combination with the gentleMACS™ Octo Dissociator (Miltenyi, 130-095-937). The resulting suspension was then filtered through a 70 μm MACS®SmartStrainers (Miltenyi, 130-098-462) with HBSS buffer with Ca^2+^ and Mg^2+^ (HBSS, Sigma-Aldrich, 55037C) to remove any remaining clumps. Following myelin and red blood cells removal, the resulting cell pellet was resuspended in 100 µl of phosphate-buffered saline (PBS) (without Ca^2+^/Mg^2+^) containing 0.5% bovine serum albumin (BSA) (Sigma-Aldrich) and processed immediately for FACS.

All steps were performed at 4 °C. To prevent unspecific antibody binding, cells were incubated with FcR Blocking Reagent mouse (Miltenyi, 130-092-575) for 10 minutes before antibody labeling. Single cell hippocampal cell-suspensions were then co-stained with ACSA-2-FITC (Miltenyi Biotec, 130-116-243) (1:50 dilution) and ACSA-1(GLAST)-PE (Miltenyi Biotec, 130-118-483) (1:50 dilution) and incubated for 1h on ice in the dark. Next, cells were washed and collected in a new tube with PBS supplemented with 0.5% BSA. Cell debris and dead cells were excluded by 4′,6-diamidino-2-phenylindole (DAPI) staining (1:1000 dilution). FACS was performed on a BD FACSAria™ III Cell Sorter using a 100 μm nozzle. Compensations were done on single-color controls and gates were set on unstained samples. For the gating, we used side scatter *versus* forward scatter to select the population and forward scatter to exclude doublets (Supplementary Fig 2). Viable astrocytes (DAPI^-^) were selected and two distinct astrocyte subpopulations were sorted: ACSA-2^+^ (A^+^) and ACSA-1^+^/ACSA-2^+^ (A^+^/G^+^) cells. Afterwards, cells were spun down at 300g for 5 min at 4°C. Supernatant was removed and 5 µL of Cell Lysis Buffer from NEBNext® Low Input RNA Library Prep Kit for Illumina® (New England Biolabs, NEB #E6420S/L) was added. Finally, cells were flashed-frozen and immediately store at −80°C until library preparation. Flow cytometry data was analysed using FlowJo Software (v10.1).

### Lectin Flow Cytometry

Hippocampal cells from young RjOrl:SWISS mice were dissociated as explained previously and labeled with lectins (1:400 FITC-conjugated L-PHA) (Vector Labs), GLAST-PE (1:100) and ACSA-2-APC (1:100) in 0.5% BSA during 30 minutes on ice and protected from light. After washing 3 times with 1x PBS, cells were re-suspended in PBS containing DAPI, which was used to exclude non-viable cells for analysis. Cells were assessed for fluorescence in a FACS MACSQuant®X (Miltenyi). For each sample a total of 10,000 cells was analysed. Mean fluorescence intensity (MFI) of lectins was quantified for A^+^/G^+^ and A^+^ cells of each mice.

### RNA isolation and RNA sequencing

The RNAseq experiment was conducted by the Multigenic Analysis Unit from the UCIM-INCLIVA (University of Valencia, Valencia, Spain). RNAseq libraries were made from total RNA isolated from sorted cells using NEBNext® Low Input RNA Library Prep Kit for Illumina® (New England Biolabs, NEB #E6420S/L) following manufacturer’s instructions. Briefly, we started the procedure directly from cells, the sample that had the least was 84 cells and the one that had the most, was 6193 cells. For the cDNA amplification, the number of cycles was adjusted to the input. cDNA was quantified using a Bioanalyzer High Sensitivity DNA Analysis (Agilent). Sequencing libraries were manually generated and adaptor’s enrichment and ligation was performed. Then, quality and quantity of libraries was assessed again with High Sensitivity DNA Analysis Chip (Agilent). Libraries were then sequenced on the Illumina® NextSeq 550 platform to generate 2 x 75 bp paired end reads using the Illumina® NSQ 500/550 Hi Output KT v2.5 (150 CYS) (Illumina, San Diego, CA), according to the manufacturer’s protocol. Libraries were sequenced on average to a depth of 14 M reads per library.

### Analysis of RNA sequencing data

Bioinformatics analysis was conducted by the Bioinformatics Unit at INCLIVA. Sequence quality control was performed with FastQC v0.11.8 (Andrews et al. 2010). Low-quality bases and reads, as well as possible adapters, were removed with fastp v0.20.1 (Chen et al. 2018). Read mapping and expression quantification per isoform were performed using the STAR alignment (Dobin et al. 2012) and RSEM (Li et al. 2011), with the default parameters and the reference genome GRCm39. Data was loaded into R using the tximport package (Soneson et al. 2015). Prior to the exploratory analysis and differential expression study, all those genes with less than 2TPM in fewer than 3 samples were removed. To assess the quality of the samples an exploratory analysis was performed on the normalized counts that were obtained with the VST function of DESeq2 v.1.38.2 (Love et al. 2014) applying the option *blind=FALSE*. The differential expression study was performed with the DESeq2 package. A Benjamini-Hochberg adjusted p-value cutoff of 0.05 was set to select differentially expressed genes. Box and bar plots were generated with ggplot2 (Wickham, 2016). The volcano plots were generated with EnhancedVolcano v1.16.0 (Blighe et al. 2022). Clustering heatmaps were prepared with the pheatmap package (Kolde, 2019). Functional enrichment was performed using Enrichr search engine (https://maayanlab.cloud/Enrichr/) (Xie et al. 2021). A circular plot was created with functional enriched terms with the circlize package v0.4.16.

### Astrocyte isolation for MACS

Cervical dislocation was used as euthanasia method MACS procedures. Brains were removed and hippocampi were dissected in cold D-PBS (GIBCO, 14287080), under a binocular microscope. The myelin present in hippocampi was discarded in order to clean the tissue as much as possible before the dissociation step. Hippocampal cell suspension was obtained using the Adult Brain Dissociation Kit (Miltenyi, 130-107-677) in combination with the gentleMACS™ Octo Dissociator (Miltenyi, 130-095-937). The resulting suspension was then filtered through a 70 μm MACS®SmartStrainers (Miltenyi, 130-098-462) with D-PBS to remove any remaining clumps. Following myelin and red blood cells removal, the resulting cell pellet was resuspended in 100 µl of phosphate-buffered saline (PBS) (without Ca^2+^/Mg^2+^) containing 0.5% bovine serum albumin (BSA) (Sigma-Aldrich) and processed immediately. Astrocytes (ACSA-2^+^ cells) were positively selected using Anti-ACSA-2 MicroBead Kit (Miltenyi, 130-097-678) following manufacturer’s procedure. Briefly, cells were incubated with 10 μL of FcR Blocking Reagent for 10 minutes at 4°C, followed by incubation with 10 μL of ACSA-2 MicroBeads for 15 minutes at 4°C. Cells were then washed with 1 mL 0.5% BSA in PBS buffer and centrifuged at 300xg for 5 minutes to remove excess beads from the solution. Pellet was resuspended with 500 µl of PBS 0.5% BSA and loaded onto an MS Column (Miltenyi), which was placed in the magnetic field of a MiniMACS™ Separator (Miltenyi). The magnetically labeled ACSA-2-positive cells were retained within the column and eluted as the positively selected cell fraction after removing the column from the magnet. After centrifugation, ACSA-2^+^ cells were resuspended with pre-warmed AstroMACS Medium (130-117-031) and seeded in 24-well laminin-coated dishes (80,000-100,000 astrocytes/well). Cell culture was maintained by replacing 50% of AstroMACS Medium every other day. After 14 days in culture, cells were fixed with 4% PFA during 10 minutes.

### Neuron Isolation for MACS

E18 RjOrl:SWISS mice were decapitated. Brains were removed and hippocampi were dissected in cold D-PBS (GIBCO, 14287080), under a binocular microscope. Hippocampal cell suspension was obtained using the Neural Tissue Dissociation Kit (Miltenyi, 130-092-628) in combination with the gentleMACS™ Octo Dissociator (Miltenyi, 130-095-937). The resulting suspension was then filtered through a 70 μm MACS®SmartStrainers (Miltenyi, 130-098-462) with D-PBS to remove any remaining clumps. Neurons and non-neurons were selected using Neuron Isolation Kit (Miltenyi, 130-115-389) following manufacturer’s procedure. Briefly, cells were incubated with 20 μL of Non-Neuronal Cell Biotin-Antibody Cocktail for 5 minutes at 4°C. Cells were then washed with 1 mL 0.5% BSA in PBS buffer and centrifuged at 300xg for 5 minutes. Pellet was resuspended with 80 µl of PBS 0.5% BSA and cells were incubated with 20 μL of Anti-Biotin MicroBeads for 10 minutes at 4°C. Then, the volume was adjusted to 500 μL and loaded onto a LS Column (Miltenyi), which was placed in the magnetic field of a MidiMACS™ Separator (Miltenyi). The magnetically labeled non-neuronal cells were retained in the column and neurons were collected from the eluted cell fraction.

### Neuron and astrocyte heterochronic co-culture

Neurons were obtained as explained before and, after negative magnetic isolation, were plated on 24-well poly-l-lysine and laminin-coated dishes (100,000 neurons/well). Cells were maintained in MACS Neuro Medium (Miltenyi, 130-093-570) supplemented with L-glutamine (0.5 mM), 2% v/v Neuro Brew-21 (Miltenyi, 130-093-566) and 1% penicillin/streptomycin (P/S). On the first day post-plating, araC (2 µM) was added to reduce non-neuronal contamination. The following day, a media change was performed to remove araC. On the third day post-seeding, astrocytes isolated from the adult hippocampus (2 and 18 months-old mice) were plated on top of neurons (100.000 ACSA-2^+^/well). Cell culture was maintained by replacing 50% of the medium every four days. After 8 days of co-culture, cells were fixed with 4% PFA during 10 minutes.

### Immunostaining

Immunohistochemistry and immunocytochemistry were performed using standard procedures. Animals were anesthetized with pentobarbital (80mg/Kg) and perfused with PB 0.1M and 4% PFA followed by 16 h postfixation in 4% PFA at 4°C. The brains were coronally sectioned in a vibratome (Leica Microsystems VT-1200-S). The resulting 40 μm free floating sections were collected sequentially generating antero-posterior reconstructions of the hippocampus conformed by 1 section every 320 μm of hippocampal structure. Stereology was performed by the analysis of at least 3 coronal sections, 40 μm each, separated 320 μm one to each other. Sampling started at first appearance of the infrapyramidal blade of the dentate gyrus.

Samples (tissue and cells) were incubated with blocking solution (10% Fetal Bovine Serum in 0.1M phosphate buffer with or without 0.5% Triton X-100 depending on the cellular location of the epitope of interest). Primary antibodies used for the staining were as follows: ATP1B2 (1:300, Alomone Labs, ANP-012), GLAST (ACSA-1) biotin (1:50, Miltenyi, 130-119-161), S100β (1:250, Synaptic Systems, 287004), GFAP (1:1000, Synaptic Systems, 173004), VGLUT1 (1:500, GeneTex, GTX133148), PSD95 (1:200, Fisher, MA1045), Anti-Pan-gamma-protocadherin (10μg, NeuroMab, clone N159/5) and Anti-gamma-protocadherin-C3 (10μg, NeuroMab, clone N174B/27). Then, samples were washed with PB and incubated with secondary antibodies at room temperature for 1 or 2 hours (cells and tissue, respectively). Secondary antibodies were Cy3 donkey anti-rabbit (1:500, Jackson, 711-165-152), Alexa Fluor 488 donkey anti-guinea pig (1:500, Jackson, 706-545-148), Alexa Fluor 647 donkey anti-guinea pig (1:500, Jackson, 706-605-148), Alexa Fluor 555 donkey anti-mouse (1:500, Invitrogen, A31570), Alexa Fluor 647 donkey anti-rabbit (1:500, Invitrogen, A31573) and streptavidin-Cy2 (green) (1:200, Jackson, 016-220-084). Nuclei were counterstained with DAPI 10 µg/ml (Sigma-Aldrich, D9542). After staining, cells and all sections were mounted and preserved with 50% Mowiol (Polysciences, 17951), 2.5% DABCO (Sigma, D2522). Images were acquired with a Leica Spectral SP8 confocal microscope with 40x or 63x Oil objectives. Images were analysed with Fiji Image J Software (1.53q).

### Synapse quantification

Excitatory synapses were visualized with presynaptic marker VGLUT1 and with postsynaptic marker PSD95. The Puncta Analyzer Plugin for ImageJ (Ippolito and Eroglu, 2010), was used to count the number of co-localized synaptic puncta. This quantification method is based on the fact that pre- and post-synaptic proteins are not within the same cellular compartments and would appear co-localized only at synapses due to their close proximity. We took images with a Leica SP8 confocal laser-scanning microscope at ×63 to image synapses. Only neurons in direct contact with young or old astrocytes (GFAP^+^) were analysed. We quantified pre- and post-synaptic co-localized puncta in 5 different areas of the preparation (15 neurons per culture and experimental condition). We reported the final data as the average number of synapses per neuron (DAPI^+^/GFAP^-^).

### RNAscope fluorescence in situ hybridization

RNAscope (Advanced Cell Diagnostics, ACD) was performed as follows. Briefly, brains were quickly frozen in Optimum Cutting Temperature compound (Tissue-Tek), using liquid nitrogen.

Fifteen-micrometer thick brain slices were prepared using a NX70 cryostat (Thermo Fisher Scientific). Sections were subsequently fixed in ice-cold 4% paraformaldehyde for 30 min. Sections were then dehydrated using a series of ethanol solutions (50–100%), before drying and incubating with Protease IV for 20 min at room temperature. Slides were washed in phosphate-buffered saline and hybridized with gene-specific probes (Supplementary Table 1) for 2 h at 40°C in a HybEZ Oven (ACD). Non-annealed probes were removed by washing sections in 1× proprietary wash buffer. Probes were then detected via sequential hybridization of proprietary amplifiers and labeled “secondary” probes (Amp 1–Amp 4). Finally, sections were stained with 4′,6-diamidino-2-phenylindole (DAPI) and mounted using ProLong Diamond Antifade Mountant (Life Technologies).

### Statistics

All statistical tests and sample sizes are included in the Figure Legends. Normality was measured with Shapiro Wilks test. Normal data are shown as mean ± SEM and non-normal data are represented as median ± interquartile range. In all cases, p-values are represented as follows: **** p<0.0001, *** p<0.001, ** p<0.01, * p<0.05. For normal data, all quantifications were statistically analysed using either one sample t-test (normalized data 2 groups), Student’s t-test-2 tails (2 groups) or 2 way-ANOVA tests. For non-normal data, Mann Whitney or Kruskal-Wallis test was employed. Statistical analysis was performed using Graphpad Prism 8. No statistical methods were used to pre-determine sample sizes.

## Data availability

The RNAseq raw data and associated metadata generated in this study have been deposited in the ArrayExpress database (EMBL-EBI) under the accession number E-MTAB-15811.

## Results

### ATP1B2 and GLAST identify two astrocyte subpopulations in the adult mouse hippocampus that are preserved during aging

To investigate astrocyte diversity in the adult hippocampus during lifespan, we quantified the number of ATP1B2^+^ single-positive and ATP1B2^+^/GLAST^+^ double-positive cells among the S100β^+^ astrocytes in young (Y, 2 months) and old (O, 18 months) wild-type mice by immunofluorescence and confocal microscopy (Figure 1a-b). We first calculated the percentage of ATP1B2^+^ (A^+^) and ATP1B2^+^/GLAST^+^(A^+^/G^+^) S100β^+^ cells in the hippocampus as a whole at both ages. The A^+^/G^+^ astrocyte fraction was higher than the A^+^ fraction in young and old mice (average±SEM, Y-A^+^/G^+^: 82.6±4.9% and O-A^+^/G^+^: 84.0±6.7 *versus* Y-A^+^: 17.4±4.9% and O-A^+^: 16.0±6.7%) (Figure 1a). We then analysed astrocyte heterogeneity across the main non-neuronal layers of the hippocampal subfields *Cornu Ammonis* area 1 (CA1) and dentate gyrus (DG), including stratum oriens (SO), stratum radiatum (SR), stratum lacunosum moleculare (SLM), molecular layer (ML) and hilus (Figure 1c-d). Again, the majority of astrocytes were A^+^/G^+^, with similar proportions across layers. There were no significant differences with aging (Figure 1d).

**Figure 1.**
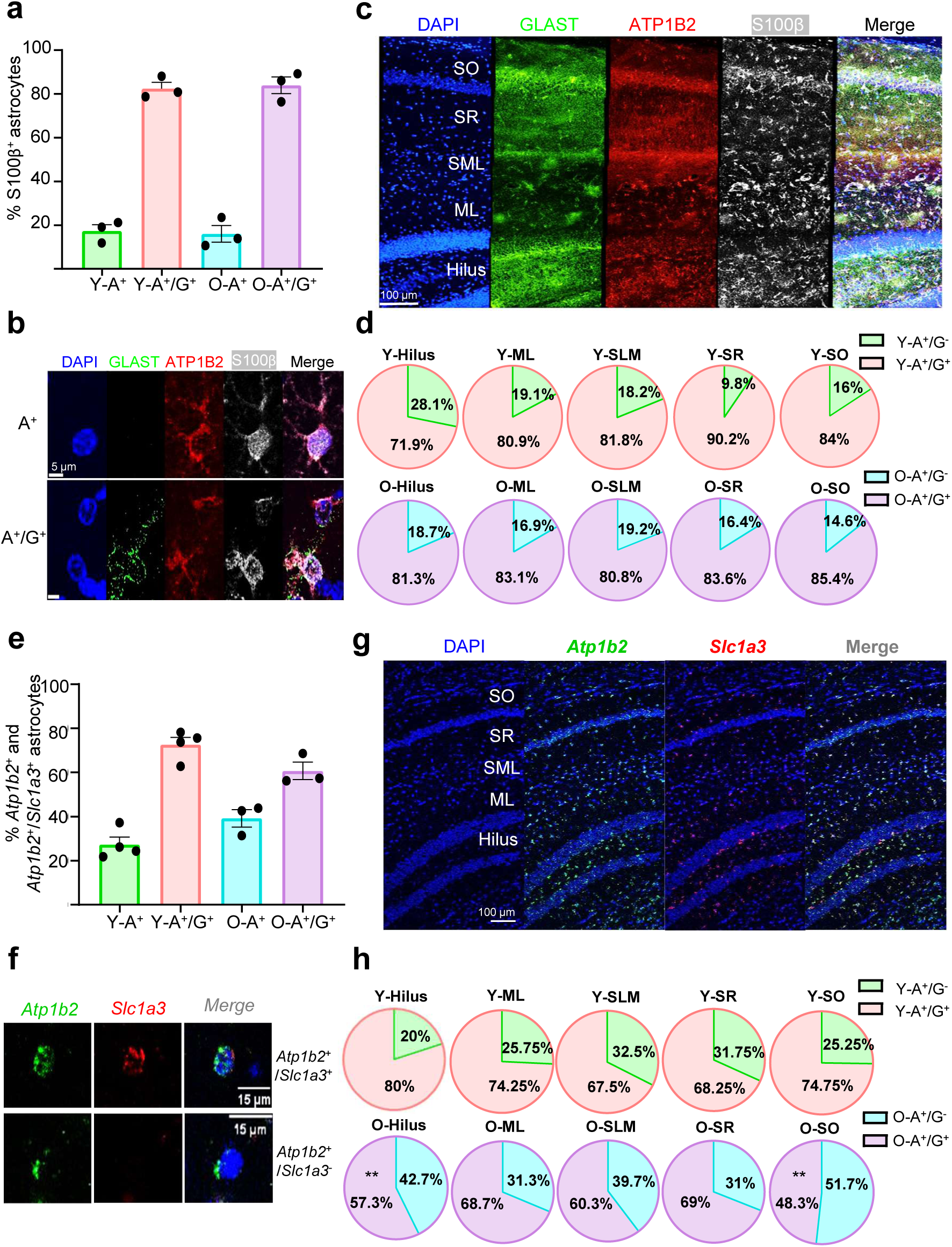
Astrocyte subpopulation analysis in the young and old hippocampus based on GLAST and ATP1B2 protein and RNA expression. **(a)** Quantification of astrocyte subsets in hippocampus of young (Y, 2m) and old (O, 18m) wild-type mice by immunohistochemistry. Percentage of S100β^+^/ATP1B2^+^/GLAST^+^ (A^+^/G^+^) and S100β^+^/ATP1B2^+^/GLAST^-^ (A^+^) astrocytes is shown as mean ± SEM. Three independent animals of each age were analysed (n=3). **(b)** Representative images of the two astrocyte subsets with S100β (white), GLAST (green) and ATP1B2 (red) biomarkers. Scale bar = 5 µm. **(c)** Representative immunohistochemistry of astrocyte subpopulations (S100β (white), GLAST (green) and ATP1B2 (red)), in the following astrocyte-enriched hippocampal layers: stratum oriens (SO), stratum radiatum (SR), stratum lacunosum moleculare (SLM), molecular layer (ML), and hilus. Scale bar = 100 µm. **(d)** Pie charts showing the proportions of the two subpopulations of astrocytes (A^+^ and A^+^/G^+^) in each hippocampal layer, in young and old mice, by immunohistochemistry. **(e)** Quantification of astrocyte subsets in hippocampal tissue of young (Y, 2m) and old wild-type mice (O, 18m) by RNAscope. Percentage of *Atp1b2*^+^/*Slc1a3*^+^ (A^+^/G^+^) and *Atp1b2*^+^/*Slc1a3*^-^ (A^+^) astrocytes is shown as mean ± SEM. At least three independent animals of each strain and age were analysed (n≥3). **(f)** Representative images of the two astrocyte subtypes identified, with *Slc1a3* (red) and *Atp1b2* (green) probes. Scale bar = 15 µm. **(g)** Representative RNAscope of astrocyte subpopulations (*Slc1a3* (red) and *Atp1b2* (green)), in the following astrocyte-enriched hippocampal layers: stratum oriens (SO), stratum radiatum (SR), stratum lacunosum moleculare (SLM), molecular layer (ML), and hilus. Scale bar = 100 µm. **(h)** Pie charts showing the proportions of the two subpopulations of astrocytes (A^+^ and A^+^/G^+^) in each hippocampal layer, in young and old mice, by RNAscope. Two-way ANOVA Sidak’s multiple comparisons test was performed in (d) and (h).). * p < 0.05, ** p < 0.01.

To confirm astrocyte diversity at the mRNA level, we performed high-resolution analysis of ATP1B2 and GLAST expression by *in situ* hybridization (RNAscope), using fluorescent probes for *Atp1b2* and *Slc1a3* (coding for GLAST) (Figure 1e-f). We quantified mRNA at the single-cell level in the whole hippocampus of young and old mice, excluding the densely packed neuronal layers. Again, the majority of *Atp1b2^+^*astrocytes expressed both astrocyte marker genes regardless of age (average±SEM, *Slc1a3^+^/Atp1b2^+^* cells in Y: 72.6±6.8% and O: 60.7±6.8%, respectively). About one-third of the *Atp1b2^+^* cells expressed only *Atp1b2^+^* (Y: 27.4±6.8 % and O: 39.3±6.8%) (Figure 1e). We then took a closer look to the different CA1 and DG layers. Cells that co-expressed both marker genes were the most abundant astrocyte type in all subregions at all ages, although a trend towards a decrease in the fraction of *Slc1a3^+^/Atp1b2^+^* double-positive cells and an increase in *Atp1b2^+^* single-positive cells was observed with age. Differences were significant in the hilus and SO layers (Figure 1g-h). To explore whether changes could reflect a reduction in *Slc1a3* expression in the aged astrocytes, we quantified the number of puncta per cell (dots representing mRNA transcripts) as a proxy for the level of single-cell gene expression. There was an overall decrease in the number of *Slc1a3^+^* puncta (and also *Atp1b2^+^* puncta) per *Atp1b2*-expressing cell during aging, while expression of control housekeeping genes like *Ubc* (coding for Ubiquitin) did not change with age (Supplementary Figure 1). We further confirmed the significant reduction in the number of puncta among the 25 top *Atp1b2*-expressing cells. The age-related decline seemed specific to the hippocampal layers, as no reduction was detected in the cortex (Supplementary Figure 1).

Hence, on the basis of ATP1B2 and GLAST mRNA and protein expression, our data confirm the existence of two subpopulations of astrocytes in the hippocampus of young and aged animals. The two subpopulations are distributed throughout all hippocampal layers, most of them being ATP1B2 and GLAST double-positive astrocytes. A moderate increase in *Atp1b2^+^* single-positive cells is observed in certain hippocampal layers in old animals.

### FACS-isolation and bulk RNA sequencing reveals transcriptional heterogeneity among hippocampal astrocyte subpopulations

We next took advantage of the cell surface expression of ATP1B2 and GLAST proteins to isolate the identified astrocyte subsets. We employed anti-astrocyte cell surface antigen ACSA-2 and ACSA-1 antibodies, that recognize ATP1B2 and GLAST epitopes respectively, conjugated to the fluorophores fluorescein isothiocyanate (FITC) and phycoerythrin (PE) (Batiuk et al., 2020; Borggrewe et al., 2021; Kantzer et al., 2021) (Figure 2a and Supplementary Figure 2). Hippocampal astrocyte diversity was confirmed by FACS, with very similar A^+^ and A^+^/G^+^ cell proportions to those detected by immunohistochemistry and RNAscope. The majority of astrocytes from both young and old hippocampus were A^+^/G^+^ (average±SEM, Y: 84.9±4.7% and O: 76.6±8.0%) compared to A^+^ astrocytes (Y: 15.1±4.7% and O: 23.4±8.0%) (Figure 2b-c). Instead, A^+^ and A^+^/G^+^ proportions in forebrain tissue were close to 50% (Y-A^+^: 49.9±1.2 and Y-A^+^/G^+^: 50.1%±1.2%) (Figure 2b), in line with published data (Borggrewe et al., 2021). Differences between hippocampal and forebrain astrocyte proportions were significant, pointing to an enrichment in A^+^/G^+^ cells in the hippocampus compared to other brain areas.

**Figure 2.**
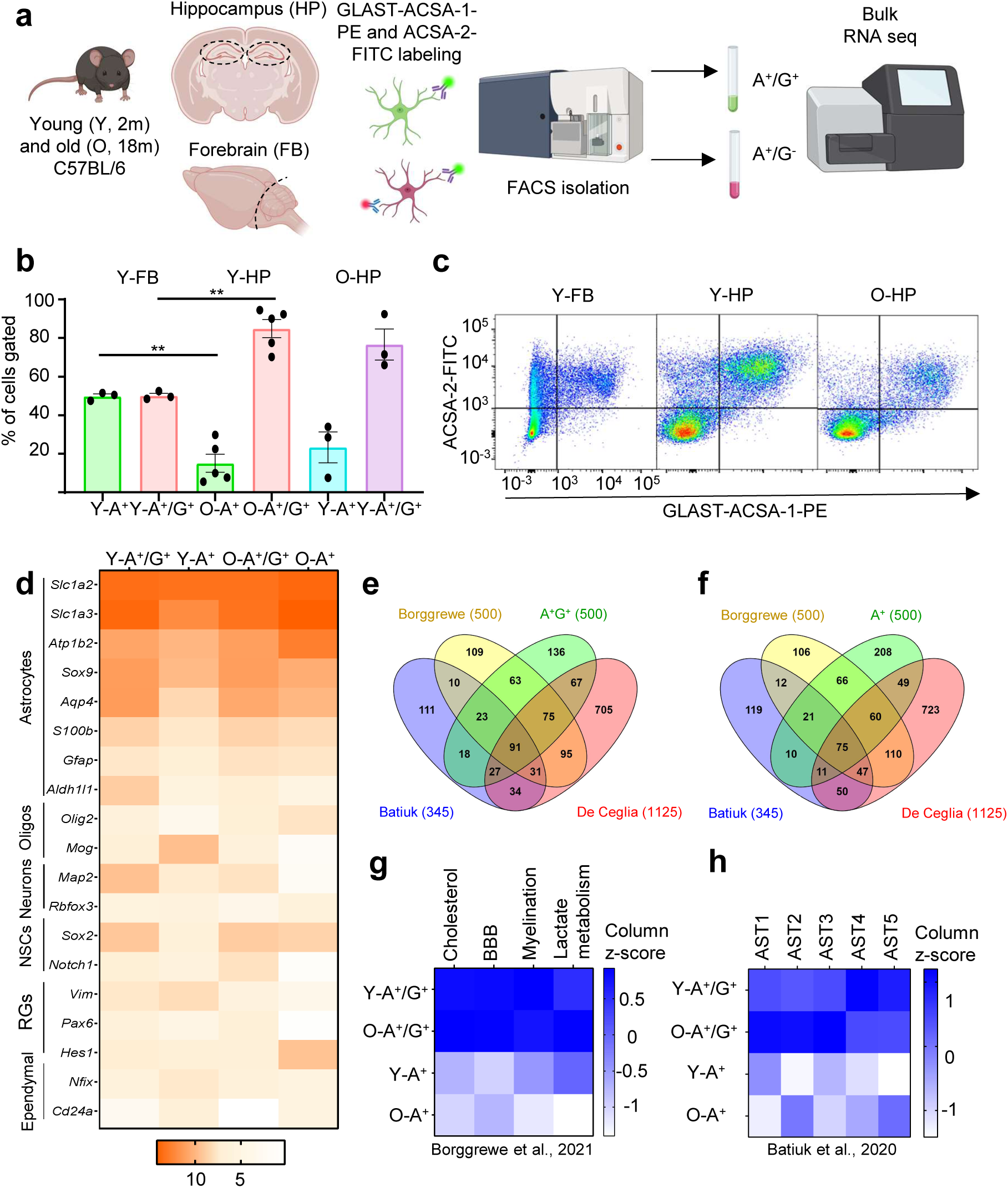
ACSA-1 (GLAST) and ACSA-2 (ATP1B2) surface expression and bulk RNAseq identify two hippocampal astrocyte subpopulations that are preserved during aging. **(a)** Diagram of the procedure followed for the isolation of astrocytes. Hippocampi (and forebrain as a control) were dissected from C57BL/6JRcc mice at 2 and 18 months of age. Cell suspensions were obtained by enzymatic digestion and mechanic dissociation. Astrocytes were labeled with GLAST-ACSA-1-PE and ACSA-2-FITC antibodies, sorted by FACS and RNA was sequenced using Illumina. **(b)** Quantification of astrocyte subpopulations in hippocampus of young (Y, 2m) and old wild-type mice (O, 18m) and forebrain of young mice by FACS. Frequency of single- and double-positive astrocytes (A^+^ and A^+^/G^+^) in the two brain regions are shown as percentages (n≥3). Bars indicate mean ± SEM. Unpaired t-test was performed. * p < 0.05, ** p < 0.01. **(c)** Representative FACS dot plots of ACSA-2 and GLAST-ACSA-1 expression in young and old hippocampus and in young forebrain. For complete gating strategies, see Supplementary Figure 2. **(d)** Gene expression heatmap of the different central nervous system cell type markers. Orange, high expression and white, no expression. **(e-f)** Overlap of top 500 expressed genes in A^+^/G^+^ and A^+^ population, respectively, with published mouse astrocyte gene sets (Batiuk et al., 2020; Borggrewe et al., 2020; De Ceglia et al., 2023) visualized in a Venn diagram. **(g)** Mean expression of genes involved in lactate metabolism, myelination, blood brain barrier, and cholesterol synthesis of published dataset (Borggrewe et al., 2020), illustrated as z-scores per group (n≥3). For lists of genes see Supplementary Table 4. **(h)** Mean expression of genes associated with astrocyte subtypes of published dataset (Batiuk et al., 2020) illustrated as z-score per group (n≥3). For list of genes see Supplementary Table 5. Representation shown in (a) was created with BioRender.com.

We next used FACS-sorting to isolate the hippocampal astrocyte subpopulations from young and old wild-type mice (Figure 2a) and performed bulk RNAseq analysis of the sorted cells. Expression of *bona fide* astrocyte marker genes (*Slc1a2*, *Slc1a3, Atp1b2*, Sox9, *Aqp4*) was high in both A^+^ and A^+^/G^+^ cells, while expression of oligodendrocyte (*Olig2, Mog*), neuron (*Map2, Rbfox3*), neural stem cell (NSC; *Sox2, Notch1*), radial glia cell (RG; *Vim, Pax6, Hes1*) and ependymal cell (*Nfix, Cd24a*) genes was low, as shown in Figure 2d heatmap. Expression of microglia (*Cx3cr1*, *Cd11b*, *Iba1*), endothelial cell (*Pecam1*, *Cd34*, *Flt1*) and additional NSC/progenitor genes (*Nes*) was not detected. These results indicate that the isolated cells were highly enriched in astrocytes and not significantly contaminated by these other cell types.

To further confirm the identity of the isolated cells, we next compared the top 500 expressed genes of A^+^ and A^+^/G^+^ populations (Supplementary Table 2 and 3) with other published astrocyte signatures (de Ceglia et al., 2023; Borggrewe et al., 2021; Batiuk et al., 2020). 296 of the top expressed genes (59%) were common for A^+^ and A^+^/G^+^ populations and high overlap was found with other datasets (Figure 2e-f). We also analysed the mean expression of genes involved in typical astrocyte functions such as: cholesterol metabolism, blood brain barrier (BBB), myelination and lactate metabolism (Borggrewe et al. 2021) (Figure 2g; Supplementary Table 4). Genes involved in the four categories were more expressed in A^+^/G^+^ than in A^+^ astrocytes regardless of age, suggesting that A^+^/G^+^ correspond to more mature astrocytes. In addition, we interrogated our bulk RNAseq dataset for the Batiuk et al. classification of five astrocyte subtypes, previously identified by sc-RNAseq in the adult mouse hippocampus and cortex (AST1-5; Supplementary Table 5) (Batiuk et al., 2020). Genes of the mature hippocampal subtypes “AST1-3” were found mostly expressed in O-A^+^/G^+^ astrocytes, whereas A^+^ astrocytes expressed intermediate levels of “AST1-3” signature genes, and low levels of “AST2” cortical signature genes. Progenitor cell subtype “AST4” and intermediate progenitor astrocyte subtype “AST5” genes were found slightly more expressed in Y-A^+^/G^+^ compared to O-A^+^/G^+^ astrocytes.

We next focused on the differentially expressed genes (DEGs) that distinguished the A^+^ and A^+^/G^+^ astrocyte transcriptome. Bulk RNAseq analysis showed 548 DEGs (p adj<0.05) when comparing the two astrocyte subpopulations in young animals (Figure 3a and Supplementary Table 6). 469 DEGs were overexpressed in Y-A^+^/G^+^ astrocytes and 79 in Y-A^+^ astrocytes, as shown in the Volcano plot (Figure 3b). Functional enrichment of the differentially up-regulated genes in Y-A^+^/G^+^ and Y-A^+^ astrocytes is shown in Figure 3c-d (GO Biological Process). Y-A^+^/G^+^ astrocyte transcriptome was significantly enriched in genes related to protein glycosylation and assembly of late multivesicular endosomes, two processes involved in membrane protein dynamics and secretion (Figure 3e; Supplementary Table 7). Glycosylation at the ER and Golgi modifies proteins directed to the cell membrane or secreted to the extracellular space. Protein glycosylation up-regulated genes in Y-A^+^/G^+^ astrocytes encoded, among other enzymes, mannosyltransferases (*Alg1, Alg3),* glucosyltransferases (*Alg6*), N-glycan branching enzymes (*Mgat2* and *Mgat4* isoforms) and sialyltransferases that add sialic acid to N-glycans (*St8sia5*). To confirm that overexpression of the enzymes indeed resulted in elevated glycosylation of cell surface proteins, we took advantage of the fluorescein-conjugated leukocyte-phytohemagglutinin (L-PHA), a plant lectin highly specific for branched N-glycans (Yale et al. 2018). Lectin flow cytometry with L-PHA combined with ACSA-2 and ACSA-1 detected a significant raise in L-PHA binding in Y-A^+^/G^+^ astrocytes compared to Y-A^+^ astrocytes, reflecting an increased level of N-glycosylation in the ATP1B2 and GLAST double-positive astrocyte population (Figure 3f).

**Figure 3.**
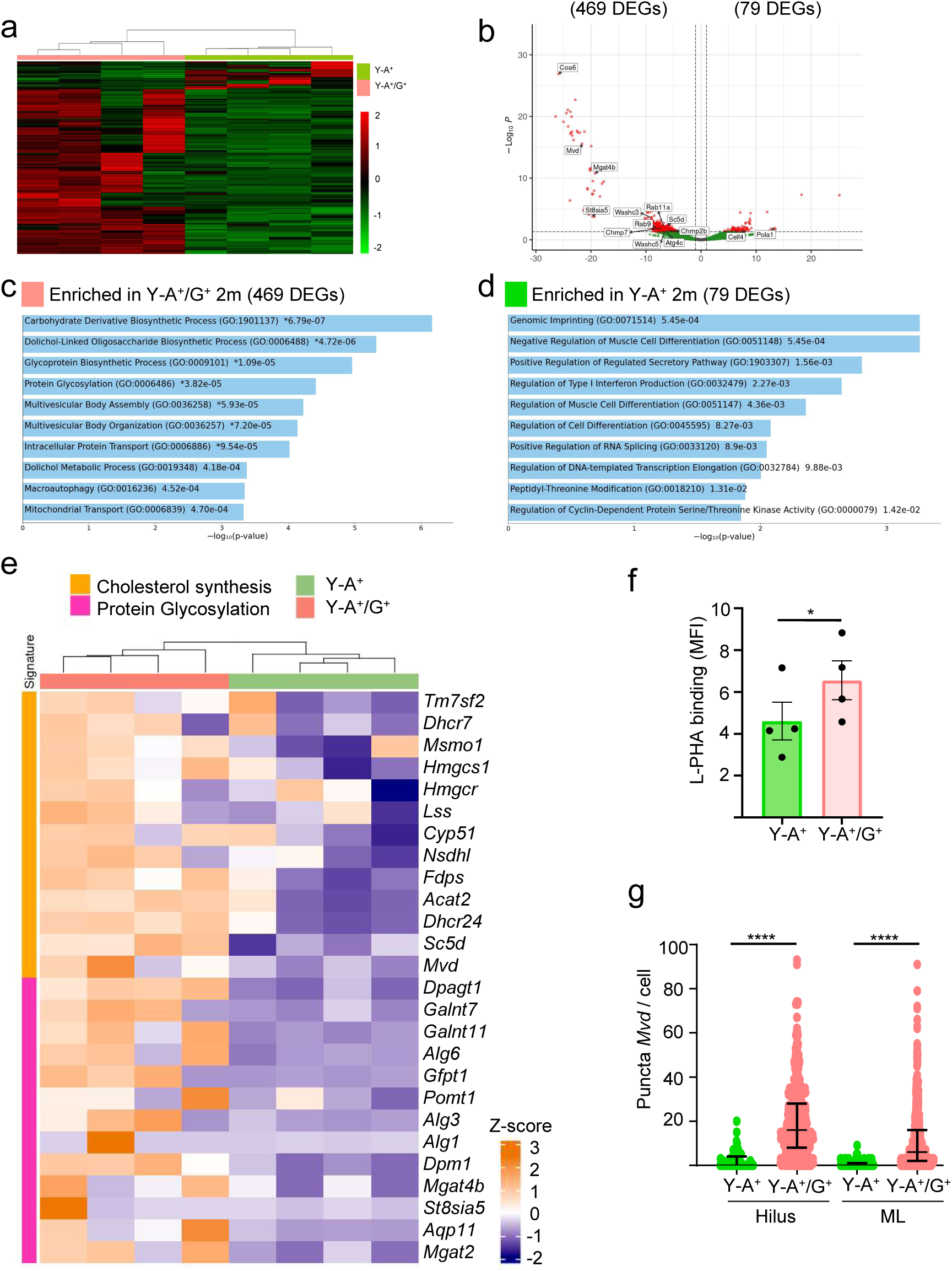
A^+^ and A^+^/G^+^ astrocytes show differential gene expression in the young hippocampus. **(a)** Unsupervised clustering of all genes differentially expressed in each astrocyte subpopulation in young mice (2m), illustrated as row z-scores of normalized counts. **(b)** Volcano plot of DEGs between Y-A^+^ and Y-A^+^/G^+^ astrocytes. Some representative genes are labeled. **(c)** GO analysis of enriched genes in Y-A^+^/G^+^ astrocytes. **(d)** GO analysis of enriched genes in Y-A^+^ astrocytes. **(e)** Clustering heatmap of RNAseq data showing VST normalized counts of Protein Glycosylation (GO:0006486) GO term and Cholesterol Synthesis (PMID: 29279367) genes in Y-A^+^ and Y-A^+^/G^+^ astrocytes. Values were compared with the DESeq2 package. **(f)** L-PHA mean Fluorescence intensity in Y-A^+^ and Y-A^+^/G^+^ astrocytes from young mice (5m) measured by flow cytometry. Data is shown as mean ± SEM. Paired t-test was performed. **(g)** Quantification of *Mvd* expression in Y-A^+^ and Y-A^+^/G^+^ astrocytes by RNAscope in hilus and molecular layer (ML) of hippocampus. Data is shown as median ± interquartile range. Mann-Whitney test was performed. * p < 0.05, ** p < 0.01, *** p < 0.001 and **** p < 0.0001.

Multivesicular body assembly and late endosome to vacuole transport categories enriched in Y-A^+^/G^+^ astrocytes included charged multivesicular body protein genes *Chmp7* and *Chmp2b* and several vacuolar protein sorting (VPS) genes. Other genes coding for WASH complex proteins, involved in regulating endosomal protein sorting, such as *Washc3* and *Washc5* were also overexpressed in Y-A^+^/G^+^ astrocytes (Figure 3b). Expression of Rab family of small GTPases genes (*Rab9, Rab11a*) controlling intracellular vesicular transport and receptor recycling back to the plasma membrane was increased as well. Macroautophagy/Lysosomal-related genes implicated in the turnover of cytoplasmic components, such as *Atg4c*, were upregulated in Y-A^+^/G^+^ astrocytes. Interestingly, Y-A^+^/G^+^ astrocytes were also enriched in mitochondrial transport genes and greatly overexpressed cytochrome c oxidase mitochondrial respiratory complex IV assembly factor gene *Coa6* (Figure 3b). *Coa6* was the coding gene with highest normalized fold change in expression in Y-A^+^/G^+^ compared to Y-A^+^, pointing to differences in mitochondrial activity between the two astrocyte subpopulations.

In addition, Y-A^+^/G^+^ astrocytes showed increased expression of a subset of cholesterol-related genes compared to Y-A^+^ astrocytes (Figure 3e), including *Sc5d* and *Mvd* that encode sterol-C5-desaturase and mevalonate diphosphatase decarboxylase, respectively, two crucial enzymes in cholesterol biosynthesis. We quantified *Mvd* expression per cell by RNAscope technique in two hippocampal layers containing abundant astrocytes, the hilus and molecular layer (ML). In both layers, the number of *Mvd* puncta per cell was higher in *Slc1a3^+^/Atp1b2^+^*cells than in *Atp1b2^+^* cells (Figure 3g). These results confirm the increase in *Mvd* mRNA in ATP1B2 and GLAST double-positive astrocytes compared to ATP1B2 single-positive astrocytes.

Regarding the transcriptomic signature of ATP1B2 single-positive astrocytes, as mentioned above only 79 genes were overexpressed in Y-A^+^ cells compared to Y-A^+^/G^+^ (Figure 3b and Supplementary Table 6). Functional enrichment (GO Biological Process) of the Y-A^+^ upregulated genes highlighted functions such as negative regulation of muscle cell differentiation and imprinting (*Igf2*), mRNA splicing (e.g. *Scnm1, Celf4*) and regulation of microtubule assembly in mitosis and spindle checkpoint signalling (e.g. *Drg1, Hspa1a*). The DNA polymerase gene *Pola1* implicated in DNA replication and cell division, and the cyclin-dependent kinase regulators *Ccnt1* and *Cdk5rap1* were also overexpressed in Y-A^+^ cells.

In summary, GLAST allows to distinguish two ATP1B2-expressing astrocyte subpopulations in the adult hippocampus, A^+^ and A^+^/G^+^, that are transcriptionally distinct. Their gene expression profile is shared in part with astrocyte subtypes identified in previous studies. Functional enrichment analysis of the transcriptomic signatures and validation studies suggest that ATP1B2/GLAST double-positive astrocytes correspond to glial cells with increased protein glycosylation and more mature functions.

### Age-related transcriptional changes differ between hippocampal astrocyte subpopulations

We next analysed the transcriptomic changes of the FACS sorted hippocampal astrocytes in old compared to young animals. Interestingly, while the A^+^ astrocyte transcriptome profoundly changed during aging (1935 DEGs in O-A^+^ *versus* Y-A^+^, Figure 4a-b; Supplementary Table 8), the A^+^/G^+^ astrocyte transcriptome remained quite stable (only 67 DEGs in O-A^+^/G^+^ *versus* Y-A^+^/G^+^, Figure 4c-d; Supplementary Table 9). For both astrocyte populations, most DEGs corresponded to down-regulated genes (70% in O-A^+^ and 88% in O-A^+^/G^+^). Common down-regulated targets during aging included the ubiquitination and proteasomal degradation-related gene *Skp2*, the metalloproteinase *Adamts20* involved in tissue-remodelling and cilium-related genes. One shared gene (C*tu2*) involved in translation fidelity was up-regulated both in O-A^+^/G^+^ and O-A^+^ cells compared to their young counterparts (Figure 4e).

**Figure 4.**
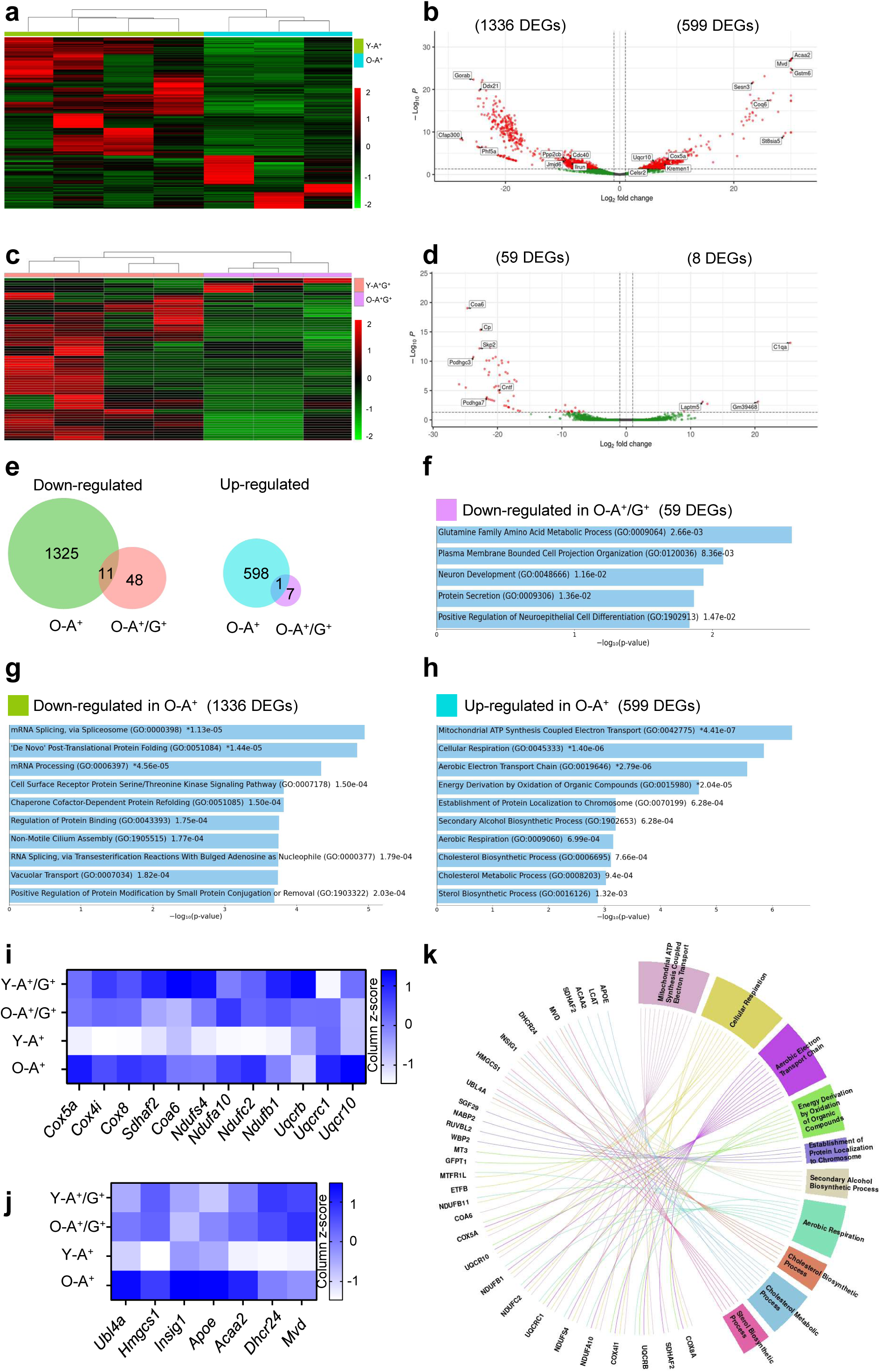
Differential transcriptional changes in A^+^ and A^+^/G^+^ hippocampal astrocytes during aging. **(a)** Unsupervised clustering of all genes differentially expressed in A^+^ astrocytes with age, illustrated as row z-scores of normalized counts. **(b)** Volcano plot of DEGs between Y-A^+^ and O-A^+^ astrocytes. Some representative genes are labeled. **(c)** Unsupervised clustering of all genes differentially expressed in A^+^/G^+^ astrocytes with age, illustrated as row z-scores of normalized counts. **(d)** Volcano plot of DEGs between Y-A^+^/G^+^ and O-A^+^/G^+^ astrocytes. Some representative genes are labeled. **(e)** Venn diagrams showing the number of significantly (p < 0.05) down-regulated and up-regulated genes in the two astrocyte subpopulations of old mice, determined by DESeq2 analysis. Comparisons between O-A^+^ and O-A^+^/G^+^ astrocytes show little overlap in the age-related transcriptional changes. **(f)** GO analysis of age-related down-regulated genes in A^+^/G^+^ astrocytes. **(g)** GO analysis of age-related down-regulated genes in A^+^ astrocytes. **(h)** GO analysis of age-related up-regulated genes in A^+^ astrocytes. **(i)** Heatmap comparing the mean expression of mitochondrial and cholesterol metabolism genes in the astrocyte subpopulations with age. **(j)** Circular plot depicting GO annotations of up-regulated genes in A^+^ astrocytes isolated from the old hippocampus compared to the young hippocampus.

Biological functions enriched in the down-regulated A^+^/G^+^ gene list during aging were related to plasma membrane cell projection organization and neuron development, amino acid metabolism and protein secretion (Figure 4f). Further inspection of the list uncovered several members of the cell adhesion γ-protocadherin (*Pcdhg*) gene cluster, ciliary neurotrophic factor (*Cntf*) and ceruloplasmin (*Cp*) (Figure 4d, Supplementary Table 9). Functional enrichment was not followed-up in up-regulated O-A^+^/G^+^ targets due to the limited number of DEGs (only 8, including the complement gene *C1qa*, lncRNA *Gm39468* and the lysosomal-associated protein coding gene *Laptm5*).

Regarding the differentially expressed genes in A^+^ astrocytes during aging, functional enrichment of the down-regulated targets identified processes such as: protein folding (several DnaJ heat shock protein (HSP) genes and representative members of the stress-inducible HSP70 family genes), protein modification, vacuolar transport, mRNA Processing / Splicing (including *Celf4* and *Scnm1* overexpressed in Y-A^+^ compared to Y-A^+^/G^+^) and cell surface receptor Serine/Threonine kinase signalling (Figure 4g). Other repressed genes were linked to protein trafficking from the ER to the Golgi (*Gorab*), genome stability (RNA helicase *Ddx21)*, cell cycle progression/proliferation modulators (e.g. *Ppp2cb*, *Cdc40*, *Jumjd6*) and cytokine production (*Ilrun*) (Figure 4b).

As for the up-regulated genes in A^+^ astrocytes during aging, many were involved in mitochondrial electron transport-mediated ATP synthesis (e.g. *Cox5a*, *Coa6*, *Uqcr10*) and cholesterol metabolism (e.g. *Acaa2, Mvd*) (Figure 4b and 4h-k). Other biological functions uncovered in O-A^+^ astrocytes include chromatin-related processes (up-regulation of transcriptional co-activators, SAGA-complex and DNA repair related proteins) and Wnt signalling pathway genes (e.g. *Kremen1, Celsr2*) (Figure 4b and 4h).

Thus, hippocampal A^+^ astrocytes in old mice down-regulate genes linked to cellular processes that distinguish them from A^+^/G^+^ astrocytes in young mice (i.e. splicing) and rewire their metabolism, increasing the expression of genes involved in mitochondrial ATP synthesis and cholesterol biosynthesis. In addition, some of the up-regulated targets in the single-positive A^+^ astrocytes during aging were protein glycosylation genes overexpressed in Y-A^+^/G^+^ (*St8sia5*, Figure 4b), altogether pointing to the acquisition of a more mature state.

### ATP1B2 and GLAST double-positive astrocytes repress γ-protocadherin genes during aging and fail to support excitatory synapse formation

Since ATP1B2 and GLAST double-positive astrocytes predominate in the hippocampus (80% of the astrocytes, Figure 1 and Figure 2b), we next explored the O-A^+^/G^+^ RNAseq dataset for DEGs that could account for the age-related decline in astrocyte physiological functions. We focused on the loss of synaptogenic function previously suggested for senescent astrocytes (Boisvert et al., 2018; Matias et al., 2021). Astrocytes promote excitatory synapse formation by secreting synaptogenic proteins and also through physical cell-to-cell contacts with neurons. Protocadherin proteins localize to perisynaptic astrocyte processes and have been reported to facilitate synaptogenesis (Garret and Weiner, 2009). As shown in Figure 5a, the RNAseq analysis identified four γ-protocadherin coding genes significantly down-regulated during aging in O-A^+^/G^+^ astrocytes (but not in O-A^+^ astrocytes, Supplementary Figure 3). Inspection of available transcriptomic datasets (Zhang et al., 2024; Zhang et al., 2016) indicated that three of these *Pcdhg* genes are expressed at equal levels in neurons and astrocytes, while expression of *Pcdhgc3* encoding γ-C3 protocadherin is highly enriched in astrocytes (Figure 5b)(Lee et al., 2025).

**Figure 5.**
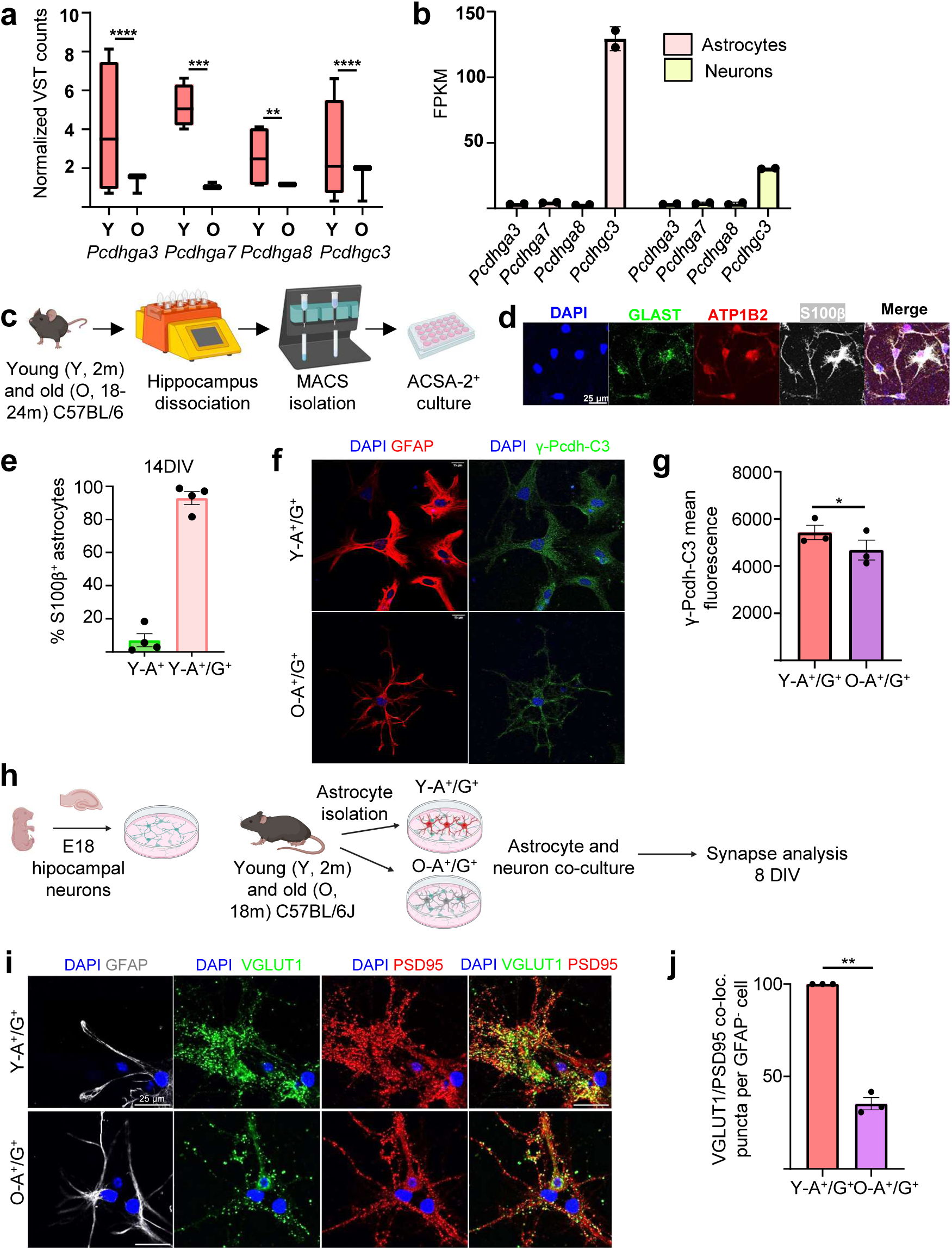
A^+^/G^+^ astrocytes transcriptome changes during aging and its effect in neuron synapse promotion. **(a)** Box and whiskers plot of RNAseq data showing VST normalized counts of *Pcdhga3*, *Pcdhga7, Pcdhga8* and *Pcdhgc3* genes in Y-A^+^/G^+^ and O-A^+^/G^+^ astrocytes. Values were compared with the DESeq2 package. **(b)** Bar plots of RNAseq data showing FPKM expression of four protocadherin-related genes (*Pcdhga3*, *Pcdhga7*, *Pcdhga8*, and *Pcdhgc3*) in cortical astrocytes and neurons. FPKM, fragments per kilobase of exon per million mapped fragments. **(c)** Schematic diagram showing the experimental strategy for isolation of astrocytes (ACSA-2^+^) from the hippocampus of young (Y, 2m) and old (O, 18-24m) wild-type mice by MACS and subsequent culture *in vitro*. **(d)** Representative immunostaining of GLAST (green), ATP1B2 (red) and S100β (white) in ACSA-2^+^ astrocytes from young (2m) adult mice, cultured during 14 DIV. Scale bar = 25 µm. **(e)** Quantification of A^+^/G^+^ and A^+^ astrocytes in young hippocampal astrocyte primary cultures at 14DIV. Percentage of S100β^+^/ATP1B2^+^/GLAST^+^ (A^+^/G^+)^ and S100β^+^/ATP1B2^+^/GLAST^-^ (A^+^) astrocytes is shown as mean ± SEM. **(f)** Representative immunofluorescence images of GFAP (red) and γ-Pcdh-C3 (green) in Y-A^+^/G^+^ (2m) and O-A^+^/G^+^ (24m) astrocytes at 14 DIV. Scale bar = 15 µm. **(g)** γ-Pcdh-C3 mean fluorescence quantification in Y-A^+^/G^+^ and O-A^+^/G^+^ astrocytes. Data is shown as mean ± SEM. Paired t-test was performed. **(h)** Schematic diagram showing the experimental procedure followed to obtained neurons (E18) and astrocytes (2 and 18m) for the heterochronic co-culture. **(i)** Immunostaining of GFAP (white), VGLUT1 (green) and PSD95 (red) in neurons co-cultured with young or old astrocytes at 8 DIV. Scale bar = 25 µm. **(j)** Quantification of excitatory pre-(VGLUT1) and postsynaptic (PSD95) vesicles colocalization in hippocampal neurons co-cultured with hippocampal young (2m) and old (18m) astrocytes. Three independent experiments were performed (n=3). Data are presented as mean ± SEM. One-sample t-test was performed. * p < 0.05, ** p < 0.01, *** p < 0.001 and **** p < 0.0001. Data in (b) were obtained from the database produced by the Barres lab (Zhang et al. 2014 available at https://brainrnaseq.org/). Representations shown in (c) and (h) were created with BioRender.com.

To confirm γ-C3 protein expression in hippocampal A^+^/G^+^ astrocytes and γ-C3 reduction during aging, astrocytes were isolated from the hippocampus of young and old mice by magnetic-activated cell sorting (MACS) using ACSA-2. Astrocytes were cultured *in vitro* in defined medium without serum, a procedure that maintains the *in vivo* astrocyte properties while avoiding the induction of a reactive astrocyte signature (Figure 5c) (Caldwell et al., 2022). The vast majority of the S100β^+^ astrocytes expressed both ATP1B2 and GLAST proteins (92.9±3.9% A^+^/G^+^) (Figure 5d-e). Thus, we considered these primary cultures as highly-enriched in hippocampal A^+^/G^+^ astrocytes. Quantitative immunofluorescence analysis demonstrated γ-C3 protein expression in Y-A^+^/G^+^ cultures and decreased γ-C3 signal intensity in O-A^+^/G^+^ cultures (Figure 5f-g). γ-C3 signal was poorly detected in MACS-isolated primary hippocampal neurons (MAP2^+^cells, Supplementary Figure 3). However, pan-γ-protocadherin signal was detected in both neurons and astrocytes (Supplementary Figure 3). This confirmed the presence γ-protocadherins in the astrocytic and neuronal compartments, the predominant γ-C3 expression in young astrocytes and its reduction in old astrocytes.

Astrocyte-neuron γ-protocadherin contacts are necessary for proper neuronal synapse formation and stabilization (Garret and Weiner, 2009; LaMassa et al., 2021), so we reasoned that γ-protocadherin reduction in aged astrocytes could affect synaptogenesis. We designed a functional heterochronic *in vitro* assay to compare the synaptogenic capacity of young and old hippocampal astrocytes (Figure 5h). The co-culture system allowed for direct neuron-astrocyte contacts between young (2m) or old (18m) primary hippocampal astrocytes and young hippocampal neurons (E18). We assessed excitatory synapse formation in the heterochronic system through immunofluorescence and co-localization image analysis of the pre-synaptic vesicle marker VGLUT1 (Vesicular Glutamate Transporter 1) and the post-synaptic marker PSD95 (Post-Synaptic Density protein 95) (Ippolito and Eroglu, 2010). Excitatory synapses were reduced by 50% in old astrocyte co-cultures compared to young ones (Figure 5i-j). γ-protocadherin puncta localized to many sites adjacent to pre-synaptic VGLUT1 puncta, in line with published studies (Garret and Weiner, 2009; LaMassa et al., 2021; not shown).

Thus, O-A^+^/G^+^ hippocampal astrocytes fail to support excitatory synaptogenesis in hippocampal neuronal cultures, a dysfunction that may be partly explained by the age-related decrease in cell adhesion molecules of the γ-protocadherin protein family, including γ-C3.c

## Discussion

Astrocyte diversity in the mammalian brain is an active research area that is challenging the long-held conception of astrocytes as a homogeneous cell population (Boisvert et al., 2018; Morel et al., 2017: Bayraktar et al., 2020; Batiuk et al. 2020; Borggrewe et al. 2021; Kantzer et al. 2021; Karpf et al., 2022, De Ceglia et al. 2023; Viana et al. 2023; Zhou et al., 2025). Astrocytes perform many essential functions in brain homeostasis, but it is currently unclear whether different astrocytes from the brain’s collection perform similarly all tasks (Hennes et al., 2025). As a complementary approach for studying astrocyte diversity in the hippocampus throughout aging, in this study we have taken advantage of the cell surface proteins GLAST and ATP1B2, since combined and single expression of these markers has previously allowed for the prospective isolation and transcriptomic analysis of astrocyte subpopulations from other brain regions.

In line with findings in forebrain, spinal cord, hindbrain and cerebellum (Borggrewe et al. 2021; Kantzer et al. 2021), here we demonstrate that GLAST and ATP1B2 expression discriminates between two astrocyte subsets in the hippocampus: the double-positive ATP1B2^+^/GLAST^+^ and the single-positive ATP1B2^+^ astrocytes, with non-detectable GLAST levels. While previous FACS analysis indicated that ATP1B2^+^/GLAST^+^ astrocyte proportions ranged from 40-60% in forebrain, to 20% in hindbrain or 1% in spinal cord (Borggrewe et al. 2021), we found that the frequency of the double-positive cells in the hippocampus was strikingly high (>80%) across all layers, in young and old mice, highlighting the regional heterogeneity of GLAST expression. A recent study explored to a limited extent GLAST and ATP1B2 expression in the hippocampal dentate gyrus of young mice (Karpf et al. 2022). ATP1B2 was found to be highly present in hilus astrocytes compared to granular zone or molecular layer astrocytes. In our study, ATP1B2^+^ enrichment was only detected in hilar astrocytes when analysed at the mRNA level in 18-month old mice. Collectively, our data employing three independent techniques (immunofluorescence, RNAscope and FACS) reveal that ATP1B2^+^/GLAST^+^ astrocytes are plentiful across the hippocampus, sharing territories with a less abundant subset of astroglial cells classified as ATP1B2^+^ astrocytes.

We found substantial differences in the gene expression profiles of ATP1B2^+^/GLAST^+^ and ATP1B2^+^ hippocampal astrocytes by bulk RNAseq. Two studies (Borggrewe et al. 2021; Kantzer et al. 2021) previously compared the transcriptome of single- and double-positive astrocytes in other regions. Borggrewe et al. failed to detect DEGs when comparing GLAST^+^ versus GLAST^-^ forebrain astrocytes. In hindbrain, the GLAST^-^ astrocyte transcriptome was enriched in glial cell differentiation and myelination genes. Overall mean expression of genes involved in cholesterol metabolism and blood brain barrier was higher in GLAST^+^ compared to GLAST^-^ astrocytes, while lactate metabolism genes were more expressed in GLAST^-^ astrocytes (Borggrewe et al. 2021). Moreover, the “AST5” astrocyte intermediate progenitor signature (Batiuk et al., 2020) was more associated to hindbrain and forebrain GLAST^-^ astrocytes. Expanding these observations, the transcriptomic signature of hippocampal GLAST^-^ astrocytes (ATP1B2^+^ only) was reminiscent of a more immature cell, although it did not clearly overlap with the “AST5” gene expression profile. As discussed below, functional enrichment analysis of the transcriptomic data and validation assays suggest that ATP1B2^+^/GLAST^+^ double-positive hippocampal astrocytes represent a more mature cellular state compared to ATP1B2^+^ astrocytes.

Mature astrocytes are highly secretory cells with a finely regulated exocytotic system (Verkhratsky et al., 2016). They communicate with neighbouring cells, including neurons, through the release of neuroactive compounds, neurotransmitters, signalling molecules, and through cell adhesion proteins and cell-to-cell contacts as well. Interestingly, hippocampal ATP1B2^+^/GLAST^+^ double-positive astrocytes expressed higher levels of protein glycosylation genes, particularly N-linked glycosylation, and were highly stained for L-PHA, a specific lectin for branched N-glycans. The addition of sugar molecules is a very common modification for secreted and membrane-bound proteins. Thus, the increased glycosylation detected in ATP1B2^+^/GLAST^+^ double-positive astrocytes reinforces their view as more mature cells, actively involved in interactions with other brain cell types. Compared to ATP1B2^+^ single-positive astrocytes, the double-positive ones also overexpressed genes involved in vesicular transport, cholesterol synthesis and metabolism, a set of functions that characterize mature astrocytes (Pfrieger et al., 2011; Verkhratsky et al., 2016).

Conversely, ATP1B2^+^ astrocytes differentially overexpressed a limited number of genes when compared to ATP1B2^+^/GLAST^+^ astrocytes. These genes were involved in DNA replication, RNA processing and cell division. Thus, the transcriptomic data tentatively suggest that, compared to double-positive astrocytes, ATP1B2^+^ astrocytes are less mature and perhaps more susceptible to undergo cell cycle reactivation, a hypothesis that remains to be tested.

We also found remarkable differences in the transcriptomic response of the two hippocampal astrocyte subpopulations during aging. The ATP1B2^+^ transcriptome was very plastic and vast changes in gene expression were detected in 18-months old compared to 2-months old animals. This cell response was defined by the up-regulation of a battery of genes involved in mitochondrial respiration and metabolic rewiring. The shift highlights the capacity of the ATP1B2^+^ cells to respond to the aged brain environment. In contrast, the ATP1B2^+^/GLAST^+^ transcriptomic signature was very robust. Thus, the RNAseq data indicate that double-positive astrocytes withstand aging without undergoing dramatic transcriptomic changes, thereby maintaining stable levels of gene expression. Currently, we do not know whether different epigenetic states underlie the plastic *versus* stable gene expression pattern identified in the two astrocyte subpopulations during aging. Although ATP1B2^+^/GLAST^+^ astrocytes undergo little transcriptomic changes in 18-months old mice compared to younger animals, a subset of DEGs (protocadherin genes, complement cascade genes) point to an impaired function of the old hippocampal double-positive astrocytes in synapse formation and maintenance.

During normal aging, the number of synapses decreases from 15 to 50% depending on the species and brain area, compromising neuronal connectivity (Pannese, 2011). In certain hippocampal subregions, aged rodents with learning deficits show reduced synaptic markers (Smith et al., 2000). Moreover, loss of synaptogenic function has been previously suggested for old and senescent astrocytes (Boisvert et al., 2018; Matias et al., 2021). Astrocytes from a variety of brain regions other than hippocampus (visual cortex, motor cortex, cerebellum and hypothalamus) showed marked alterations in the expression of several synapse-regulating genes during aging (Boisvert et al., 2018). This finding is in line with the poor supportive role of old hippocampal astrocytes in the heterochronic synaptogenesis assays performed in the current study. While complement cascade components involved in synapse elimination were up-regulated in a coordinated manner in aged astrocytes from all the aforementioned brain areas, the synapse inducing genes *Thbs1* and *Thbs4* showed region-specific changes and were significantly decreased in astrocytes from the aging cerebellum and hypothalamus, respectively (Boisvert et al., 2018; Clarke et al., 2018). We add the γ-protocadherin (*Pcdhg*) gene cluster to this list of synapse regulatory genes altered during aging in astroglial cells.

Protocadherins are expressed by both neurons and astrocytes and have relevant roles in developmental synapse formation, maturation and stabilization (Peek et al., 2017; LaMassa et al., 2021). γ-Pcdh proteins establish adhesive homophilic trans-interactions and promiscuous cis-interactions with other γ-Pcdh proteins through their extracellular domains (Peek et al., 2017). Combinatorial γ-Pcdh expression in neurons orchestrates dendritic arborization and synaptic connectivity in the neocortex, allowing to discriminate between self and non-self (Molumby et al., 2016; Zhu et al., 2024). In astrocytes, γ-Pcdh proteins localize to perisynaptic astrocyte processes and γ-Pcdh cell-cell contacts between astrocytes and neurons facilitate synaptogenesis (Garret and Weiner, 2009). In vivo, hippocampal γ-Pcdh immunopositive synapses are larger and more mature (LaMassa et al., 2021) while restricted mutation of the *Pcdhg* gene cluster in astrocytes delays synapse formation, at least in the spinal cord (Garret and Weiner, 2009). In our study, decreased γ-C3 levels in hippocampal astrocytes from old mice correlates with their loss of excitatory synaptogenic capacity, as shown in heterochronic neuron-astrocyte co-cultures that allow cell-to-cell contacts. Recent data demonstrate γ-C3 expression by virtually all cortical *Slc1a3^+^* (GLAST^+^) astrocytes and uncover a cell-autonomous role of γ-C3 in astrocyte self-recognition during astrocyte morphogenesis (Lee et al., 2025). Interestingly, astrocyte morphology in the cortex and hippocampus of γC3-KO mice show reduced territory size and volume (Lee et al. 2025). Given astrocytes shrink during aging (Popov et al., 2021), it will be interesting to explore in the future whether γ-C3 downregulation in ATP1B2^+^/GLAST^+^ hippocampal astrocytes from old mice influences astrocyte morphology.

The loss of cell adhesion molecules, including protocadherins, may not only affect the interaction of perisynaptic processes between astrocytes and neurons, but may also have other negative consequences on the structural complexity and function of astrocytes in pathological aging (Lee et al. 2025). Neurodegenerative diseases are characterized simultaneously by increased astrocyte hypertrophy (associated to the reactive phenotype) and astrocyte atrophy (Verkhratsky et al., 2014). Since impaired astrocyte function can disrupt neuronal circuitry and drive pathogenesis, brain repair could benefit from therapeutic strategies directed towards the astrocytic compartment, such as transplantation of engineered astrocytes, reprogramming of endogenous astrocytes to restore youthful capabilities or recruiting specific astrocyte subsets to fulfill lost functions (Pihlaja et al., 2008; Giralt et al., 2010, Proschel et al., 2014; Lattke et al., 2022; Chierzi et al., 2023). A key goal for the future is to leverage molecular diversity data to develop tools that allow targeted manipulation of astrocyte subtypes.

In summary, our work identifies two astrocyte subpopulations in the adult mouse hippocampus that are preserved during aging and provides a new resource for understanding the transcriptional differences between ATP1B2^+^ single-positive and ATP1B2^+^/GLAST^+^ double-positive astrocytes. We also highlight the contrasting transcriptomic changes observed during aging in these two astrocyte subpopulations, which point to distinct cellular responses and to a specific loss of synaptogenic support of the more abundant and mature double-positive astrocytes. How these responses influence the progression of age-related diseases remains to be elucidated. A deeper understanding of astrocyte heterogeneity will open new avenues for modulating processes related to aging and brain pathology.

## Supporting information

Supplementary informatiin

## Acknowledgments

We thank technical assistance from Rosa Viana at the IBV Confocal Microscopy Core and Úrsula Estada Gimeno at the Genòmica i Epigenètica Unit (UCIM), Universitat de València. This work was supported by grants PID2019-111225RB-I00 and PID2022-141707NBI00 from Spanish Ministry of Science and Innovation, and CIAICO/2022/74 from Generalitat Valenciana to H.M.

---

**Supplementary Figure 1. (a)** Quantification of the expression of *Atp1b2*, *Slc1a3* and *Ubc* genes by measuring total puncta (mRNA) per cell in the hippocampus of young and old wild-type mice by RNAscope. **(b)** Quantification of *Slc1a3* puncta (mRNA) per cell in astrocyte-enriched hippocampal layers of young and old wild-type mice and in cortex by RNAscope. **(c)** Quantification of *Atp1b2* puncta (mRNA) per cell in astrocyte-enriched hippocampal layers of young and old wild-type mice and in cortex by RNAscope. Data in (b) and (c) correspond to the 25 top *Atp1b2*-expressing cells in 2 m and 18 m wild-type animals (n≥3). Data is shown as median ± interquartile range. Mann-Whitney test was performed is (a). Kruskal-Wallis test multiple comparisons was performed in (b) and (c). * p < 0.05, ** p < 0.01, *** p < 0.001 and **** p < 0.0001.

**Supplementary Figure 2. Gating strategies in FACS procedure. (a)** Dot plot showing side scatter versus forward scatter to select the population avoiding debris. **(b)** Cell doublets were excluded using forward scatter width *vs*. height plot. **(c)** Dead cells were excluded based on DAPI staining. For the gating, only DAPI^-^ (live) cells were selected and analysed afterwards. Percentages represent the fraction of total cells present within the gate.

**Supplementary Figure 3. (a)** Box and whiskers plot of RNAseq data showing VST normalized counts of *Pcdhga3*, *Pcdhga7, Pcdhga8* and *Pcdhgc3* genes in Y-A^+^ and O-A^+^ astrocytes. Values were compared with the DESeq2 package. **(b)** Representative immunofluorescence images of MAP2 (white) and Pan-γ-Pcdh (red) signal in E18 cultured hippocampal neurons at 11 DIV. Scale bar = 25 and 10 µm. **(c)** Representative immunofluorescence images of MAP2 (green) and γ-Pcdh-C3 (red) signal in E18 cultured hippocampal neurons at 11 DIV. Scale bar = 25 and 10 µm. **(d)** Representative immunofluorescence images of GFAP (white) and Pan-γ-Pcdh (red) signal in cultured young (2 m) hippocampal astrocytes at 14 DIV. Scale bar = 25 and 10 µm.

## Notes

### Competing Interest Statement

The authors have declared no competing interest.

