## Supplementary informatiin for "Cell surface markers identify astrocyte subpopulations in the adult hippocampus with a heterogeneous response to aging"

### Supplementary Figure 1

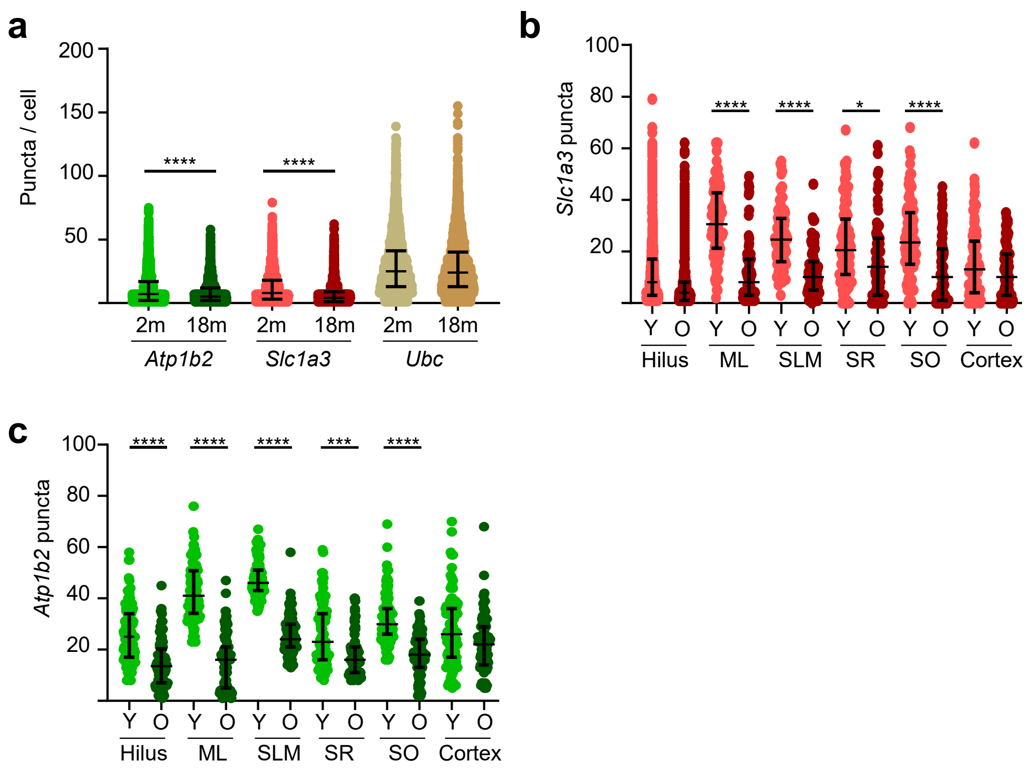

### Supplementary Figure 2

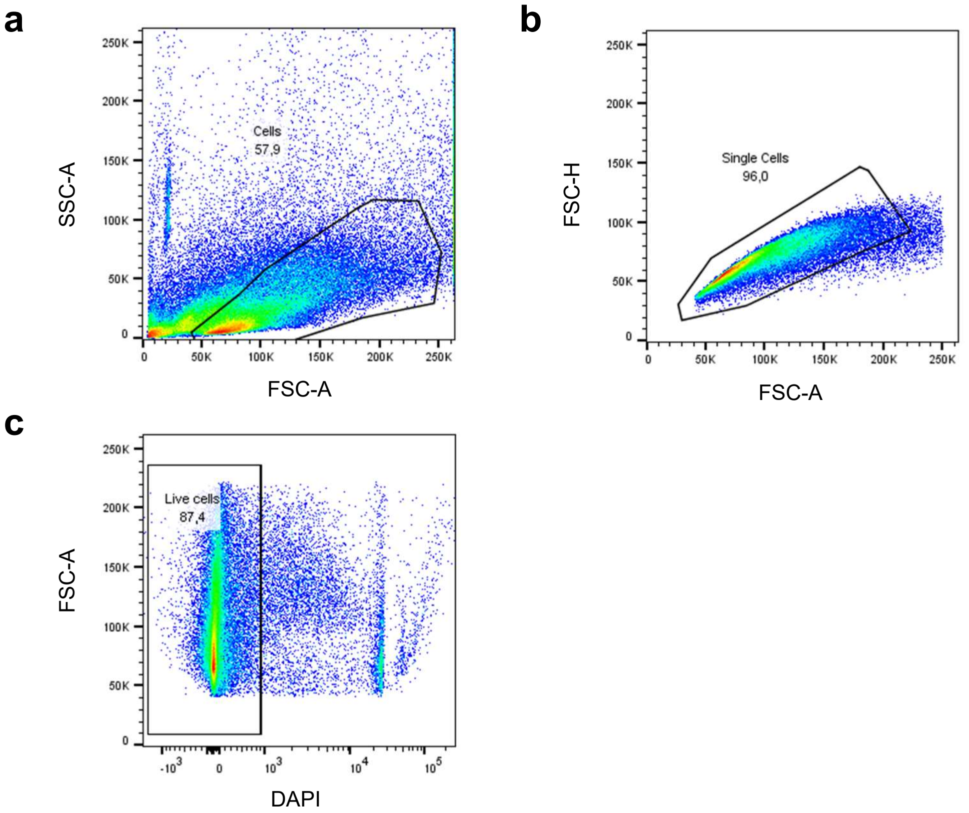

### Supplementary Figure 3

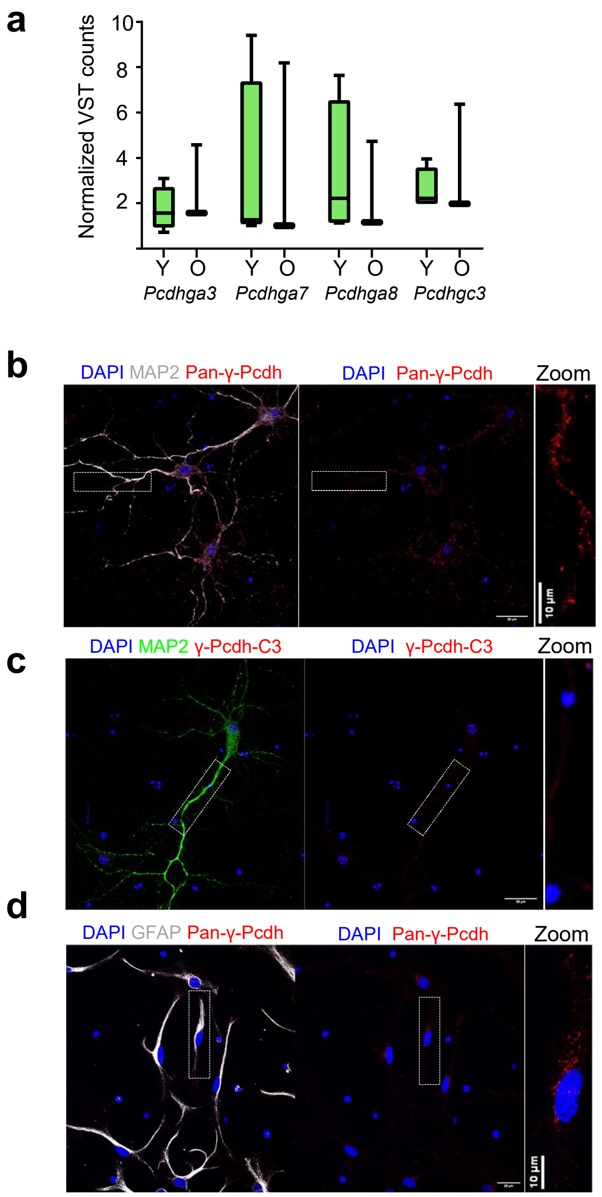

**Table S1.** RNAscope probes used for in situ hybridization.

| Gene name | ACD product code | Fluorescence channel | mRNA accession number |
|---|---|---|---|
| *Slc1a3* | 430781-C2 | C2 | NM_148938.3 |
| *Atp1b2* | 417131-C1 | C1 | NM_013415.5 |
| *Ubc* | 310771-C3 | C3 | NM_019639.4 |
| *Mvd* | 1005611-C3 | C3 | NM_138656.2 |

**Table S2.** Top 500 highest expressed genes among $A^+$ astrocytes.

| Gene | 2m-N2-$A^+$ | 2m-N3-$A^+$ | 2m-N5-$A^+$ | 2m-N6-$A^+$ | 18m-N2-$A^+$ | 18m-N3-$A^+$ | 18m-N4-$A^+$ | Mean expression |
|---|---|---|---|---|---|---|---|---|
| Malat1 | 17,59182119 | 17,0541673 | 17,31906036 | 18,05203251 | 21,68000006 | 18,12372925 | 16,84442244 | 18,0950333 |
| mt-Rnr2 | 18,06393355 | 15,10883311 | 15,93140522 | 17,10533937 | 20,20486643 | 18,80337561 | 15,57207657 | 17,25568998 |
| mt-Rnr1 | 17,13883845 | 14,36617911 | 14,82322788 | 16,09801568 | 20,10126436 | 16,41142888 | 14,57934949 | 16,21690055 |
| COX1 | 16,20626933 | 14,19085929 | 14,77786042 | 15,70577893 | 20,38115462 | 15,4492544 | 14,81468078 | 15,9322654 |
| ATP6 | 15,65476013 | 13,83018321 | 14,1534589 | 15,59871594 | 19,90528006 | 15,36619976 | 14,94231398 | 15,63584457 |
| CYTB | 15,31380988 | 13,57936853 | 14,58958643 | 14,5164619 | 20,50425423 | 15,42531076 | 14,79361572 | 15,53177249 |
| Cst3 | 14,20553366 | 14,01807383 | 13,97059863 | 14,84322193 | 21,24247295 | 13,50696644 | 15,09401885 | 15,26869804 |
| ND4 | 15,32187435 | 13,50548828 | 13,92757702 | 15,30276336 | 19,32429916 | 15,14850611 | 14,17973111 | 15,24431991 |
| COX3 | 15,97631135 | 13,36394874 | 13,73902543 | 15,37847931 | 18,64438431 | 14,54112799 | 14,30539479 | 15,13552456 |
| COX2 | 14,80221434 | 12,73288179 | 13,32141402 | 15,34167958 | 19,41063004 | 14,4612935 | 13,92378253 | 14,85627083 |
| Slc1a2 | 12,88080499 | 13,67303985 | 12,77944778 | 14,65924697 | 18,89005598 | 13,73777412 | 13,58665944 | 14,31528988 |
| Apoe | 12,62497146 | 13,66402449 | 13,20492033 | 13,93009788 | 18,09070716 | 14,90165878 | 13,63695237 | 14,29333321 |
| Atp1a2 | 13,24304252 | 13,78565256 | 13,42197688 | 13,41759502 | 16,46613577 | 15,57366088 | 13,54524362 | 14,20761532 |
| Plpp3 | 12,60616442 | 12,86896404 | 12,5957882 | 13,84551275 | 17,72819176 | 15,41560336 | 13,5197785 | 14,08285757 |
| ND1 | 13,52548544 | 12,01259235 | 12,73204458 | 13,59249346 | 18,82097883 | 14,38057919 | 13,45696473 | 14,07444837 |
| ND2 | 14,5263981 | 12,01626454 | 12,69495491 | 14,36534603 | 19,6818231 | 11,61288762 | 12,99151001 | 13,98416919 |
| Ptn | 11,94122253 | 12,6516276 | 12,30083967 | 14,30766233 | 19,49654256 | 13,21283999 | 13,06908379 | 13,85425978 |
| Clu | 11,04087181 | 13,53858388 | 12,29236394 | 13,03837066 | 19,54541585 | 12,77286493 | 13,28493868 | 13,64477282 |
| Slc1a3 | 11,17108987 | 13,61642821 | 12,88476997 | 10,40013291 | 18,37972232 | 13,04790505 | 14,16308586 | 13,38044774 |
| Scd2 | 13,03274188 | 11,54473354 | 12,27183671 | 13,16521067 | 17,05776844 | 13,22944613 | 12,08277227 | 13,19778709 |
| ND5 | 13,73509929 | 11,10791013 | 12,4000238 | 13,85353897 | 18,23984521 | 11,24039597 | 11,17989456 | 13,10810113 |
| Gja1 | 11,66643326 | 11,91970407 | 11,95712584 | 13,69007185 | 18,037879 | 11,5119335 | 12,76126076 | 13,07777261 |
| Cpe | 11,38619283 | 12,07646308 | 11,6344133 | 13,28766993 | 18,38835298 | 12,57105969 | 11,97027914 | 13,04491871 |
| Sparcl1 | 12,86109272 | 13,63680617 | 12,95766793 | 12,12477509 | 12,38530756 | 14,28376303 | 13,0427376 | 13,04173573 |
| ND6 | 12,84626484 | 10,31261057 | 11,24292743 | 13,27674089 | 17,68586773 | 13,83895427 | 11,69859141 | 12,98599388 |
| Ckb | 11,68754271 | 11,77740849 | 12,08050615 | 11,96325311 | 16,8144562 | 12,1629995 | 12,94587361 | 12,77600568 |
| Plp1 | 16,85502352 | 12,52447573 | 13,78045636 | 14,38671516 | 10,57068491 | 7,945448092 | 12,90505903 | 12,70969469 |
| Scg3 | 11,62659361 | 11,05773763 | 11,32071569 | 11,79719494 | 18,62383268 | 12,54990709 | 11,83871765 | 12,68781418 |
| OC11856734 | 10,07495706 | 12,37131998 | 12,46278209 | 12,83352166 | 15,83907175 | 13,1856352 | 11,27027302 | 12,57679439 |

| | | | | | | | | |
|---|---|---|---|---|---|---|---|---|
| Son | 11,35182154 | 12,1627001 | 12,30668666 | 12,56673408 | 14,25607535 | 12,79598191 | 11,89068638 | 12,47581229 |
| ATP8 | 12,67669328 | 10,68092758 | 10,98233394 | 12,04485544 | 16,44550653 | 11,80104368 | 11,84641224 | 12,35396753 |
| Prnp | 11,34367387 | 11,09185991 | 11,47525719 | 11,50579955 | 16,97105594 | 12,06272662 | 11,95154456 | 12,34313109 |
| Aldoc | 10,41845786 | 11,90231383 | 11,3012823 | 9,61401775 | 18,68384038 | 10,43916534 | 13,07392788 | 12,20471505 |
| Gpm6b | 11,50365898 | 12,61133598 | 11,38714579 | 13,33666772 | 11,43294055 | 12,97730138 | 12,15256372 | 12,20023059 |
| Gpm6a | 12,4314026 | 12,38590707 | 11,00830986 | 10,55933872 | 12,90168715 | 12,50367752 | 13,28857353 | 12,15412807 |
| Htra1 | 10,01634898 | 12,00086209 | 11,47355487 | 8,105268247 | 17,6989776 | 12,52332671 | 13,09595911 | 12,13061394 |
| Calm1 | 13,16943021 | 10,59747146 | 11,62989181 | 11,69165755 | 16,54423657 | 8,019433442 | 12,45179726 | 12,01484547 |
| Car2 | 12,49090318 | 9,724078625 | 11,51957558 | 11,95351527 | 18,33110293 | 8,535288706 | 11,18487451 | 11,96276269 |
| Ccdc88a | 11,28675307 | 11,51891954 | 11,32655903 | 10,83269072 | 14,496652 | 12,24230618 | 11,31357162 | 11,85963602 |
| Ncoa4 | 11,16151291 | 11,89168012 | 9,711846133 | 13,31939169 | 11,61048707 | 14,19376391 | 10,94808911 | 11,83382442 |
| Glul | 11,31187256 | 9,836720054 | 11,23514694 | 10,69344157 | 18,47869115 | 9,084474658 | 12,06962638 | 11,81571047 |
| Pcdh9 | 13,59149054 | 10,9690909 | 12,90345992 | 11,09568754 | 12,93495093 | 11,41197877 | 9,646191785 | 11,79326434 |
| Slc7a10 | 9,925243915 | 9,943770176 | 9,634916315 | 12,51137623 | 17,36542374 | 11,99884087 | 11,00905855 | 11,76980426 |
| Lsamp | 10,62355436 | 10,53644361 | 10,96969594 | 10,97846763 | 18,19549936 | 9,897010712 | 11,18785409 | 11,76978939 |
| Klf13 | 11,75321492 | 10,6513141 | 9,691016349 | 12,72723146 | 14,97520591 | 10,89415077 | 11,47549677 | 11,7382329 |
| Hnrnpa2b1 | 11,39439243 | 11,74862338 | 11,50876409 | 10,44256504 | 12,85904388 | 12,19390592 | 11,97661359 | 11,7319869 |
| Tmsb4x | 11,67101533 | 9,443500304 | 11,42817257 | 11,68421036 | 17,03094833 | 10,11529109 | 10,37459816 | 11,67824802 |
| Mt1 | 11,27500349 | 9,9687012 | 10,86898347 | 10,9253063 | 16,86012532 | 9,565829349 | 11,83200002 | 11,61370702 |
| Eef1a1 | 12,5595633 | 10,06417935 | 11,92664026 | 11,94392908 | 12,00077887 | 10,822739 | 11,77425725 | 11,58458387 |
| OC11856827 | 10,09799465 | 9,401741286 | 11,53204295 | 11,11959092 | 15,33150303 | 13,45726433 | 9,876365292 | 11,54521464 |
| Gstm1 | 10,95309506 | 10,07279662 | 10,74440285 | 7,209279557 | 17,8774487 | 11,69941787 | 12,03767954 | 11,51344574 |
| Gm33649 | 9,927873296 | 10,99891854 | 11,56446552 | 12,85721499 | 15,08669689 | 12,15478403 | 7,794087472 | 11,48343439 |
| Luc7l3 | 11,29526945 | 10,26614827 | 10,06402855 | 12,30380884 | 17,01453109 | 8,86144505 | 10,19186313 | 11,42815634 |
| Ntrk2 | 9,793526267 | 11,09151752 | 11,01176061 | 9,495474268 | 13,38216964 | 12,44867543 | 11,63182468 | 11,26499263 |
| Tspan7 | 10,2113272 | 12,91553023 | 11,2327277 | 12,15032425 | 8,132245398 | 12,9523862 | 11,20949732 | 11,25771976 |
| Slc4a4 | 8,278060654 | 11,65652156 | 10,5757844 | 11,77047991 | 12,02584257 | 12,6991191 | 11,72364243 | 11,24706438 |
| Hepacam | 9,691640427 | 9,67031256 | 9,027189898 | 10,85331082 | 17,68752485 | 10,85428447 | 10,83582558 | 11,23144123 |
| Dbi | 12,46185876 | 10,75477047 | 11,54030964 | 11,63988996 | 17,25349734 | 4,203018797 | 10,71815527 | 11,22450004 |
| Slc6a1 | 7,861686042 | 10,99243265 | 10,23943835 | 10,7753156 | 14,89414717 | 12,68121163 | 10,87981669 | 11,18914973 |
| Nrxn1 | 10,99425923 | 8,952775043 | 10,09136107 | 11,47052683 | 14,40372932 | 10,96943504 | 11,33559743 | 11,17395485 |
| Gm30618 | 10,59070363 | 8,997581165 | 8,573836763 | 12,13512206 | 14,85669495 | 12,57171697 | 10,26965107 | 11,14218666 |
| Cryab | 13,4301451 | 9,896943297 | 11,27275239 | 12,71052228 | 18,27172769 | 1,78350001 | 10,59958223 | 11,13788186 |

| | | | | | | | | |
|---|---|---|---|---|---|---|---|---|
| Hsp90aa1 | 15,05231825 | 11,1428814 | 12,27324307 | 13,37273218 | 1,566251539 | 13,68364289 | 10,70710105 | 11,11402434 |
| Pnisr | 9,153169282 | 11,6198429 | 11,13386398 | 11,31123823 | 11,99026601 | 12,14587856 | 10,2495078 | 11,08625239 |
| Brd7 | 10,46407876 | 8,118197257 | 9,440788825 | 11,16101364 | 16,08470718 | 11,26508022 | 10,90739756 | 11,06303764 |
| Gpr37l1 | 11,50585086 | 11,76307271 | 9,414990855 | 11,79039533 | 8,578203729 | 12,72815052 | 11,51886935 | 11,04279048 |
| Gstm5 | 9,951236063 | 10,44626122 | 9,948379076 | 12,27673892 | 16,03193275 | 7,535490943 | 11,08616853 | 11,03945821 |
| Syt11 | 11,4550662 | 9,150023411 | 11,11355911 | 10,83982267 | 11,7818308 | 11,88551645 | 10,95305189 | 11,02555293 |
| Atp1b2 | 9,441400925 | 10,7870128 | 10,46622663 | 5,709433669 | 18,27872986 | 9,808362798 | 12,63602343 | 11,01817002 |
| Zbtb20 | 10,49828304 | 12,09268015 | 9,82454914 | 12,36624158 | 10,21448621 | 12,08507344 | 9,871277846 | 10,99322734 |
| Gstp1 | 10,99870964 | 6,34995433 | 11,06022768 | 11,86900854 | 16,57288603 | 10,41991732 | 9,457963867 | 10,9612382 |
| Actb | 12,78175069 | 10,62103278 | 11,38328663 | 11,02600564 | 11,54664483 | 8,149614153 | 10,8801254 | 10,91263716 |
| Irf2bpl | 11,42360424 | 8,149002347 | 8,7210758 | 12,21681226 | 15,13890612 | 9,850877328 | 10,65032044 | 10,87865693 |
| Fam171b | 9,515441646 | 9,760149525 | 10,02557393 | 11,73641132 | 14,69023927 | 9,45068569 | 10,95746838 | 10,87656711 |
| Hspa8 | 12,68422312 | 12,33486573 | 11,43811433 | 12,13463129 | 5,173536434 | 11,2650953 | 11,08654409 | 10,87385861 |
| Srrm2 | 10,39455861 | 9,229119286 | 9,711422983 | 8,72439952 | 16,48641967 | 11,82654376 | 9,73163131 | 10,87201359 |
| Fgfr3 | 9,167983607 | 10,81354519 | 9,957212966 | 11,46109887 | 14,12857086 | 9,762335998 | 10,67081328 | 10,85165154 |
| Qk | 9,179807721 | 11,81503756 | 11,66659092 | 10,22427259 | 12,92954798 | 9,479953504 | 10,58607226 | 10,84018322 |
| Mt3 | 10,46821655 | 9,211970543 | 9,362373464 | 10,64045132 | 14,99864377 | 10,70864646 | 10,44283837 | 10,83330578 |
| Ubc | 11,26309947 | 9,870038037 | 8,186920901 | 11,66499521 | 12,99704726 | 10,67311234 | 10,8906069 | 10,79226002 |
| Srsf11 | 8,51314955 | 9,269847377 | 9,470412406 | 9,940259489 | 16,21667822 | 11,7770398 | 10,24463336 | 10,77600289 |
| Gapdh | 8,619581319 | 8,627181423 | 9,884820343 | 10,39775322 | 17,03423166 | 10,68861668 | 10,10092981 | 10,76473064 |
| S1pr1 | 9,955517279 | 12,39716667 | 11,12338055 | 8,913794186 | 9,251634058 | 11,63045613 | 11,9394878 | 10,74449095 |
| Id2 | 9,510358072 | 10,89023621 | 9,49431701 | 12,55506322 | 13,21562791 | 10,24073303 | 9,201151928 | 10,72964105 |
| Rbm39 | 11,19247012 | 11,85538988 | 10,50256944 | 12,47571206 | 5,890413279 | 12,76814802 | 10,29713509 | 10,71169113 |
| Itm2b | 11,38351485 | 10,72186359 | 11,15504958 | 10,12104816 | 17,34230472 | 3,008922739 | 11,1875295 | 10,70289045 |
| Ubb | 13,71743354 | 13,44965554 | 12,88997681 | 11,24347696 | 2,315742648 | 9,433423481 | 11,77267347 | 10,68891178 |
| ND3 | 11,38461881 | 7,977494779 | 9,685715729 | 11,62423892 | 12,43435222 | 11,92849866 | 9,735340148 | 10,68146561 |
| Rbm25 | 10,13994915 | 8,717187429 | 9,201001993 | 10,43800746 | 18,3073657 | 7,547031243 | 10,25399406 | 10,65779101 |
| Clmn | 10,80784489 | 10,12638519 | 8,428734498 | 12,59263409 | 14,05920533 | 8,519268617 | 9,949618499 | 10,6405273 |
| Cspg5 | 10,22669549 | 8,492118033 | 9,35886144 | 10,2853801 | 15,86683293 | 9,666936585 | 10,55308755 | 10,63570173 |
| Atp2a2 | 10,67190559 | 8,092646846 | 10,12462109 | 10,56990635 | 14,67208685 | 11,16399807 | 8,212139188 | 10,50104343 |
| Gria2 | 11,27090511 | 12,4618422 | 10,61844928 | 12,13905532 | 1,489534757 | 14,79581304 | 10,67093637 | 10,4923623 |
| Cd81 | 12,14170263 | 12,53962275 | 10,80126875 | 10,37301125 | 4,479335813 | 11,43419015 | 11,58318624 | 10,47890251 |
| Tpt1 | 11,22338401 | 8,528141615 | 10,11386644 | 10,36080814 | 15,54640179 | 7,279376693 | 10,27689196 | 10,47555295 |

| | | | | | | | | |
|---|---|---|---|---|---|---|---|---|
| Gas7 | 10,58706679 | 9,251579162 | 8,93168293 | 11,41311534 | 13,19572376 | 10,08142711 | 9,859514939 | 10,47430143 |
| Atp5b | 11,16159384 | 8,828212256 | 10,44868484 | 9,384581155 | 18,46764958 | 4,694168184 | 10,28284897 | 10,46681983 |
| Fth1 | 13,11831118 | 12,17039241 | 11,37235625 | 11,21537008 | 6,97930818 | 7,44934211 | 10,92281265 | 10,46112755 |
| Tcf4 | 10,37623711 | 11,26098868 | 10,26883499 | 13,51055078 | 10,14970923 | 7,758154927 | 9,521862959 | 10,40661981 |
| Gm52229 | 7,541426077 | 10,66193813 | 10,15052374 | 11,84332628 | 10,01773531 | 13,68570354 | 8,906096782 | 10,40096427 |
| Luc7l2 | 10,23994299 | 8,026111069 | 9,633875238 | 11,16908203 | 14,0994231 | 10,31681764 | 9,27214343 | 10,39391364 |
| Ktn1 | 10,82759403 | 10,44302718 | 10,63715568 | 11,37661207 | 9,523899295 | 10,81560035 | 9,00465039 | 10,37550557 |
| Mt2 | 9,34620461 | 8,380295152 | 9,067041241 | 11,10396001 | 15,77241612 | 9,268760284 | 9,570934685 | 10,35851601 |
| Acsl3 | 10,13595323 | 11,71006703 | 10,82765305 | 9,92134447 | 8,489387015 | 10,60520407 | 10,56571409 | 10,32218899 |
| Prdx6 | 8,360232239 | 9,867234403 | 10,33480224 | 6,69559057 | 17,15867374 | 8,868649334 | 10,49135866 | 10,2537916 |
| OC11856734 | 7,579794423 | 9,961560549 | 7,462522287 | 11,001798 | 12,78100888 | 12,74133566 | 10,21077674 | 10,24839951 |
| Slc25a5 | 6,886956744 | 10,4242997 | 9,443149636 | 9,483673756 | 16,40788665 | 9,663640377 | 9,035641303 | 10,19217831 |
| Insig1 | 8,076936074 | 9,313539148 | 9,062253285 | 11,2953769 | 15,74133318 | 7,970477643 | 9,830051404 | 10,18428109 |
| Gm15521 | 10,55064415 | 10,99331278 | 11,15419858 | 8,414933694 | 9,545834989 | 11,83644034 | 8,769200399 | 10,18065213 |
| Ddx5 | 11,09957974 | 12,65917595 | 10,90369883 | 12,08069796 | 1,854343835 | 12,11172237 | 10,49490876 | 10,17201821 |
| Ddhd1 | 8,484035009 | 10,11555064 | 10,30212827 | 9,140269722 | 11,89806718 | 11,43996633 | 9,541243914 | 10,13160872 |
| Syne1 | 12,19778389 | 12,09274282 | 8,75243986 | 12,18176953 | 7,410284941 | 6,709535747 | 11,56198937 | 10,1295066 |
| Gnas | 10,22064612 | 11,09844884 | 10,08856733 | 11,32244461 | 6,127260845 | 12,07552784 | 9,945475633 | 10,1254816 |
| Hsp90ab1 | 13,74897147 | 10,90915454 | 11,12200284 | 11,24816558 | 4,373848877 | 8,71501397 | 10,64825233 | 10,10934423 |
| Bcan | 10,49397486 | 10,40315521 | 9,922384808 | 4,459024283 | 11,46476588 | 12,3143684 | 11,51346491 | 10,08159119 |
| Ptprz1 | 10,83693884 | 12,4984477 | 9,95705424 | 9,824116588 | 1,815046859 | 13,99261558 | 11,62904966 | 10,07903849 |
| Macf1 | 10,35629717 | 11,85471217 | 10,62423367 | 11,43836085 | 4,969521949 | 10,22547924 | 11,03972271 | 10,07261825 |
| Chchd2 | 9,168292647 | 8,199309576 | 8,35900421 | 7,420135652 | 16,53634145 | 9,874643102 | 10,80113829 | 10,05126642 |
| Slc27a1 | 9,245013508 | 9,37127571 | 7,756087527 | 9,897568157 | 15,09615172 | 7,152909938 | 11,61589663 | 10,01927188 |
| Arhgap5 | 13,88316853 | 11,7982646 | 10,42158226 | 10,39069197 | 2,071732111 | 10,74231798 | 10,73297395 | 10,00581877 |
| Arglu1 | 10,4773786 | 5,624263014 | 9,591171305 | 10,73456002 | 14,04065328 | 10,30041345 | 9,231385535 | 9,999975029 |
| Ahcyl1 | 4,839679671 | 10,69637498 | 9,473550656 | 9,129485252 | 15,10052908 | 9,790760788 | 10,80646527 | 9,976692242 |
| Psap | 11,54490072 | 8,676148784 | 10,82591158 | 8,80428033 | 9,456721698 | 11,42607308 | 9,069776338 | 9,971973219 |
| Ptma | 12,99635149 | 10,8449438 | 11,0271584 | 10,27968483 | 4,726226904 | 9,727760699 | 10,04316208 | 9,949326885 |
| Srsf2 | 7,780452273 | 9,999567332 | 10,48284078 | 9,337516383 | 12,39245806 | 10,6112755 | 8,996043833 | 9,942879166 |
| Pcbp2 | 9,960399 | 9,404057098 | 10,05503009 | 8,396699133 | 16,16340138 | 5,994175142 | 9,622026004 | 9,942255407 |
| OC11548946 | 8,541398385 | 9,949720325 | 9,085015751 | 10,14379679 | 11,22310196 | 12,33474342 | 8,159112418 | 9,91955558 |
| S100a1 | 7,772143729 | 9,905605246 | 8,621925221 | 11,04933868 | 17,38604807 | 4,777691847 | 9,685866627 | 9,88551706 |

| | | | | | | | | |
|---|---|---|---|---|---|---|---|---|
| Atrx | 10,48676786 | 9,039354965 | 9,244507864 | 10,13512812 | 10,30406281 | 9,695812179 | 10,09638366 | 9,857431066 |
| Sparc | 11,87973977 | 10,35315573 | 10,36962939 | 10,81304004 | 1,849823661 | 12,95118464 | 10,73623715 | 9,850401483 |
| Aldoa | 10,86252307 | 8,221444393 | 10,2064433 | 7,957488276 | 19,53668096 | 1,518442021 | 10,39750135 | 9,814360483 |
| Ank2 | 10,52506454 | 11,17748069 | 10,27909679 | 9,559810964 | 3,176105524 | 12,42035092 | 11,38194781 | 9,788551034 |
| Mapre1 | 3,952248667 | 10,39494371 | 8,756314089 | 10,032142 | 13,45772185 | 11,15809891 | 10,53301797 | 9,754926741 |
| Gjb6 | 8,087773806 | 12,36925274 | 9,521255637 | 12,24346166 | 1,826896382 | 13,70453214 | 10,39799845 | 9,735881544 |
| Mfge8 | 9,967656019 | 11,82109009 | 9,748809744 | 11,47754355 | 1,630722107 | 12,44088459 | 10,80583419 | 9,698934327 |
| Uros | 11,44911535 | 6,873166511 | 8,068250545 | 10,73261015 | 7,393332398 | 13,60369883 | 9,760777001 | 9,697278683 |
| Serinc1 | 11,14611212 | 10,15296657 | 9,807638082 | 10,53468284 | 6,670527401 | 10,64395057 | 8,869587258 | 9,68935212 |
| Dst | 7,47536789 | 9,089679227 | 9,668773407 | 12,32979978 | 11,61508559 | 10,33213305 | 7,310720376 | 9,688794189 |
| Calm2 | 11,17848989 | 7,90717105 | 9,755668035 | 9,324633345 | 18,42556383 | 1,367655022 | 9,715599406 | 9,667825797 |
| Kif1b | 11,77969393 | 11,0826933 | 10,96580379 | 11,33738464 | 1,688368716 | 10,87994759 | 9,890887331 | 9,660682757 |
| Gm51419 | 8,565612354 | 9,512956519 | 10,11575074 | 10,75099046 | 9,876348783 | 11,8858088 | 6,898815664 | 9,658040474 |
| F3 | 10,62311789 | 10,79275603 | 10,93273779 | 3,944650956 | 6,279178143 | 13,64018263 | 11,31318513 | 9,64654408 |
| Aplp1 | 12,21832313 | 9,311989631 | 10,80087511 | 8,952145926 | 6,543165536 | 10,25360356 | 9,409235271 | 9,641334025 |
| Paqr8 | 9,255153187 | 10,52339487 | 9,749775193 | 10,22859718 | 6,364558172 | 11,95774367 | 9,373681503 | 9,63612911 |
| Cadm1 | 10,04837436 | 10,3913635 | 8,525772726 | 11,99270304 | 4,232455718 | 11,31343186 | 10,93173262 | 9,633690546 |
| Pde4b | 10,01418305 | 7,637109749 | 9,619891537 | 8,121121791 | 12,76121488 | 9,995397742 | 9,27122434 | 9,631449012 |
| Dclk1 | 8,997269012 | 12,48813549 | 11,59532827 | 4,593153098 | 11,05789532 | 7,848314835 | 10,82964789 | 9,629963416 |
| Gm6145 | 8,510014912 | 7,887772233 | 10,24358464 | 10,06422256 | 12,61932419 | 10,0062425 | 7,986663428 | 9,616832067 |
| Gpc5 | 7,538321197 | 9,445406359 | 10,06029749 | 8,855260228 | 12,4014423 | 10,76287391 | 8,252382383 | 9,616569123 |
| Prdx1 | 9,634892814 | 10,22724471 | 10,24119552 | 7,081836115 | 18,03662312 | 1,203598178 | 10,80943304 | 9,604974786 |
| Fos | 13,38120827 | 10,13921334 | 10,61574183 | 12,07966047 | 4,52434987 | 6,954931689 | 9,384445989 | 9,582793066 |
| Ntm | 9,672541319 | 9,590771899 | 9,374021548 | 9,905125658 | 6,609640361 | 12,40236044 | 9,290776846 | 9,549319724 |
| Ash1l | 6,660571273 | 10,7568208 | 5,934342654 | 12,07112088 | 12,84202959 | 8,752964906 | 9,751259057 | 9,538444166 |
| Gnao1 | 9,177484892 | 9,985682726 | 9,543502676 | 4,440357244 | 14,66636421 | 8,018673741 | 10,89763944 | 9,53281499 |
| Etfa | 11,01771998 | 6,639456855 | 8,327251824 | 9,915705962 | 13,04369702 | 8,579466307 | 9,149401251 | 9,524671314 |
| Mal | 14,78037729 | 10,12038842 | 11,59772198 | 12,52920271 | 2,676094516 | 4,459530528 | 10,4452817 | 9,515513876 |
| Rpl6 | 12,80291695 | 10,21918358 | 9,061785221 | 12,44922856 | 1,740004178 | 10,82783041 | 9,421204586 | 9,503164784 |
| Cfl1 | 6,519314278 | 9,657517775 | 7,938214435 | 8,514372901 | 17,31995097 | 7,208779151 | 9,280147585 | 9,4911853 |
| Camk2g | 10,15764165 | 9,319152565 | 9,838691389 | 4,41473219 | 13,03879335 | 9,863095421 | 9,806054045 | 9,491165801 |
| Ppia | 10,13687625 | 11,40845105 | 11,14216648 | 3,244125863 | 18,10980838 | 0,646152028 | 11,70749694 | 9,485010998 |
| Hnrnpu | 10,78932672 | 10,74598813 | 10,52638591 | 11,81658115 | 1,525086314 | 12,40201101 | 8,532786603 | 9,476880834 |

| | | | | | | | | |
|---|---|---|---|---|---|---|---|---|
| Nfia | 9,42967306 | 9,364436491 | 10,22308578 | 8,206251077 | 8,358974386 | 10,54268416 | 10,15442626 | 9,46850446 |
| Eif1 | 10,13474521 | 9,740938889 | 10,57473601 | 8,798249472 | 15,956699 | 0,526554958 | 10,52251931 | 9,464920407 |
| Myo10 | 8,990617069 | 9,569454469 | 10,81652636 | 5,30165142 | 12,5706221 | 10,58663091 | 8,189768458 | 9,43218154 |
| Appl2 | 9,030079406 | 10,1519765 | 10,92043543 | 9,036591034 | 9,612485357 | 9,705263904 | 7,470381719 | 9,418173336 |
| Ndrg2 | 8,405940249 | 12,19316452 | 10,27523226 | 10,74334515 | 5,564097712 | 8,624941439 | 10,06490739 | 9,410232674 |
| Serpine2 | 7,579754767 | 10,81863187 | 9,621476722 | 9,057754867 | 9,837387715 | 8,379653374 | 10,51507455 | 9,401390551 |
| ND4L | 9,578025342 | 8,215136009 | 8,448995612 | 10,62186145 | 14,49054927 | 6,070278037 | 8,353156867 | 9,396857513 |
| Myo6 | 11,0716607 | 11,55799997 | 8,235641399 | 12,14337239 | 1,593467405 | 11,63347481 | 9,324468258 | 9,365726421 |
| Kirrel3 | 7,0816903 | 11,40928065 | 8,294570414 | 8,051053569 | 9,640524408 | 12,37074625 | 8,703518291 | 9,364483412 |
| Cmtm5 | 11,09401831 | 9,333391891 | 8,307986897 | 9,135103175 | 9,665745246 | 7,269453953 | 10,7319592 | 9,362522667 |
| Prpf4b | 9,481631189 | 4,86481621 | 8,33042212 | 10,48735166 | 13,24126667 | 10,54963263 | 8,516283281 | 9,35305768 |
| Daam2 | 8,824551086 | 8,825198468 | 7,480430885 | 10,15550938 | 15,05762922 | 5,770262036 | 9,349366382 | 9,351849636 |
| Prrc2c | 11,05546912 | 5,436937974 | 9,529170788 | 11,87542173 | 6,173229124 | 12,96352165 | 8,415376644 | 9,349875289 |
| Itm2c | 9,797409014 | 8,940127517 | 8,171823723 | 8,232498293 | 11,38141383 | 8,304859018 | 10,53514208 | 9,337610496 |
| Tuba1a | 9,289956369 | 11,19759392 | 9,228280135 | 10,79024585 | 9,354462279 | 5,797720042 | 9,654397091 | 9,330379383 |
| Reep3 | 10,59309408 | 5,397665685 | 9,372888777 | 12,58362144 | 7,869337003 | 10,75236831 | 8,720194377 | 9,327024239 |
| Ldhb | 9,192635428 | 11,94924699 | 9,634075921 | 8,662593226 | 1,785458715 | 13,16307968 | 10,78770321 | 9,310684738 |
| Uqcr10 | 9,157522092 | 7,076041114 | 8,344172723 | 6,930918847 | 15,84757692 | 7,755849473 | 10,04140473 | 9,307640842 |
| Luzp2 | 11,09205919 | 10,41717735 | 10,87024002 | 8,934677591 | 6,270161639 | 8,420694368 | 9,129442933 | 9,30492187 |
| Gm51889 | 9,098111106 | 7,149946802 | 7,233141392 | 10,71228884 | 12,10166447 | 11,13513761 | 7,685304183 | 9,302227772 |
| Plcd4 | 9,139178018 | 8,839747309 | 8,88774727 | 7,332994759 | 14,41925809 | 6,761495428 | 9,687335293 | 9,295393739 |
| Ttyh1 | 9,236399595 | 11,45430318 | 9,533205271 | 7,476004528 | 1,47640703 | 14,56967767 | 11,30267275 | 9,292667147 |
| Gm3764 | 9,206245765 | 8,987526097 | 9,244165488 | 9,825106364 | 9,20447421 | 9,935774099 | 8,583259714 | 9,283793105 |
| Kcnq1ot1 | 8,701378978 | 10,19490807 | 9,208281124 | 12,07704773 | 5,80263937 | 11,38992972 | 7,594482969 | 9,28123828 |
| Ubr5 | 8,132563316 | 11,01957747 | 7,844733556 | 12,37764566 | 3,830413008 | 12,93001531 | 8,803723823 | 9,276953163 |
| Lamc1 | 9,164097504 | 5,904055943 | 7,74192698 | 10,45898987 | 13,65167426 | 9,829395773 | 8,181477971 | 9,275945472 |
| Brd9 | 8,525109664 | 7,66588435 | 8,119857865 | 12,81448837 | 12,81409736 | 7,45151252 | 7,448696113 | 9,262806606 |
| Gabrb1 | 7,413651813 | 11,46806446 | 9,913536784 | 8,528635607 | 4,702600036 | 12,07358319 | 10,66983335 | 9,252843607 |
| Ednrb | 5,110046935 | 12,68641055 | 9,515280733 | 11,07703129 | 2,364319678 | 14,51960612 | 9,386347685 | 9,237006141 |
| Snrnp70 | 8,74808067 | 8,614758377 | 7,97586817 | 10,40626449 | 11,53008802 | 9,521374495 | 7,77718379 | 9,224802574 |
| Sox9 | 11,00312659 | 9,707533063 | 9,284323404 | 9,881790672 | 1,896363731 | 12,33214315 | 10,41104315 | 9,216617679 |
| Ankrd12 | 4,005915233 | 9,165176002 | 9,781327402 | 9,53906968 | 12,28708077 | 11,31032221 | 8,410432857 | 9,214189165 |
| Zfp148 | 7,222001148 | 4,138530119 | 6,880570819 | 12,28176758 | 18,38094551 | 6,917009755 | 8,649523249 | 9,210049741 |

| | | | | | | | | |
|---|---|---|---|---|---|---|---|---|
| Hsp90b1 | 10,97113022 | 7,136752147 | 10,01030482 | 11,41667685 | 2,575438529 | 13,14096464 | 9,200171992 | 9,207348458 |
| App | 13,38549235 | 5,743460097 | 10,22409626 | 10,84712131 | 2,528029115 | 12,67115177 | 8,952716866 | 9,193152539 |
| Septin7 | 11,09224607 | 11,28647891 | 10,76501457 | 10,71335217 | 2,307032112 | 8,562802265 | 9,502379211 | 9,175615044 |
| Tmem47 | 10,46111318 | 12,17062365 | 9,868635452 | 11,38620153 | 2,271345476 | 8,734164558 | 9,138029009 | 9,147158978 |
| Gm19417 | 8,91349414 | 9,535030063 | 9,118588656 | 8,242102964 | 7,505494982 | 11,62680382 | 9,082646286 | 9,146308702 |
| Hnrnpa0 | 7,208597152 | 6,940100114 | 7,50219332 | 8,900324671 | 17,3033793 | 7,534863569 | 8,577928325 | 9,138198064 |
| Eif4g2 | 10,6604674 | 8,735675688 | 9,066573302 | 10,51831729 | 5,040652741 | 10,72422596 | 9,192109087 | 9,134003066 |
| Gm39752 | 8,275018067 | 9,990186824 | 9,575549381 | 8,748284025 | 8,996166986 | 11,24031729 | 6,989282059 | 9,116400662 |
| Slc39a12 | 6,279950929 | 10,99314214 | 9,756273139 | 3,887067772 | 10,32478575 | 11,28759607 | 11,2679172 | 9,113818999 |
| Gm46110 | 8,008890053 | 7,769285403 | 10,5062593 | 7,170151523 | 10,68444623 | 11,06308654 | 8,59456311 | 9,113811737 |
| Gm46892 | 8,956024565 | 7,902832328 | 6,451432912 | 10,00875018 | 10,21815596 | 11,15209113 | 9,045415889 | 9,104957567 |
| Ncam1 | 10,14479341 | 10,66386437 | 9,677536268 | 9,241833205 | 4,478953478 | 10,38891319 | 9,107192907 | 9,100440975 |
| Cxcl14 | 0,980659624 | 9,717480674 | 8,997528911 | 8,142915268 | 15,54943939 | 11,58398367 | 8,629716034 | 9,08596051 |
| Laptm4a | 9,683545269 | 9,742469101 | 9,239286997 | 9,091010316 | 8,442813556 | 7,579416166 | 9,662506387 | 9,063006827 |
| Hspa5 | 9,785507321 | 9,186712142 | 9,769573888 | 11,58006303 | 2,516542055 | 10,82304304 | 9,630078838 | 9,04164576 |
| Tnrc6b | 9,858892791 | 6,148296075 | 8,103047844 | 8,068706999 | 11,82309349 | 10,51756151 | 8,764783095 | 9,040625973 |
| Pitpnc1 | 7,335507319 | 8,441328558 | 8,961310195 | 7,579333686 | 10,12274983 | 11,31988395 | 9,483950264 | 9,034866257 |
| Srrm1 | 10,70658736 | 6,789804211 | 9,800385064 | 6,186013723 | 15,06383857 | 5,120089917 | 9,520368633 | 9,026726782 |
| Trim9 | 6,733169184 | 11,40398602 | 9,981277997 | 9,902480123 | 3,27842326 | 12,20386355 | 9,672812938 | 9,025144724 |
| Tcf25 | 8,305116749 | 7,215415365 | 8,560974447 | 4,33996625 | 16,81040548 | 7,83351179 | 10,1028655 | 9,024036512 |
| Rpl21 | 10,10982256 | 9,302642434 | 8,896078412 | 10,17510945 | 3,977830183 | 11,83813363 | 8,843335263 | 9,020421705 |
| Ghitm | 6,951744682 | 7,136223873 | 6,383032238 | 13,37754948 | 9,879721198 | 11,19028318 | 8,213563351 | 9,018874 |
| Zfp207 | 8,28969849 | 8,846906206 | 8,801423166 | 8,303314509 | 8,508324797 | 12,95900369 | 7,344430529 | 9,007585913 |
| Rsrp1 | 10,71202408 | 10,95605081 | 9,182046013 | 9,317638739 | 4,277177477 | 8,872310088 | 9,734996199 | 9,007463345 |
| Rpl31 | 8,082833474 | 9,082291597 | 7,826374552 | 10,3525409 | 16,2717144 | 2,545457199 | 8,885871035 | 9,006726165 |
| Abca1 | 9,756520007 | 8,618991914 | 8,964726901 | 3,778932993 | 9,706634085 | 10,65243409 | 11,5647163 | 9,006136613 |
| Tspan15 | 6,131640774 | 0,868074797 | 5,585379553 | 12,19909326 | 18,28821664 | 11,69969308 | 8,212332405 | 8,997775787 |
| Gm41284 | 10,63669716 | 11,25410225 | 9,87225578 | 10,30339505 | 3,124354995 | 11,86256999 | 5,757776095 | 8,973021616 |
| Pla2g7 | 8,220881996 | 13,15304597 | 9,548197675 | 10,45858229 | 2,260199754 | 8,828977007 | 10,29518123 | 8,966437988 |
| Msmo1 | 11,30035091 | 8,994775289 | 9,878326736 | 8,718738298 | 1,699375987 | 12,05089709 | 10,09212737 | 8,962084526 |
| Nfib | 8,663122973 | 9,496976073 | 8,018693715 | 8,460932641 | 14,81702545 | 5,378348384 | 7,83833872 | 8,95334828 |
| St3gal4 | 9,249938473 | 9,094288353 | 9,163605103 | 7,598897312 | 13,68556453 | 3,843444528 | 10,03411907 | 8,952836768 |
| Sfxn1 | 8,82591485 | 3,661970995 | 8,920231824 | 9,714598785 | 11,76214134 | 10,58327211 | 9,030976488 | 8,928443771 |

| | | | | | | | | |
|---|---|---|---|---|---|---|---|---|
| Crebbp | 11,53811615 | 9,624366774 | 8,901393814 | 11,24101305 | 1,224891244 | 10,72639347 | 9,169467059 | 8,917948795 |
| H2az2 | 8,716217398 | 7,20507238 | 8,495164518 | 3,759916299 | 16,2484247 | 9,124744664 | 8,805978729 | 8,907931242 |
| Dmd | 3,780498841 | 11,32918332 | 9,91555658 | 8,566962346 | 7,479469257 | 12,72083095 | 8,431514662 | 8,889145137 |
| Ntsr2 | 7,417365538 | 10,17393097 | 9,362972917 | 2,106149181 | 16,38141257 | 5,328882476 | 11,3628433 | 8,87622242 |
| Acot13 | 9,669336155 | 5,554128419 | 7,346306213 | 9,612556837 | 16,95679726 | 3,007966341 | 9,843614176 | 8,855815057 |
| Rpl32 | 10,74224567 | 8,838868801 | 9,165211526 | 7,109583148 | 16,57564156 | 0,822036872 | 8,661087572 | 8,844953593 |
| Mertk | 9,537774842 | 9,58033036 | 9,758998603 | 8,814187822 | 1,496343047 | 12,97043986 | 9,750443906 | 8,844074062 |
| Cnp | 12,00897496 | 8,687829238 | 10,17751947 | 10,88806903 | 7,852305266 | 3,189964734 | 9,007370625 | 8,830290474 |
| Akap9 | 9,966963415 | 10,48165585 | 9,922973907 | 9,915024033 | 13,09371263 | 3,116573388 | 5,248375747 | 8,820754139 |
| Tmem151a | 13,38565205 | 8,792442128 | 7,2847265 | 11,6166202 | 7,417582369 | 3,591482474 | 9,525249283 | 8,801965 |
| Tril | 3,475533455 | 11,14832246 | 8,933232794 | 9,536584053 | 4,61551113 | 13,21521358 | 10,51971044 | 8,777729702 |
| Sf3b1 | 12,03599426 | 8,995497027 | 9,300417585 | 10,25159043 | 1,609703998 | 9,321790032 | 9,868389697 | 8,769054718 |
| Actg1 | 11,57963423 | 10,25879052 | 9,608677024 | 9,997566468 | 1,71133036 | 9,773082013 | 8,427781887 | 8,765266071 |
| Mlc1 | 10,12413523 | 8,357121717 | 10,12705058 | 3,683370684 | 10,11542422 | 8,977091147 | 9,871147068 | 8,750762949 |
| Rcn2 | 2,01753739 | 11,9047039 | 7,922667979 | 11,55814761 | 4,569015224 | 13,499052 | 9,780289813 | 8,750201987 |
| Glud1 | 5,066107786 | 6,323115341 | 9,067984022 | 7,689302528 | 9,874619367 | 13,39415041 | 9,76380725 | 8,73986953 |
| Gm32337 | 9,010099677 | 8,786711373 | 8,751784089 | 10,44722419 | 5,141857024 | 11,37802922 | 7,654280142 | 8,738569387 |
| Srpk2 | 11,23880348 | 10,59774496 | 9,418643945 | 10,95690737 | 1,585250845 | 8,089290688 | 9,256730607 | 8,734767415 |
| Dlgap4 | 10,76071637 | 9,362682939 | 9,709741244 | 7,299869332 | 14,61527876 | 4,412954946 | 4,934196583 | 8,727920025 |
| Cox7c | 8,883972173 | 9,447266611 | 9,240413835 | 9,500762358 | 5,671568998 | 8,700162945 | 9,587217724 | 8,718766378 |
| Rpl4 | 11,0416851 | 10,20879093 | 8,939535362 | 9,45901799 | 3,736460577 | 9,213259747 | 8,412750826 | 8,715928648 |
| Fam172a | 11,84972969 | 3,766132915 | 8,235068386 | 8,881424799 | 18,24808744 | 0,633999416 | 9,392491822 | 8,715276353 |
| Parp14 | 6,939909251 | 6,78673243 | 6,324429687 | 9,576520138 | 16,28622242 | 7,107805466 | 7,95439616 | 8,710859365 |
| Fnbp1 | 10,20852225 | 8,396251142 | 8,083875765 | 9,425718362 | 4,492139911 | 11,66594844 | 8,661242439 | 8,704814043 |
| Rpl18 | 9,027423479 | 6,941525167 | 9,686925118 | 4,262265921 | 11,76371354 | 8,596300979 | 10,61823173 | 8,699483705 |
| Mmd2 | 10,26760195 | 11,10333312 | 10,35798496 | 5,906878343 | 1,067728867 | 11,61321182 | 10,49469071 | 8,68734711 |
| Add1 | 5,697785024 | 2,944127413 | 8,577620485 | 10,2396596 | 14,73162386 | 10,03261697 | 8,584021529 | 8,686779268 |
| Ptges3 | 12,04263717 | 3,493195054 | 9,006278806 | 11,52979662 | 14,35314421 | 2,161936478 | 8,180353306 | 8,681048806 |
| U2surp | 10,31795436 | 1,300256657 | 9,96260007 | 3,84588297 | 15,19179597 | 10,15558538 | 9,975411456 | 8,678498124 |
| Saraf | 9,497169964 | 9,112319412 | 9,269463281 | 5,875948444 | 14,52301556 | 2,268458237 | 10,18549877 | 8,675981953 |
| Ftl1 | 9,972815246 | 9,110342877 | 10,80801288 | 2,560315654 | 16,28019384 | 1,522092622 | 10,46757134 | 8,674477781 |
| Sub1 | 6,482481348 | 8,294929997 | 10,50564784 | 4,58931846 | 17,55571462 | 4,170944015 | 9,022724114 | 8,660251486 |
| 930402H24R | 11,40301585 | 10,76481606 | 10,60600923 | 6,707464364 | 4,985248025 | 7,137025191 | 8,832471818 | 8,633721506 |

| | | | | | | | | |
|---|---|---|---|---|---|---|---|---|
| Hacd2 | 7,11423029 | 9,746856572 | 9,018512284 | 7,489524776 | 4,678303835 | 12,66730827 | 9,652577968 | 8,623902 |
| Fau | 9,768673433 | 6,518583056 | 9,037458044 | 10,43440109 | 13,57232702 | 4,717271274 | 6,309672825 | 8,622626678 |
| Npas3 | 9,836475102 | 9,920424838 | 9,777394228 | 7,815346853 | 3,665013729 | 10,23638724 | 9,100231741 | 8,621610533 |
| OC11856771 | 10,01587645 | 6,711025096 | 9,651632643 | 7,568404223 | 10,9408532 | 7,901461982 | 7,516479181 | 8,615104681 |
| Chka | 0,22933109 | 7,468809389 | 9,044899376 | 10,24866289 | 14,44116436 | 9,757018759 | 9,085409576 | 8,610756491 |
| Pura | 9,251059461 | 9,382627201 | 7,21228341 | 11,00781411 | 4,746297579 | 9,701185897 | 8,683539457 | 8,569258159 |
| Nnat | 11,17036816 | 9,842195436 | 9,74900521 | 10,07570712 | 1,360564085 | 9,21374609 | 8,447423076 | 8,551287024 |
| Scaf11 | 7,694299597 | 6,788602167 | 8,886693808 | 11,16768527 | 7,964141282 | 11,02787819 | 6,238444415 | 8,538249247 |
| Sfpq | 12,4783127 | 6,847182993 | 8,029786687 | 10,23739776 | 2,1595639 | 11,39983625 | 8,586039133 | 8,53401706 |
| Nrcam | 10,60180403 | 10,05998535 | 8,824783858 | 10,13792978 | 1,600527632 | 9,885652602 | 8,598171924 | 8,529836452 |
| 330159H11R | 7,756410252 | 11,40843461 | 8,505777324 | 10,20078649 | 1,563575549 | 11,95610013 | 8,239762841 | 8,518692455 |
| Mbp | 13,56662975 | 10,37349059 | 10,31371944 | 10,08783583 | 4,410408591 | 1,834278466 | 8,990154331 | 8,510931 |
| Ptgds | 13,71650616 | 8,481750388 | 11,66874807 | 12,24789169 | 2,031057401 | 2,031057401 | 9,31719758 | 8,49917267 |
| Dpysl2 | 11,7704983 | 11,72774638 | 9,010470473 | 11,86467162 | 2,62168639 | 2,62168639 | 9,831564554 | 8,49261773 |
| Nfe2l2 | 7,030512075 | 7,332639092 | 9,116608734 | 8,270297956 | 17,48209491 | 2,792368617 | 7,419386746 | 8,491986875 |
| Auts2 | 9,317596294 | 6,199841677 | 6,357363767 | 5,048533364 | 11,13859947 | 11,42094618 | 9,9046996 | 8,48394005 |
| Snx3 | 9,003523496 | 8,41645822 | 6,792701173 | 9,366202853 | 16,75022767 | 1,108401179 | 7,922983802 | 8,480071199 |
| Rpl13 | 10,54810189 | 7,642010466 | 9,329359082 | 10,68034271 | 1,108306787 | 10,24449854 | 9,807082281 | 8,479957393 |
| Ogt | 9,700728149 | 10,55287634 | 10,40047765 | 6,004110303 | 1,267089861 | 11,35018522 | 10,07412108 | 8,478512658 |
| Pds5a | 2,530348906 | 8,342747111 | 10,23511904 | 4,361033521 | 13,0719422 | 11,72174827 | 9,0656867 | 8,475517965 |
| Gm51810 | 8,352945934 | 6,018153845 | 6,275568658 | 9,988477946 | 11,49906422 | 8,698875875 | 8,40213298 | 8,462174208 |
| Stag1 | 8,010799759 | 8,537770335 | 8,250543836 | 10,1064542 | 11,5908097 | 5,35959925 | 7,34539817 | 8,457339322 |
| Mat2a | 10,91395119 | 9,473323198 | 9,175678644 | 9,549722807 | 3,553565169 | 7,952121919 | 8,472824988 | 8,441598274 |
| Klhl9 | 3,467894302 | 9,885131668 | 9,026558977 | 5,315717119 | 18,99950539 | 3,315761863 | 9,077944577 | 8,44121627 |
| Adgrl1 | 8,6095286 | 7,357105604 | 7,369434961 | 5,156776225 | 14,00359941 | 7,675764409 | 8,890157312 | 8,437480931 |
| Fermt2 | 7,918538865 | 7,868698421 | 9,10841795 | 4,152205388 | 16,37150848 | 3,034363227 | 10,46893978 | 8,417524587 |
| Acat1 | 11,02285274 | 1,142769105 | 7,194673058 | 11,42533756 | 8,47498614 | 10,6193579 | 8,950484296 | 8,404351543 |
| Celf1 | 5,936207702 | 5,482845015 | 6,650742887 | 12,85455944 | 15,21747843 | 5,861040642 | 6,815140053 | 8,402573453 |
| Rpl36 | 9,072174223 | 7,309489209 | 7,612120908 | 9,978931417 | 15,86489323 | 1,064811775 | 7,801942207 | 8,386337566 |
| Rtn3 | 11,39585096 | 10,94194494 | 9,609845728 | 7,812163613 | 1,66406746 | 6,978633411 | 10,28611927 | 8,38408934 |
| Rplp1 | 9,982741661 | 7,383601909 | 8,040821521 | 8,007458239 | 16,16958944 | 0,727984366 | 8,309404059 | 8,374514456 |
| OC11856734 | 8,411968434 | 7,57677944 | 7,596879962 | 9,727470711 | 12,42633443 | 4,736806174 | 8,129564127 | 8,372257611 |
| Sat1 | 7,941967999 | 12,60214976 | 9,199385244 | 10,08773792 | 1,987648247 | 7,545657518 | 9,219053612 | 8,369085758 |

| | | | | | | | | |
|---|---|---|---|---|---|---|---|---|
| Grm3 | 8,958904578 | 11,02708659 | 9,317288974 | 9,606761203 | 2,25529734 | 8,805999299 | 8,599441847 | 8,367254262 |
| Tra2a | 9,109829119 | 7,376794572 | 8,657357565 | 11,85952351 | 1,489532843 | 12,41690342 | 7,604229463 | 8,359167213 |
| Ywhae | 9,353133239 | 9,779873744 | 11,38490454 | 4,205978692 | 4,577018671 | 9,205965103 | 10,00603076 | 8,358986394 |
| Set | 8,88878593 | 10,01656243 | 8,019075148 | 11,59107138 | 10,92498395 | 1,559445885 | 7,483271102 | 8,35474226 |
| H3f3b | 8,971681382 | 8,603732372 | 10,41510573 | 10,09566255 | 1,532925907 | 10,99057167 | 7,862704155 | 8,353197681 |
| Hmgb1 | 12,55683192 | 9,536952031 | 8,953404776 | 7,165867781 | 1,605850572 | 8,987414583 | 9,654390713 | 8,35153034 |
| Gm32262 | 8,395938975 | 9,233438886 | 9,622668135 | 9,709574964 | 3,77027346 | 12,06760007 | 5,635799958 | 8,347899207 |
| Atxn7l3b | 3,615687074 | 7,292019281 | 8,263752637 | 10,83802493 | 9,916822765 | 11,4319179 | 7,072799372 | 8,347289138 |
| Rorb | 8,494844266 | 9,846187933 | 8,631853612 | 11,23489945 | 5,041868167 | 8,944884463 | 6,188758621 | 8,34047093 |
| Gm51670 | 4,357721832 | 7,538640713 | 9,015688883 | 7,69631916 | 10,44306036 | 10,97334159 | 8,349033079 | 8,339115088 |
| Rplp2 | 9,676726899 | 7,179219583 | 7,616460991 | 9,975958644 | 4,237855745 | 10,95634381 | 8,711701961 | 8,336323948 |
| Tnik | 10,5777512 | 9,104276111 | 10,45845691 | 7,877752843 | 11,53612101 | 0,660808145 | 8,103892918 | 8,331294162 |
| Gramd1b | 8,615628361 | 8,98776565 | 6,893013018 | 8,860791121 | 12,0856336 | 4,289582249 | 8,520664119 | 8,321868302 |
| Arhgef12 | 7,803486493 | 8,155073909 | 9,363031944 | 6,985706949 | 9,337154643 | 7,270033319 | 9,318655999 | 8,319020465 |
| Rhob | 11,05477992 | 3,906745023 | 8,063871389 | 10,83560109 | 16,28094521 | 1,729701999 | 6,338732191 | 8,315768117 |
| Ppp3ca | 9,381911429 | 9,812046492 | 7,834978704 | 9,74714568 | 1,564446278 | 10,71211435 | 9,118149743 | 8,310113239 |
| Tmed10 | 10,37157723 | 11,54581037 | 8,286617681 | 12,263783 | 2,214521946 | 4,628378957 | 8,853088141 | 8,309111046 |
| Trf | 12,62806893 | 8,959671761 | 10,18070066 | 8,830682565 | 5,259116724 | 2,699163806 | 9,597541557 | 8,30784943 |
| Brwd1 | 6,910187181 | 7,175382749 | 9,353019013 | 5,93645932 | 9,363415513 | 11,35165069 | 8,059866664 | 8,307140161 |
| Notch2 | 8,966592875 | 7,192446803 | 7,340266763 | 8,313817924 | 7,075590114 | 11,64584647 | 7,58437091 | 8,302704552 |
| Gcc2 | 11,29373356 | 2,422203062 | 7,230067259 | 10,79537547 | 7,388624026 | 11,67448585 | 7,30178741 | 8,300896663 |
| Ctsb | 9,770898392 | 9,270745572 | 9,530238797 | 7,992637733 | 4,266136204 | 8,775385719 | 8,483562833 | 8,298515036 |
| Fcho2 | 1,752292603 | 10,25693221 | 8,373973034 | 6,828455873 | 15,3759884 | 7,601797885 | 7,898494798 | 8,298276401 |
| Rps25 | 10,14073314 | 7,387499493 | 8,164645699 | 9,337336443 | 13,58704966 | 1,191311275 | 8,235214806 | 8,291970074 |
| Spry2 | 10,24819889 | 7,257190794 | 7,824562013 | 11,0724995 | 1,815659084 | 12,91744215 | 6,860980282 | 8,285218959 |
| Thoc2 | 5,831989916 | 9,280919484 | 9,188481479 | 9,69139987 | 1,838805483 | 13,06737608 | 9,096424899 | 8,285056744 |
| Scrg1 | 5,198521922 | 8,725274468 | 8,800354642 | 4,319303106 | 16,03623882 | 5,116898164 | 9,743430655 | 8,277145969 |
| Slc25a4 | 3,070374349 | 10,87783351 | 8,297277848 | 9,766649044 | 5,827164008 | 12,35739779 | 7,699939536 | 8,270948013 |
| Tmbim6 | 10,03182889 | 10,29270008 | 10,51489588 | 6,373095268 | 0,835522575 | 11,14205971 | 8,700974192 | 8,270153799 |
| Enah | 8,036236206 | 9,861961248 | 9,004751736 | 9,174902075 | 3,652577123 | 8,844853992 | 9,293790625 | 8,267010429 |
| Rps24 | 9,976428014 | 8,641905218 | 8,094576981 | 11,48104095 | 6,10669677 | 5,549645317 | 8,001400764 | 8,264527717 |
| Luc7l | 9,460534124 | 7,070585187 | 8,217950879 | 9,526439918 | 14,00549767 | 0,513346119 | 9,042100965 | 8,262350695 |
| Mobp | 12,09378264 | 7,337796691 | 8,93582592 | 12,04629966 | 7,085124678 | 1,960714683 | 8,338784703 | 8,256904139 |

| | | | | | | | | |
|---|---|---|---|---|---|---|---|---|
| Timp3 | 9,811737584 | 9,351561919 | 8,864439562 | 8,119747846 | 1,439606908 | 10,73191808 | 9,42612278 | 8,249304954 |
| Celf2 | 8,295139935 | 11,17009647 | 9,583628371 | 5,461167345 | 5,474916767 | 8,319725123 | 9,398116017 | 8,243255719 |
| Slco1c1 | 9,328807507 | 9,884926165 | 9,881825207 | 3,413382909 | 3,818070827 | 11,34467731 | 10,01348027 | 8,240738599 |
| Kmt2e | 11,33851188 | 5,483794765 | 9,152456102 | 7,301418783 | 4,115363717 | 13,15554659 | 7,052603779 | 8,228527945 |
| Gpr37 | 4,055484475 | 11,00907221 | 8,191570605 | 10,54032406 | 2,630779555 | 12,57899368 | 8,556171643 | 8,223199461 |
| Pmm1 | 8,499621956 | 7,456092834 | 7,302799254 | 2,852902116 | 16,02793975 | 4,895698963 | 10,48099356 | 8,216578348 |
| Rora | 8,063865231 | 10,71603674 | 8,99573599 | 5,883327762 | 6,077135009 | 9,918135247 | 7,817871144 | 8,210301017 |
| Rpl7 | 10,02290506 | 9,328754688 | 8,446994394 | 11,2703947 | 0,810294055 | 9,179728741 | 8,388718472 | 8,206827158 |
| Npc1 | 11,83889072 | 9,862750398 | 8,321942253 | 8,611988361 | 7,38105808 | 0,85738912 | 10,56113089 | 8,205021404 |
| Eif5 | 7,409849413 | 11,20773974 | 9,437600135 | 6,157640057 | 3,687615249 | 10,60863574 | 8,893061824 | 8,200306024 |
| Gnai2 | 9,914148278 | 5,490172922 | 8,541316597 | 7,849565138 | 9,304341812 | 9,896673028 | 6,370366231 | 8,195226287 |
| Fut9 | 9,526229944 | 11,32371727 | 8,490275688 | 10,0558328 | 4,575272924 | 4,424045725 | 8,867297237 | 8,180381655 |
| Cnn3 | 9,697928218 | 8,83892443 | 9,372650113 | 2,397195463 | 10,29552118 | 7,493393923 | 9,097221363 | 8,170404955 |
| S100a16 | 8,698475232 | 8,131989728 | 7,379142736 | 2,093451919 | 16,76259087 | 3,639908723 | 10,46885973 | 8,167774134 |
| Rab2a | 4,555679999 | 10,73151404 | 8,479240686 | 9,707051406 | 9,99060878 | 4,12473628 | 9,540175897 | 8,161286726 |
| Ankrd11 | 11,52068819 | 9,422337858 | 9,500088649 | 11,36308758 | 3,684464139 | 8,061071556 | 3,567853053 | 8,159941575 |
| Ndufc2 | 2,233278173 | 3,74600622 | 7,511788138 | 10,36144784 | 18,22131646 | 6,201860643 | 8,823472838 | 8,15702433 |
| Tnrc6a | 9,65145047 | 7,090443667 | 8,838414935 | 11,22471318 | 1,695943262 | 11,61604463 | 6,969349163 | 8,155194187 |
| Golgb1 | 10,2400363 | 7,773098077 | 9,40148111 | 8,583993509 | 4,120507155 | 8,75117348 | 8,215215288 | 8,155072132 |
| Trps1 | 11,81075051 | 7,517837003 | 9,511693776 | 10,69958267 | 1,568473898 | 7,387646893 | 8,581776742 | 8,153965927 |
| Taok1 | 10,9285434 | 9,596551251 | 8,033715831 | 9,453197605 | 2,160772856 | 7,316043368 | 9,572797966 | 8,151660325 |
| Rps2 | 8,480682049 | 8,647701909 | 9,96300431 | 5,381634318 | 16,35523547 | 0,414501793 | 7,79648418 | 8,148463433 |
| Uba1 | 10,97295812 | 1,475464082 | 7,60653922 | 10,94670983 | 17,77961385 | 1,475464082 | 6,680195631 | 8,133849258 |
| Calr | 10,31043661 | 8,161486561 | 9,369742703 | 8,582517646 | 4,19326395 | 6,623910418 | 9,62992593 | 8,124469117 |
| Capza2 | 6,321200888 | 8,634582103 | 9,295242814 | 10,26334813 | 9,536943799 | 4,499387905 | 8,190099861 | 8,105829357 |
| Septin4 | 13,41236239 | 2,704143434 | 10,22425655 | 10,92549947 | 5,376222275 | 4,095088912 | 9,996779126 | 8,104907451 |
| Frmd4a | 9,582669394 | 7,987817184 | 7,047454393 | 7,186231974 | 5,267521186 | 9,067239284 | 10,42977041 | 8,081243403 |
| Ptp4a2 | 9,321159502 | 7,756937232 | 9,625563439 | 6,893972991 | 12,27730636 | 1,63241143 | 9,019634305 | 8,075283608 |
| Cdc5l | 9,743519281 | 9,211474571 | 9,307794988 | 10,01162903 | 0,860217378 | 9,189840267 | 8,183899968 | 8,072625069 |
| Mbnl2 | 11,10177395 | 10,01192057 | 9,652329754 | 10,84174486 | 4,602078327 | 3,828337395 | 6,459996096 | 8,071168707 |
| Chmp2a | 10,87714558 | 6,957093868 | 8,478321158 | 10,31136518 | 0,8708253 | 10,23073223 | 8,769053283 | 8,070648085 |
| Rbm26 | 8,386781394 | 9,677749168 | 8,112609689 | 11,21038748 | 4,213866658 | 7,334361164 | 7,47329267 | 8,058435461 |
| Ndufa2 | 9,688352528 | 9,638496553 | 7,509483244 | 8,834411721 | 1,216988273 | 11,22273665 | 8,18797923 | 8,042635457 |

| | | | | | | | | |
|---|---|---|---|---|---|---|---|---|
| Spag9 | 2,003577264 | 11,79365724 | 7,745301077 | 11,92602668 | 2,003577264 | 12,02047128 | 8,791332203 | 8,040563287 |
| Kmt2c | 5,955297095 | 10,77438389 | 8,699694534 | 9,194032833 | 4,497100369 | 10,1848977 | 6,976473331 | 8,040268536 |
| R3hdm1 | 9,825984735 | 9,182325382 | 8,72772266 | 9,460925762 | 1,317723203 | 11,50077734 | 6,220946049 | 8,033772161 |
| Map4k4 | 5,307789405 | 9,308674595 | 7,780480301 | 10,51323465 | 13,1392905 | 2,265329258 | 7,909523834 | 8,032046078 |
| Ube2d3 | 9,293776338 | 5,226600531 | 7,918593366 | 10,8967296 | 4,481405224 | 11,09994215 | 7,276886169 | 8,027704768 |
| Emsy | 7,513718545 | 9,212404443 | 9,334233048 | 3,907392271 | 3,660211858 | 12,79134893 | 9,685812136 | 8,015017319 |
| Slc1a4 | 9,711932413 | 10,02500065 | 7,999939205 | 7,781361603 | 1,221432756 | 9,956058544 | 9,328227953 | 8,003421874 |
| Ndufb11 | 9,263012015 | 8,858564309 | 9,353892713 | 3,182658756 | 15,69518594 | 0,712753117 | 8,949146466 | 8,00217333 |
| Hmgcs1 | 6,348149622 | 7,218930779 | 8,513323162 | 5,338049918 | 15,82397601 | 2,976400178 | 9,788394263 | 8,001031991 |
| Kif5b | 9,660935885 | 5,641470355 | 9,032226362 | 9,395206436 | 4,05139393 | 12,07062171 | 6,135592312 | 7,998206713 |
| Arap2 | 7,655091865 | 10,4203898 | 9,143427411 | 4,769472844 | 13,96343939 | 0,620172804 | 9,414896107 | 7,998127175 |
| Ubb-ps | 9,884968663 | 9,583212673 | 11,19825291 | 1,427193386 | 4,166307628 | 9,598316579 | 10,10954166 | 7,99539907 |
| Mycbp2 | 9,781672539 | 9,137020947 | 9,003145298 | 4,934904979 | 3,741802279 | 10,60253002 | 8,708584354 | 7,987094345 |
| Eea1 | 11,11557723 | 9,319998098 | 6,30885499 | 9,065090638 | 10,0917315 | 1,818919894 | 8,189096256 | 7,987038372 |
| Rn7sk | 9,239862345 | 11,77317036 | 8,355266584 | 10,94819511 | 2,338370468 | 4,072295487 | 9,172879042 | 7,985719914 |
| Oxct1 | 9,720046252 | 8,522241542 | 6,421150531 | 9,933248656 | 1,006708072 | 10,89798567 | 9,397850357 | 7,98560444 |
| Spcs1 | 9,397565071 | 8,841180642 | 9,089406284 | 4,538161155 | 16,73259297 | 0,308637766 | 6,98515027 | 7,984670594 |
| Ppargc1a | 10,25270854 | 9,021824791 | 7,223070619 | 4,321814593 | 11,98775377 | 3,098244461 | 9,97737912 | 7,983256556 |
| Pcdh10 | 5,906532455 | 11,30107817 | 5,668161219 | 10,29733741 | 1,527815029 | 12,3769892 | 8,679454703 | 7,965338312 |
| Cox4i1 | 7,11290923 | 8,463190499 | 10,08422791 | 3,082784319 | 16,95790292 | 0,7836931 | 9,264459896 | 7,96416684 |
| OC11548862 | 11,00003485 | 6,859853217 | 6,81446484 | 9,041476976 | 3,238997056 | 10,14500247 | 8,633818625 | 7,961949719 |
| Aqp4 | 7,128476933 | 12,3222901 | 9,557237777 | 4,294965785 | 1,491864202 | 10,94491294 | 9,93753514 | 7,953897553 |
| Itprid2 | 2,050638628 | 2,050638628 | 6,996216991 | 9,747714971 | 12,12289358 | 14,31580894 | 8,303472562 | 7,941054899 |
| Hmgn2 | 11,88773696 | 10,78455169 | 8,30596421 | 11,29244974 | 2,15964449 | 2,15964449 | 8,982294283 | 7,938897981 |
| Kidins220 | 3,735021366 | 6,472689832 | 6,183412942 | 7,655993229 | 11,74102364 | 10,15236442 | 9,5985647 | 7,934152876 |
| Cdc73 | 2,362272779 | 8,190535472 | 7,541462209 | 10,06624119 | 18,30825614 | 2,362272779 | 6,692203414 | 7,931891999 |
| Smarca4 | 9,220664629 | 6,753454458 | 8,123835404 | 10,92191835 | 12,19809761 | 1,240735878 | 7,039098424 | 7,928257822 |
| Grin2c | 9,585796226 | 9,105824823 | 7,285499821 | 10,15800031 | 1,408389619 | 9,62018367 | 8,300509954 | 7,923457775 |
| Slc38a3 | 7,721058046 | 9,61134229 | 8,289398883 | 7,095198827 | 4,526777539 | 10,26837827 | 7,87118807 | 7,91190599 |
| Gtf2i | 9,323507016 | 1,266517155 | 8,350163815 | 9,657071824 | 18,49478475 | 1,266517155 | 7,021499942 | 7,911437379 |
| Gm52669 | 4,245378818 | 6,395922322 | 10,17193246 | 9,154841875 | 11,52926029 | 9,328006442 | 4,535419185 | 7,908680199 |
| Ttc3 | 7,818946791 | 9,372310683 | 7,940737172 | 10,33711895 | 11,45675355 | 1,709650923 | 6,724092612 | 7,908515811 |
| Zfp318 | 8,18551236 | 8,331771785 | 9,205462273 | 5,080577696 | 14,92014435 | 0,357716412 | 9,242569571 | 7,903393493 |

| | | | | | | | | |
|---|---|---|---|---|---|---|---|---|
| Sorbs1 | 10,0891166 | 11,94430775 | 6,858703946 | 11,41269125 | 4,692625342 | 2,102301855 | 8,202727045 | 7,900353399 |
| Zfp36l1 | 9,726370135 | 5,92706026 | 8,809861807 | 9,435467324 | 2,344126453 | 9,713602561 | 9,313209074 | 7,895671088 |
| Rtn4 | 9,059759264 | 8,246616673 | 10,11185743 | 8,712693943 | 6,741671206 | 4,263530027 | 8,11024592 | 7,892339209 |
| 820431F20R | 8,279262845 | 9,218228449 | 9,933133093 | 6,696632987 | 3,880643798 | 8,874727588 | 8,324976033 | 7,886800685 |
| Rps9 | 5,684101568 | 8,36894504 | 7,820850231 | 9,475229663 | 3,648324071 | 11,98209436 | 8,158537035 | 7,876868852 |
| Arxes2 | 1,279683004 | 10,65214033 | 9,501873736 | 2,603879235 | 18,79626623 | 3,069337805 | 9,220769878 | 7,874850032 |
| Egr1 | 12,9295366 | 9,233938797 | 8,173689416 | 12,01635538 | 1,850562787 | 1,850562787 | 9,049461708 | 7,872015354 |
| Rplp0 | 10,0049453 | 9,732719088 | 8,518681138 | 10,21182889 | 5,420766541 | 2,103735476 | 9,096323812 | 7,869857177 |
| Gabbr1 | 7,555057034 | 6,630071785 | 7,797860152 | 7,133658823 | 13,76448607 | 4,887207777 | 7,313808455 | 7,868878585 |
| Ube3a | 12,24708107 | 6,609809144 | 7,825380172 | 6,46277717 | 1,516423659 | 10,50157246 | 9,908447015 | 7,867355813 |
| Eif3a | 10,80932026 | 9,330715335 | 8,551814803 | 10,38501736 | 2,395760791 | 4,192064139 | 9,304160441 | 7,852693305 |
| Ubtd2 | 8,294059282 | 6,516967468 | 6,160067421 | 8,938977171 | 5,693168339 | 10,98083049 | 8,38002522 | 7,852013627 |
| Gnb1 | 1,465784804 | 9,915153415 | 10,00392244 | 2,416278928 | 12,04475246 | 10,70434549 | 8,358829047 | 7,844152369 |
| Uqcrq | 9,677856426 | 7,490673191 | 9,599816054 | 4,353315026 | 14,86485578 | 0,540759758 | 8,356433532 | 7,840529967 |
| Atp5j2 | 9,316621728 | 4,225497073 | 7,971850393 | 7,817871972 | 16,40270884 | 1,318474564 | 7,776096162 | 7,832731533 |
| Zeb1 | 7,767339658 | 9,666327076 | 7,875141111 | 9,719635788 | 1,027532447 | 11,8268867 | 6,920165056 | 7,829003977 |
| Klf7 | 10,51934015 | 7,650642016 | 7,89670481 | 8,250178612 | 1,098036948 | 11,66640457 | 7,687628204 | 7,824133617 |
| Uba52 | 8,82661686 | 7,592851184 | 7,144874634 | 6,994409669 | 3,396127751 | 11,30475436 | 9,497154941 | 7,822398485 |
| Rpl18a | 10,41734937 | 7,851109483 | 8,347201543 | 5,314054103 | 3,959536782 | 9,240287991 | 9,599810719 | 7,81847857 |
| Csnk1a1 | 8,764649824 | 10,96294682 | 8,814266235 | 9,952954198 | 1,896124701 | 5,707481658 | 8,627374127 | 7,817971081 |
| OC11548692 | 0,593379555 | 9,583634357 | 10,28000229 | 3,28737308 | 13,84488514 | 10,45982589 | 6,639421496 | 7,812645973 |
| Chd6 | 10,34107751 | 2,98749979 | 8,450462873 | 10,04228937 | 15,12737029 | 1,961033147 | 5,771393401 | 7,811589484 |
| Acsl6 | 8,07812418 | 12,48631191 | 8,94531358 | 5,784264491 | 5,225663969 | 3,483689082 | 10,65657915 | 7,808563766 |
| Limch1 | 9,954436284 | 5,962424356 | 8,310502059 | 9,108082586 | 8,388007954 | 4,32434008 | 8,606143718 | 7,807705291 |
| Mindy2 | 7,383704066 | 8,205894318 | 6,446464268 | 10,25125475 | 1,439258975 | 11,87094887 | 9,051212538 | 7,80696254 |
| Arl6ip1 | 11,1662656 | 10,64365403 | 6,848882186 | 10,39750995 | 2,046232529 | 4,456653024 | 9,019733435 | 7,796990109 |
| Trim2 | 8,673120967 | 9,252984271 | 10,13480054 | 7,41928511 | 9,43470232 | 1,092032538 | 8,570792262 | 7,796816858 |
| Chst2 | 3,203004805 | 10,61192115 | 8,12080512 | 6,373547101 | 4,343327284 | 12,53832399 | 9,380442324 | 7,795910253 |
| Tsc22d3 | 7,328732978 | 10,20720755 | 8,5031089 | 7,3492207 | 4,357338498 | 8,479681641 | 8,334094169 | 7,794197776 |
| Kat6b | 5,230329889 | 7,359524171 | 9,51779732 | 9,387336562 | 5,309411863 | 9,370192842 | 8,380053554 | 7,793520886 |
| Phactr3 | 9,370902382 | 10,46085812 | 9,016979876 | 4,605742873 | 3,15598744 | 7,270846561 | 10,65113851 | 7,790350823 |
| Nufip2 | 8,690549759 | 8,187076823 | 9,242171034 | 11,16908357 | 0,917477843 | 11,18786819 | 5,131529087 | 7,789393759 |
| AY036118 | 9,480837224 | 6,96550778 | 4,878284275 | 8,720464153 | 6,662274507 | 10,55852435 | 7,17904803 | 7,777848617 |

| | | | | | | | | |
|---|---|---|---|---|---|---|---|---|
| Gatm | 10,09829224 | 9,884076196 | 9,480905753 | 9,904105235 | 2,97889839 | 2,97889839 | 9,107536379 | 7,776101797 |
| Tinagl1 | 7,956069172 | 6,938069254 | 6,25006479 | 9,049378911 | 9,891516298 | 6,982798988 | 7,346024752 | 7,773417452 |
| Il33 | 11,54524952 | 10,19455028 | 10,408926 | 7,92744557 | 2,223018759 | 4,017133804 | 8,085311578 | 7,771662216 |
| Selenos | 10,89192886 | 10,16124545 | 8,053425064 | 10,04858855 | 4,548287887 | 1,999728005 | 8,662405173 | 7,76651557 |
| Gnaq | 5,817045994 | 9,804131497 | 7,518838969 | 11,86472197 | 1,351159451 | 11,23269057 | 6,759756046 | 7,764049213 |
| Chpt1 | 5,627385933 | 8,200184212 | 7,215713377 | 9,533425573 | 2,540200509 | 11,93975799 | 9,283417904 | 7,762869357 |
| Cdc42bpa | 9,502011155 | 6,863172403 | 8,680438537 | 8,255099985 | 2,188222265 | 10,01428118 | 8,820309921 | 7,760505063 |
| Msi2 | 10,68725967 | 9,546539163 | 7,612127193 | 10,59496515 | 6,504247212 | 1,142801333 | 8,174983806 | 7,751846219 |
| Vcp | 9,31360036 | 7,646975796 | 8,031988561 | 8,013496551 | 2,125906757 | 10,70870445 | 8,402577149 | 7,74903566 |
| Plxnb1 | 10,90382453 | 9,031716826 | 8,525365719 | 9,087004703 | 2,89457794 | 3,923333019 | 9,848572862 | 7,744913657 |
| Cldn10 | 2,153218887 | 11,71587485 | 8,920902245 | 7,657762774 | 2,153218887 | 11,83607337 | 9,75841475 | 7,742209394 |
| Psat1 | 8,09937577 | 6,386353673 | 9,549536832 | 4,367316732 | 14,13109386 | 2,937025142 | 8,708199378 | 7,739843056 |
| Aco1 | 5,969148339 | 6,368724373 | 4,728863051 | 11,0501567 | 12,17465231 | 5,609224342 | 8,08961013 | 7,712911321 |
| Dcun1d5 | 6,261285503 | 8,725440604 | 6,122935153 | 11,4191088 | 4,542437775 | 8,693096399 | 8,214598833 | 7,711271867 |
| Arl13b | 9,793323875 | 7,42795662 | 8,910372375 | 9,584766527 | 15,02033717 | 0,427504505 | 2,765955607 | 7,704316669 |
| Enpp2 | 11,12842484 | 8,674530746 | 9,875598067 | 10,8193528 | 2,424231145 | 2,424231145 | 8,574318031 | 7,702955253 |
| Trpm3 | 4,025010215 | 10,03426245 | 7,304996441 | 6,9863437 | 5,186509355 | 10,36259803 | 10,01170798 | 7,701632596 |
| Rpl41 | 11,58182413 | 8,544461784 | 8,057041274 | 10,2654246 | 1,683564026 | 5,908542772 | 7,860938224 | 7,700256686 |
| Hnrnpk | 5,523190312 | 10,38078756 | 8,055967681 | 10,5133837 | 3,951129656 | 5,516986796 | 9,942516634 | 7,697708905 |
| Atp6v1f | 8,749422878 | 2,979734711 | 8,2586789 | 3,784878295 | 17,17222544 | 2,503432992 | 10,41630586 | 7,694954153 |
| Tppp3 | 5,850748588 | 3,308212299 | 8,114966007 | 9,521469027 | 17,2161051 | 0,700925165 | 9,129054556 | 7,691640106 |
| Id4 | 10,1279584 | 8,62378402 | 7,951000253 | 6,770640582 | 1,59556374 | 9,112874428 | 9,632309385 | 7,687732973 |
| Asph | 5,851743472 | 8,311684114 | 8,565747222 | 5,742420512 | 5,740754546 | 9,749704538 | 9,850119764 | 7,687453453 |
| Elovl2 | 2,861883622 | 5,463403923 | 5,994460254 | 7,066805324 | 19,41444838 | 4,272719956 | 8,65647832 | 7,675742826 |
| Hp1bp3 | 6,486442227 | 10,61875142 | 8,781921231 | 10,79737947 | 1,892787747 | 7,695532262 | 7,403229256 | 7,668006231 |
| Rpl36a | 10,13079102 | 7,275625841 | 7,908063409 | 11,41406494 | 1,265022338 | 7,200580953 | 8,419294575 | 7,659063297 |
| Myo9a | 7,610910559 | 2,560077683 | 7,584899734 | 7,378719139 | 17,86937682 | 2,560077683 | 8,003833244 | 7,652556408 |
| Strbp | 9,006442742 | 10,04813477 | 8,748167602 | 9,974603418 | 6,232437444 | 3,733308781 | 5,804289995 | 7,649626392 |
| Sptbn1 | 6,409906065 | 4,962236008 | 9,205335461 | 9,741587751 | 6,294723728 | 9,02923467 | 7,844854511 | 7,641125456 |
| Dhrs3 | 1,690096442 | 10,05753487 | 7,828840926 | 9,889152088 | 1,690096442 | 13,64693266 | 8,679834397 | 7,640355403 |
| Plekhb1 | 11,15707659 | 2,032443308 | 10,30873142 | 7,823640606 | 2,032443308 | 10,45305408 | 9,652220057 | 7,637087052 |
| Pou3f3 | 9,662416082 | 9,348066631 | 7,686641829 | 11,01427089 | 1,638227276 | 6,272949973 | 7,780865235 | 7,62906256 |
| OC11856850 | 4,656397048 | 6,5103611 | 7,329491899 | 5,171480476 | 12,62874435 | 10,95279566 | 6,093669612 | 7,620420021 |

| | | | | | | | | |
|---|---|---|---|---|---|---|---|---|
| Ndufa4 | 7,528862255 | 8,630634382 | 9,549009346 | 1,554770072 | 4,015275446 | 12,028545 | 9,982243121 | 7,612762804 |
| Rpl8 | 11,88053852 | 8,300458422 | 8,25336937 | 9,649255496 | 4,780303107 | 2,241595135 | 8,117564966 | 7,60329786 |
| Tmco1 | 9,403501152 | 11,4854934 | 8,122072506 | 9,214367747 | 1,93841612 | 3,734283752 | 9,299044198 | 7,599596982 |
| Rpl10 | 6,356010387 | 10,26240005 | 7,941353244 | 7,506028774 | 5,362822131 | 6,77016525 | 8,946984927 | 7,592252109 |
| Ywhah | 4,132339875 | 9,055933583 | 7,184971564 | 9,178347408 | 3,654173873 | 11,04537434 | 8,88491946 | 7,590865729 |
| Ube2b | 3,944051152 | 7,599304789 | 8,980246237 | 10,50811518 | 3,463427903 | 10,27840762 | 8,348092759 | 7,58880652 |
| Slc4a8 | 1,150833009 | 10,89297339 | 4,794875097 | 10,66955035 | 15,71527746 | 1,150833009 | 8,742955342 | 7,588185379 |
| Eif2s3y | 5,562821119 | 9,286026739 | 9,514075393 | 10,4266525 | 1,324929921 | 10,7646162 | 6,222064454 | 7,58588376 |
| Fasn | 10,08087067 | 8,077676831 | 8,273624488 | 4,629101847 | 1,008705566 | 11,40594882 | 9,597635669 | 7,581937699 |
| Tor1aip1 | 6,022811916 | 5,066596716 | 7,833669115 | 11,03182609 | 9,59107047 | 8,565376645 | 4,937402727 | 7,578393382 |
| Cab39 | 10,38970068 | 0,788585383 | 8,914247548 | 5,668647129 | 13,65589685 | 7,739659379 | 5,885559689 | 7,577470951 |
| Slc38a2 | 8,466848834 | 7,68044892 | 7,185081093 | 10,87263066 | 4,799945975 | 6,238170363 | 7,769483751 | 7,573229943 |
| Qdpr | 11,96305274 | 9,357809508 | 9,719673072 | 8,106639291 | 1,636887842 | 3,42513922 | 8,799269235 | 7,572638701 |
| Sdc4 | 4,312869151 | 10,38738911 | 8,850543212 | 7,178248168 | 10,44645002 | 3,507455486 | 8,285553601 | 7,56692982 |
| Nckap1 | 9,511706401 | 7,433647831 | 8,818993006 | 6,260451584 | 0,731880788 | 11,62284235 | 8,568353052 | 7,563982144 |
| Ccar1 | 12,09421766 | 2,423065128 | 7,94729283 | 8,003961198 | 1,636685994 | 12,67491894 | 8,162318015 | 7,563208537 |
| Uqcrc1 | 5,574916153 | 7,855536688 | 8,689199473 | 7,17850612 | 16,00792427 | 0,14898281 | 7,48541558 | 7,56292587 |
| Atp5g3 | 6,034583755 | 8,761944995 | 6,252874904 | 8,253709195 | 5,067605705 | 10,19744763 | 8,352218304 | 7,560054927 |
| Canx | 8,726853303 | 11,21728686 | 9,203032955 | 10,18250423 | 2,492827678 | 2,492827678 | 8,585848895 | 7,557311658 |
| Itgav | 9,054127552 | 10,84552108 | 6,31676317 | 10,38032125 | 1,845224028 | 6,005900191 | 8,426454986 | 7,553473179 |
| Tbc1d19 | 6,942496311 | 1,429409322 | 8,81675934 | 11,07202801 | 19,41337416 | 2,536113205 | 2,632728118 | 7,548986925 |
| OC11856831 | 7,421696911 | 7,026057632 | 5,271608373 | 8,771482649 | 9,625880651 | 8,63503479 | 6,037661477 | 7,541346069 |
| Sfxn5 | 8,40734684 | 8,850878524 | 8,762723065 | 5,555449252 | 4,288846472 | 8,131384919 | 8,791126039 | 7,541107873 |
| Gm51968 | 6,148860836 | 9,383392791 | 7,099342959 | 11,15065576 | 2,273393931 | 10,52949164 | 6,160415487 | 7,535079058 |
| Rbbp6 | 10,34686228 | 5,934463775 | 7,551714385 | 10,24984625 | 1,901526186 | 10,30906129 | 6,434418991 | 7,532556165 |
| Rpl23a | 7,841143491 | 9,135320926 | 8,791034803 | 10,55280793 | 5,358181267 | 2,081388645 | 8,918524315 | 7,525485911 |
| Rps3a1 | 10,89815762 | 5,058394383 | 8,978488477 | 11,01004053 | 2,189070784 | 6,169145628 | 8,355980085 | 7,522753929 |
| Top1 | 8,634671428 | 8,187653455 | 7,886667434 | 6,688091068 | 3,417556887 | 10,16079062 | 7,674787481 | 7,521459767 |
| Slc3a2 | 2,622677018 | 11,1160344 | 8,827542804 | 9,276473343 | 6,310080995 | 5,541479792 | 8,949495967 | 7,520540617 |
| Gm35760 | 9,561568177 | 8,297729912 | 6,426334236 | 9,331146094 | 6,443129774 | 4,574534329 | 7,967197619 | 7,51452002 |
| Cd47 | 7,073530024 | 8,31675662 | 9,322566458 | 3,682165519 | 7,204968807 | 9,311286203 | 7,680925139 | 7,513171253 |
| Lhx2 | 10,44423832 | 8,49102685 | 5,583458944 | 9,350747277 | 6,896118819 | 3,100516164 | 8,720926868 | 7,51243332 |
| Rrbp1 | 10,02742211 | 9,551550971 | 7,921866592 | 7,477062466 | 2,052692474 | 8,086688323 | 7,437284625 | 7,507795365 |

| | | | | | | | | |
|---|---|---|---|---|---|---|---|---|
| Dtna | 8,178327769 | 9,642665251 | 9,253304061 | 5,289035564 | 1,141262243 | 10,89262694 | 8,147960098 | 7,506454561 |
| Dnajb6 | 8,192081191 | 8,322801532 | 8,153423564 | 6,013655549 | 9,935511886 | 3,847719036 | 8,056366358 | 7,503079874 |
| Atp5j | 9,672771841 | 9,373053872 | 9,321717696 | 5,512075043 | 4,243970778 | 5,727165008 | 8,663971004 | 7,502103606 |
| Xrn1 | 4,680073746 | 2,580495295 | 4,730527286 | 9,957109699 | 19,69050507 | 4,499191085 | 6,373885773 | 7,501683993 |
| Hmgn3 | 8,522762158 | 11,07174034 | 8,860188325 | 11,02455757 | 4,68742192 | 2,009639865 | 6,224944907 | 7,485893584 |
| Thra | 1,558007727 | 8,967776261 | 4,977946899 | 8,770373793 | 8,035039796 | 11,30624576 | 8,723279192 | 7,476952776 |
| Rpl9 | 8,934281105 | 9,257055512 | 7,845668835 | 10,03695258 | 4,361813444 | 3,606765842 | 8,276512037 | 7,474149908 |
| Trim8 | 9,143051839 | 9,045560704 | 5,984755154 | 8,385531548 | 7,605122842 | 2,818507733 | 9,333129099 | 7,47366556 |

**Table S3.** Top 500 highest expressed genes among $A^+/G^+$ astrocytes.

| Gene | 2m-N2-$A^+/G^+$ | 2m-N3-$A^+/G^+$ | 2m-N5-$A^+/G^+$ | 2m-N6-$A^+/G^+$ | 18m-N2-$A^+/G^+$ | 18m-N3-$A^+/G^+$ | 18m-N4-$A^+/G^+$ | Mean expression |
|---|---|---|---|---|---|---|---|---|
| Malat1 | 15,76958452 | 16,04674635 | 17,23411095 | 16,72961315 | 16,24752454 | 15,57102476 | 16,81234367 | 16,34 |
| mt-Rnr2 | 14,86285186 | 15,02012127 | 16,23419729 | 16,12535695 | 14,09853484 | 14,24542868 | 16,34860694 | 15,28 |
| ATP6 | 15,15109525 | 14,90951168 | 15,66922349 | 15,72491014 | 14,20473774 | 13,40461841 | 15,29753537 | 14,91 |
| CYTB | 14,66321636 | 14,47643075 | 15,69726045 | 15,23439764 | 14,14787056 | 13,09667546 | 15,28977911 | 14,66 |
| Cst3 | 14,55316234 | 14,17009089 | 15,89393307 | 14,94156432 | 14,74402622 | 13,29696571 | 14,27435388 | 14,55 |
| COX1 | 14,29485181 | 14,28786527 | 15,24732946 | 15,31663605 | 13,81899569 | 13,52495613 | 15,22168027 | 14,53 |
| COX3 | 14,36193603 | 14,29951063 | 14,74894418 | 15,01177771 | 13,40956435 | 13,13097475 | 14,900241 | 14,27 |
| ND4 | 14,0126394 | 14,06015156 | 14,40878732 | 15,17691641 | 13,06822523 | 12,71226138 | 14,50999694 | 13,99 |
| mt-Rnr1 | 13,27155862 | 13,46176361 | 15,27055277 | 14,97386637 | 13,59019573 | 12,68423636 | 14,618721 | 13,98 |
| COX2 | 13,87264691 | 13,87465022 | 14,15836987 | 14,50008806 | 12,76981858 | 12,67519309 | 14,20531637 | 13,72 |
| Slc1a3 | 13,427645 | 12,94025644 | 15,08545568 | 14,09336836 | 13,20017121 | 13,00554065 | 13,67235321 | 13,63 |
| Slc1a2 | 13,33634894 | 12,75711957 | 14,555475 | 13,55496962 | 12,6754371 | 13,27931571 | 13,52826249 | 13,38 |
| Apoe | 12,44739884 | 12,67632065 | 14,14888142 | 12,63234133 | 14,0119283 | 12,94276944 | 14,45893532 | 13,33 |
| ND2 | 13,3617929 | 13,28535184 | 13,8551115 | 14,05481745 | 12,01232994 | 12,35024804 | 13,50181746 | 13,20 |
| Sparcl1 | 12,99520385 | 12,81561882 | 13,92025231 | 13,34296335 | 13,04064628 | 12,95515956 | 13,26199777 | 13,19 |
| ND1 | 13,12108804 | 12,96971746 | 14,00746478 | 13,70308828 | 12,44467627 | 12,03133312 | 13,57716313 | 13,12 |
| Plpp3 | 12,93688688 | 12,49325387 | 13,73467622 | 13,95003564 | 12,29695192 | 13,03450889 | 13,30229177 | 13,11 |
| Atp1a2 | 12,74583582 | 12,49759074 | 14,25632166 | 13,59673619 | 11,96646192 | 13,3231695 | 13,30408935 | 13,10 |
| Plp1 | 12,43320607 | 14,2754418 | 13,1077793 | 11,04128016 | 13,79605263 | 13,37109532 | 12,76551855 | 12,97 |
| Gja1 | 13,11488316 | 12,36832263 | 13,70834525 | 13,83988814 | 12,89817813 | 12,83356626 | 11,78495173 | 12,94 |
| Ptn | 12,66726147 | 12,18601698 | 13,06126549 | 13,7117996 | 12,47642356 | 12,67618356 | 12,91033215 | 12,81 |
| Clu | 11,82257804 | 12,28983191 | 13,65406259 | 12,58983859 | 12,8922707 | 11,67647227 | 12,99649557 | 12,56 |
| Gpm6a | 12,49874935 | 11,88853707 | 13,04947726 | 13,24618024 | 12,575128 | 11,94714185 | 12,68121124 | 12,56 |
| Ptprz1 | 12,93824941 | 12,70811946 | 12,45966806 | 12,57042878 | 11,61916329 | 12,88986437 | 11,65716119 | 12,41 |
| Tspan7 | 12,34238397 | 11,98375773 | 12,82704502 | 11,7463205 | 13,31602248 | 11,74861102 | 11,48757148 | 12,21 |
| Gpm6b | 12,5228861 | 12,33805508 | 12,51410279 | 11,88887779 | 12,19288887 | 11,8233038 | 11,94751011 | 12,18 |
| Cpe | 11,89149134 | 11,60788318 | 13,37663166 | 12,79885237 | 11,79608936 | 11,09943886 | 11,85553869 | 12,06 |
| ND5 | 11,90735997 | 12,3068749 | 13,10753127 | 12,31994784 | 11,7809117 | 10,29677201 | 12,63121969 | 12,05 |
| Pcdh9 | 10,6119109 | 11,69410312 | 13,2020227 | 12,58808007 | 11,97876691 | 11,55147875 | 12,10040483 | 11,96 |

| | | | | | | | | |
|---|---|---|---|---|---|---|---|---|
| Qk | 12,15025044 | 11,85046144 | 12,3693388 | 11,82865908 | 11,95032221 | 11,58442036 | 11,56607161 | 11,90 |
| OC11856734 | 11,23828923 | 11,83518013 | 11,72163852 | 12,59395597 | 11,99574502 | 11,46320443 | 11,9024413 | 11,82 |
| Pla2g7 | 12,35098627 | 12,26413917 | 12,28195529 | 10,68979066 | 13,07830726 | 12,00008558 | 10,04649919 | 11,82 |
| Gria2 | 11,99025131 | 11,90775492 | 11,09620596 | 11,8493674 | 11,59959333 | 12,82412665 | 11,1293542 | 11,77 |
| ATP8 | 11,96097365 | 11,75581119 | 12,63043934 | 12,57871068 | 11,23311467 | 9,955834825 | 11,99988734 | 11,73 |
| Scd2 | 11,39467525 | 11,41821093 | 12,9336988 | 12,00988249 | 10,21917934 | 11,7582427 | 11,80684573 | 11,65 |
| Zbtb20 | 11,40459704 | 11,62124802 | 12,5680875 | 11,84504088 | 11,35285037 | 10,98980325 | 11,70949567 | 11,64 |
| Gpr37l1 | 11,45342944 | 10,7510364 | 12,01592961 | 10,83880953 | 11,93587058 | 11,94415887 | 12,13427056 | 11,58 |
| Son | 11,16759329 | 11,55555099 | 11,40931606 | 11,97183117 | 11,24935351 | 11,93360825 | 11,71414659 | 11,57 |
| Aldoc | 10,74376347 | 10,72905047 | 13,22161798 | 10,59379335 | 12,55628093 | 10,62194943 | 12,37384586 | 11,55 |
| ND6 | 11,56863292 | 11,62932656 | 11,95637048 | 12,57991382 | 9,954423971 | 10,66344756 | 12,34841704 | 11,53 |
| Fth1 | 11,19963181 | 11,83031402 | 11,67619678 | 10,39785135 | 13,04848099 | 11,11343008 | 11,11512426 | 11,48 |
| Prnp | 10,91221074 | 10,70064459 | 12,45113174 | 11,22132824 | 11,52138111 | 11,44482816 | 12,09460564 | 11,48 |
| Ubb | 12,02818172 | 12,52943892 | 13,29177373 | 7,571325734 | 13,04824463 | 10,79121345 | 11,067756 | 11,48 |
| Gstm5 | 11,9030955 | 11,79412343 | 12,36340115 | 10,5648744 | 11,83267229 | 10,5763693 | 11,10805612 | 11,45 |
| S1pr1 | 11,80780636 | 11,19814554 | 11,0016992 | 12,14333995 | 11,25691844 | 11,26273256 | 11,39369927 | 11,44 |
| Cd81 | 11,34428898 | 11,58385796 | 11,49425616 | 11,61623371 | 12,20128173 | 10,99878764 | 10,46713825 | 11,39 |
| Dclk1 | 11,08395169 | 10,88071189 | 12,28854242 | 12,19673532 | 10,97959904 | 10,58843339 | 11,48970513 | 11,36 |
| Htra1 | 10,81870497 | 10,4617355 | 12,70996674 | 12,5577108 | 10,43895145 | 10,90085221 | 11,52345804 | 11,34 |
| Calm1 | 11,50512076 | 11,4243364 | 12,66374961 | 9,698343506 | 11,44085397 | 10,87632454 | 11,74816427 | 11,34 |
| Hnrnpa2b1 | 10,86395997 | 11,07673769 | 11,53896934 | 12,54384579 | 11,17790789 | 10,98817223 | 10,92007828 | 11,30 |
| Itm2b | 11,31807741 | 11,19325072 | 11,58815962 | 11,14736069 | 11,72996865 | 11,18875526 | 10,90126257 | 11,30 |
| Scg3 | 11,01522534 | 10,38892999 | 12,59281471 | 11,43606693 | 10,7812522 | 10,62588651 | 12,15799232 | 11,29 |
| Tmem47 | 12,16731602 | 11,4042102 | 11,00334911 | 10,38171332 | 11,36533045 | 11,53582656 | 10,48782724 | 11,19 |
| Gstm1 | 9,862595771 | 10,11544877 | 12,88114902 | 10,79675096 | 11,68886859 | 9,814167283 | 13,03153705 | 11,17 |
| Hspa8 | 11,71473786 | 10,77304823 | 12,59532481 | 9,302879884 | 12,52623727 | 10,8593197 | 10,35249373 | 11,16 |
| Luzp2 | 11,11521135 | 10,95127443 | 11,53056018 | 10,95639661 | 11,37489173 | 11,54670507 | 10,6396694 | 11,16 |
| Ntrk2 | 10,7529809 | 10,59471795 | 11,82416959 | 11,7382144 | 11,1506638 | 10,95180509 | 10,95717513 | 11,14 |
| Slc6a1 | 11,09771985 | 10,75683166 | 11,52203039 | 11,6194793 | 10,49360715 | 11,25546773 | 11,19670506 | 11,13 |
| Ddx5 | 11,80714591 | 11,43322186 | 11,11274993 | 10,70283015 | 11,60993931 | 11,51421554 | 9,699561712 | 11,13 |
| Mfge8 | 10,97798613 | 10,92426074 | 10,29504832 | 9,016197591 | 12,82078367 | 12,43950588 | 11,22515927 | 11,10 |
| Slc4a4 | 11,39471589 | 11,16486688 | 10,96809152 | 9,910075692 | 10,95789516 | 11,60145637 | 11,37081427 | 11,05 |
| Acsl3 | 11,63249858 | 10,91737241 | 11,12926555 | 11,34213163 | 10,91355333 | 10,73818061 | 10,68907083 | 11,05 |

| | | | | | | | | |
|---|---|---|---|---|---|---|---|---|
| Glul | 11,13555959 | 10,38944987 | 11,71143577 | 11,16913813 | 11,28338353 | 10,59591554 | 11,04993579 | 11,05 |
| Dbi | 11,61631818 | 11,41394769 | 11,8612347 | 10,38664288 | 11,20717982 | 9,652693114 | 11,0541527 | 11,03 |
| Atp1b2 | 10,18069276 | 9,805969282 | 11,95813398 | 11,39856006 | 11,07339329 | 10,16326912 | 12,53789518 | 11,02 |
| Tcf4 | 11,55236898 | 11,57744203 | 10,61766953 | 11,10675753 | 10,7792031 | 11,39021503 | 9,964196167 | 11,00 |
| Ldhb | 10,9636116 | 10,83201751 | 11,35123732 | 10,43082 | 12,33523952 | 10,7150633 | 10,35580619 | 11,00 |
| Aqp4 | 11,2203335 | 10,69304212 | 11,23290361 | 11,79053758 | 10,60446543 | 10,98857942 | 10,44374928 | 11,00 |
| Ednrb | 11,97852295 | 11,41415352 | 11,0063921 | 10,92820753 | 10,6853821 | 11,66714091 | 9,228543447 | 10,99 |
| Ppia | 10,89787357 | 10,44598369 | 12,19345969 | 10,88942636 | 10,77102747 | 9,572249 | 11,90712613 | 10,95 |
| Ckb | 10,37456864 | 10,20186703 | 12,91611461 | 8,365592789 | 11,9264222 | 10,65882801 | 11,87241951 | 10,90 |
| Eef1a1 | 11,04285169 | 11,15573157 | 12,01633143 | 9,476833751 | 11,20170565 | 10,99580886 | 10,33020249 | 10,89 |
| Mt1 | 11,08418027 | 10,79475473 | 11,74254301 | 10,710967 | 11,14323439 | 9,750247715 | 10,9838483 | 10,89 |
| F3 | 10,5458889 | 9,564411188 | 11,64772804 | 12,49812224 | 10,82569351 | 10,10910365 | 10,99109507 | 10,88 |
| Cldn10 | 11,78812229 | 11,44946779 | 9,085163464 | 10,16749317 | 12,21029252 | 11,42580279 | 10,01080065 | 10,88 |
| Msmo1 | 11,23032195 | 11,01911655 | 10,90731967 | 10,24986724 | 11,83119421 | 10,67513928 | 10,12891047 | 10,86 |
| Sox9 | 11,331935 | 11,00304222 | 11,36907989 | 9,886099248 | 11,35969371 | 11,12729531 | 9,914962539 | 10,86 |
| Septin7 | 11,52931081 | 11,25141515 | 10,16519179 | 9,787156151 | 12,31718907 | 10,6942201 | 10,23479476 | 10,85 |
| Macf1 | 11,29999846 | 11,46716723 | 10,4799831 | 9,918239453 | 11,31305215 | 11,59097759 | 9,897811223 | 10,85 |
| Gjb6 | 11,40281648 | 11,2453891 | 9,927914197 | 10,73599963 | 11,78615688 | 11,13855527 | 9,337522632 | 10,80 |
| Selenop | 11,18979619 | 11,1510014 | 10,7866895 | 9,288681169 | 11,972649 | 10,82744207 | 10,32619942 | 10,79 |
| Ttyh1 | 11,07584892 | 10,41931823 | 11,30269614 | 10,15885449 | 10,51545807 | 11,49717556 | 10,51301853 | 10,78 |
| Npas3 | 10,90030217 | 10,8024049 | 9,777079717 | 10,82818638 | 10,45410145 | 11,46252679 | 11,25498274 | 10,78 |
| Fgfr3 | 10,62760003 | 10,66217239 | 10,65793589 | 11,0234607 | 10,13924245 | 11,33096652 | 10,63843923 | 10,73 |
| Actb | 10,64409975 | 10,79327416 | 11,80612308 | 10,64121598 | 10,78132958 | 10,68187637 | 9,687251598 | 10,72 |
| Car2 | 10,25129541 | 10,37286295 | 11,58693236 | 9,71635102 | 11,33319933 | 10,1944805 | 11,55923567 | 10,72 |
| Ndrg2 | 11,05001878 | 11,13896888 | 10,27848563 | 9,208171886 | 12,01794205 | 11,04551863 | 10,27348565 | 10,72 |
| Gm52229 | 10,46652837 | 10,15888974 | 11,07785802 | 11,69026161 | 10,77813884 | 10,97461695 | 9,372110026 | 10,65 |
| Arhgap5 | 11,53448075 | 10,84490634 | 10,44748688 | 10,68502231 | 9,76517861 | 10,68271157 | 10,48502378 | 10,63 |
| Serpine2 | 10,68993485 | 10,96750774 | 10,27233424 | 10,51732296 | 11,63879003 | 10,29833094 | 10,00221181 | 10,63 |
| Prdx6 | 9,9026545 | 9,56946122 | 12,20241468 | 10,94885626 | 11,15750993 | 9,399490087 | 10,93924184 | 10,59 |
| Id2 | 10,6119705 | 10,04048308 | 10,15953931 | 10,72007554 | 10,98574156 | 9,866249499 | 11,54606365 | 10,56 |
| Ccdc88a | 10,37626877 | 10,47270725 | 10,87728257 | 11,82571008 | 9,347422492 | 10,0757564 | 10,86180159 | 10,55 |
| Hsp90aa1 | 11,49754668 | 11,4944208 | 11,18953577 | 8,721218943 | 10,29527777 | 10,53545824 | 10,05604206 | 10,54 |
| Cadm2 | 11,9525836 | 11,49241705 | 9,258480832 | 9,582233922 | 11,40690864 | 11,5575084 | 8,459677259 | 10,53 |

| | | | | | | | | |
|---|---|---|---|---|---|---|---|---|
| Nrxn1 | 10,54061138 | 10,63407868 | 10,77448093 | 10,9589422 | 10,90429475 | 9,815292002 | 10,0805881 | 10,53 |
| Prdx1 | 10,65890492 | 10,67421477 | 10,78703537 | 10,73227261 | 10,54281383 | 9,558186137 | 10,62586579 | 10,51 |
| Fos | 11,62425463 | 11,67645163 | 10,0500665 | 10,30178495 | 10,78625162 | 9,478792875 | 9,31812262 | 10,46 |
| Ntsr2 | 9,429187964 | 9,439466641 | 11,31761961 | 11,39147183 | 10,45563813 | 9,816442738 | 11,33964784 | 10,46 |
| Syt11 | 10,4625725 | 10,58309047 | 10,96739283 | 9,09408582 | 10,14102793 | 10,82732252 | 11,09171608 | 10,45 |
| Gabrb1 | 10,49461153 | 10,60987481 | 10,23047457 | 10,71987234 | 10,91573052 | 10,93213875 | 9,087455383 | 10,43 |
| Bcan | 9,117787957 | 8,995480521 | 11,87108058 | 10,98564308 | 9,549528645 | 10,60687942 | 11,7835056 | 10,42 |
| Rbm39 | 10,37861354 | 10,96342653 | 8,744790848 | 11,42312489 | 10,31734667 | 10,63731378 | 10,44225291 | 10,42 |
| Serinc1 | 10,50198102 | 9,854007181 | 10,3248739 | 11,78244335 | 10,61402652 | 10,16848927 | 9,582981163 | 10,40 |
| Glud1 | 11,26871435 | 10,42369628 | 10,11950986 | 10,11542727 | 10,21471704 | 11,05391713 | 9,571952284 | 10,40 |
| Cd63 | 11,76354392 | 11,27672498 | 10,20650948 | 8,328679572 | 11,79575076 | 11,05204834 | 8,333831147 | 10,39 |
| Tril | 11,06048646 | 11,19107717 | 9,50227765 | 9,307080754 | 10,76584845 | 11,60187189 | 9,276695305 | 10,39 |
| OC1185682 | 9,682335167 | 10,44280711 | 11,000194 | 11,60812372 | 10,13212579 | 9,642850802 | 10,17920165 | 10,38 |
| Gm33649 | 9,188726999 | 9,977623016 | 10,75582479 | 12,08403792 | 10,61011888 | 9,528372556 | 10,51959769 | 10,38 |
| Tmem59 | 10,19201941 | 10,35755557 | 10,76263147 | 9,900045166 | 11,29151982 | 10,10185679 | 10,0316335 | 10,38 |
| Mmd2 | 10,32518611 | 9,894600633 | 10,73663416 | 9,613693966 | 11,32455575 | 10,28835813 | 10,38388436 | 10,37 |
| Cadm1 | 10,6271437 | 10,72651362 | 10,42110001 | 8,018389334 | 10,91193955 | 11,54106892 | 10,19428796 | 10,35 |
| Sparc | 10,15208352 | 10,23205861 | 10,28443757 | 10,29455092 | 11,96050571 | 10,71205426 | 8,64208177 | 10,33 |
| Hsp90b1 | 11,25613806 | 10,94168542 | 9,855078549 | 9,336167144 | 10,81026633 | 10,77596937 | 9,216756302 | 10,31 |
| Id4 | 11,11994003 | 10,45711957 | 10,34392679 | 9,867652943 | 9,712728824 | 10,62762794 | 9,938774168 | 10,30 |
| Luc7l3 | 9,483258068 | 10,0704161 | 9,717023769 | 11,27523704 | 9,842595838 | 10,22818535 | 11,09776079 | 10,24 |
| Slc39a12 | 9,815323824 | 9,574895033 | 10,68214714 | 11,30994063 | 10,19518861 | 9,784144711 | 10,2988528 | 10,24 |
| Hacd3 | 11,45066991 | 11,2321918 | 9,624905809 | 7,31242407 | 11,63896137 | 12,04903887 | 8,234238752 | 10,22 |
| Hspa5 | 11,39918812 | 10,57186742 | 10,05050472 | 9,626677201 | 11,25326229 | 10,61264596 | 7,910009957 | 10,20 |
| Fut9 | 10,9808597 | 10,90959856 | 9,295429966 | 9,544578657 | 10,19341875 | 10,47783489 | 9,983705179 | 10,20 |
| Fam107a | 10,98205028 | 10,04949608 | 10,62597485 | 9,792403104 | 10,26885129 | 10,05681344 | 9,483800612 | 10,18 |
| Actg1 | 10,15128601 | 10,30938117 | 10,21134255 | 10,2038871 | 10,76412026 | 10,11348956 | 9,461443297 | 10,17 |
| Sat1 | 10,74000373 | 10,77297044 | 9,314733815 | 9,002559473 | 10,50966796 | 11,10338903 | 9,693634379 | 10,16 |
| Lrrc58 | 11,11708606 | 10,00888671 | 10,32502726 | 10,11635802 | 9,991055582 | 10,33525288 | 9,20744305 | 10,16 |
| Ogt | 9,99716978 | 10,70197848 | 9,074133994 | 10,83887286 | 10,55922125 | 9,916739283 | 9,948865178 | 10,15 |
| Fabp7 | 10,29531748 | 9,762184646 | 11,49902474 | 10,68820082 | 7,734444933 | 9,576218921 | 11,32092446 | 10,13 |
| Klf13 | 10,45449555 | 10,32420933 | 10,02485154 | 12,29620393 | 8,412409934 | 9,898480529 | 9,447574766 | 10,12 |
| Canx | 11,28079885 | 11,31792981 | 9,819428431 | 9,489540781 | 10,40684225 | 11,00231135 | 7,497827399 | 10,12 |

| | | | | | | | | |
|---|---|---|---|---|---|---|---|---|
| Atp5b | 9,594036727 | 9,169719548 | 11,46776757 | 9,806557651 | 10,91788785 | 9,825633726 | 9,904299066 | 10,10 |
| Rcn2 | 10,99653165 | 10,99903335 | 10,4229995 | 8,326330087 | 10,72553959 | 10,88296479 | 8,323741824 | 10,10 |
| Celf2 | 10,32810525 | 10,01001878 | 10,12016436 | 11,23823913 | 9,718031494 | 9,4938286 | 9,655255898 | 10,08 |
| Kif1b | 10,58904203 | 10,62947299 | 8,694524312 | 8,828722795 | 10,23051433 | 11,15227856 | 10,2755173 | 10,06 |
| Ptma | 10,44665349 | 10,26607144 | 10,61552599 | 7,789447785 | 10,34949798 | 9,870889746 | 10,925678 | 10,04 |
| Slc25a4 | 10,12161008 | 10,3248376 | 9,964379888 | 9,66640298 | 11,03160921 | 9,722257539 | 9,294234077 | 10,02 |
| Syne1 | 10,4011693 | 9,998370553 | 9,85069766 | 9,568785776 | 10,44621669 | 10,48455629 | 9,215587772 | 10,00 |
| Dpysl2 | 10,64529654 | 11,25079918 | 9,10156515 | 8,540734334 | 11,41152328 | 10,22235319 | 8,764734194 | 9,99 |
| Brd7 | 10,65291973 | 9,796794653 | 9,197885972 | 12,4017201 | 8,374726233 | 9,745846488 | 9,4507189 | 9,95 |
| Grm3 | 10,76528553 | 10,08677643 | 10,11848713 | 9,867778995 | 10,14742957 | 10,1209425 | 8,51119513 | 9,95 |
| Etfa | 9,77129376 | 8,622586829 | 11,15866402 | 10,34180004 | 10,94867605 | 9,529215695 | 9,131852964 | 9,93 |
| Mertk | 10,1515276 | 9,904847811 | 9,551310334 | 8,526185096 | 10,5612103 | 10,67395828 | 9,993380904 | 9,91 |
| Ank2 | 10,17319517 | 10,14264084 | 9,250924161 | 9,75406795 | 10,29139836 | 10,45936814 | 9,172524701 | 9,89 |
| Gnas | 10,71773879 | 10,14061334 | 10,63526709 | 9,041516114 | 9,405024619 | 9,720736244 | 9,536038088 | 9,89 |
| Slco1c1 | 9,394157172 | 9,370153954 | 10,30218765 | 10,25197522 | 9,998291062 | 9,473383417 | 10,40023639 | 9,88 |
| Dtna | 9,769058128 | 10,14504706 | 12,02794036 | 8,478493449 | 9,909234378 | 9,346793515 | 9,511742018 | 9,88 |
| Acsl6 | 10,37583017 | 10,18157222 | 9,443768834 | 9,179760901 | 9,768546803 | 10,54590973 | 9,681873655 | 9,88 |
| Trf | 8,770853654 | 10,95412447 | 10,26014729 | 7,451252806 | 12,18073399 | 10,36861092 | 9,098722795 | 9,87 |
| Cpeb4 | 9,786100453 | 9,665901743 | 10,17453963 | 10,88489536 | 8,764946363 | 9,935192718 | 9,86762005 | 9,87 |
| Fermt2 | 9,615458945 | 9,497062207 | 10,23765417 | 9,752100499 | 9,985564147 | 9,610281696 | 10,26895431 | 9,85 |
| Nfia | 9,812947046 | 10,10694388 | 10,775561 | 9,015691059 | 9,458630316 | 10,22523515 | 9,508394412 | 9,84 |
| Appl2 | 9,579401425 | 9,783861048 | 8,574779086 | 10,50701885 | 8,980383905 | 10,0786224 | 11,34786655 | 9,84 |
| Tspan3 | 10,41473604 | 10,31633469 | 9,94630113 | 9,087222337 | 11,04298115 | 9,716178587 | 8,310653223 | 9,83 |
| Prxl2a | 9,89673796 | 9,3773553 | 10,14157535 | 10,38813719 | 10,84999598 | 8,792847015 | 9,333852188 | 9,83 |
| Arl6ip1 | 10,13252894 | 9,653234989 | 9,320972232 | 8,451761876 | 10,85490786 | 10,48972056 | 9,83644154 | 9,82 |
| Gatm | 10,00817912 | 11,61512131 | 9,584495768 | 6,941929114 | 12,37378348 | 9,201587932 | 8,929722459 | 9,81 |
| Acsbg1 | 10,18927918 | 10,16437898 | 9,650065796 | 8,019861169 | 11,60852655 | 10,47715671 | 8,442325007 | 9,79 |
| Selenok | 10,36580295 | 10,69157395 | 9,826679149 | 7,841900294 | 11,19897972 | 10,19021636 | 8,423317737 | 9,79 |
| Aldoa | 9,311788219 | 8,847479403 | 10,73776418 | 9,540744531 | 11,09543219 | 9,43200769 | 9,494376176 | 9,78 |
| Gm41284 | 9,135259617 | 9,470037593 | 11,11521884 | 10,49254156 | 9,1363682 | 9,005599191 | 9,84118413 | 9,74 |
| Gm15521 | 8,276500628 | 9,21634121 | 11,15922599 | 11,36431691 | 9,425755721 | 9,599743224 | 9,142779433 | 9,74 |
| Mal | 9,808385057 | 11,65065395 | 9,13422047 | 6,642544339 | 11,41249091 | 10,51747108 | 8,929598234 | 9,73 |
| Clk1 | 10,73019099 | 10,68499726 | 7,524568656 | 9,883725191 | 10,30081344 | 9,895395857 | 9,031028171 | 9,72 |

| | | | | | | | | |
|---|---|---|---|---|---|---|---|---|
| Ndufc2 | 11,21750306 | 10,62021088 | 8,625657308 | 8,549734589 | 11,12533642 | 9,488925017 | 8,27152428 | 9,70 |
| Pnisr | 9,403642822 | 9,578221439 | 9,813885582 | 9,731768307 | 9,507655852 | 9,948729665 | 9,869847511 | 9,69 |
| Hnrnph1 | 10,66513754 | 10,82649486 | 7,374328215 | 10,50403889 | 10,02326334 | 10,41641051 | 7,984401413 | 9,68 |
| Ugp2 | 10,78128623 | 10,51611162 | 9,523417587 | 7,542250643 | 10,018769 | 10,58230895 | 8,802441618 | 9,68 |
| Ddx3x | 9,623057959 | 9,631545001 | 9,468463172 | 9,47873631 | 9,798989307 | 10,021571 | 9,720676112 | 9,68 |
| Ddhd1 | 9,370782245 | 9,721511582 | 9,795783399 | 10,5505545 | 8,930785799 | 9,559049886 | 9,80004837 | 9,68 |
| Zfp36l1 | 10,77701432 | 11,61100326 | 8,541181965 | 8,500384534 | 9,739227947 | 9,866427482 | 8,5843974 | 9,66 |
| Paqr8 | 10,21245979 | 9,383770088 | 10,37600186 | 10,2047568 | 9,217318103 | 9,797385823 | 8,406556551 | 9,66 |
| Atp5j | 10,169531 | 9,937500407 | 10,00092917 | 9,742667866 | 9,43419459 | 9,189297144 | 9,064943261 | 9,65 |
| Ctsb | 9,831749736 | 9,335165112 | 9,706474218 | 9,695211029 | 10,43336215 | 9,545537603 | 8,98964578 | 9,65 |
| ND3 | 10,3261019 | 10,00831897 | 9,603769814 | 10,73088204 | 8,242924935 | 9,001714184 | 9,607350823 | 9,65 |
| Rtn3 | 10,25186233 | 9,873090206 | 10,80661592 | 8,96925789 | 9,655650746 | 10,07588555 | 7,849648277 | 9,64 |
| Gnao1 | 8,605804008 | 8,922659392 | 10,64666463 | 9,633165606 | 8,73592665 | 9,522338044 | 11,39496754 | 9,64 |
| Omg | 9,181962362 | 9,556449041 | 9,091544102 | 10,40212607 | 9,763195378 | 9,021102757 | 10,44445615 | 9,64 |
| Lamp1 | 10,0883986 | 10,26878818 | 10,03805442 | 6,975641043 | 11,06336432 | 10,10996318 | 8,896515363 | 9,63 |
| Mlc1 | 8,840181682 | 8,846558572 | 11,9260909 | 8,177989401 | 10,1773562 | 8,673216293 | 10,79051859 | 9,63 |
| Rsrp1 | 9,161866897 | 9,757113504 | 8,796054585 | 9,943310617 | 10,57232333 | 9,700292899 | 9,487059708 | 9,63 |
| Sorbs1 | 9,985839151 | 9,586157501 | 9,888647891 | 8,357938837 | 10,78712285 | 9,594709547 | 9,201232063 | 9,63 |
| Dmd | 9,578144957 | 9,935369814 | 9,887021738 | 10,49707955 | 9,922672715 | 9,390319624 | 8,142564149 | 9,62 |
| Tmsb4x | 9,39977252 | 9,780759373 | 10,22435358 | 9,918651408 | 9,375322046 | 8,643378565 | 9,987616564 | 9,62 |
| Hepacam | 8,510479756 | 8,733449177 | 11,0562973 | 9,795824966 | 9,766884268 | 9,459207251 | 9,953069039 | 9,61 |
| Psap | 8,781747484 | 8,923468518 | 10,35458605 | 9,074432836 | 9,964333578 | 9,774608537 | 10,39138315 | 9,61 |
| Sec62 | 10,3979446 | 10,14953496 | 9,244462356 | 8,504740911 | 9,717715553 | 10,01383377 | 9,235028821 | 9,61 |
| Pitpnc1 | 9,401430413 | 9,375143182 | 10,52880789 | 10,10747628 | 9,088878014 | 9,959078035 | 8,793705714 | 9,61 |
| Mt3 | 9,168422819 | 9,086163986 | 9,873380896 | 9,403377867 | 10,08420192 | 8,786340042 | 10,83747049 | 9,61 |
| Gm3764 | 8,813810962 | 9,751358061 | 9,633689996 | 9,404537803 | 9,657032948 | 10,09035593 | 9,849139808 | 9,60 |
| Tmed5 | 9,689982938 | 9,164596718 | 9,7425667 | 11,16826642 | 9,342054987 | 9,424757983 | 8,652404667 | 9,60 |
| S100a1 | 9,848059946 | 9,677325836 | 9,946737348 | 10,12930119 | 9,740690677 | 8,974997254 | 8,845721535 | 9,59 |
| Ncam1 | 9,334714396 | 9,741322037 | 10,48903063 | 8,617844423 | 9,586564692 | 9,894366727 | 9,461043401 | 9,59 |
| Rnf13 | 9,980322369 | 10,18022999 | 9,28074161 | 9,57492618 | 10,75497523 | 8,950576003 | 8,390039313 | 9,59 |
| Ntm | 9,309627635 | 9,317138253 | 9,270026381 | 10,29499333 | 9,147288762 | 10,46422903 | 9,305416595 | 9,59 |
| Cryab | 9,194740405 | 10,106626 | 9,897672792 | 7,58345482 | 11,11863368 | 8,8960358 | 10,30840575 | 9,59 |
| Lsamp | 8,943026054 | 8,974547706 | 11,30763288 | 8,186998147 | 9,860234675 | 9,20159112 | 10,61539668 | 9,58 |

| | | | | | | | | |
|---|---|---|---|---|---|---|---|---|
| Gm32337 | 9,713954151 | 10,18291853 | 8,577817287 | 9,896125656 | 9,52863467 | 9,193896793 | 9,982478097 | 9,58 |
| Cox7c | 10,24094153 | 10,05098329 | 10,30123898 | 9,633669506 | 10,0734509 | 9,425103401 | 7,308234596 | 9,58 |
| Il18 | 10,1585801 | 9,954216779 | 8,513281509 | 9,522090972 | 10,50043524 | 10,07240208 | 8,261419459 | 9,57 |
| Rps14 | 10,32839262 | 10,40856297 | 9,367423596 | 9,775868422 | 9,422663543 | 8,809470767 | 8,687086491 | 9,54 |
| Wsb1 | 9,810200139 | 10,74741082 | 8,831276027 | 7,571447686 | 10,48426718 | 10,42622393 | 8,919826701 | 9,54 |
| Ncoa4 | 10,84456491 | 10,72836566 | 9,695231341 | 12,19998874 | 7,573403518 | 6,706931343 | 8,868953074 | 9,52 |
| H3f3a | 10,2981293 | 9,773010815 | 9,19556406 | 9,445497858 | 10,27293333 | 9,833681418 | 7,789825575 | 9,52 |
| Cycs | 10,82292695 | 10,34249806 | 9,387519042 | 9,473465804 | 10,37700318 | 9,152546903 | 6,99402503 | 9,51 |
| Eif5 | 9,083425272 | 9,510089191 | 10,94166704 | 8,391809773 | 9,270692637 | 9,799705098 | 9,502052496 | 9,50 |
| Slc25a3 | 9,608590606 | 9,490568103 | 9,157626857 | 9,633162973 | 10,24147582 | 9,67201027 | 8,660360587 | 9,49 |
| Nnat | 10,11221185 | 9,758816962 | 9,410032724 | 7,763967532 | 11,19066991 | 8,575789553 | 9,636621608 | 9,49 |
| Rab14 | 9,138057416 | 9,734537353 | 10,25750406 | 9,079492956 | 9,922817623 | 9,347818714 | 8,907016969 | 9,48 |
| Cspg5 | 9,332188809 | 9,038515052 | 8,61008218 | 10,04701533 | 8,93761425 | 9,764849566 | 10,65616286 | 9,48 |
| Cmtm5 | 9,215438424 | 9,591984521 | 9,104470108 | 8,419147019 | 9,994607987 | 9,383022129 | 10,62554992 | 9,48 |
| Timp3 | 9,576426348 | 9,56970504 | 9,424874957 | 8,828489769 | 10,08957087 | 9,879749378 | 8,948208906 | 9,47 |
| Myo6 | 9,434940535 | 10,36454939 | 9,066580899 | 9,101273462 | 8,691431004 | 10,05437103 | 9,573615706 | 9,47 |
| 330159H11F | 9,249620704 | 10,48727119 | 8,579779037 | 9,070997483 | 9,650161095 | 9,325644617 | 9,91627517 | 9,47 |
| Klhl9 | 9,482702957 | 9,240614523 | 10,10557334 | 11,22395046 | 8,989433546 | 8,891203239 | 8,323415106 | 9,47 |
| Itgav | 10,08538049 | 10,00629492 | 8,473825436 | 8,973687692 | 9,336323366 | 9,831351557 | 9,467989458 | 9,45 |
| Ndfip1 | 10,57614339 | 10,2197788 | 9,124595393 | 7,479850397 | 10,6147859 | 9,812787847 | 8,169457849 | 9,43 |
| Irf2bpl | 9,954281057 | 9,640352835 | 8,810612283 | 11,97086988 | 7,851992869 | 9,258117729 | 8,485529273 | 9,42 |
| Hsp90ab1 | 10,25888447 | 9,609048352 | 10,53494526 | 6,176426197 | 9,777784836 | 9,748286315 | 9,742603884 | 9,41 |
| Lgr4 | 10,43929028 | 10,22636042 | 8,888382271 | 10,36489873 | 9,72770924 | 9,501529957 | 6,599374654 | 9,39 |
| Ptgds | 9,764783041 | 11,95849619 | 10,4967599 | 2,473150534 | 11,15669235 | 10,88335028 | 9,010120325 | 9,39 |
| Prex2 | 10,72989478 | 9,917490358 | 8,119915583 | 9,642633706 | 8,887632449 | 9,590063062 | 8,848607446 | 9,39 |
| Hmgb1 | 9,601484294 | 8,842102631 | 9,906856455 | 8,66887589 | 10,5487017 | 8,498135684 | 9,54743504 | 9,37 |
| Ndufa12 | 10,12276083 | 9,492041126 | 9,755687093 | 9,605054124 | 9,649431985 | 8,487779662 | 8,487325593 | 9,37 |
| Hnrnpa3 | 9,817561488 | 9,993408669 | 8,659219942 | 8,498434389 | 9,79425289 | 10,42301282 | 8,395756063 | 9,37 |
| Degs1 | 9,145141056 | 9,745598425 | 9,902535101 | 8,916320657 | 10,54841377 | 9,290111556 | 7,930375283 | 9,35 |
| Trim9 | 9,720460299 | 10,55050399 | 8,672980308 | 6,53397848 | 9,715645717 | 9,893443271 | 10,36092689 | 9,35 |
| Ptges3 | 10,66224251 | 9,865822207 | 8,822983312 | 9,132552916 | 9,406304079 | 9,718987855 | 7,819046969 | 9,35 |
| Slc25a5 | 8,780796377 | 8,612719286 | 10,8770911 | 9,439506921 | 9,937377192 | 8,127714244 | 9,628680949 | 9,34 |
| Sdc4 | 9,878130624 | 9,635250786 | 9,672059328 | 8,136698994 | 9,901695102 | 9,491616995 | 8,670122672 | 9,34 |

| | | | | | | | | |
|---|---|---|---|---|---|---|---|---|
| Ddah1 | 10,03646715 | 9,607449634 | 10,08768186 | 8,416116423 | 9,589021195 | 9,169300638 | 8,475215452 | 9,34 |
| Lxn | 9,736711576 | 9,259810903 | 10,40090579 | 8,867960141 | 10,55105688 | 8,874175283 | 7,632955524 | 9,33 |
| Glrb | 10,31810801 | 9,444848714 | 8,094003972 | 9,668498951 | 10,09676044 | 9,103347465 | 8,584583459 | 9,33 |
| Arxes2 | 9,480674839 | 7,656008427 | 10,95397725 | 9,154636091 | 9,285258014 | 9,845880995 | 8,918206863 | 9,33 |
| Tmed10 | 10,28014037 | 9,79678179 | 10,26012271 | 5,471730612 | 11,1285437 | 9,261569268 | 9,047460611 | 9,32 |
| Tmem229a | 10,24011286 | 10,10811389 | 9,86514271 | 9,239525213 | 9,400278253 | 8,699750784 | 7,655740333 | 9,32 |
| Rora | 8,672569707 | 8,649786904 | 10,25198968 | 9,997294453 | 8,874933695 | 8,91760161 | 9,806464657 | 9,31 |
| Fjx1 | 9,065533784 | 8,450515046 | 10,19426227 | 9,635030528 | 8,460185997 | 10,10628552 | 9,258123968 | 9,31 |
| Rn7sk | 10,37307618 | 10,22457937 | 8,426668756 | 8,881488232 | 9,888253419 | 9,762050201 | 7,606907923 | 9,31 |
| Mt2 | 9,800107221 | 9,15173043 | 9,934738595 | 9,343436263 | 8,956206283 | 8,396183344 | 9,547719306 | 9,30 |
| Lix1 | 9,368687386 | 9,660585556 | 9,798750567 | 8,79035385 | 9,750090237 | 9,802063756 | 7,951717278 | 9,30 |
| Kmt2e | 8,806943158 | 8,945180966 | 9,663944743 | 10,17257418 | 8,936868118 | 9,200269413 | 9,364447857 | 9,30 |
| Gm30618 | 9,745844427 | 8,867137604 | 9,062802625 | 11,77086101 | 5,763712553 | 10,7039873 | 9,172388098 | 9,30 |
| Dazap2 | 10,38427331 | 9,956068627 | 9,823337817 | 7,799338783 | 11,14343815 | 8,319786133 | 7,608664177 | 9,29 |
| Ubc | 9,056686086 | 7,588809234 | 10,6070297 | 8,707979825 | 10,05625892 | 9,374040709 | 9,618403111 | 9,29 |
| App | 9,351282733 | 10,3702016 | 9,633333329 | 7,818953704 | 8,810621568 | 10,47685969 | 8,535813343 | 9,29 |
| Aplp1 | 8,916489689 | 9,401503101 | 9,537824938 | 7,607246848 | 9,119013391 | 9,767266353 | 10,6449487 | 9,28 |
| Myo10 | 8,122667059 | 8,063105118 | 11,10209136 | 10,84683362 | 8,124017615 | 9,156851907 | 9,538348522 | 9,28 |
| Kmt2c | 8,911559175 | 9,230209882 | 9,588554667 | 9,39325563 | 9,555926521 | 9,743525909 | 8,511186203 | 9,28 |
| Atp6v0e | 10,01547593 | 9,619435086 | 9,67582913 | 9,171857502 | 9,617697636 | 8,533214596 | 8,297109911 | 9,28 |
| Eno1 | 9,066002734 | 9,63194251 | 10,02676234 | 5,634225904 | 11,53882244 | 8,897310043 | 10,12618142 | 9,27 |
| Zeb1 | 8,876744246 | 9,304842202 | 8,234502649 | 10,80747072 | 9,420082671 | 9,862725635 | 8,382325858 | 9,27 |
| Dst | 9,055862572 | 9,244378051 | 10,80009558 | 8,033705325 | 9,601585857 | 9,443123369 | 8,709928797 | 9,27 |
| Spag9 | 10,30073301 | 10,13624599 | 8,655137684 | 5,72686679 | 10,53635641 | 10,58424705 | 8,916654749 | 9,27 |
| Spcs2 | 9,876940299 | 9,307946304 | 10,52420096 | 10,22945535 | 9,911089882 | 9,101820503 | 5,89299564 | 9,26 |
| Ptch1 | 9,828230203 | 9,612285664 | 6,305027952 | 9,601929379 | 8,920850647 | 10,38477378 | 10,18365009 | 9,26 |
| Fabp5 | 9,750822503 | 10,1627499 | 8,569581491 | 9,128243443 | 9,531188913 | 9,963923173 | 7,703789779 | 9,26 |
| Rps27 | 10,57459807 | 10,3030553 | 9,146425218 | 8,113389096 | 9,202351401 | 9,310583946 | 8,138680275 | 9,26 |
| Srsf1 | 9,748117104 | 9,427009674 | 9,325920947 | 9,571539051 | 9,263209839 | 9,985654805 | 7,404456103 | 9,25 |
| Arglu1 | 9,445153463 | 9,838127221 | 8,324798191 | 11,06557058 | 8,053040112 | 9,519175308 | 8,476872186 | 9,25 |
| Atp2a2 | 9,223351673 | 8,778906 | 9,547260158 | 10,66704346 | 7,677204842 | 8,915796167 | 9,89588807 | 9,24 |
| Slc27a1 | 8,85402133 | 9,083775636 | 8,154027686 | 9,511888279 | 9,455258384 | 9,655751097 | 9,990024843 | 9,24 |
| Chpt1 | 10,21516291 | 9,927857801 | 10,11968389 | 6,049908556 | 9,683457794 | 10,07242581 | 8,623397739 | 9,24 |

| | | | | | | | | |
|---|---|---|---|---|---|---|---|---|
| Timp4 | 9,679460379 | 9,38702727 | 9,55007056 | 8,185262805 | 10,56411158 | 9,361446396 | 7,95235966 | 9,24 |
| Atp5c1 | 10,32459137 | 10,55738223 | 8,647554399 | 7,653534871 | 10,62856427 | 9,482604402 | 7,353527321 | 9,24 |
| Hnrnpu | 9,647052878 | 9,693775791 | 10,75448376 | 6,507389602 | 9,274716715 | 9,792816964 | 8,918006228 | 9,23 |
| Cox6c | 10,27285762 | 9,63803839 | 9,223379729 | 8,851601636 | 9,537852798 | 9,346219394 | 7,592329098 | 9,21 |
| Eif4g2 | 9,891132469 | 9,582004354 | 9,062203987 | 8,831535145 | 9,127194919 | 9,103200711 | 8,848515717 | 9,21 |
| )30402H24F | 9,068484312 | 9,159927833 | 9,447882968 | 8,727291759 | 9,098542297 | 8,988482769 | 9,952874119 | 9,21 |
| Nedd4 | 10,49106254 | 9,697482374 | 8,079394229 | 8,604559191 | 8,977529491 | 10,15564113 | 8,386783781 | 9,20 |
| Gabrg1 | 9,82223748 | 8,382530666 | 9,264970413 | 8,888419431 | 9,453693196 | 9,3336902 | 9,218222023 | 9,19 |
| Hadhb | 9,701110457 | 8,580966472 | 10,70996924 | 6,709188642 | 11,52814762 | 10,41986254 | 6,701287361 | 9,19 |
| Thrsp | 9,391806458 | 8,452882999 | 10,89094234 | 9,673306544 | 9,381503094 | 7,611680671 | 8,871633272 | 9,18 |
| Insig1 | 9,054972595 | 9,200230442 | 9,114338586 | 9,834132935 | 9,131395561 | 9,368020483 | 8,530244442 | 9,18 |
| Itgb8 | 11,0363275 | 10,61380841 | 7,572752978 | 6,428636042 | 10,74420038 | 9,699259358 | 8,137414275 | 9,18 |
| Neat1 | 8,754156114 | 9,623540811 | 9,624625701 | 7,643469943 | 7,671379924 | 10,62129505 | 10,29388152 | 9,18 |
| Taok1 | 9,70062914 | 9,589786791 | 9,37911006 | 8,068775016 | 9,297473326 | 9,813461568 | 8,381611646 | 9,18 |
| Cbx3 | 10,17683867 | 10,61701619 | 8,92163977 | 9,590934582 | 9,717956078 | 8,477524165 | 6,685646965 | 9,17 |
| Chst2 | 9,827878927 | 9,116595694 | 9,098712421 | 8,7035828 | 9,367523718 | 9,316672827 | 8,749546435 | 9,17 |
| Ywhaz | 9,392446133 | 9,483734305 | 9,58827543 | 8,223686664 | 9,81743051 | 8,482795229 | 9,186999387 | 9,17 |
| Sub1 | 9,289194908 | 9,012668927 | 10,80174442 | 8,432519501 | 8,951883396 | 8,199020266 | 9,478590758 | 9,17 |
| Megf10 | 9,800259079 | 9,619351294 | 8,111700877 | 8,926436247 | 9,670665108 | 10,26432346 | 7,724852346 | 9,16 |
| Uros | 8,896289793 | 8,368919089 | 9,448120904 | 10,75304284 | 9,68172384 | 9,280403563 | 7,657902047 | 9,16 |
| Hmgn3 | 9,777769603 | 9,480278166 | 7,903286741 | 8,897454383 | 10,00433968 | 9,503126112 | 8,502616322 | 9,15 |
| Grina | 8,323824307 | 9,213425625 | 9,232634792 | 7,755142104 | 10,34447133 | 9,305244591 | 9,803111183 | 9,14 |
| Ahcyl1 | 9,426461029 | 9,33085222 | 9,581786397 | 7,149776685 | 9,792004355 | 8,595623303 | 10,05207018 | 9,13 |
| Tmem30a | 9,960794235 | 9,726517914 | 9,221474598 | 8,925854769 | 9,095826137 | 10,00909373 | 6,98426607 | 9,13 |
| Slc3a2 | 9,736593886 | 9,905115474 | 8,160809731 | 6,22407704 | 10,7474188 | 10,73469946 | 8,38598195 | 9,13 |
| Tmco1 | 9,480093543 | 9,44544516 | 9,519880323 | 7,51830421 | 10,32210885 | 9,885942771 | 7,721262901 | 9,13 |
| Srsf5 | 9,078968385 | 9,789855593 | 7,071871657 | 10,10869678 | 10,21394643 | 9,801565014 | 7,805440139 | 9,12 |
| Fam171b | 9,387507353 | 9,385224021 | 8,019850561 | 8,92292572 | 8,649315057 | 9,706373688 | 9,798509231 | 9,12 |
| Ywhaq | 9,288871395 | 10,01811265 | 8,159299219 | 9,091785448 | 10,98781461 | 8,147975913 | 8,169001206 | 9,12 |
| Rpl5 | 9,917408063 | 9,962916384 | 8,73312983 | 7,961343063 | 8,765468715 | 9,89634807 | 8,620406156 | 9,12 |
| Smpdl3a | 8,226835284 | 8,29341862 | 10,64268403 | 10,79186471 | 8,538374006 | 8,294453088 | 9,061281853 | 9,12 |
| Chchd2 | 8,68732336 | 8,445022083 | 10,60912549 | 8,771997637 | 9,124217601 | 7,825731993 | 10,38409229 | 9,12 |
| Tuba1a | 9,374052116 | 10,29494447 | 10,94329555 | 6,516870898 | 9,575749427 | 8,224695369 | 8,830307293 | 9,11 |

| | | | | | | | | |
|---|---|---|---|---|---|---|---|---|
| Ctsd | 8,019083118 | 7,926786895 | 10,87336942 | 9,48151823 | 9,293757048 | 9,582802906 | 8,525959664 | 9,10 |
| Tmed7 | 9,758433303 | 9,858616614 | 6,310578558 | 9,771039517 | 8,185473193 | 9,824248745 | 9,992643398 | 9,10 |
| Vapa | 9,345229815 | 8,711021162 | 8,956744041 | 10,27263044 | 9,44509029 | 9,464354582 | 7,481624836 | 9,10 |
| Aplp2 | 8,761366611 | 9,028924591 | 10,5479312 | 8,841391434 | 9,105461441 | 8,596711576 | 8,75252551 | 9,09 |
| Hacd2 | 9,634632971 | 7,850179055 | 8,689303914 | 9,441925319 | 10,50545845 | 8,950090287 | 8,527976904 | 9,09 |
| Rac1 | 10,05512036 | 10,49002219 | 8,465567112 | 6,79442195 | 9,519314744 | 9,719427922 | 8,542360643 | 9,08 |
| Bnip3l | 10,04351758 | 9,749637848 | 8,107355575 | 7,790151283 | 10,73089113 | 10,0219759 | 7,142560935 | 9,08 |
| Abhd3 | 9,496155962 | 8,885652281 | 5,983329695 | 9,295312543 | 10,74679854 | 9,887782772 | 9,279875622 | 9,08 |
| Mbnl2 | 9,138910666 | 9,611116073 | 10,47890022 | 8,83322891 | 9,03248753 | 9,423272512 | 7,02622992 | 9,08 |
| Nrcam | 9,255393926 | 9,977681097 | 10,00964908 | 8,01798737 | 8,175455139 | 9,780331284 | 8,301197141 | 9,07 |
| Luc7l2 | 8,663136705 | 9,113121799 | 8,938637371 | 9,5367707 | 8,851208931 | 8,980851786 | 9,424967713 | 9,07 |
| OC11548692 | 8,657587393 | 8,567788486 | 9,649294619 | 10,08197687 | 8,192696855 | 8,780513685 | 9,577346486 | 9,07 |
| Srsf2 | 9,11806568 | 9,13135029 | 9,43492514 | 10,13875527 | 9,123176957 | 8,002758954 | 8,545339408 | 9,07 |
| Ubb-ps | 8,138968607 | 8,99138318 | 11,66925079 | 7,155911027 | 9,311143998 | 8,585455935 | 9,582014173 | 9,06 |
| Dynll1 | 9,863067916 | 9,273510394 | 9,118760704 | 8,398278582 | 9,78275427 | 8,959091696 | 8,038599209 | 9,06 |
| Alcam | 10,05121024 | 10,38341753 | 5,258991356 | 9,663567211 | 9,555072679 | 10,25019438 | 8,254322261 | 9,06 |
| OC11856734 | 8,925398437 | 8,78548917 | 9,516363268 | 9,703316242 | 8,957160763 | 8,152384787 | 9,281350389 | 9,05 |
| Hmgcs1 | 9,535734423 | 8,915490906 | 10,01669124 | 7,857950866 | 8,865257304 | 9,218647831 | 8,872964307 | 9,04 |
| Mbp | 7,98938184 | 9,218876862 | 9,775734989 | 5,109200013 | 11,89336039 | 9,863344548 | 9,406958243 | 9,04 |
| H3f3b | 9,639815084 | 9,409819853 | 8,060149757 | 6,72056776 | 10,45702527 | 9,559354098 | 9,407406491 | 9,04 |
| Selenos | 9,44507876 | 9,482639962 | 10,37975 | 7,078134878 | 10,00276596 | 9,217227166 | 7,537345247 | 9,02 |
| Scp2 | 8,841327602 | 9,132820638 | 9,721038641 | 9,111783401 | 8,837012293 | 7,888802271 | 9,601552935 | 9,02 |
| Cdh2 | 9,455188687 | 8,44849375 | 9,106493383 | 9,587891396 | 9,278304489 | 8,986564614 | 8,241666536 | 9,01 |
| Ndufa13 | 9,478028582 | 9,251047549 | 10,13860094 | 8,864715416 | 9,303710022 | 8,184981821 | 7,873312112 | 9,01 |
| Srsf11 | 8,202462089 | 8,940800513 | 10,09604525 | 9,8095778 | 8,201635913 | 8,849190651 | 8,985112368 | 9,01 |
| Rock1 | 10,80458205 | 10,71252581 | 7,661899963 | 6,522334533 | 10,19537677 | 9,812807695 | 7,321925078 | 9,00 |
| Ywhae | 8,978198004 | 8,986394526 | 11,22471079 | 4,352532705 | 10,13143623 | 8,638568944 | 10,71756829 | 9,00 |
| Gapdh | 8,554810066 | 8,255191215 | 10,01965273 | 8,368329453 | 9,692625132 | 8,655657873 | 9,474292071 | 9,00 |
| Ktn1 | 9,394019473 | 8,826457436 | 10,2625791 | 8,617943796 | 8,2468765 | 8,948545081 | 8,710415829 | 9,00 |
| Slc15a2 | 8,677612351 | 8,058131795 | 9,013723592 | 9,847054083 | 9,497392323 | 8,363160827 | 9,533493282 | 9,00 |
| Agpat5 | 9,58626344 | 9,657199238 | 9,512052189 | 8,464248802 | 9,471848604 | 9,726796723 | 6,56540021 | 9,00 |
| Cd164 | 9,284160394 | 9,206122909 | 9,451562947 | 7,016243734 | 9,746840681 | 8,955939905 | 9,313921334 | 9,00 |
| Scrg1 | 8,791860952 | 8,659392646 | 9,557994254 | 6,935690282 | 9,080686403 | 8,828815813 | 11,10916973 | 8,99 |

| | | | | | | | | |
|---|---|---|---|---|---|---|---|---|
| Cox4i1 | 9,071603931 | 8,467717604 | 9,673606183 | 9,519515708 | 9,383324469 | 7,951605988 | 8,840268697 | 8,99 |
| Gm51419 | 7,920992689 | 9,077921019 | 11,25252451 | 8,360017101 | 8,682221163 | 8,868597995 | 8,70632253 | 8,98 |
| Tpt1 | 9,336033431 | 8,678268521 | 9,082599899 | 9,405458407 | 8,246413469 | 9,140679345 | 8,931864707 | 8,97 |
| Ctsl | 8,913295094 | 9,103922423 | 8,715848853 | 10,01873286 | 9,327921173 | 9,079785691 | 7,645042359 | 8,97 |
| Ndufa4 | 8,867415848 | 8,788423247 | 10,64991557 | 9,338023687 | 8,527544875 | 8,172174853 | 8,432239763 | 8,97 |
| Zfp207 | 7,953960747 | 8,268856682 | 10,29154062 | 10,69931003 | 8,914016946 | 8,209663929 | 8,398949504 | 8,96 |
| Atp6v0c | 9,78915102 | 9,663403737 | 7,54362019 | 8,233555174 | 10,90930997 | 8,141374451 | 8,412295715 | 8,96 |
| Ndufb11 | 8,584512592 | 8,366146189 | 9,524566954 | 9,936307442 | 8,716967593 | 7,75244378 | 9,795635678 | 8,95 |
| Ddx3y | 9,952275869 | 9,071689087 | 8,484314874 | 7,448080799 | 9,294023283 | 9,780258033 | 8,623986187 | 8,95 |
| Tecr | 8,144864822 | 8,966659525 | 9,413153611 | 7,951987402 | 9,950132729 | 8,708297364 | 9,481357843 | 8,95 |
| Dnm1l | 9,609471983 | 9,355445011 | 8,758374649 | 8,932413208 | 9,249095283 | 9,299514749 | 7,411492711 | 8,95 |
| Atp5o | 8,934556098 | 8,903035805 | 9,607938097 | 9,058772949 | 8,567014126 | 8,951901992 | 8,590600451 | 8,94 |
| Hspa2 | 10,16463183 | 9,621356842 | 8,574628544 | 9,421932751 | 8,779348825 | 8,08475556 | 7,960144768 | 8,94 |
| Cox8a | 8,468840392 | 8,453142632 | 9,619562916 | 10,19509696 | 8,537081405 | 7,732786254 | 9,59044587 | 8,94 |
| Chd9 | 9,203969974 | 9,198100611 | 8,040142202 | 9,706222058 | 9,059193247 | 8,908454458 | 8,475590446 | 8,94 |
| Tmx2 | 9,168449875 | 9,154368499 | 9,735620665 | 5,552572808 | 10,96251037 | 9,468948239 | 8,538016237 | 8,94 |
| Add3 | 8,851008069 | 8,671547829 | 9,718538474 | 9,526115866 | 9,281910772 | 8,858087017 | 7,66923552 | 8,94 |
| Pura | 9,923913142 | 9,045538728 | 9,657842483 | 9,053769414 | 7,724618545 | 9,5072804 | 7,659723591 | 8,94 |
| Plcd4 | 8,039939085 | 8,33044688 | 9,349071427 | 9,568458439 | 9,08189189 | 8,872350172 | 9,288442048 | 8,93 |
| Actr2 | 10,62421416 | 10,16238707 | 7,741948521 | 8,606695185 | 9,859076962 | 8,545104799 | 6,961531949 | 8,93 |
| Sfxn5 | 8,598146269 | 8,691444341 | 9,290193918 | 8,186531741 | 9,46959938 | 9,361681947 | 8,899318818 | 8,93 |
| Tmem151a | 10,37922902 | 9,811016592 | 7,721802776 | 10,74951432 | 8,586760384 | 7,485438798 | 7,762139724 | 8,93 |
| Selenof | 10,19153298 | 9,407941137 | 8,156961703 | 6,544653737 | 11,02140916 | 8,608120199 | 8,559824415 | 8,93 |
| Cxcl14 | 8,27334225 | 8,709250036 | 8,688241812 | 8,028929859 | 10,30734046 | 8,637868609 | 9,776937579 | 8,92 |
| Sox2 | 9,356182534 | 8,92275731 | 9,625688155 | 7,532360122 | 8,834728956 | 9,235490772 | 8,911308978 | 8,92 |
| Akap9 | 8,073528171 | 8,474213525 | 9,626124682 | 9,902452303 | 8,804034484 | 8,026920235 | 9,494584149 | 8,91 |
| Atl2 | 9,176718892 | 9,698193887 | 7,261405125 | 8,111174507 | 10,57635014 | 9,01849546 | 8,512497715 | 8,91 |
| Aco2 | 8,702542096 | 8,703962254 | 8,882838243 | 8,077054256 | 10,31946787 | 9,324639432 | 8,325878676 | 8,91 |
| Nr2f1 | 9,691267054 | 9,017956124 | 9,746222815 | 9,740581122 | 5,763118589 | 9,161184639 | 9,161774618 | 8,90 |
| Rgcc | 8,920214416 | 8,091906913 | 8,081285989 | 9,57890364 | 9,084645663 | 8,839078021 | 9,676282038 | 8,90 |
| Dag1 | 9,803389748 | 9,726506632 | 7,946738918 | 6,894311294 | 9,321294904 | 10,38348404 | 8,184797574 | 8,89 |
| Tsc22d3 | 9,681768443 | 7,627579433 | 8,60517308 | 7,780169053 | 10,41678186 | 8,569601797 | 9,533320235 | 8,89 |
| Ermn | 7,013637796 | 9,748195361 | 10,76896601 | 7,459129443 | 10,12280073 | 9,173403794 | 7,926429708 | 8,89 |

| | | | | | | | | |
|---|---|---|---|---|---|---|---|---|
| Spred1 | 9,495691524 | 8,835020944 | 10,29173799 | 6,980654959 | 9,532695409 | 9,102060815 | 7,921607349 | 8,88 |
| Psma3 | 9,021567009 | 8,600846906 | 8,35830552 | 9,229603268 | 9,199548226 | 8,707284171 | 9,018658847 | 8,88 |
| Atrx | 8,381753806 | 9,641173892 | 7,66078255 | 9,601156796 | 9,088876662 | 8,413762654 | 9,335479881 | 8,87 |
| Trpm3 | 9,112561028 | 9,278088825 | 8,40278136 | 8,832788459 | 9,145635086 | 9,544694397 | 7,800269871 | 8,87 |
| Gas7 | 9,643569287 | 9,775601769 | 9,024220548 | 10,81593174 | 5,717339031 | 8,786761798 | 8,335760744 | 8,87 |
| Itm2c | 8,615598011 | 8,336712723 | 7,650234384 | 9,670777195 | 9,340764773 | 8,682443168 | 9,792983566 | 8,87 |
| Dynlrb1 | 8,989350045 | 9,509725517 | 9,882539213 | 7,920140807 | 8,872458292 | 8,617742045 | 8,280329269 | 8,87 |
| Bsg | 9,076557019 | 9,516305195 | 9,616263791 | 6,707334335 | 10,34131035 | 8,371139844 | 8,430360657 | 8,87 |
| Prkar1a | 9,387437109 | 9,298968891 | 10,14750677 | 7,075172715 | 10,27879507 | 8,571965987 | 7,286110997 | 8,86 |
| Sfpq | 9,776382948 | 9,318294469 | 9,863062291 | 7,970877985 | 7,717548855 | 9,063726919 | 8,323585119 | 8,86 |
| Prelp | 9,060524384 | 8,400142177 | 9,472849802 | 8,07581459 | 10,85189526 | 8,402413427 | 7,752210041 | 8,86 |
| Ppp3ca | 9,138406859 | 9,093948236 | 9,183544968 | 6,617122672 | 8,886969914 | 9,732795368 | 9,362653956 | 8,86 |
| Eps15 | 9,231040423 | 8,132842026 | 8,064398163 | 8,969389773 | 9,678932312 | 9,442648814 | 8,485971354 | 8,86 |
| Mtdh | 10,11999879 | 9,727499827 | 8,506160382 | 6,802383635 | 8,104115122 | 10,06466945 | 8,660344543 | 8,86 |
| Il33 | 8,704266563 | 9,378579031 | 7,680329963 | 8,758249251 | 11,49543012 | 7,597374575 | 8,366938 | 8,85 |
| Enpp2 | 9,113230032 | 11,26140666 | 9,036798101 | 4,870361365 | 10,10811494 | 9,661838402 | 7,916501241 | 8,85 |
| Crot | 9,269602146 | 8,618806798 | 9,766698442 | 8,65167882 | 10,20178164 | 7,56102031 | 7,894498569 | 8,85 |
| Astn1 | 8,723473891 | 8,848177099 | 10,11975769 | 8,77987377 | 9,066418962 | 7,799030856 | 8,594081606 | 8,85 |
| Slc7a11 | 9,757794868 | 9,312384853 | 6,894289221 | 8,751586276 | 8,604131923 | 8,610171852 | 9,993590809 | 8,85 |
| Hmgcr | 10,25286803 | 10,47384795 | 7,202061949 | 8,725249065 | 8,894146358 | 9,7762168 | 6,598516533 | 8,85 |
| Ube3a | 9,288738257 | 8,743723295 | 8,555710661 | 10,46879574 | 8,361660766 | 9,019009368 | 7,473402129 | 8,84 |
| Nfib | 8,674385445 | 8,293262432 | 9,335684128 | 10,6019304 | 7,60553298 | 8,203366646 | 9,196677542 | 8,84 |
| Naaa | 9,91931903 | 9,215796391 | 9,400556838 | 6,901470293 | 9,744346554 | 8,912143617 | 7,807953731 | 8,84 |
| Sdc2 | 10,33587647 | 9,668677829 | 6,785025265 | 8,165838888 | 10,43009489 | 9,041079407 | 7,433767535 | 8,84 |
| Tprkb | 9,524924955 | 9,46339395 | 7,191670922 | 6,898892181 | 9,479530418 | 9,947340139 | 9,352598823 | 8,84 |
| Mat2a | 9,298432089 | 8,882638882 | 9,8658462 | 6,353113913 | 9,76560851 | 8,527139849 | 9,144380583 | 8,83 |
| Atp5pb | 9,079763623 | 8,427305254 | 9,666966544 | 8,172300623 | 9,552242318 | 7,54420802 | 9,336583842 | 8,83 |
| Dnajb9 | 8,477152931 | 9,545482979 | 8,225495699 | 7,482544934 | 10,14128746 | 9,791330064 | 8,10685753 | 8,82 |
| Atp2c1 | 8,933609612 | 9,033505634 | 9,182686603 | 8,432972396 | 9,259248661 | 9,840586597 | 7,050554612 | 8,82 |
| Map2 | 9,691803176 | 9,243831688 | 9,53312135 | 9,31421229 | 8,863184794 | 7,810439949 | 7,251134008 | 8,82 |
| Ftl1 | 7,929357129 | 8,74792523 | 9,92678173 | 6,84429237 | 9,085191387 | 7,984132223 | 11,18401208 | 8,81 |
| Ssr3 | 8,920832672 | 8,318195056 | 8,600731291 | 8,746484501 | 9,916105967 | 8,875399713 | 8,321546682 | 8,81 |
| Atp6ap2 | 10,61560882 | 10,34464342 | 8,171973477 | 6,809541391 | 8,009855425 | 10,27733603 | 7,449579501 | 8,81 |

| | | | | | | | | |
|---|---|---|---|---|---|---|---|---|
| Asah1 | 9,199969206 | 8,206955622 | 10,77698503 | 8,02115109 | 9,333420176 | 8,485616764 | 7,639336134 | 8,81 |
| Atp5h | 9,335448329 | 9,387076827 | 8,712229036 | 7,485064907 | 9,324771133 | 7,793826167 | 9,596460213 | 8,80 |
| Fads1 | 9,032466333 | 9,115119606 | 8,724264702 | 8,1903384 | 9,6125946 | 9,991792265 | 6,91569144 | 8,80 |
| Tmbim6 | 8,519730757 | 8,778094394 | 10,1084619 | 6,014576949 | 9,522376656 | 8,744936519 | 9,87996663 | 8,80 |
| Hmgn2 | 9,801865515 | 9,671124251 | 9,696689625 | 6,610351369 | 10,29146339 | 8,322398916 | 7,164148971 | 8,79 |
| Jam2 | 8,976486789 | 8,738589685 | 9,668318273 | 9,450251798 | 9,463055965 | 8,4327055 | 6,818424316 | 8,79 |
| Tlcd1 | 8,952076292 | 8,853224551 | 9,016894668 | 8,102074539 | 9,754974807 | 9,629974934 | 7,234340136 | 8,79 |
| OC11856829 | 9,290447586 | 9,252620728 | 6,911800435 | 8,58765064 | 9,565628096 | 9,473642993 | 8,461329887 | 8,79 |
| Rida | 9,671874071 | 8,940720577 | 8,99996814 | 9,929297004 | 9,456435298 | 7,670832074 | 6,873591154 | 8,79 |
| Apc | 9,496890587 | 9,427756776 | 9,386147198 | 9,097174852 | 7,324963662 | 9,455867741 | 7,346314469 | 8,79 |
| Grk3 | 8,893952598 | 9,212291026 | 9,073222351 | 6,407030722 | 10,04237765 | 9,906555 | 7,954869353 | 8,78 |
| Fmn2 | 8,487341719 | 9,271979204 | 9,302585494 | 9,548677138 | 6,968152403 | 9,02269828 | 8,849565611 | 8,78 |
| S100a16 | 7,852839352 | 7,926022363 | 9,136516886 | 9,943954688 | 8,533575429 | 7,329335327 | 10,70604078 | 8,78 |
| Elob | 8,799005886 | 9,156023967 | 9,221938693 | 9,148027034 | 7,969515526 | 8,717579235 | 8,415052157 | 8,78 |
| Clk4 | 8,837848441 | 8,550074305 | 9,22415953 | 8,498532858 | 9,08242083 | 9,215253183 | 8,006862881 | 8,77 |
| Csnk1a1 | 9,829199261 | 9,131694283 | 9,121522545 | 7,756937062 | 9,440288853 | 9,298147715 | 6,825754949 | 8,77 |
| Slc33a1 | 9,189079545 | 9,005989991 | 8,993240044 | 9,261864419 | 8,789475065 | 8,957426403 | 7,168862105 | 8,77 |
| Map1lc3b | 8,440457565 | 8,88278785 | 10,51660343 | 9,107289618 | 9,838570813 | 7,574727451 | 6,962962201 | 8,76 |
| Tmem9b | 9,332501955 | 9,191755725 | 8,437720879 | 7,138364539 | 9,987039919 | 8,774774393 | 8,421544517 | 8,75 |
| Pcdh10 | 8,960697764 | 8,996128356 | 8,67698361 | 7,472926151 | 8,37456452 | 9,367001213 | 9,43050155 | 8,75 |
| Rpl10 | 9,217391874 | 9,43966871 | 8,977999963 | 7,379838879 | 8,52056131 | 9,159698726 | 8,562326402 | 8,75 |
| Rorb | 8,804856214 | 8,975930798 | 10,51255537 | 6,970357172 | 8,851921674 | 8,530959269 | 8,57517337 | 8,75 |
| Pcdh7 | 8,708455589 | 8,329630161 | 9,067595815 | 9,41338337 | 8,313433726 | 9,536366451 | 7,846741494 | 8,75 |
| Rpl4 | 8,726359018 | 8,595351039 | 8,445508112 | 9,613408146 | 8,468946717 | 8,988523046 | 8,376881576 | 8,74 |
| Elovl5 | 8,881345288 | 9,225765044 | 10,55899425 | 5,997201836 | 7,648894827 | 9,749330925 | 9,144548594 | 8,74 |
| Atp5k | 9,851951996 | 9,517808542 | 9,051948149 | 8,155681245 | 8,862462769 | 9,376774401 | 6,367667302 | 8,74 |
| Saraf | 8,860037225 | 8,071352956 | 10,31074043 | 5,801137537 | 9,102857949 | 8,705304393 | 10,3270117 | 8,74 |
| Tial1 | 9,711348383 | 9,686798113 | 8,677942398 | 7,407896299 | 9,201249549 | 9,474238561 | 7,017715074 | 8,74 |
| Pnrc2 | 9,044929283 | 8,164820404 | 9,093925962 | 7,667808992 | 10,46303102 | 9,496789829 | 7,239171276 | 8,74 |
| Gpam | 9,49894392 | 9,4034463 | 7,578723823 | 7,932415934 | 9,616015328 | 9,678978868 | 7,455245552 | 8,74 |
| Cnn3 | 8,05247084 | 7,548514927 | 11,43469378 | 8,984183631 | 8,765744067 | 7,157412338 | 9,22065096 | 8,74 |
| Tardbp | 9,064663783 | 9,489498737 | 8,08874881 | 8,482557652 | 9,520291354 | 8,82923671 | 7,684676716 | 8,74 |
| Hsbp1 | 8,590859156 | 8,712872823 | 10,69119224 | 7,50603898 | 8,759913558 | 8,156374063 | 8,710837058 | 8,73 |

| | | | | | | | | |
|---|---|---|---|---|---|---|---|---|
| Agt | 9,003288095 | 9,402188309 | 7,968234424 | 4,956910803 | 10,94846172 | 9,707540959 | 9,134419898 | 8,73 |
| Gli3 | 8,993894563 | 8,163379476 | 10,25820214 | 6,915320019 | 8,952580382 | 8,873231685 | 8,955380909 | 8,73 |
| Rbm25 | 8,180582921 | 8,188052816 | 9,003293415 | 8,704810708 | 8,553424949 | 8,741934454 | 9,729878164 | 8,73 |
| Reep5 | 8,03157207 | 8,792862114 | 7,026723818 | 9,686198348 | 9,940384999 | 8,760427806 | 8,863067471 | 8,73 |
| Kcnk1 | 9,120236882 | 8,581890871 | 9,541917314 | 7,414511585 | 10,03510942 | 9,136352213 | 7,269588827 | 8,73 |
| Ncan | 8,626622829 | 8,674086791 | 10,24387291 | 6,509084586 | 8,672682978 | 9,202193805 | 9,14219052 | 8,72 |
| Tiparp | 8,197887074 | 9,151991109 | 9,284839455 | 8,86055956 | 9,560581653 | 7,746373645 | 8,242307346 | 8,72 |
| Rpn1 | 8,867663099 | 8,192461 | 9,494467062 | 8,980478037 | 8,771456459 | 9,159508178 | 7,541811416 | 8,72 |
| Dnajc3 | 9,280252825 | 10,05533609 | 9,122208544 | 8,036499889 | 8,068023362 | 9,344246001 | 7,078503345 | 8,71 |
| P4ha1 | 10,10049133 | 9,300706909 | 7,728666583 | 7,364743149 | 9,95741445 | 9,886573524 | 6,637810198 | 8,71 |
| Slc1a4 | 8,848044506 | 8,152137958 | 9,915004563 | 7,715214692 | 9,000568646 | 9,362179238 | 7,936088234 | 8,70 |
| Strn3 | 9,490213643 | 10,28471266 | 8,894903461 | 7,587974725 | 8,173420259 | 9,300266929 | 7,133146051 | 8,69 |
| Gnb1 | 8,31962716 | 8,024115569 | 9,790485988 | 9,077681239 | 9,128319962 | 7,743861717 | 8,779045033 | 8,69 |
| Spry2 | 9,023553654 | 8,826767195 | 8,612250626 | 9,637604226 | 7,519873916 | 9,534321651 | 7,689263746 | 8,69 |
| Bclaf1 | 9,391799297 | 8,522496682 | 8,905443443 | 9,127038415 | 8,369008437 | 8,437644628 | 8,050317049 | 8,69 |
| Eva1a | 8,637880569 | 8,506396574 | 9,426714921 | 9,286991352 | 9,217211562 | 8,767683112 | 6,956088648 | 8,69 |
| Selenow | 9,170272774 | 9,182848396 | 10,36623661 | 3,819100429 | 9,554437157 | 8,373253488 | 10,30046367 | 8,68 |
| Kif5b | 9,307599853 | 8,89049427 | 9,1486331 | 7,7783449 | 8,487737156 | 8,237748339 | 8,897725431 | 8,68 |
| Btg1 | 9,479385007 | 9,102943512 | 7,741894947 | 10,59148235 | 7,153754141 | 8,251418554 | 8,427099674 | 8,68 |
| Clcn3 | 9,589093454 | 9,352907179 | 7,464073551 | 8,187985805 | 8,467334013 | 9,064898075 | 8,606866828 | 8,68 |
| Uqcr11 | 9,189965325 | 8,747610805 | 9,730469733 | 7,819639955 | 8,702804025 | 8,432135504 | 8,077954854 | 8,67 |
| Ndufc1 | 10,27412536 | 9,384327538 | 8,70635852 | 7,952768878 | 9,869578726 | 8,804606577 | 5,664021635 | 8,67 |
| Arid4b | 8,666608038 | 8,875278308 | 8,746827088 | 8,099576461 | 9,089183763 | 8,697621836 | 8,475859024 | 8,66 |
| Fkbp3 | 8,897303908 | 8,744230262 | 10,17499234 | 9,053279873 | 8,583435645 | 6,28453126 | 8,845627496 | 8,65 |
| Uqcr10 | 8,690431657 | 8,333061218 | 10,36258731 | 9,636642551 | 8,048120336 | 7,481226901 | 8,012033076 | 8,65 |
| Eif2s2 | 9,897228754 | 9,287804382 | 7,797994454 | 8,759559707 | 8,725682876 | 8,714791028 | 7,365820087 | 8,65 |
| Slc7a10 | 8,013902426 | 7,926450327 | 8,563837643 | 8,560328943 | 8,73488938 | 8,869031175 | 9,879214807 | 8,65 |
| Sptbn1 | 10,27440424 | 9,798878695 | 6,085165876 | 7,382227113 | 9,004104493 | 9,577302171 | 8,381159624 | 8,64 |
| Uqcrh | 9,557382263 | 9,24972754 | 7,275373952 | 8,885990082 | 8,314022886 | 8,327459107 | 8,872136702 | 8,64 |
| Rpl21 | 9,287835529 | 8,631450572 | 9,632866746 | 7,454143008 | 8,678948835 | 8,336451048 | 8,452511698 | 8,64 |
| Tm9sf2 | 9,602838251 | 9,464661898 | 7,303275438 | 7,999685544 | 9,780645866 | 8,556384417 | 7,74225369 | 8,64 |
| Calm2 | 8,820059313 | 9,087111272 | 9,397366375 | 7,132797678 | 8,879392002 | 7,644051855 | 9,479506517 | 8,63 |
| Hsd11b1 | 9,108657614 | 9,342209701 | 8,99164881 | 9,992532837 | 9,279782516 | 7,616120002 | 6,099803937 | 8,63 |

| | | | | | | | | |
|---|---|---|---|---|---|---|---|---|
| Myorg | 9,073914139 | 8,753389726 | 7,76740789 | 8,350843664 | 9,036910564 | 8,840899755 | 8,596288858 | 8,63 |
| Gsk3b | 8,808120497 | 8,588765483 | 8,814739391 | 9,341435126 | 8,557919436 | 8,579776089 | 7,715834922 | 8,63 |
| Camk2g | 7,612904111 | 7,292220104 | 9,588105998 | 10,80336443 | 7,632912223 | 8,108221693 | 9,318225618 | 8,62 |
| Dnaja2 | 9,906360105 | 9,435254945 | 7,758739472 | 9,361652894 | 8,794840264 | 9,40637244 | 5,679390552 | 8,62 |
| Eci2 | 8,45693916 | 9,041412907 | 8,906212028 | 7,392972383 | 9,954188366 | 8,582432378 | 7,952826895 | 8,61 |
| Prdx2 | 9,104746878 | 8,682430528 | 8,931162739 | 7,900096173 | 9,641704644 | 8,004161324 | 8,008862577 | 8,61 |
| Rpl19 | 9,023656943 | 8,930418598 | 9,44051652 | 7,891020645 | 8,798044336 | 8,490508186 | 7,694562858 | 8,61 |
| Gm2a | 8,177592343 | 8,599762205 | 11,03546439 | 7,345046602 | 9,067400996 | 8,069873202 | 7,945551609 | 8,61 |
| Uqcrb | 8,381894026 | 7,887321504 | 9,686802572 | 10,26856402 | 7,374658819 | 7,613449218 | 9,009996679 | 8,60 |
| Nadk2 | 9,207160495 | 8,947174955 | 8,96087891 | 9,503656356 | 9,233563143 | 7,716995442 | 6,626492612 | 8,60 |
| OC11548940 | 7,988083928 | 8,323088358 | 8,626336612 | 9,550462658 | 8,392103813 | 7,618533777 | 9,672638454 | 8,60 |
| Nlgn1 | 8,905935882 | 9,009927137 | 7,248771707 | 8,563849798 | 8,60350237 | 9,127955065 | 8,711293301 | 8,60 |
| Nfat5 | 8,908376242 | 8,931392915 | 8,643008851 | 6,729916357 | 8,649836611 | 9,79062868 | 8,512883911 | 8,60 |
| Ncor1 | 8,969353146 | 9,709087187 | 8,177073999 | 7,697735334 | 7,986056683 | 9,844856319 | 7,766188119 | 8,59 |
| Gldc | 9,817232466 | 9,336967161 | 8,868705359 | 8,541089876 | 9,003063035 | 8,996998021 | 5,569502312 | 8,59 |
| Grin2c | 8,413242306 | 8,564202436 | 8,657531644 | 7,594142214 | 8,869697517 | 9,924520106 | 8,091304353 | 8,59 |
| Pou3f2 | 8,264894398 | 8,125564306 | 9,211697264 | 8,943767294 | 7,992460402 | 8,605201746 | 8,966458672 | 8,59 |
| Nipbl | 9,232499388 | 9,475787933 | 7,804876288 | 7,83322571 | 8,235960869 | 9,262833904 | 8,248332274 | 8,58 |
| Lamp2 | 9,07501135 | 9,028329933 | 7,129931887 | 9,073775762 | 8,812901147 | 8,4961215 | 8,430026324 | 8,58 |
| Cfl1 | 8,32572199 | 7,826455555 | 9,192224223 | 9,630906389 | 9,344484933 | 6,852895839 | 8,872876602 | 8,58 |
| Gm2026 | 8,208876779 | 8,571382173 | 9,245546824 | 9,291625285 | 8,948393464 | 7,421566491 | 8,358007812 | 8,58 |
| Pgk1 | 8,274258188 | 9,008154899 | 9,422039669 | 9,791645448 | 9,23232634 | 7,526843552 | 6,785697862 | 8,58 |
| Gdi2 | 8,249956119 | 8,580028838 | 9,64485087 | 7,872319197 | 8,163042001 | 9,043863932 | 8,479368397 | 8,58 |
| Nkain4 | 8,935256151 | 8,675970441 | 9,576294595 | 7,285484228 | 9,997948902 | 8,119978982 | 7,430536791 | 8,57 |
| Tob1 | 9,322237022 | 8,20263181 | 8,170864831 | 9,927709887 | 8,099687942 | 8,512565359 | 7,760045529 | 8,57 |
| Acox1 | 9,06273591 | 8,882119119 | 9,371399908 | 8,747961335 | 9,363342452 | 8,588774655 | 5,965874214 | 8,57 |
| Pcm1 | 9,299980561 | 9,005046865 | 8,101151328 | 9,417118727 | 8,449679801 | 8,684441159 | 6,99524951 | 8,56 |
| Mrps33 | 9,709121261 | 9,719092592 | 7,871968762 | 6,444862582 | 9,526918618 | 8,886546875 | 7,79157992 | 8,56 |
| Cd47 | 8,134324801 | 8,02557593 | 9,92652135 | 9,343425583 | 8,722969515 | 7,600246902 | 8,195441923 | 8,56 |
| Sash1 | 9,432578671 | 7,996140545 | 8,490369111 | 10,07830015 | 7,07494595 | 8,440771798 | 8,419468875 | 8,56 |
| ND4L | 8,836632744 | 8,853639713 | 8,891151991 | 7,050895436 | 8,834468551 | 7,174530648 | 10,28465337 | 8,56 |
| Limch1 | 8,552512644 | 8,97739961 | 9,382585778 | 5,778777534 | 9,41212701 | 9,801183028 | 8,006804021 | 8,56 |
| Mbnl1 | 8,377239797 | 9,073160972 | 8,38313737 | 10,77302734 | 7,368620372 | 8,814706051 | 7,101043681 | 8,56 |

| | | | | | | | | |
|---|---|---|---|---|---|---|---|---|
| Clcn4 | 8,381766526 | 9,255771638 | 9,033196641 | 8,301784913 | 8,487877705 | 8,697367661 | 7,712713091 | 8,55 |
| Klf9 | 10,6769448 | 9,774289141 | 4,003335118 | 7,845900982 | 9,506657024 | 9,994895003 | 8,053084121 | 8,55 |
| Atp5l | 8,952071262 | 8,890143196 | 9,206347954 | 7,878478125 | 8,387913178 | 7,545006161 | 8,972298457 | 8,55 |
| Tmem41b | 8,6604049 | 8,072888566 | 9,841274673 | 8,70149056 | 8,719852468 | 8,706606371 | 7,090915137 | 8,54 |
| Apod | 7,667622148 | 9,935383146 | 9,903953508 | 6,881454076 | 8,583041632 | 7,996030098 | 8,797160858 | 8,54 |
| Gpc5 | 8,453549967 | 8,163597064 | 5,861802288 | 9,430201929 | 8,672752506 | 7,975732786 | 11,19900285 | 8,54 |
| Zfand5 | 8,909423438 | 9,634990018 | 7,664284709 | 8,773758563 | 8,370307784 | 8,910112734 | 7,472347026 | 8,53 |
| Slc7a2 | 9,52417534 | 9,540815112 | 7,703878863 | 8,249184886 | 8,519622787 | 9,396643102 | 6,790517505 | 8,53 |
| Dner | 10,07820125 | 8,873418474 | 7,462114076 | 7,66258297 | 8,707543096 | 9,357391172 | 7,5624377 | 8,53 |

**Table S4.** List of astrocyte function genes by Borggrewe et al. 2021.

| **Cholesterol_synthesis (PMID: 29279367)** | **Blood_brain_barrier** | **Lactate_metabolism** | **Myelination** |
|---|---|---|---|
| Hmgcs1 | Kcnj10 | Slc16a1 | Cntf |
| Hmgcr | Aqp4 | Slc16a3 | Lif |
| Mvk | Slc2a1 | Ldha | Fgf2 |
| Pmvk | Abcb1b | Ldhb | Slc40a1 |
| Mvd | Vegfa | Ldhc | Scap |
| Fdps | Gdnf | Slc2a1 | Pdgfa |
| Idi1 | Fgf2 | Gsk3a | Igf1 |
| Fdft1 | Angpt1 | Gsk3b | Ntf3 |
| Sqle | P2rx1 | Pygb | |
| Lss | Pld2 | Glo1 | |
| Cyp51 | Ptgs1 | Glul | |
| Msmo1 | Gja1 | Slc16a11 | |
| Nsdhl | Slc16a6 | Slc16a12 | |
| Sc5d | Slc16a7 | Slc16a13 | |
| Dhcr7 | Slc16a8 | Slc16a4 | |
| Hsd17b7 | Slc5a12 | Slc16a5 | |
| Acat2 | | | |
| Tm7sf2 | | | |
| Dhcr24 | | | |

**Table S5.** List of astrocyte subtypes genes by Batiuk et al. 2020.

| **AST1** | **AST2** | **AST3** | **AST4** | **AST5** |
|---|---|---|---|---|
| Gfap | Uncl13c | Gfap | Frzb | Fam107a |
| Agt | Slc1a3 | Slc1a3 | Ascl1 | Slc1a3 |
| Slc1a3 | | | Slc1a3 | |

**Table S6.** Differentially expressed genes between Y-A+ and Y-A+/G+ astrocytes.

| Gene | Entrez_ID | baseMean | nonNorm_log2FoldChange | nonNorm_lfcSE | pvalue | padj |
|---|---|---|---|---|---|---|
| Sesn3 | 75747 | 69,874756 | -24,64846744 | 2,059598828 | 5,25E-33 | 5,22E-29 |
| Coa6 | 67892 | 75,7412459 | -25,88614517 | 2,223124385 | 2,46E-31 | 1,22E-27 |
| Galnt11 | 231050 | 33,8332754 | -22,84439224 | 2,124445582 | 5,73E-27 | 1,90E-23 |
| Gfpt1 | 14583 | 50,159263 | -24,19706307 | 2,333823971 | 3,47E-25 | 8,62E-22 |
| Mad2l2 | 71890 | 49,2458473 | -23,73899895 | 2,308569469 | 8,41E-25 | 1,67E-21 |
| Acaa2 | 52538 | 26331,3434 | -24,45513108 | 2,394201642 | 1,71E-24 | 2,84E-21 |
| Coq6 | 217707 | 711,27833 | -26,42748988 | 2,623295159 | 7,19E-24 | 1,02E-20 |
| Rpain | 69723 | 50,9926279 | -23,50453267 | 2,338326852 | 9,02E-24 | 1,12E-20 |
| 3230217C12Ril | 68127 | 76,9594703 | -24,98391257 | 2,536312998 | 6,82E-23 | 7,54E-20 |
| Gstm6 | 14867 | 8691,58278 | -24,15065127 | 2,498883011 | 4,26E-22 | 4,24E-19 |
| Heph | 15203 | 23,0999121 | -22,52670017 | 2,374055965 | 2,34E-21 | 2,12E-18 |
| Adrm1 | 56436 | 198,938858 | -21,20079817 | 2,243468659 | 3,39E-21 | 2,81E-18 |
| Nat8f5 | 69049 | 110,971415 | -23,44352798 | 2,484884414 | 3,93E-21 | 3,01E-18 |
| Tmem176a | 66058 | 3794,47614 | -22,39745264 | 2,382270824 | 5,37E-21 | 3,81E-18 |
| Nbdy | 70123 | 45,0489264 | -22,04182356 | 2,350934341 | 6,87E-21 | 4,31E-18 |
| Tsr1 | 104662 | 68,5586335 | -23,65141356 | 2,522858222 | 6,93E-21 | 4,31E-18 |
| Gm17750 | 553095 | 8205,04951 | -23,71901244 | 2,543895754 | 1,12E-20 | 6,56E-18 |
| 9330159M07Ril | 319673 | 44,5171768 | -23,39533378 | 2,52802357 | 2,15E-20 | 1,19E-17 |
| Alkbh7 | 66400 | 39,8052381 | -23,1719187 | 2,556711383 | 1,27E-19 | 6,63E-17 |
| Mvd | 192156 | 23,0550399 | -21,6210044 | 2,429884055 | 5,69E-19 | 2,83E-16 |
| Tcof1 | 21453 | 34,6942802 | -19,94491939 | 2,267285214 | 1,41E-18 | 6,66E-16 |
| Ccnq | 69109 | 35,3834834 | -21,98586064 | 2,546619778 | 5,96E-18 | 2,69E-15 |
| Timm9 | 30056 | 15,1395592 | -22,0465398 | 2,569641767 | 9,52E-18 | 4,12E-15 |
| Tap2 | 21355 | 14,3839848 | -20,09494251 | 2,57633471 | 6,20E-15 | 2,57E-12 |
| Pkdcc | 106522 | 123,424918 | -20,16765981 | 2,598440059 | 8,40E-15 | 3,34E-12 |
| BC065397 | 436230 | 18,0999261 | -19,6529186 | 2,537378051 | 9,53E-15 | 3,65E-12 |
| Gm527 | 217648 | 18,3005583 | -20,17898736 | 2,625466816 | 1,52E-14 | 5,60E-12 |
| Mgat4b | 103534 | 24,6650985 | -19,24295692 | 2,552188783 | 4,71E-14 | 1,67E-11 |
| Cenpw | 66311 | 10,566935 | -19,6486529 | 2,748808034 | 8,80E-13 | 3,02E-10 |

| | | | | | | |
|---|---|---|---|---|---|---|
| Gm27239 | 102641244 | 81,9299513 | -17,67505194 | 2,481188092 | 1,05E-12 | 3,48E-10 |
| Slc37a4 | 14385 | 18,9611654 | -17,97216269 | 2,562715228 | 2,33E-12 | 7,49E-10 |
| Zc4h2 | 245522 | 14,1868869 | -20,66691273 | 3,060554701 | 1,45E-11 | 4,51E-09 |
| C920006O11Ril | 320295 | 12,6846404 | -20,59335162 | 3,077465913 | 2,21E-11 | 6,65E-09 |
| Gm29508 | 102632488 | 17,0284709 | -19,42927238 | 2,938690758 | 3,80E-11 | 1,11E-08 |
| Zfp128 | 243833 | 11,4419924 | -19,77775723 | 3,076123131 | 1,28E-10 | 3,64E-08 |
| Ankrd13d | 68423 | 11,8499242 | -19,66225078 | 3,067861102 | 1,46E-10 | 4,04E-08 |
| C1qa | 12259 | 17,4012236 | 18,25462033 | 2,864750688 | 1,86E-10 | 5,01E-08 |
| Gm39468 | 105243583 | 19,6590805 | 25,11417901 | 3,961840295 | 2,31E-10 | 6,05E-08 |
| Gal3st4 | 330217 | 17,1479168 | -18,36460948 | 2,930650405 | 3,70E-10 | 9,42E-08 |
| Rgs1 | 50778 | 42,0004473 | -23,80120709 | 3,952159259 | 1,72E-09 | 4,27E-07 |
| Nudt19 | 110959 | 356,515407 | -12,12273943 | 2,217698844 | 4,59E-08 | 1,11E-05 |
| Ranbp9 | 56705 | 117,065708 | -9,531526459 | 1,752327969 | 5,35E-08 | 1,27E-05 |
| Alg1 | 208211 | 19,9370055 | -21,35195507 | 3,953606176 | 6,64E-08 | 1,54E-05 |
| Zfyve16 | 218441 | 73,1236878 | -8,884771475 | 1,661547324 | 8,93E-08 | 2,02E-05 |
| Glrx5 | 73046 | 2234,82064 | -20,481206 | 3,853316314 | 1,07E-07 | 2,35E-05 |
| Dhx40 | 67487 | 78,9412539 | -9,247541423 | 1,756062786 | 1,39E-07 | 3,01E-05 |
| Gm38504 | 102637959 | 32,3518827 | 11,99114507 | 2,28156537 | 1,47E-07 | 3,12E-05 |
| Zfp57 | 22715 | 305,33732 | 7,105945694 | 1,361660303 | 1,80E-07 | 3,74E-05 |
| Zfp639 | 67778 | 103,952184 | -9,777059369 | 1,876459251 | 1,88E-07 | 3,78E-05 |
| Fam162a | 70186 | 80,6856021 | -9,643499019 | 1,851346905 | 1,90E-07 | 3,78E-05 |
| Vtn | 22370 | 552,32815 | 8,984796801 | 1,741089022 | 2,46E-07 | 4,80E-05 |
| Pabpn1 | 54196 | 32,6892704 | -20,42912599 | 4,013883815 | 3,59E-07 | 6,86E-05 |
| Gm36298 | 102640165 | 19,5455076 | -20,29041031 | 3,992943901 | 3,74E-07 | 7,02E-05 |
| Decr1 | 67460 | 150,845819 | -8,486123254 | 1,706460583 | 6,59E-07 | 0,000121 |
| St8sia5 | 225742 | 24,2889551 | -19,70712266 | 4,014557416 | 9,16E-07 | 0,000166 |
| Gm26788 | 108167617 | 87,8030699 | -19,3790863 | 3,964180544 | 1,02E-06 | 0,00018 |
| Alkbh3 | 69113 | 65,12643 | -8,931396175 | 1,846079657 | 1,31E-06 | 0,000229 |
| Washc3 | 67282 | 142,10164 | -9,068970587 | 1,883872233 | 1,48E-06 | 0,000254 |
| Ramac | 67148 | 125,308495 | -9,656022299 | 2,009825786 | 1,55E-06 | 0,000262 |
| Gm36742 | 102640746 | 134,315251 | -10,2514473 | 2,154250796 | 1,95E-06 | 0,000323 |
| Capn2 | 12334 | 70,5828111 | -10,01031561 | 2,112265071 | 2,15E-06 | 0,00035 |
| Ccnt1 | 12455 | 520,506617 | 8,863278231 | 1,883779215 | 2,54E-06 | 0,000407 |

| | | | | | | |
|---|---|---|---|---|---|---|
| Bckdhb | 12040 | 80,3946114 | -9,019423839 | 1,928829215 | 2,92E-06 | 0,000462 |
| Ncbp2 | 68092 | 118,869871 | -9,026945 | 1,949630343 | 3,66E-06 | 0,000568 |
| Gramd3 | 107022 | 122,777991 | -7,353764006 | 1,599839581 | 4,30E-06 | 0,000657 |
| Asf1a | 66403 | 58,0135851 | -9,588964991 | 2,103982672 | 5,18E-06 | 0,00078 |
| Exosc3 | 66362 | 64,1444148 | -8,669214759 | 1,910354527 | 5,68E-06 | 0,000836 |
| Gorab | 98376 | 439,114405 | 8,912771592 | 1,96458496 | 5,71E-06 | 0,000836 |
| Gulp1 | 70676 | 52,6454463 | -7,92625295 | 1,752698536 | 6,12E-06 | 0,000882 |
| E130311K13Ri[illegible] | 329659 | 58,3996387 | -9,620484595 | 2,136637828 | 6,71E-06 | 0,000954 |
| Psme2 | 19188 | 80,2362373 | -8,774806709 | 1,954543218 | 7,14E-06 | 0,001 |
| Atp13a5 | 268878 | 283,133282 | 8,673314028 | 1,960802449 | 9,72E-06 | 0,001342 |
| Tor2a | 30933 | 66,5245635 | -7,403709823 | 1,677102196 | 1,01E-05 | 0,001379 |
| 4833439L19Ri[illegible] | 97820 | 77,0019836 | -8,030315806 | 1,838283901 | 1,25E-05 | 0,001682 |
| Rnf34 | 80751 | 77,209464 | -9,745774714 | 2,239994795 | 1,36E-05 | 0,001739 |
| Jagn1 | 67767 | 56,8317584 | -8,095352621 | 1,858290324 | 1,32E-05 | 0,001739 |
| Gm867 | 333670 | 156,550035 | 8,151157698 | 1,872699095 | 1,35E-05 | 0,001739 |
| Pde9a | 18585 | 44,1748317 | -7,634020279 | 1,755115588 | 1,36E-05 | 0,001739 |
| Brix1 | 67832 | 71,566121 | -8,708052667 | 2,006258148 | 1,42E-05 | 0,00179 |
| Lpar6 | 67168 | 124,217597 | -8,553949239 | 1,972116032 | 1,44E-05 | 0,001792 |
| Tmem218 | 66279 | 48,1995765 | -8,545682926 | 1,984185043 | 1,66E-05 | 0,001984 |
| Usp37 | 319651 | 82,5823709 | -8,196749939 | 1,903157884 | 1,66E-05 | 0,001984 |
| Slc33a1 | 11416 | 294,427843 | -6,778031369 | 1,573220073 | 1,64E-05 | 0,001984 |
| Eno4 | 226265 | 752,721333 | 8,728716673 | 2,030698518 | 1,72E-05 | 0,002037 |
| Mcu | 215999 | 54,4236336 | -7,555079521 | 1,762626665 | 1,82E-05 | 0,002126 |
| N4bp2 | 333789 | 109,777311 | -7,655427759 | 1,788273047 | 1,86E-05 | 0,002152 |
| Tmsb10 | 19240 | 175,720902 | 8,054880885 | 1,884759575 | 1,92E-05 | 0,002198 |
| Slc8b1 | 170756 | 49,7366107 | -8,062623004 | 1,889172963 | 1,97E-05 | 0,002231 |
| Nepro | 212547 | 39,2130142 | -8,196014709 | 1,92555414 | 2,08E-05 | 0,002318 |
| Kctd12 | 239217 | 140,879676 | -7,741465075 | 1,819711526 | 2,10E-05 | 0,002318 |
| Bambi | 68010 | 92,9116434 | -7,452674571 | 1,755197086 | 2,18E-05 | 0,002357 |
| Tlk1 | 228012 | 55,1886235 | -7,421399497 | 1,748031101 | 2,18E-05 | 0,002357 |
| Dr1 | 13486 | 98,429642 | -9,479556994 | 2,240571335 | 2,33E-05 | 0,002489 |
| Dapp1 | 26377 | 51,4109351 | -8,013442352 | 1,896583961 | 2,39E-05 | 0,002526 |
| Agbl3 | 76223 | 57,3128192 | -8,141037889 | 1,929402107 | 2,45E-05 | 0,002564 |

| | | | | | | |
|---|---|---|---|---|---|---|
| Tppp3 | 67971 | 14927,802 | 8,893801163 | 2,115593005 | 2,62E-05 | 0,002717 |
| Map2k7 | 26400 | 46,540771 | -7,894773055 | 1,879505903 | 2,66E-05 | 0,002731 |
| Ifnar2 | 15976 | 135,320125 | -8,802829635 | 2,097083725 | 2,70E-05 | 0,002737 |
| Anapc10 | 68999 | 46,0186704 | -9,334497414 | 2,226088528 | 2,75E-05 | 0,002763 |
| 1700019D03Ril | 67080 | 72,2261044 | -8,507855844 | 2,03640811 | 2,94E-05 | 0,002926 |
| Eaf2 | 106389 | 55,1937407 | -9,547407818 | 2,295383957 | 3,19E-05 | 0,003142 |
| Rpp14 | 67053 | 348,750917 | -9,633719498 | 2,320469771 | 3,30E-05 | 0,003219 |
| Entr1 | 68112 | 46,4723275 | -8,12930748 | 1,960437675 | 3,37E-05 | 0,003257 |
| Gm46406 | 108168072 | 27,9285517 | -10,79682144 | 2,618035836 | 3,72E-05 | 0,00356 |
| Lcat | 16816 | 994,180602 | -6,081516196 | 1,485659765 | 4,25E-05 | 0,004014 |
| Peli1 | 67245 | 226,903916 | -5,74313765 | 1,403535971 | 4,28E-05 | 0,004014 |
| St7l | 229681 | 64,2566662 | -7,088971402 | 1,743303453 | 4,77E-05 | 0,004438 |
| Dner | 227325 | 215,553168 | -6,970871126 | 1,719269838 | 5,02E-05 | 0,004625 |
| Dmac2l | 68055 | 50,8042654 | -9,223140571 | 2,282775347 | 5,34E-05 | 0,004804 |
| Zmat1 | 215693 | 141,221986 | -8,896450475 | 2,202492462 | 5,36E-05 | 0,004804 |
| Sc5d | 235293 | 257,359648 | -6,09846865 | 1,508993118 | 5,31E-05 | 0,004804 |
| Gle1 | 74412 | 37,8398336 | -7,660978529 | 1,90123387 | 5,59E-05 | 0,004964 |
| Uhrf2 | 109113 | 31,8101454 | -8,662384625 | 2,154769004 | 5,82E-05 | 0,00512 |
| Pcgf6 | 71041 | 60,0629514 | -8,104107623 | 2,020673972 | 6,06E-05 | 0,005238 |
| Dpagt1 | 13478 | 77,665973 | -7,028635185 | 1,751943271 | 6,02E-05 | 0,005238 |
| F8 | 14069 | 18,2938322 | -9,195752585 | 2,295329933 | 6,17E-05 | 0,005288 |
| Tmem29 | 382245 | 193,243415 | -7,814364932 | 1,951741556 | 6,23E-05 | 0,005299 |
| Syap1 | 67043 | 45,6105225 | -8,789380225 | 2,199492751 | 6,44E-05 | 0,005427 |
| Ppp3r1 | 19058 | 34,4724752 | -8,519112424 | 2,1394708 | 6,84E-05 | 0,005676 |
| Rps6ka5 | 73086 | 68,0978305 | -7,816837304 | 1,963296363 | 6,85E-05 | 0,005676 |
| Herc4 | 67345 | 37,5633439 | -7,44979649 | 1,872705306 | 6,95E-05 | 0,00571 |
| Sptssa | 104725 | 149,421926 | -6,841974855 | 1,728025235 | 7,51E-05 | 0,006125 |
| Lyn | 17096 | 31,8481792 | -7,613133732 | 1,92397818 | 7,59E-05 | 0,006137 |
| Atf1 | 11908 | 27,0602825 | -7,121944025 | 1,802754814 | 7,80E-05 | 0,006252 |
| Rab11a | 53869 | 158,169995 | -6,771987657 | 1,715957861 | 7,93E-05 | 0,00631 |
| Calcrl | 54598 | 100,58661 | -7,094489165 | 1,80087016 | 8,17E-05 | 0,006394 |
| Gid4 | 66771 | 57,3797181 | -5,826732423 | 1,478994694 | 8,16E-05 | 0,006394 |
| Mrps36 | 66128 | 62,7317631 | -6,329892311 | 1,608759739 | 8,33E-05 | 0,006474 |

| | | | | | | |
|---|---|---|---|---|---|---|
| Mrpl46 | 67308 | 46,0865544 | -9,502533098 | 2,419265892 | 8,57E-05 | 0,006476 |
| Zfp30 | 22693 | 41,4030158 | -7,993203597 | 2,036316651 | 8,66E-05 | 0,006476 |
| Slc46a3 | 71706 | 48,9293462 | -8,211848421 | 2,091400731 | 8,62E-05 | 0,006476 |
| Bcar3 | 29815 | 59,7515268 | -6,695333396 | 1,703846057 | 8,51E-05 | 0,006476 |
| Timm22 | 56322 | 58,448314 | -7,416733642 | 1,88845541 | 8,59E-05 | 0,006476 |
| Chic1 | 12212 | 89,6574869 | -5,330603722 | 1,359509445 | 8,82E-05 | 0,006545 |
| Nsmaf | 18201 | 56,3083106 | -7,468891413 | 1,91190731 | 9,36E-05 | 0,006823 |
| Tapt1 | 231225 | 62,8385241 | -6,959121001 | 1,780919424 | 9,32E-05 | 0,006823 |
| Zkscan17 | 268417 | 123,794605 | 7,45832307 | 1,909653115 | 9,40E-05 | 0,006823 |
| Osgin2 | 209212 | 102,893985 | -7,596354979 | 1,947726534 | 9,61E-05 | 0,006929 |
| Elf2 | 69257 | 47,3625649 | -6,837527039 | 1,754620499 | 9,74E-05 | 0,006972 |
| Gm4631 | 102633156 | 51,4868885 | -7,050345376 | 1,810659078 | 9,87E-05 | 0,00701 |
| Tmem50b | 77975 | 151,123607 | -6,401910487 | 1,647184758 | 0,000102 | 0,007171 |
| Tnip1 | 57783 | 45,9880306 | -9,634528398 | 2,482439635 | 0,000104 | 0,007283 |
| Polr2j | 20022 | 53,3233445 | -8,119387185 | 2,096933331 | 0,000108 | 0,007506 |
| Lifr | 16880 | 222,443324 | -6,182160258 | 1,598077787 | 0,00011 | 0,007563 |
| Cmtm6 | 67213 | 67,4043009 | -8,205974671 | 2,124233396 | 0,000112 | 0,007681 |
| Smc5 | 226026 | 209,830958 | -7,78320654 | 2,017058068 | 0,000114 | 0,007765 |
| Naprt | 223646 | 35,4830818 | -8,077963047 | 2,102174954 | 0,000122 | 0,007963 |
| Nek6 | 59126 | 38,6267352 | -7,68970728 | 1,99709676 | 0,000118 | 0,007963 |
| Mgat2 | 217664 | 52,0973287 | -7,132362815 | 1,855594212 | 0,000121 | 0,007963 |
| Zfp788 | 67607 | 43,0712953 | -6,951603838 | 1,808674895 | 0,000121 | 0,007963 |
| Btf3l4 | 70533 | 64,0486991 | -6,963954023 | 1,81208453 | 0,000122 | 0,007963 |
| Usp42 | 76800 | 35,4182304 | -5,828642036 | 1,516148808 | 0,000121 | 0,007963 |
| Rabggtb | 19352 | 34,8423053 | -7,875577843 | 2,054623686 | 0,000127 | 0,008214 |
| Ucp2 | 22228 | 48,2305453 | -9,0352621 | 2,357953427 | 0,000127 | 0,008214 |
| Cand2 | 67088 | 111,52521 | -8,215533759 | 2,149445144 | 0,000132 | 0,008488 |
| Dnah3 | 381917 | 53,7806745 | 5,715842929 | 1,496574468 | 0,000134 | 0,008532 |
| Zfpm2 | 22762 | 30,2596339 | -6,928540902 | 1,815272031 | 0,000135 | 0,008563 |
| Rbbp9 | 26450 | 49,5471686 | -6,887337823 | 1,80524004 | 0,000136 | 0,008565 |
| Cerox1 | 72834 | 133,167099 | 7,82566572 | 2,058672894 | 0,000144 | 0,009002 |
| Pigm | 67556 | 64,2709697 | -7,013172687 | 1,846382732 | 0,000146 | 0,009055 |
| Stard3 | 59045 | 28,089765 | -6,709339534 | 1,770022944 | 0,00015 | 0,009285 |

| | | | | | | |
|---|---|---|---|---|---|---|
| Polr2e | 66420 | 55,0976594 | -7,246338946 | 1,912590888 | 0,000151 | 0,009295 |
| Stam2 | 56324 | 64,4084504 | -7,727056564 | 2,04120719 | 0,000153 | 0,009358 |
| C030037D09Ril | 193280 | 29,3603172 | -7,585776676 | 2,006435811 | 0,000156 | 0,009376 |
| Galnt6 | 207839 | 21,142137 | 7,902215172 | 2,090251117 | 0,000157 | 0,009376 |
| Nab1 | 17936 | 59,0560289 | -6,188458499 | 1,636304391 | 0,000156 | 0,009376 |
| Ndufs4 | 17993 | 151,648885 | -6,70390561 | 1,776676877 | 0,000161 | 0,009594 |
| Marchf2 | 224703 | 51,5547787 | -6,940568727 | 1,840851572 | 0,000163 | 0,009652 |
| Mrpl21 | 353242 | 53,3165798 | -7,851064174 | 2,087093967 | 0,000169 | 0,009872 |
| Septin1 | 54204 | 35,6436971 | 8,211958845 | 2,182340681 | 0,000168 | 0,009872 |
| Scnm1 | 69269 | 508,216986 | 6,966453088 | 1,858768278 | 0,000178 | 0,010371 |
| Ptp4a3 | 19245 | 37,6480657 | 6,677074825 | 1,783722162 | 0,000182 | 0,0105 |
| 1110008P14Rik | 73737 | 26,4700594 | -8,645473179 | 2,320184936 | 0,000194 | 0,010799 |
| Naga | 17939 | 27,7043702 | -7,389554042 | 1,981601308 | 0,000192 | 0,010799 |
| Saal1 | 78935 | 57,2348112 | -8,271719892 | 2,220702258 | 0,000195 | 0,010799 |
| Mrpl22 | 216767 | 79,8365957 | -7,896057619 | 2,119543232 | 0,000195 | 0,010799 |
| Cklf | 75458 | 42,3612588 | -7,582115178 | 2,035087602 | 0,000195 | 0,010799 |
| Gm16523 | 100042584 | 72,6673649 | -6,837381535 | 1,832525168 | 0,000191 | 0,010799 |
| Kdm3b | 277250 | 33,3928083 | -5,989848116 | 1,604663281 | 0,000189 | 0,010799 |
| Cdc123 | 98828 | 47,9783577 | -9,51577385 | 2,551835344 | 0,000192 | 0,010799 |
| Tgfbr1 | 21812 | 59,9149986 | -6,772636346 | 1,819165432 | 0,000197 | 0,01082 |
| Med23 | 70208 | 46,8888821 | -8,370753643 | 2,251876276 | 0,000201 | 0,011006 |
| Acyp2 | 75572 | 55,1834083 | -6,559550265 | 1,766393347 | 0,000204 | 0,011107 |
| Necap2 | 66147 | 36,4623446 | -6,092145894 | 1,641729149 | 0,000207 | 0,011166 |
| Slc39a11 | 69806 | 58,2011947 | -9,08489175 | 2,451634936 | 0,000211 | 0,011335 |
| Tmem101 | 76547 | 32,5794672 | -8,812355626 | 2,381492669 | 0,000215 | 0,011385 |
| Gm35116 | 102638587 | 38,4540663 | -7,451758064 | 2,014476392 | 0,000216 | 0,011385 |
| Tbk1 | 56480 | 166,084852 | 5,96854147 | 1,612198745 | 0,000214 | 0,011385 |
| Ginm1 | 215751 | 239,881013 | -6,537047415 | 1,767188375 | 0,000216 | 0,011385 |
| Sgf29 | 75565 | 41,6871579 | -8,630910186 | 2,334280154 | 0,000218 | 0,011398 |
| Cdc37 | 12539 | 42,0733072 | -7,39680272 | 2,003346646 | 0,000222 | 0,011455 |
| Aph1b | 208117 | 99,4919413 | -6,770100729 | 1,833203137 | 0,000222 | 0,011455 |
| Rnf121 | 75212 | 58,213175 | -8,675209137 | 2,348278711 | 0,000221 | 0,011455 |
| Ngly1 | 59007 | 79,5132461 | -7,69222398 | 2,085475409 | 0,000226 | 0,011513 |

| | | | | | | |
|---|---|---|---|---|---|---|
| Inpp1 | 16329 | 77,4384977 | -6,093174238 | 1,65202219 | 0,000226 | 0,011513 |
| Cnih1 | 12793 | 116,575229 | -5,920195805 | 1,606462037 | 0,000228 | 0,011593 |
| Ccdc138 | 76138 | 46,5800456 | -7,766886504 | 2,114777309 | 0,00024 | 0,011704 |
| Golph3l | 229593 | 61,6604562 | -7,036443685 | 1,915923731 | 0,00024 | 0,011704 |
| Cep57 | 74360 | 84,4100964 | -6,611592489 | 1,796213224 | 0,000232 | 0,011704 |
| Tomm70a | 28185 | 37,7457589 | -6,809090044 | 1,852358713 | 0,000237 | 0,011704 |
| Ppp6c | 67857 | 127,472417 | -6,001625386 | 1,631715451 | 0,000235 | 0,011704 |
| Tfam | 21780 | 37,2267148 | -6,534488553 | 1,779100536 | 0,00024 | 0,011704 |
| 0610040J01Rik | 76261 | 29,0908396 | -8,622630665 | 2,347008554 | 0,000239 | 0,011704 |
| Rab9 | 56382 | 109,930264 | -8,285689629 | 2,255139244 | 0,000239 | 0,011704 |
| Ctif | 269037 | 225,760939 | 5,814701154 | 1,586145233 | 0,000246 | 0,011955 |
| Fbxo30 | 71865 | 94,4600618 | -6,601372335 | 1,806220912 | 0,000257 | 0,012412 |
| Col5a3 | 53867 | 114,635851 | 8,305367857 | 2,273054066 | 0,000258 | 0,012412 |
| Dcaf7 | 71833 | 62,9242597 | -6,656307405 | 1,825323962 | 0,000266 | 0,012703 |
| Ndc1 | 72787 | 27,4731605 | -7,187986153 | 1,972340291 | 0,000268 | 0,01271 |
| Cdr1 | 631990 | 100,51556 | 6,254166656 | 1,71627082 | 0,000268 | 0,01271 |
| Iqca | 74918 | 106,548431 | 9,11376892 | 2,503033692 | 0,000271 | 0,012796 |
| H2aw | 319162 | 33,0252217 | -8,890096821 | 2,443108343 | 0,000274 | 0,012847 |
| Eif3g | 53356 | 85,5415283 | -7,704630168 | 2,119007228 | 0,000277 | 0,012931 |
| Cops3 | 26572 | 80,3326964 | -5,618910638 | 1,547361896 | 0,000282 | 0,013103 |
| Pola1 | 18968 | 129,064437 | 13,34519029 | 3,676232607 | 0,000283 | 0,013103 |
| Slc25a16 | 73132 | 50,5434936 | -8,092687288 | 2,230626603 | 0,000286 | 0,013151 |
| Snrpe | 20643 | 89,1556491 | -7,89039531 | 2,190443829 | 0,000316 | 0,013793 |
| Cinp | 67236 | 18,9477093 | -7,725295347 | 2,143443595 | 0,000313 | 0,013793 |
| Med7 | 66213 | 41,250424 | -7,342570788 | 2,035739623 | 0,00031 | 0,013793 |
| Usp25 | 30940 | 62,6440805 | -6,909512056 | 1,914151345 | 0,000307 | 0,013793 |
| Herc6 | 67138 | 55,2368717 | -6,837321138 | 1,89719411 | 0,000313 | 0,013793 |
| D030047H15Ril | 100037396 | 33,2191084 | -6,898660813 | 1,912667301 | 0,00031 | 0,013793 |
| Ilrun | 224647 | 76,8422653 | 6,745185113 | 1,872813824 | 0,000316 | 0,013793 |
| Chmp7 | 105513 | 25,9267842 | -7,194134977 | 1,995845483 | 0,000313 | 0,013793 |
| Xpo5 | 72322 | 24,371346 | -6,127701455 | 1,700030807 | 0,000313 | 0,013793 |
| Erg28 | 58520 | 109,8795 | -6,19788611 | 1,716373879 | 0,000305 | 0,013793 |
| Mphosph10 | 67973 | 68,8506605 | -5,607498086 | 1,552741071 | 0,000305 | 0,013793 |

| | | | | | | |
|---|---|---|---|---|---|---|
| Zbed6 | 667118 | 49,8983844 | 5,815492618 | 1,612912248 | 0,000311 | 0,013793 |
| BC064078 | 408064 | 28,9179434 | -8,793565977 | 2,444302652 | 0,000321 | 0,013949 |
| Arhgap18 | 73910 | 36,6541456 | -7,075917022 | 1,968791772 | 0,000326 | 0,014025 |
| Slc39a10 | 227059 | 69,4251501 | -5,336999942 | 1,485018589 | 0,000326 | 0,014025 |
| Agtrap | 11610 | 35,9276326 | -7,662486294 | 2,135640609 | 0,000333 | 0,014054 |
| Lin52 | 217708 | 36,226206 | -7,70458769 | 2,146305505 | 0,000331 | 0,014054 |
| Lats1 | 16798 | 32,9275743 | -7,419191911 | 2,066111543 | 0,00033 | 0,014054 |
| Catsper2 | 212670 | 43,9535982 | -7,477973062 | 2,083860388 | 0,000333 | 0,014054 |
| Mthfsl | 100039707 | 69,7613173 | -5,951079387 | 1,658706422 | 0,000334 | 0,014054 |
| Vps13b | 666173 | 76,0793366 | -6,090555155 | 1,699452467 | 0,000339 | 0,01418 |
| Ddx47 | 67755 | 38,4210562 | -8,848024738 | 2,469270573 | 0,000339 | 0,01418 |
| Gabpa | 14390 | 64,9279593 | -7,666729767 | 2,142006451 | 0,000345 | 0,01434 |
| Angel2 | 52477 | 116,572148 | -5,30953074 | 1,484730795 | 0,000349 | 0,014452 |
| Parp2 | 11546 | 108,730345 | -6,39186016 | 1,787946668 | 0,00035 | 0,014454 |
| Tomm22 | 223696 | 68,0253319 | -7,155072938 | 2,003278194 | 0,000355 | 0,014576 |
| Dpm2 | 13481 | 48,2334202 | -8,426121719 | 2,363224273 | 0,000363 | 0,014862 |
| Arl6 | 56297 | 49,2520686 | -7,309988222 | 2,053481915 | 0,000371 | 0,015127 |
| Mfsd8 | 72175 | 99,748978 | -8,251710689 | 2,320888445 | 0,000377 | 0,015319 |
| Alms1-ps2 | 623273 | 16,1031228 | -7,547313343 | 2,124852568 | 0,000382 | 0,015431 |
| Rb1 | 19645 | 213,826637 | -6,123124119 | 1,724166147 | 0,000383 | 0,015431 |
| Med28 | 66999 | 52,5100025 | -6,649482065 | 1,873900052 | 0,000387 | 0,015538 |
| Tmem8b | 242409 | 166,314246 | 5,886434894 | 1,659389856 | 0,000389 | 0,015542 |
| Cnpy4 | 66455 | 61,5847209 | -7,332600791 | 2,068617618 | 0,000393 | 0,015637 |
| Smpd1 | 20597 | 336,270264 | -6,067326744 | 1,71348356 | 0,000399 | 0,015798 |
| Galnt7 | 108150 | 38,839826 | -6,16575762 | 1,74388246 | 0,000407 | 0,016053 |
| Borcs7 | 66439 | 31,0421817 | -6,409989908 | 1,814271855 | 0,000411 | 0,016081 |
| Pfn2 | 18645 | 155,785801 | -6,339642717 | 1,79413218 | 0,00041 | 0,016081 |
| 1300002E11Ril | 100043489 | 103,705797 | -6,754366362 | 1,913650729 | 0,000416 | 0,016131 |
| Hibadh | 58875 | 58,2115381 | -6,481199304 | 1,836342755 | 0,000416 | 0,016131 |
| Mlst8 | 56716 | 182,881636 | 7,533647757 | 2,134681177 | 0,000417 | 0,016131 |
| Pik3r4 | 75669 | 27,5757262 | -6,508498618 | 1,845075428 | 0,00042 | 0,016171 |
| Nfu1 | 56748 | 24,5782495 | -7,313771663 | 2,075360396 | 0,000425 | 0,016316 |
| Ncald | 52589 | 57,403402 | -7,055591192 | 2,006245289 | 0,000437 | 0,016706 |

| | | | | | | |
|---|---|---|---|---|---|---|
| Gbp7 | 229900 | 77,4622186 | -6,254685717 | 1,781686142 | 0,000447 | 0,016808 |
| Igfbp5 | 16011 | 61,7027886 | -6,289960544 | 1,791787245 | 0,000447 | 0,016808 |
| Ptprt | 19281 | 47,3749964 | -5,169910439 | 1,473271679 | 0,00045 | 0,016808 |
| Tdp2 | 56196 | 115,062329 | -7,110291383 | 2,024265074 | 0,000444 | 0,016808 |
| Bbs7 | 71492 | 274,592671 | 8,141886757 | 2,319371974 | 0,000447 | 0,016808 |
| Phtf1 | 18685 | 174,789572 | 12,72906907 | 3,626888545 | 0,000449 | 0,016808 |
| Arsa | 11883 | 33,3614885 | -7,83250187 | 2,235250666 | 0,000458 | 0,017002 |
| Pomt1 | 99011 | 48,4909517 | -8,84986829 | 2,52528454 | 0,000457 | 0,017002 |
| Nmd3 | 97112 | 29,2579868 | -7,630529485 | 2,181492672 | 0,000469 | 0,017324 |
| Wrnip1 | 78903 | 29,4919738 | -4,90879033 | 1,403669902 | 0,00047 | 0,017324 |
| Hddc3 | 68695 | 80,4693908 | -6,159374574 | 1,762467201 | 0,000475 | 0,017413 |
| Uchl5 | 56207 | 74,8464391 | -7,336004726 | 2,102515236 | 0,000485 | 0,017651 |
| Prrg1 | 546336 | 65,8952344 | -5,519144035 | 1,581446527 | 0,000483 | 0,017651 |
| Pld3 | 18807 | 74,0684652 | -5,983299659 | 1,719292838 | 0,000501 | 0,018193 |
| Drg1 | 13494 | 78,9218504 | 7,637314736 | 2,196521728 | 0,000507 | 0,018293 |
| Ppp1r32 | 67752 | 296,857672 | 13,5003315 | 3,883113524 | 0,000508 | 0,018293 |
| Cops4 | 26891 | 91,9702423 | -6,494165713 | 1,868459602 | 0,00051 | 0,018295 |
| Lrrc59 | 98238 | 56,859125 | -8,814897635 | 2,537130072 | 0,000512 | 0,018319 |
| Dexi | 58239 | 42,1844649 | -7,068644448 | 2,037193106 | 0,000521 | 0,018368 |
| Sfmbt1 | 54650 | 16,7777222 | -7,052074128 | 2,031313128 | 0,000517 | 0,018368 |
| Chd1 | 12648 | 68,5457565 | -6,010333017 | 1,732112678 | 0,000521 | 0,018368 |
| Pak5 | 241656 | 60,5491136 | 8,520319458 | 2,454720032 | 0,000519 | 0,018368 |
| Cfap100 | 243538 | 229,820741 | 5,706226783 | 1,645036593 | 0,000523 | 0,018375 |
| Slc4a8 | 59033 | 4127,95911 | 6,902130563 | 1,994491103 | 0,000539 | 0,018808 |
| Dlg1 | 13383 | 113,639755 | -5,968489433 | 1,724448499 | 0,000538 | 0,018808 |
| C87436 | 232196 | 18,2772503 | -6,162837292 | 1,782930369 | 0,000547 | 0,019024 |
| Abca8a | 217258 | 42,1763479 | -6,820976182 | 1,974597373 | 0,000552 | 0,019114 |
| Fars2 | 69955 | 71,3612273 | -6,29466743 | 1,822805472 | 0,000554 | 0,019124 |
| Aspscr1 | 68938 | 80,6141931 | 5,540169207 | 1,605667959 | 0,00056 | 0,019263 |
| Gm30908 | 102632966 | 35,7273191 | -7,058267866 | 2,048825459 | 0,000571 | 0,019581 |
| Slc38a10 | 72055 | 53,3444555 | -6,822904912 | 1,984786814 | 0,000587 | 0,019786 |
| Leng1 | 69757 | 31,0800169 | -5,740894316 | 1,670036717 | 0,000587 | 0,019786 |
| Cabin1 | 104248 | 166,28781 | 4,890995943 | 1,42229645 | 0,000584 | 0,019786 |

| | | | | | | |
|---|---|---|---|---|---|---|
| Cdan1 | 68968 | 1742,19716 | -7,805294331 | 2,269288994 | 0,000583 | 0,019786 |
| Cideb | 12684 | 32,1023386 | -8,433998563 | 2,45188637 | 0,000582 | 0,019786 |
| Dpm1 | 13480 | 65,9962473 | -6,073434074 | 1,768133974 | 0,000593 | 0,019913 |
| Sft2d3 | 67158 | 38,3905086 | -6,664087415 | 1,941582102 | 0,000598 | 0,02004 |
| Igf2 | 16002 | 84,0928836 | 6,155721854 | 1,794581554 | 0,000603 | 0,020063 |
| Arhgef28 | 110596 | 80,1956713 | 7,477384598 | 2,179752705 | 0,000603 | 0,020063 |
| Zfp12 | 231866 | 34,3413918 | -7,416174213 | 2,163153234 | 0,000607 | 0,020126 |
| Nr1d2 | 353187 | 223,881618 | -6,876017162 | 2,00617064 | 0,000609 | 0,020132 |
| Lyrm1 | 73919 | 16,2852667 | -6,448943677 | 1,883126631 | 0,000616 | 0,020209 |
| Mospd1 | 70380 | 82,523131 | 6,796893182 | 1,984459214 | 0,000615 | 0,020209 |
| Arel1 | 68497 | 31,6848469 | -7,345002828 | 2,145763783 | 0,000619 | 0,020259 |
| Lrig2 | 269473 | 446,396297 | 6,315028965 | 1,845626678 | 0,000622 | 0,020298 |
| Slc30a9 | 109108 | 95,33804 | -7,572253538 | 2,214833223 | 0,000629 | 0,020352 |
| Zfp825 | 235956 | 24,5031152 | -6,805421463 | 1,991434244 | 0,000632 | 0,020352 |
| Zyg11b | 414872 | 52,0268486 | -5,512902683 | 1,613175036 | 0,000632 | 0,020352 |
| Arf4 | 11843 | 221,938854 | -5,750674548 | 1,681936513 | 0,000628 | 0,020352 |
| Mapk1ip1 | 69546 | 33,7999973 | -7,693470706 | 2,255228731 | 0,000646 | 0,020394 |
| Rab29 | 226422 | 12,6135083 | -6,846709204 | 2,005831739 | 0,000642 | 0,020394 |
| Atp13a1 | 170759 | 49,5338861 | -6,67235724 | 1,955222581 | 0,000644 | 0,020394 |
| Wdr77 | 70465 | 26,0102984 | -7,116461695 | 2,085038763 | 0,000642 | 0,020394 |
| Pdpn | 14726 | 116,450572 | -5,438354725 | 1,592746073 | 0,000639 | 0,020394 |
| C330018D20Ril | 77422 | 51,5406905 | -8,103934863 | 2,37604429 | 0,000648 | 0,020394 |
| Mcat | 223722 | 85,100317 | 8,409443283 | 2,464837528 | 0,000645 | 0,020394 |
| Ssbp2 | 66970 | 210,946433 | -5,265643546 | 1,5459933 | 0,000659 | 0,020682 |
| Spats2l | 67198 | 28,9659674 | -7,128202649 | 2,094902777 | 0,000667 | 0,020871 |
| Sox21 | 223227 | 116,367477 | -6,009744929 | 1,770385811 | 0,000687 | 0,021412 |
| Shtn1 | 71653 | 70,7008541 | 7,245577589 | 2,134865553 | 0,000689 | 0,021412 |
| Gm41212 | 105245822 | 19,3925211 | -7,529762946 | 2,219655216 | 0,000693 | 0,021458 |
| Cyb5d1 | 327951 | 45,9994227 | -5,291916229 | 1,560284685 | 0,000695 | 0,021458 |
| Rgl2 | 19732 | 16,3602859 | -7,35511815 | 2,170128459 | 0,000701 | 0,021534 |
| Ddx51 | 69663 | 65,7899874 | -6,358670888 | 1,876281442 | 0,000702 | 0,021534 |
| Hnrnpll | 72692 | 75,9743252 | -6,379006055 | 1,882947531 | 0,000705 | 0,021561 |
| Rab3il1 | 74760 | 87,4115239 | -6,745789313 | 1,992810846 | 0,000712 | 0,021608 |

| | | | | | | |
|---|---|---|---|---|---|---|
| Ahnak | 66395 | 218,031879 | 6,178307117 | 1,825805747 | 0,000715 | 0,021608 |
| Pip4k2c | 117150 | 23,8345966 | -6,304383369 | 1,863051635 | 0,000715 | 0,021608 |
| Cct2 | 12461 | 199,874864 | -5,921650915 | 1,749984217 | 0,000715 | 0,021608 |
| Nipa1 | 233280 | 33,9970446 | -6,636452104 | 1,965652443 | 0,000735 | 0,022148 |
| Magi2 | 50791 | 102,654576 | -6,597840114 | 1,954933428 | 0,000738 | 0,022181 |
| Efs | 13644 | 23,3062116 | -5,900023844 | 1,750465251 | 0,00075 | 0,022472 |
| Zmym1 | 68310 | 64,9367504 | 6,983936475 | 2,072730703 | 0,000753 | 0,022495 |
| Dnlz | 52838 | 64,5161976 | -5,447374543 | 1,61716448 | 0,000756 | 0,022506 |
| Jak2 | 16452 | 55,9605428 | -6,003019996 | 1,784365474 | 0,000768 | 0,022786 |
| Cluap1 | 76779 | 34,4981446 | -6,650878747 | 1,977861048 | 0,000772 | 0,022809 |
| Txnl1 | 53382 | 39,0180482 | -7,447499134 | 2,214993492 | 0,000773 | 0,022809 |
| Cand1 | 71902 | 78,8434473 | -5,364870684 | 1,59663522 | 0,000779 | 0,022924 |
| Adrb1 | 11554 | 22,9545478 | -5,688963902 | 1,696941075 | 0,000801 | 0,023496 |
| Pikfyve | 18711 | 56,2141153 | -5,900952933 | 1,760762492 | 0,000804 | 0,023507 |
| Fam89b | 17826 | 27,7435154 | -7,784312292 | 2,323179577 | 0,000806 | 0,023507 |
| Ino80c | 225280 | 44,8606394 | -7,07033184 | 2,113573431 | 0,000822 | 0,023676 |
| Pop7 | 74097 | 34,8733102 | -7,419890664 | 2,216554675 | 0,000815 | 0,023676 |
| Tenm3 | 23965 | 464,135051 | -5,2926012 | 1,582384755 | 0,000824 | 0,023676 |
| Glrb | 14658 | 393,145774 | -5,278122841 | 1,577662557 | 0,000821 | 0,023676 |
| Rnase1 | 19752 | 60,8944849 | -7,157440956 | 2,139682721 | 0,000823 | 0,023676 |
| Sec24c | 218811 | 54,7509615 | -6,191748012 | 1,852669921 | 0,000832 | 0,023833 |
| Vbp1 | 22327 | 176,625002 | -6,839016087 | 2,047689546 | 0,000838 | 0,023953 |
| Sorcs2 | 81840 | 80,9651214 | -5,94125604 | 1,781657852 | 0,000854 | 0,024335 |
| Faap20 | 67513 | 59,4069345 | -7,097738762 | 2,131488838 | 0,000869 | 0,024611 |
| Ptgs1 | 19224 | 75,4806138 | -6,704620338 | 2,013126695 | 0,000867 | 0,024611 |
| Ccny | 67974 | 17,6179704 | -6,254035375 | 1,879160383 | 0,000874 | 0,024704 |
| Scfd2 | 212986 | 34,9891761 | -8,126140415 | 2,442752686 | 0,000879 | 0,024765 |
| Tfb1m | 224481 | 64,9009416 | -8,069263653 | 2,426900754 | 0,000884 | 0,024788 |
| 2510002D24Rik | 72307 | 470,21798 | 7,372481019 | 2,217429005 | 0,000885 | 0,024788 |
| Pcgf5 | 76073 | 98,9437702 | -6,218680495 | 1,871184498 | 0,000889 | 0,024843 |
| Aqp11 | 66333 | 31,2150636 | -7,872087455 | 2,369697337 | 0,000894 | 0,024899 |
| Snapc1 | 75627 | 82,3091821 | -6,144869239 | 1,850609456 | 0,000899 | 0,024904 |
| Serpinb1a | 66222 | 65,6701979 | -7,313472555 | 2,202608032 | 0,000899 | 0,024904 |

| | | | | | | |
|---|---|---|---|---|---|---|
| Ints14 | 69882 | 27,4696345 | 8,186221469 | 2,467353674 | 0,000907 | 0,025063 |
| Lypla1 | 18777 | 55,0237579 | -6,558155782 | 1,983123026 | 0,000943 | 0,02598 |
| Thoc7 | 66231 | 62,1562991 | -5,092637918 | 1,54356459 | 0,000969 | 0,026631 |
| Rbsn | 78287 | 91,1486101 | -6,012200679 | 1,822867184 | 0,000973 | 0,026647 |
| Casc3 | 192160 | 54,8415414 | -5,466573772 | 1,657769685 | 0,000975 | 0,026647 |
| Entpd6 | 12497 | 29,3294567 | -6,540506197 | 1,984441475 | 0,000981 | 0,026731 |
| Slc25a27 | 74011 | 155,103898 | 5,36962249 | 1,63492318 | 0,001022 | 0,027626 |
| Pias1 | 56469 | 192,796959 | -4,878734568 | 1,485307078 | 0,001021 | 0,027626 |
| Sh3bp5l | 79566 | 120,215634 | 6,601148009 | 2,009739704 | 0,001021 | 0,027626 |
| Fam120aos | 68128 | 174,951245 | -5,406423033 | 1,647236523 | 0,00103 | 0,027768 |
| Cabcoco1 | 73287 | 144,364321 | 7,580282291 | 2,310805134 | 0,001037 | 0,027866 |
| Fam181b | 58238 | 138,511572 | -6,600883704 | 2,013304386 | 0,001043 | 0,027962 |
| Ppp6r1 | 243819 | 16,6062731 | -7,514010416 | 2,293281695 | 0,001051 | 0,028084 |
| Egln1 | 112405 | 21,8052956 | -7,011654189 | 2,140383666 | 0,001053 | 0,028084 |
| Ccdc191 | 212153 | 63,4329429 | -6,483458515 | 1,98337717 | 0,00108 | 0,028656 |
| Lrrc4b | 272381 | 101,880721 | -4,698691297 | 1,437494836 | 0,001081 | 0,028656 |
| Tbca | 21371 | 257,745531 | -5,84120782 | 1,788383215 | 0,00109 | 0,028831 |
| Mir22hg | 100042498 | 50,5185007 | -6,47142215 | 1,985327491 | 0,001116 | 0,029429 |
| Bace2 | 56175 | 165,739405 | 6,460136213 | 1,984610806 | 0,001133 | 0,029768 |
| Golt1b | 66964 | 44,9877579 | -5,236792651 | 1,609280472 | 0,001137 | 0,029768 |
| Rbm43 | 71684 | 18,7544245 | -7,488553169 | 2,300875771 | 0,001135 | 0,029768 |
| Krt10 | 16661 | 66,0390512 | -6,056440266 | 1,864268865 | 0,001159 | 0,030262 |
| Gm3055 | 100040944 | 31,9346497 | -7,628581996 | 2,353148321 | 0,001188 | 0,030916 |
| Foxd1 | 15229 | 17,4389073 | 8,719853126 | 2,692408961 | 0,001201 | 0,031009 |
| Twnk | 226153 | 71,2667021 | 6,511525215 | 2,010949826 | 0,001204 | 0,031009 |
| Letmd1 | 68614 | 47,9356053 | -5,539428464 | 1,710692341 | 0,001203 | 0,031009 |
| 3230311B06Ril | 381914 | 29,617506 | -7,81083769 | 2,412190938 | 0,001203 | 0,031009 |
| Celf4 | 108013 | 122,993668 | 7,999987691 | 2,471216784 | 0,001207 | 0,031012 |
| Il1rapl1 | 331461 | 82,0142538 | -4,867298677 | 1,504229816 | 0,001213 | 0,031098 |
| Rab11fip4 | 268451 | 31,3641167 | -5,481943265 | 1,694909172 | 0,001219 | 0,031104 |
| Msl3 | 17692 | 22,2710836 | -6,993110568 | 2,162715953 | 0,001223 | 0,031104 |
| Cox16 | 66272 | 75,6369433 | -5,888471731 | 1,820832741 | 0,001221 | 0,031104 |
| Dennd4a | 102442 | 97,2679773 | -5,127386783 | 1,586153348 | 0,001227 | 0,031122 |

| | | | | | | |
|---|---|---|---|---|---|---|
| Stx5a | 56389 | 42,7320585 | -5,569562794 | 1,731382183 | 0,001296 | 0,0328 |
| Polr1c | 20016 | 51,0804577 | -6,926952844 | 2,156037069 | 0,001314 | 0,033125 |
| Mospd2 | 76763 | 29,5201807 | -7,423201087 | 2,310696693 | 0,001316 | 0,033125 |
| Nsa2 | 59050 | 110,685502 | -4,893819588 | 1,523924363 | 0,001321 | 0,033181 |
| Dclre1c | 227525 | 49,5624845 | -5,842073312 | 1,82300355 | 0,001352 | 0,033622 |
| Naa20 | 67877 | 61,4437946 | -5,183793017 | 1,617405914 | 0,001351 | 0,033622 |
| Mtm1 | 17772 | 30,0903454 | -7,061569237 | 2,202531187 | 0,001345 | 0,033622 |
| Zswim7 | 69747 | 35,9081013 | -7,829901059 | 2,443056668 | 0,001351 | 0,033622 |
| Gm46430 | 108168101 | 71,3800426 | -6,141603524 | 1,917516642 | 0,001361 | 0,033719 |
| Rcan3 | 53902 | 13,6291835 | -8,623541373 | 2,692862242 | 0,001363 | 0,033719 |
| Pcx | 18563 | 111,601408 | -5,757881253 | 1,801105392 | 0,001389 | 0,033865 |
| Sema6a | 20358 | 96,0186317 | -4,39153044 | 1,372808643 | 0,001379 | 0,033865 |
| Tmem129 | 68366 | 72,2695404 | -6,089713129 | 1,904580374 | 0,001387 | 0,033865 |
| Vps25 | 28084 | 43,45409 | -7,514991783 | 2,34894938 | 0,001378 | 0,033865 |
| Chmp2b | 68942 | 116,457517 | -5,920078883 | 1,850236551 | 0,001376 | 0,033865 |
| Gpx8 | 69590 | 109,176688 | -7,504193696 | 2,346415499 | 0,001383 | 0,033865 |
| Ip6k1 | 27399 | 205,226042 | -5,653963935 | 1,769226084 | 0,001395 | 0,033915 |
| Thyn1 | 77862 | 57,9547609 | -8,453540275 | 2,646414678 | 0,001402 | 0,033996 |
| Smim8 | 66291 | 36,5008941 | -5,097365579 | 1,597222945 | 0,001416 | 0,034184 |
| Trim30d | 209387 | 19,1998645 | -7,521190397 | 2,356754702 | 0,001416 | 0,034184 |
| Aopep | 72061 | 98,221146 | 6,306059675 | 1,976661241 | 0,001421 | 0,034229 |
| Cog2 | 76332 | 26,6980774 | -6,803599637 | 2,133954181 | 0,001431 | 0,034383 |
| Sdhd | 66925 | 137,526893 | -5,182711858 | 1,626515097 | 0,001441 | 0,034522 |
| Ethe1 | 66071 | 73,785581 | -6,090073021 | 1,912936968 | 0,001454 | 0,034565 |
| Asb8 | 78541 | 95,8851415 | -5,46663996 | 1,716363148 | 0,001447 | 0,034565 |
| Tyw5 | 68736 | 62,5403002 | -6,510978698 | 2,045384106 | 0,001456 | 0,034565 |
| Cetn2 | 26370 | 712,27652 | 5,628687773 | 1,76763256 | 0,001451 | 0,034565 |
| Taf4 | 228980 | 37,0675188 | -6,507376922 | 2,044832929 | 0,001461 | 0,034591 |
| Alg6 | 320438 | 37,5237164 | -6,23448186 | 1,960842278 | 0,001475 | 0,034678 |
| Mark2 | 13728 | 72,4352321 | 4,295070918 | 1,350541194 | 0,001471 | 0,034678 |
| Raver1 | 71766 | 16,1657264 | -7,842108898 | 2,466497962 | 0,001476 | 0,034678 |
| Cfap300 | 234912 | 171,502139 | 12,34365014 | 3,883026278 | 0,001478 | 0,034678 |
| Bcat2 | 12036 | 67,238003 | -5,094160151 | 1,605315447 | 0,001507 | 0,035039 |

| | | | | | | |
|---|---|---|---|---|---|---|
| Hnrnpc | 15381 | 168,431692 | -4,430250098 | 1,396168929 | 0,001508 | 0,035039 |
| Mcm4 | 17217 | 93,6032561 | -6,698980435 | 2,110011222 | 0,001499 | 0,035039 |
| Idh3g | 15929 | 84,1823721 | -7,241376368 | 2,28143631 | 0,001503 | 0,035039 |
| Smug1 | 71726 | 19,6371139 | -6,266886439 | 1,97759346 | 0,00153 | 0,035467 |
| Slc19a2 | 116914 | 49,0903106 | -6,054001909 | 1,911045697 | 0,001535 | 0,035512 |
| Dusp22 | 105352 | 53,8606859 | -5,685980933 | 1,795423557 | 0,001541 | 0,035548 |
| Gm43305 | 108168162 | 21,3901046 | 8,055809317 | 2,545052324 | 0,001549 | 0,035667 |
| P2rx4 | 18438 | 163,098534 | 5,898081502 | 1,864130619 | 0,001556 | 0,035743 |
| Alg3 | 208624 | 25,9080053 | -6,87371816 | 2,173817242 | 0,001567 | 0,035899 |
| Xpo4 | 57258 | 44,3158976 | -5,231612721 | 1,655624313 | 0,001578 | 0,036061 |
| Retreg3 | 67998 | 52,0679111 | -7,632231333 | 2,415720966 | 0,001581 | 0,036061 |
| Erich1 | 234086 | 101,785873 | 7,771877994 | 2,461436001 | 0,001592 | 0,036219 |
| Ppil3 | 70225 | 19,4641932 | -6,078241728 | 1,925743888 | 0,001598 | 0,036279 |
| 5031425E22Rik | 269630 | 44,1531052 | -6,098060413 | 1,93415039 | 0,001617 | 0,036575 |
| Ireb2 | 64602 | 63,7732115 | -6,003888692 | 1,904820825 | 0,001622 | 0,036575 |
| Farsb | 23874 | 37,6932211 | -6,376351669 | 2,022904394 | 0,001621 | 0,036575 |
| Toe1 | 68276 | 19,3102369 | -6,912703566 | 2,19657181 | 0,001649 | 0,03711 |
| Rragc | 54170 | 18,4695304 | -6,907020685 | 2,195304542 | 0,001654 | 0,037124 |
| Smc2 | 14211 | 30,8702284 | -6,715289231 | 2,137437572 | 0,001679 | 0,037617 |
| Syt10 | 54526 | 124,507392 | 6,648354615 | 2,119803867 | 0,001711 | 0,03815 |
| Fbxo22 | 71999 | 527,768778 | -6,792885024 | 2,16584804 | 0,001711 | 0,03815 |
| Eif4enif1 | 74203 | 55,8934421 | -6,071668707 | 1,93672234 | 0,001718 | 0,038232 |
| Slc25a37 | 67712 | 25,367877 | -5,52146217 | 1,762254321 | 0,001729 | 0,038387 |
| Cep68 | 216543 | 43,79174 | -6,741086421 | 2,152638268 | 0,001739 | 0,038515 |
| Strn | 268980 | 98,2258458 | -5,880239075 | 1,878557312 | 0,001747 | 0,038535 |
| Kbtbd3 | 69149 | 40,4159476 | -6,194357726 | 1,980614694 | 0,001763 | 0,038535 |
| Celrr | 98452 | 19,9567575 | -6,661331629 | 2,13022212 | 0,001766 | 0,038535 |
| Zfp62 | 22720 | 198,275944 | -4,954825914 | 1,583413769 | 0,001753 | 0,038535 |
| Csgalnact1 | 234356 | 173,367471 | -4,931768647 | 1,576128379 | 0,001754 | 0,038535 |
| Azi2 | 27215 | 36,0216653 | -5,557379023 | 1,776968051 | 0,001763 | 0,038535 |
| Gm19935 | 100503868 | 193,540501 | 7,218745647 | 2,308634152 | 0,001767 | 0,038535 |
| Dnajb9 | 27362 | 225,862248 | -4,846023453 | 1,551888723 | 0,001792 | 0,039002 |
| Ube2k | 53323 | 110,34931 | -5,54356262 | 1,778542644 | 0,001828 | 0,039602 |

| | | | | | | |
|---|---|---|---|---|---|---|
| Dag1 | 13138 | 314,380825 | -4,342787901 | 1,393314897 | 0,001828 | 0,039602 |
| Nus1 | 52014 | 48,500298 | -6,03338257 | 1,936380892 | 0,001834 | 0,03966 |
| Hgf | 15234 | 97,232297 | -6,469388421 | 2,077827304 | 0,001849 | 0,039804 |
| Atad5 | 237877 | 58,7958951 | -7,147997151 | 2,295841972 | 0,001849 | 0,039804 |
| Atg4c | 242557 | 44,0021632 | -6,354231402 | 2,042657249 | 0,001866 | 0,040081 |
| Ccdc84 | 382073 | 14,8930098 | -6,42341563 | 2,065403689 | 0,001871 | 0,040098 |
| Nrp1 | 18186 | 105,936968 | -5,016607536 | 1,613707573 | 0,001879 | 0,040183 |
| Armc9 | 78795 | 68,410885 | -5,554461856 | 1,788777191 | 0,001902 | 0,040584 |
| Zfp738 | 408068 | 11,563403 | -8,701035034 | 2,805381589 | 0,001925 | 0,040996 |
| Acad9 | 229211 | 60,0745076 | -6,081605208 | 1,961681737 | 0,001934 | 0,041095 |
| Prrt1 | 260297 | 23,5301108 | -6,130701136 | 1,979432652 | 0,001954 | 0,041162 |
| Coa8 | 68020 | 21,7154163 | -5,71784145 | 1,845909229 | 0,001951 | 0,041162 |
| E2f3 | 13557 | 83,2672155 | -5,257280327 | 1,697129475 | 0,00195 | 0,041162 |
| Gk | 14933 | 111,002058 | -6,037263926 | 1,949015623 | 0,001951 | 0,041162 |
| Alg2 | 56737 | 63,1307519 | -6,507733812 | 2,102101712 | 0,001963 | 0,041266 |
| Dph7 | 67228 | 15,0481661 | -7,297081348 | 2,358522562 | 0,001975 | 0,041357 |
| Dmxl2 | 235380 | 57,6094647 | -5,489854655 | 1,774110545 | 0,001972 | 0,041357 |
| Slc31a1 | 20529 | 84,445771 | -5,473237278 | 1,771048859 | 0,001999 | 0,041697 |
| Taf9 | 108143 | 31,2365345 | -5,957297663 | 1,927779815 | 0,002 | 0,041697 |
| Slc9a7 | 236727 | 79,3508961 | -6,221334907 | 2,015199053 | 0,00202 | 0,042037 |
| Cox6b2 | 333182 | 50,8335625 | 6,219243619 | 2,016761488 | 0,002044 | 0,042394 |
| Lrrcc1 | 71710 | 392,898076 | 5,751913187 | 1,865790924 | 0,00205 | 0,042394 |
| Ypel2 | 77864 | 362,509493 | 4,443321777 | 1,441156323 | 0,002048 | 0,042394 |
| Zdhhc12 | 66220 | 29,0427428 | -7,440704443 | 2,41592817 | 0,002071 | 0,042731 |
| Stk38l | 232533 | 52,5689572 | -4,70579339 | 1,528275576 | 0,002076 | 0,042743 |
| Rasa3 | 19414 | 71,7080566 | -5,802275092 | 1,885801111 | 0,002092 | 0,04299 |
| Noc3l | 57753 | 20,7666353 | -5,690016664 | 1,850698429 | 0,002108 | 0,043233 |
| Mrpl2 | 27398 | 38,9894699 | -7,172772292 | 2,334132203 | 0,002119 | 0,043366 |
| Gm35254 | 102638770 | 75,4479849 | -5,98057125 | 1,948013858 | 0,00214 | 0,0437 |
| Washc5 | 223593 | 11,2118214 | -6,551875016 | 2,13631094 | 0,002163 | 0,044077 |
| Mbd3 | 17192 | 37,4353486 | -5,715689231 | 1,864201173 | 0,002169 | 0,044117 |
| Plekhb2 | 226971 | 301,839937 | 5,9505627 | 1,941786257 | 0,00218 | 0,044255 |
| Bnip3 | 12176 | 48,1228438 | -5,488849911 | 1,791509351 | 0,002185 | 0,044263 |

| | | | | | | |
|---|---|---|---|---|---|---|
| Ppp2r3c | 59032 | 63,9555858 | -5,906563773 | 1,928652941 | 0,002195 | 0,044273 |
| Arl1 | 104303 | 149,125952 | -5,347616896 | 1,746085051 | 0,002194 | 0,044273 |
| Hmces | 232210 | 110,366014 | 7,456277332 | 2,435582368 | 0,002203 | 0,044352 |
| Mrps26 | 99045 | 37,5301892 | -7,499698528 | 2,450966475 | 0,002214 | 0,044484 |
| Lpcat2 | 270084 | 15,6077008 | -7,089177474 | 2,318223453 | 0,002228 | 0,044673 |
| 4930522L14Rik | 100041734 | 25,6549978 | -6,60142051 | 2,160069641 | 0,002242 | 0,044868 |
| Gask1a | 245050 | 56,3750326 | -6,916206164 | 2,263945038 | 0,002251 | 0,044888 |
| Mgst1 | 56615 | 204,615982 | -6,061107245 | 1,984142135 | 0,002252 | 0,044888 |
| Trp53rka | 381406 | 16,5994643 | -7,193137181 | 2,358043375 | 0,002285 | 0,045446 |
| Rbpms2 | 71973 | 12,852507 | -7,258228485 | 2,381057207 | 0,002301 | 0,045589 |
| Mfsd9 | 211798 | 42,4636972 | -7,460840761 | 2,447207952 | 0,002298 | 0,045589 |
| Zfp512b | 269401 | 24,4820926 | -5,512486755 | 1,808909266 | 0,002308 | 0,045637 |
| Fgf4 | 14175 | 18,7336529 | -4,940825498 | 1,622619171 | 0,002327 | 0,045918 |
| Tm2d3 | 68634 | 113,031122 | -6,507503979 | 2,137747411 | 0,002334 | 0,04596 |
| Elovl6 | 170439 | 68,4782698 | -5,065561143 | 1,66479293 | 0,002344 | 0,045986 |
| Pxmp4 | 59038 | 19,7992214 | -5,310358993 | 1,745259524 | 0,002344 | 0,045986 |
| Brd9 | 105246 | 1839,61331 | 4,662556846 | 1,533621477 | 0,002364 | 0,046098 |
| 1810014B01Rik | 66263 | 16,945864 | -6,694067827 | 2,201724124 | 0,002363 | 0,046098 |
| Slmap | 83997 | 150,92831 | -5,641480304 | 1,855018284 | 0,002356 | 0,046098 |
| Arpc3 | 56378 | 23,4191586 | -5,74208674 | 1,889722669 | 0,002377 | 0,046259 |
| Dedd2 | 67379 | 179,523167 | 6,687889597 | 2,203329526 | 0,002403 | 0,046666 |
| Pcmt1 | 18537 | 134,022496 | -4,98659563 | 1,644580883 | 0,002428 | 0,047075 |
| Slc39a13 | 68427 | 16,5008431 | -5,419085458 | 1,788044945 | 0,00244 | 0,047202 |
| Tmed7 | 66676 | 596,851871 | -4,96724468 | 1,639406433 | 0,002446 | 0,04724 |
| Map4k5 | 399510 | 45,4049955 | -5,545471104 | 1,831433126 | 0,002462 | 0,047456 |
| Alpl | 11647 | 89,2563145 | -4,713761309 | 1,557351806 | 0,002472 | 0,047546 |
| Nell2 | 54003 | 14,155993 | -7,342848504 | 2,427172768 | 0,002484 | 0,047603 |
| Cdk5rap1 | 66971 | 148,948062 | 6,909586517 | 2,283972115 | 0,002484 | 0,047603 |
| Rcc1 | 100088 | 25,6686152 | -8,212090991 | 2,715520892 | 0,002493 | 0,047688 |
| Rgl1 | 19731 | 51,9764862 | -6,398927731 | 2,116680825 | 0,002502 | 0,04776 |
| Zfp938 | 237411 | 49,8641026 | -4,790253284 | 1,585802768 | 0,002522 | 0,047861 |
| Rest | 19712 | 138,390358 | -4,794537306 | 1,587091796 | 0,00252 | 0,047861 |
| Hspa1a | 193740 | 265,86313 | 5,128832222 | 1,697677496 | 0,002519 | 0,047861 |

| | | | | | | |
|---|---|---|---|---|---|---|
| Rnasek | 52898 | 193,765931 | -6,079059656 | 2,013436294 | 0,002534 | 0,048002 |
| Ndufaf5 | 69487 | 13,554355 | -5,633211301 | 1,868027934 | 0,002565 | 0,048131 |
| Brip1os | 74038 | 45,5749839 | -5,5006666 | 1,823833797 | 0,002561 | 0,048131 |
| Ddx42 | 72047 | 58,721696 | -4,895467265 | 1,622500971 | 0,002551 | 0,048131 |
| Ier2 | 15936 | 316,670892 | 6,516313538 | 2,159519776 | 0,002549 | 0,048131 |
| Gpx7 | 67305 | 20,3240776 | -8,030072368 | 2,662878805 | 0,002565 | 0,048131 |
| Fzd9 | 14371 | 34,3898915 | -6,687100768 | 2,218446427 | 0,002576 | 0,048238 |
| Vps4a | 116733 | 42,6880152 | -5,57546822 | 1,851752917 | 0,002605 | 0,048573 |
| Gm39941 | 105244306 | 166,680214 | 4,98575772 | 1,655634372 | 0,002601 | 0,048573 |
| Wipf1 | 215280 | 29,4397324 | -7,018875293 | 2,331462485 | 0,002608 | 0,048573 |
| 1700025G04Rik | 69399 | 14,4680936 | -5,769388437 | 1,917259082 | 0,002619 | 0,048693 |
| 1700066M21Rik | 73467 | 22,7728871 | -6,853068596 | 2,280543884 | 0,002656 | 0,049274 |
| Coro7 | 78885 | 14,8603802 | -7,386367888 | 2,45880036 | 0,002664 | 0,049338 |
| 2410004P03Rik | 73667 | 138,260203 | 6,29683294 | 2,098601454 | 0,002695 | 0,049431 |
| Fgf11 | 14166 | 40,8999002 | -6,246867148 | 2,080610792 | 0,002678 | 0,049431 |
| 2210408I21Rik | 72371 | 213,693467 | 5,629563946 | 1,876430266 | 0,002699 | 0,049431 |
| Rab3gap2 | 98732 | 73,7199352 | -5,599968113 | 1,865951013 | 0,00269 | 0,049431 |
| Tomm34 | 67145 | 28,0616447 | -4,413099648 | 1,470076275 | 0,002683 | 0,049431 |
| Ccdc97 | 52132 | 16,8129138 | -6,591310832 | 2,197031264 | 0,002699 | 0,049431 |
| Arfip2 | 76932 | 44,1500795 | -6,289112358 | 2,098167174 | 0,002723 | 0,049682 |
| Coq7 | 12850 | 30,9893856 | -6,643028843 | 2,21617368 | 0,002722 | 0,049682 |
| Kcna1 | 16485 | 290,893535 | 4,972090624 | 1,659901425 | 0,002741 | 0,049835 |
| Gm35251 | 102638766 | 18,2430307 | 7,584084289 | 2,53192685 | 0,002741 | 0,049835 |
| Trmt10c | 52575 | 69,6183971 | -6,488126293 | 2,166647707 | 0,002749 | 0,04988 |

**Table S7.** List of genes in the Protein Glycosylation (GO:0006486) GO term.

*Dpagt1*
*Galnt7*
*Galnt11*
*Alg6*
*Gfpt1*
*Pomt1*
*Alg3*
*Alg1*
*Dpm1*
*Mgat4b*
*St8sia5*
*Aqp11*
*Mgat2*

**Table S8.** Differentially expressed genes between Y-A$^{+}$ and O-A$^{+}$ astrocytes.

| Gene | Entrez_ID | baseMean | nonNorm_log2FoldChange | nonNorm_lfcSE | pvalue | padj |
|---|---|---|---|---|---|---|
| Tmem176a | 66058 | 3794,4761 | -30 | 2,553086529 | 7,02E-32 | 6,98E-28 |
| Acaa2 | 52538 | 26331,343 | -30 | 2,568291233 | 1,60E-31 | 7,93E-28 |
| Mvd | 192156 | 23,05504 | -29,99851336 | 2,586770683 | 4,27E-31 | 1,42E-27 |
| Gstm6 | 14867 | 8691,5828 | -30 | 2,683705528 | 5,19E-29 | 1,29E-25 |
| Gm17750 | 553095 | 8205,0495 | -30 | 2,733219914 | 4,98E-28 | 9,91E-25 |
| Adrm1 | 56436 | 198,93886 | -25,37831069 | 2,355948314 | 4,67E-27 | 7,74E-24 |
| Gorab | 98376 | 439,11441 | 25,58777829 | 2,417126435 | 3,46E-26 | 4,91E-23 |
| Pdia6 | 71853 | 530,74824 | 24,04072301 | 2,279927298 | 5,39E-26 | 6,69E-23 |
| Sesn3 | 75747 | 69,874756 | -23,36730159 | 2,241361529 | 1,90E-25 | 2,10E-22 |
| Lrig2 | 269473 | 446,3963 | 24,15649204 | 2,320619449 | 2,24E-25 | 2,23E-22 |
| Ddx21 | 56200 | 624,52051 | 24,58651357 | 2,466043882 | 2,06E-23 | 1,73E-20 |
| Trappc1 | 245828 | 366,60334 | 23,38643905 | 2,345989371 | 2,09E-23 | 1,73E-20 |
| Tap2 | 21355 | 14,383985 | -26,89459963 | 2,740565113 | 9,85E-23 | 7,53E-20 |
| Tcof1 | 21453 | 34,69428 | -22,86691899 | 2,352958515 | 2,52E-22 | 1,79E-19 |
| Gm29508 | 1,03E+08 | 17,028471 | -29,99913959 | 3,102880649 | 4,12E-22 | 2,73E-19 |
| Ppil6 | 73075 | 152,17176 | 21,67180596 | 2,25679277 | 7,77E-22 | 4,83E-19 |
| Drg1 | 13494 | 78,92185 | 24,85076141 | 2,618166076 | 2,27E-21 | 1,31E-18 |
| Rab5a | 271457 | 439,09287 | 23,31304665 | 2,457224272 | 2,37E-21 | 1,31E-18 |
| Gm27239 | 1,03E+08 | 81,929951 | -24,35195852 | 2,570143842 | 2,67E-21 | 1,40E-18 |
| Mlst8 | 56716 | 182,88164 | 24,4692724 | 2,584568281 | 2,87E-21 | 1,43E-18 |
| Coq6 | 217707 | 711,27833 | -26,41609523 | 2,820472331 | 7,54E-21 | 3,57E-18 |
| Tpcn1 | 252972 | 71,565772 | 21,03641824 | 2,254827665 | 1,06E-20 | 4,60E-18 |
| Sucla2 | 20916 | 251,6104 | 21,97646372 | 2,355037932 | 1,04E-20 | 4,60E-18 |
| Zc4h2 | 245522 | 14,186887 | -29,99706091 | 3,217113521 | 1,12E-20 | 4,63E-18 |
| Rpain | 69723 | 50,992628 | -23,5572921 | 2,528873214 | 1,22E-20 | 4,84E-18 |
| Nbdy | 70123 | 45,048926 | -23,51395999 | 2,527668806 | 1,37E-20 | 5,05E-18 |
| Ankrd13d | 68423 | 11,849924 | -29,99675526 | 3,224097896 | 1,35E-20 | 5,05E-18 |
| LOC118568128 | 1,19E+08 | 86,416132 | -24,62824373 | 2,659151214 | 2,01E-20 | 7,15E-18 |
| Rabac1 | 14470 | 192,3392 | 20,98081994 | 2,272056755 | 2,60E-20 | 8,92E-18 |

| | | | | | | |
|---|---|---|---|---|---|---|
| Mad2l2 | 71890 | 49,245847 | -23,08844056 | 2,516872923 | 4,58E-20 | 1,52E-17 |
| Tmem9 | 66241 | 66,087639 | 21,64609471 | 2,36363088 | 5,29E-20 | 1,64E-17 |
| Garem2 | 242915 | 402,52089 | 23,8320161 | 2,602230182 | 5,27E-20 | 1,64E-17 |
| Nat8f5 | 69049 | 110,97141 | -24,334845 | 2,669186399 | 7,73E-20 | 2,33E-17 |
| Abhd14a | 68644 | 48,741565 | 21,83636113 | 2,398482998 | 8,69E-20 | 2,54E-17 |
| Pkdcc | 106522 | 123,42492 | -25,01594069 | 2,768974262 | 1,65E-19 | 4,69E-17 |
| Hnmt | 140483 | 171,53025 | 22,13173793 | 2,451678079 | 1,76E-19 | 4,87E-17 |
| Elp2 | 58523 | 72,181555 | 21,47197759 | 2,400905915 | 3,78E-19 | 9,89E-17 |
| Bbs7 | 71492 | 274,59267 | 24,74679908 | 2,766335666 | 3,70E-19 | 9,89E-17 |
| Afap1l2 | 226250 | 255,67979 | 23,85415426 | 2,675628002 | 4,86E-19 | 1,24E-16 |
| Tsr1 | 104662 | 68,558634 | -24,24521938 | 2,728873894 | 6,41E-19 | 1,59E-16 |
| BC065397 | 436230 | 18,099926 | -23,9783066 | 2,707496769 | 8,27E-19 | 2,01E-16 |
| Ublcp1 | 79560 | 92,721179 | 20,27193586 | 2,291784319 | 9,11E-19 | 2,16E-16 |
| Gfpt1 | 14583 | 50,159263 | -22,26754415 | 2,533769289 | 1,52E-18 | 3,51E-16 |
| Nrep | 27528 | 84,043187 | 22,0937044 | 2,523877193 | 2,06E-18 | 4,66E-16 |
| Srp14 | 20813 | 247,31934 | 22,1561595 | 2,547108022 | 3,36E-18 | 7,43E-16 |
| Herpud1 | 64209 | 185,89097 | 20,7564325 | 2,397832302 | 4,87E-18 | 1,05E-15 |
| Dcp2 | 70640 | 57,429438 | 20,72030465 | 2,397970967 | 5,58E-18 | 1,18E-15 |
| Stx12 | 100226 | 150,29882 | 20,58933498 | 2,384015587 | 5,80E-18 | 1,20E-15 |
| Rbl2 | 19651 | 233,46645 | 21,63870598 | 2,514913436 | 7,69E-18 | 1,56E-15 |
| Ocrl | 320634 | 66,965918 | 22,00008963 | 2,563631053 | 9,36E-18 | 1,86E-15 |
| C920006O11Rik | 320295 | 12,68464 | -27,76597344 | 3,236640758 | 9,60E-18 | 1,87E-15 |
| 9330159M07Rik | 319673 | 44,517177 | -23,27199351 | 2,718508243 | 1,12E-17 | 2,15E-15 |
| Hcfc2 | 67933 | 110,99657 | 22,09068791 | 2,582788965 | 1,20E-17 | 2,25E-15 |
| Ccnq | 69109 | 35,383483 | -23,32449489 | 2,743421715 | 1,86E-17 | 3,43E-15 |
| Rbm4 | 19653 | 171,61431 | 22,98868214 | 2,719021875 | 2,80E-17 | 5,06E-15 |
| Mgat4b | 103534 | 24,665099 | -22,78606084 | 2,699467292 | 3,15E-17 | 5,59E-15 |
| Hmces | 232210 | 110,36601 | 24,1667239 | 2,876517805 | 4,41E-17 | 7,70E-15 |
| Ppdpf | 66496 | 69,175185 | 22,00235978 | 2,621218425 | 4,70E-17 | 8,06E-15 |
| Spice1 | 212514 | 40,848027 | 22,29329433 | 2,656697859 | 4,81E-17 | 8,10E-15 |
| Sec63 | 140740 | 73,032169 | 21,15533392 | 2,527110555 | 5,70E-17 | 9,44E-15 |
| Nudt2 | 66401 | 25,707274 | 23,68007611 | 2,831196419 | 6,06E-17 | 9,89E-15 |
| Gm9925 | 433202 | 61,023856 | 19,3402925 | 2,314968135 | 6,57E-17 | 1,05E-14 |

| | | | | | | |
|---|---|---|---|---|---|---|
| Glrx3 | 30926 | 78,489579 | 18,41355981 | 2,211613364 | 8,37E-17 | 1,32E-14 |
| Trappc6b | 78232 | 99,006516 | 21,1074754 | 2,536128626 | 8,60E-17 | 1,34E-14 |
| Thyn1 | 77862 | 57,954761 | -23,86587048 | 2,869384956 | 8,99E-17 | 1,35E-14 |
| Plscr2 | 18828 | 209,92209 | 23,18333944 | 2,787083217 | 8,94E-17 | 1,35E-14 |
| Lamtor5 | 68576 | 196,11562 | 21,84504195 | 2,63187182 | 1,04E-16 | 1,54E-14 |
| Ankrd29 | 225187 | 68,262022 | 19,76285627 | 2,383789386 | 1,13E-16 | 1,65E-14 |
| Minpp1 | 17330 | 69,406643 | 19,58345492 | 2,368165224 | 1,35E-16 | 1,94E-14 |
| Alkbh7 | 66400 | 39,805238 | -22,73899726 | 2,762044309 | 1,83E-16 | 2,60E-14 |
| Armcx4 | 1,01E+08 | 40,627337 | 20,11095228 | 2,445303429 | 1,96E-16 | 2,75E-14 |
| Slc37a4 | 14385 | 18,961165 | -22,05357695 | 2,685888939 | 2,20E-16 | 3,03E-14 |
| Pacrg | 69310 | 178,76997 | 20,8938832 | 2,54881488 | 2,45E-16 | 3,30E-14 |
| Chil1 | 12654 | 201,14422 | 21,588475 | 2,63331528 | 2,44E-16 | 3,30E-14 |
| Pcdh19 | 279653 | 162,25594 | 22,73077408 | 2,773809239 | 2,51E-16 | 3,33E-14 |
| Fra10ac1 | 70567 | 46,742904 | 19,50593005 | 2,384719616 | 2,85E-16 | 3,73E-14 |
| Gm15764 | 1,03E+08 | 72,08412 | 18,98135718 | 2,329962103 | 3,74E-16 | 4,83E-14 |
| 0610012G03Rik | 106264 | 73,777268 | 19,41720218 | 2,38690423 | 4,12E-16 | 5,19E-14 |
| Cbfb | 12400 | 95,106696 | 20,12321094 | 2,473624574 | 4,12E-16 | 5,19E-14 |
| Bex3 | 12070 | 49,485065 | 19,42060402 | 2,395244723 | 5,15E-16 | 6,40E-14 |
| Echdc2 | 52430 | 35,264945 | 22,87293108 | 2,830426577 | 6,42E-16 | 7,88E-14 |
| Slc9a9 | 331004 | 101,69854 | 20,45502827 | 2,53419172 | 6,94E-16 | 8,41E-14 |
| Smdt1 | 69029 | 92,996036 | 19,99852835 | 2,483513613 | 8,11E-16 | 9,72E-14 |
| Gm527 | 217648 | 18,300558 | -22,69093082 | 2,826222299 | 9,85E-16 | 1,17E-13 |
| Pold3 | 67967 | 38,549443 | 19,23008937 | 2,396122697 | 1,01E-15 | 1,18E-13 |
| Ilk | 16202 | 50,441954 | 19,05530559 | 2,378287837 | 1,13E-15 | 1,30E-13 |
| Prpf4 | 70052 | 53,366917 | 22,05289533 | 2,754354034 | 1,18E-15 | 1,35E-13 |
| Gal3st4 | 330217 | 17,147917 | -25,0133477 | 3,128341313 | 1,29E-15 | 1,46E-13 |
| Tyro3 | 22174 | 51,117715 | 18,29678576 | 2,299091191 | 1,74E-15 | 1,95E-13 |
| Ugdh | 22235 | 149,41686 | 22,1055299 | 2,778432029 | 1,78E-15 | 1,96E-13 |
| Psen2 | 19165 | 52,057936 | 21,76456903 | 2,736452482 | 1,81E-15 | 1,98E-13 |
| Pycr2 | 69051 | 20,656937 | -22,28315917 | 2,803386081 | 1,89E-15 | 2,04E-13 |
| Ppp1r21 | 73825 | 59,906882 | 19,86526489 | 2,500335021 | 1,94E-15 | 2,08E-13 |
| Tmem214 | 68796 | 70,398285 | 20,41703737 | 2,571495582 | 2,03E-15 | 2,10E-13 |
| Cwf19l1 | 72502 | 95,698652 | 20,52244322 | 2,584616603 | 2,02E-15 | 2,10E-13 |

| | | | | | | |
|---|---|---|---|---|---|---|
| Fam174c | 69770 | 20,109861 | 22,02884706 | 2,773972429 | 2,00E-15 | 2,10E-13 |
| Cenpw | 66311 | 10,566935 | -23,32872536 | 2,940455553 | 2,13E-15 | 2,18E-13 |
| Stx3 | 20908 | 72,381204 | 21,77121761 | 2,751704178 | 2,53E-15 | 2,57E-13 |
| Vps51 | 68505 | 31,086603 | 19,39618596 | 2,45612086 | 2,86E-15 | 2,87E-13 |
| Aldh4a1 | 212647 | 58,941394 | 20,01847697 | 2,537115068 | 3,02E-15 | 2,97E-13 |
| Nat1 | 17960 | 120,57075 | 22,29311922 | 2,825071082 | 2,99E-15 | 2,97E-13 |
| Cars | 27267 | 27,448862 | 20,80699691 | 2,641354635 | 3,34E-15 | 3,26E-13 |
| Pctp | 18559 | 12,566715 | -21,36647266 | 2,714457566 | 3,51E-15 | 3,39E-13 |
| Ppil2 | 66053 | 54,83584 | 19,05992482 | 2,423023894 | 3,66E-15 | 3,50E-13 |
| Smim1 | 68859 | 145,05727 | 20,71981134 | 2,641316122 | 4,35E-15 | 4,12E-13 |
| Ahcy | 269378 | 24,720585 | 22,29038765 | 2,843959097 | 4,59E-15 | 4,30E-13 |
| Nfam1 | 74039 | 31,054417 | 19,98435365 | 2,551730159 | 4,81E-15 | 4,47E-13 |
| Gpx1 | 14775 | 81,797941 | 20,49760958 | 2,619308536 | 5,05E-15 | 4,65E-13 |
| Dmac1 | 66928 | 75,364161 | 19,59234156 | 2,504538371 | 5,17E-15 | 4,67E-13 |
| Agps | 228061 | 147,86741 | 22,25308228 | 2,84454355 | 5,15E-15 | 4,67E-13 |
| Gm39080 | 1,05E+08 | 85,001823 | 21,81582241 | 2,791137893 | 5,45E-15 | 4,88E-13 |
| Cpt1c | 78070 | 72,359627 | 19,85844545 | 2,541827192 | 5,60E-15 | 4,97E-13 |
| Dennd6b | 69440 | 47,636525 | 19,06894445 | 2,443089473 | 5,94E-15 | 5,23E-13 |
| Atg4b | 66615 | 71,085557 | 20,75294954 | 2,668416534 | 7,41E-15 | 6,47E-13 |
| Armc10 | 67211 | 35,949 | 18,74120664 | 2,415908335 | 8,67E-15 | 7,50E-13 |
| Ap2a1 | 11771 | 27,261097 | 18,81673728 | 2,428467853 | 9,31E-15 | 7,98E-13 |
| Adprm | 66358 | 27,429028 | 19,84238752 | 2,573312074 | 1,25E-14 | 1,06E-12 |
| Gm266 | 212539 | 51,516883 | 18,92042194 | 2,457981211 | 1,39E-14 | 1,17E-12 |
| Ntan1 | 18203 | 85,680376 | 19,2009301 | 2,496628687 | 1,46E-14 | 1,22E-12 |
| Pak5 | 241656 | 60,549114 | 21,32486276 | 2,776347197 | 1,58E-14 | 1,31E-12 |
| 1700029J07Rik | 69479 | 47,684821 | 18,3343074 | 2,394736211 | 1,92E-14 | 1,58E-12 |
| Cep20 | 66086 | 86,460273 | 19,4276191 | 2,53841668 | 1,96E-14 | 1,60E-12 |
| Vps52 | 224705 | 81,789325 | 21,1753446 | 2,768945035 | 2,05E-14 | 1,66E-12 |
| Tsr2 | 69499 | 28,48724 | 21,76404986 | 2,847379174 | 2,11E-14 | 1,69E-12 |
| Polr3e | 26939 | 21,182691 | 21,30709817 | 2,792265171 | 2,33E-14 | 1,86E-12 |
| Exosc10 | 50912 | 28,175347 | 19,52203515 | 2,568677446 | 2,96E-14 | 2,34E-12 |
| Rit1 | 19769 | 52,621613 | 18,87580918 | 2,483994691 | 2,98E-14 | 2,34E-12 |
| Lyar | 17089 | 24,16333 | 19,8876197 | 2,621089889 | 3,26E-14 | 2,53E-12 |

| | | | | | | |
|---|---|---|---|---|---|---|
| B230217C12Rik | 68127 | 76,95947 | -21,6793306 | 2,860328255 | 3,47E-14 | 2,68E-12 |
| Vps39 | 269338 | 188,2988 | 21,37101653 | 2,821409037 | 3,60E-14 | 2,76E-12 |
| Cep43 | 75296 | 25,936531 | 18,80009034 | 2,485834698 | 3,94E-14 | 2,99E-12 |
| Cul4b | 72584 | 76,525605 | 19,71692131 | 2,611730716 | 4,37E-14 | 3,30E-12 |
| Mrpl13 | 68537 | 34,370994 | 20,00622974 | 2,652669007 | 4,63E-14 | 3,46E-12 |
| H4c8 | 69386 | 55,303241 | 19,12547048 | 2,548132807 | 6,11E-14 | 4,54E-12 |
| Oaz2 | 18247 | 23,617836 | 19,78358607 | 2,647307742 | 7,83E-14 | 5,77E-12 |
| Ufd1 | 22230 | 48,398239 | 19,75056684 | 2,648246178 | 8,79E-14 | 6,42E-12 |
| Pitrm1 | 69617 | 121,32649 | 21,36075159 | 2,867421104 | 9,37E-14 | 6,75E-12 |
| Zfp438 | 240186 | 68,574748 | 22,11153257 | 2,967974766 | 9,33E-14 | 6,75E-12 |
| Tm7sf2 | 73166 | 33,857234 | 19,61575156 | 2,634344235 | 9,61E-14 | 6,84E-12 |
| Hey2 | 15214 | 57,138309 | 22,24622513 | 2,987733829 | 9,63E-14 | 6,84E-12 |
| Ppa2 | 74776 | 46,955252 | 20,57474653 | 2,765521635 | 1,01E-13 | 7,12E-12 |
| Mipep | 70478 | 83,596539 | 20,51734649 | 2,763171002 | 1,13E-13 | 7,88E-12 |
| Cyb5r3 | 109754 | 31,53867 | 20,0680998 | 2,713221678 | 1,40E-13 | 9,73E-12 |
| Inka2 | 109050 | 24,952342 | 18,82597666 | 2,545660268 | 1,41E-13 | 9,74E-12 |
| Als2 | 74018 | 69,065889 | 20,24874643 | 2,74125449 | 1,51E-13 | 1,03E-11 |
| Ccdc107 | 622404 | 107,41345 | 19,79058484 | 2,687655722 | 1,79E-13 | 1,22E-11 |
| Pagr1a | 67278 | 37,798103 | 18,82391234 | 2,556921933 | 1,81E-13 | 1,23E-11 |
| Rab22a | 19334 | 25,087846 | 20,89648865 | 2,850437952 | 2,28E-13 | 1,54E-11 |
| Trmt61b | 68789 | 20,077705 | 22,45471899 | 3,064996653 | 2,37E-13 | 1,58E-11 |
| Fxyd6 | 59095 | 31,315546 | 19,01238589 | 2,596398592 | 2,43E-13 | 1,61E-11 |
| 2010320M18Rik | 72093 | 94,623784 | 20,28510436 | 2,771029018 | 2,47E-13 | 1,63E-11 |
| Ciao2a | 68250 | 53,184343 | 20,55012382 | 2,808558845 | 2,54E-13 | 1,66E-11 |
| Pop4 | 66161 | 96,688745 | 19,75309552 | 2,699919418 | 2,55E-13 | 1,66E-11 |
| 1700030K09Rik | 72254 | 17,906323 | 19,48841772 | 2,666175196 | 2,68E-13 | 1,73E-11 |
| Lrrc28 | 67867 | 23,253042 | 20,71130219 | 2,835012866 | 2,76E-13 | 1,77E-11 |
| Cryzl1 | 66609 | 53,756821 | 18,1540357 | 2,486325515 | 2,84E-13 | 1,79E-11 |
| Fam122a | 68034 | 24,659172 | 19,02023066 | 2,605332541 | 2,87E-13 | 1,79E-11 |
| Tmem86a | 67893 | 26,007828 | 20,76463807 | 2,843722554 | 2,84E-13 | 1,79E-11 |
| Dhx32 | 101437 | 19,854602 | 21,59213565 | 2,957587817 | 2,87E-13 | 1,79E-11 |
| Ube2f | 67921 | 58,609878 | 18,93879006 | 2,599928725 | 3,23E-13 | 2,01E-11 |
| Gm4767 | 210583 | 20,301181 | 22,45487431 | 3,083779887 | 3,30E-13 | 2,04E-11 |

| | | | | | | |
|---|---|---|---|---|---|---|
| Efemp2 | 58859 | 25,159839 | 19,05702058 | 2,62055794 | 3,54E-13 | 2,17E-11 |
| Kbtbd7 | 211255 | 25,581114 | 19,35032809 | 2,664797269 | 3,83E-13 | 2,34E-11 |
| Plat | 18791 | 28,74326 | 18,88656465 | 2,609747107 | 4,59E-13 | 2,78E-11 |
| Nup37 | 69736 | 37,244944 | 19,21491897 | 2,663305154 | 5,41E-13 | 3,26E-11 |
| Ccdc124 | 234388 | 51,128015 | 18,82004097 | 2,619528871 | 6,74E-13 | 4,04E-11 |
| Hars2 | 70791 | 18,741433 | 18,5067268 | 2,579735446 | 7,29E-13 | 4,32E-11 |
| Slc35b3 | 108652 | 36,683403 | 19,46018301 | 2,712504002 | 7,27E-13 | 4,32E-11 |
| Rnf227 | 80515 | 15,793624 | 18,73427512 | 2,613527696 | 7,60E-13 | 4,47E-11 |
| Zdhhc13 | 243983 | 31,855922 | 20,87164272 | 2,917643634 | 8,45E-13 | 4,95E-11 |
| Ass1 | 11898 | 80,296783 | 19,6856584 | 2,757725671 | 9,44E-13 | 5,49E-11 |
| LOC108169181 | 1,08E+08 | 126,64971 | 20,10853549 | 2,817826977 | 9,60E-13 | 5,55E-11 |
| Gm35251 | 1,03E+08 | 18,243031 | 20,45235568 | 2,870919393 | 1,05E-12 | 6,03E-11 |
| Prmt7 | 214572 | 19,692426 | 19,84718813 | 2,788859027 | 1,11E-12 | 6,32E-11 |
| Ebp | 13595 | 14,264198 | 18,29815566 | 2,573997012 | 1,17E-12 | 6,65E-11 |
| Ap5s1 | 69596 | 26,040604 | 18,01260421 | 2,534261668 | 1,18E-12 | 6,67E-11 |
| Man1c1 | 230815 | 35,002266 | 19,48730339 | 2,74222171 | 1,19E-12 | 6,69E-11 |
| Tmem62 | 96957 | 51,292029 | 20,65348543 | 2,928526909 | 1,76E-12 | 9,82E-11 |
| Lipe | 16890 | 18,471068 | 20,34603715 | 2,889078736 | 1,89E-12 | 1,05E-10 |
| Pabpn1 | 54196 | 32,68927 | -29,99918366 | 4,278112129 | 2,35E-12 | 1,30E-10 |
| Gm36298 | 1,03E+08 | 19,545508 | -29,99333844 | 4,278050316 | 2,37E-12 | 1,30E-10 |
| Snap91 | 20616 | 42,184706 | 19,14804389 | 2,735040304 | 2,54E-12 | 1,39E-10 |
| Qars | 97541 | 107,36026 | 18,0393588 | 2,577603573 | 2,59E-12 | 1,41E-10 |
| Nol6 | 230082 | 34,500635 | 18,7712008 | 2,685769893 | 2,77E-12 | 1,49E-10 |
| Zfpl1 | 81909 | 30,350786 | 18,12141553 | 2,599474616 | 3,14E-12 | 1,69E-10 |
| Glrx5 | 73046 | 2234,8206 | -29,02836228 | 4,164745577 | 3,17E-12 | 1,69E-10 |
| Gemin7 | 69731 | 13,308441 | 18,27854399 | 2,62426478 | 3,28E-12 | 1,74E-10 |
| Aprt | 11821 | 63,648288 | 20,67525008 | 2,971954605 | 3,48E-12 | 1,84E-10 |
| Ubl7 | 69459 | 37,009895 | 18,48716113 | 2,658523666 | 3,55E-12 | 1,87E-10 |
| Mak16 | 67920 | 33,822036 | 19,65780455 | 2,834937194 | 4,09E-12 | 2,14E-10 |
| Rhbdf1 | 13650 | 54,491957 | 18,97151798 | 2,736731921 | 4,14E-12 | 2,15E-10 |
| Eftud2 | 20624 | 62,415137 | 19,21892036 | 2,772192589 | 4,13E-12 | 2,15E-10 |
| Patz1 | 56218 | 48,374971 | 17,98371855 | 2,606408766 | 5,21E-12 | 2,68E-10 |
| Larp7 | 28036 | 108,09792 | 20,37516726 | 2,953674525 | 5,26E-12 | 2,70E-10 |

| | | | | | | |
|---|---|---|---|---|---|---|
| Prpf3 | 70767 | 18,695077 | 20,19752221 | 2,928300055 | 5,30E-12 | 2,70E-10 |
| Gins4 | 109145 | 37,670464 | 17,93904664 | 2,605368786 | 5,76E-12 | 2,92E-10 |
| Scyl3 | 240880 | 48,938427 | 20,12084772 | 2,926499595 | 6,18E-12 | 3,12E-10 |
| Timm9 | 30056 | 15,139559 | -19,79223396 | 2,880225079 | 6,34E-12 | 3,17E-10 |
| Setd6 | 66083 | 20,41908 | 20,3946076 | 2,96778092 | 6,33E-12 | 3,17E-10 |
| Adi1 | 104923 | 67,299677 | 17,24083806 | 2,512809837 | 6,83E-12 | 3,40E-10 |
| Manbal | 69161 | 29,405754 | 19,29403299 | 2,817326055 | 7,47E-12 | 3,70E-10 |
| Hprt | 15452 | 55,456158 | 20,09177077 | 2,943109043 | 8,69E-12 | 4,28E-10 |
| Psmd5 | 66998 | 34,65648 | 19,16329378 | 2,821825341 | 1,11E-11 | 5,44E-10 |
| Tmem126b | 68472 | 31,062013 | 19,14390943 | 2,819177717 | 1,12E-11 | 5,44E-10 |
| Plpp5 | 71910 | 17,175169 | 20,87801699 | 3,080006845 | 1,21E-11 | 5,89E-10 |
| 5530601H04Rik | 71445 | 13,008602 | 19,5308276 | 2,883812555 | 1,27E-11 | 6,11E-10 |
| Trpc1 | 22063 | 21,764798 | 17,28435858 | 2,554492038 | 1,32E-11 | 6,35E-10 |
| Cib2 | 56506 | 33,272116 | 18,99405068 | 2,808181879 | 1,34E-11 | 6,43E-10 |
| 2510009E07Rik | 72190 | 22562,322 | -14,18133269 | 2,100359022 | 1,46E-11 | 6,95E-10 |
| Rangrf | 57785 | 20,831589 | 19,36304728 | 2,875669448 | 1,66E-11 | 7,85E-10 |
| Enoph1 | 67870 | 23,072011 | 18,59578928 | 2,768745705 | 1,86E-11 | 8,78E-10 |
| Sap30l | 50724 | 8150,1771 | -17,6034488 | 2,621652039 | 1,89E-11 | 8,84E-10 |
| St8sia5 | 225742 | 24,288955 | -28,70930763 | 4,278337887 | 1,94E-11 | 9,06E-10 |
| Slc7a6os | 66432 | 10,735588 | 18,94857568 | 2,825283588 | 1,99E-11 | 9,25E-10 |
| Ccdc58 | 381045 | 58,614898 | 18,93387382 | 2,8302268 | 2,23E-11 | 1,03E-09 |
| Fam234a | 106581 | 41,661062 | 19,11649574 | 2,874021918 | 2,90E-11 | 1,34E-09 |
| Rpgrip1l | 244585 | 16,790507 | 20,43692777 | 3,078192729 | 3,15E-11 | 1,44E-09 |
| Kin | 16588 | 33,238762 | 18,97133457 | 2,882115036 | 4,63E-11 | 2,11E-09 |
| Cep78 | 208518 | 37,944175 | 18,28463242 | 2,778873493 | 4,71E-11 | 2,13E-09 |
| Igbp1 | 18518 | 31,909823 | 19,39517643 | 2,947381593 | 4,69E-11 | 2,13E-09 |
| Snhg4 | 1,01E+08 | 43,305246 | 18,38154747 | 2,801985409 | 5,37E-11 | 2,42E-09 |
| Amn1 | 232566 | 56,642795 | 17,30956405 | 2,647908167 | 6,27E-11 | 2,81E-09 |
| Nme6 | 54369 | 14,61106 | 22,44482361 | 3,435939879 | 6,47E-11 | 2,89E-09 |
| Inpp5k | 19062 | 38,485126 | 19,13275905 | 2,930954008 | 6,67E-11 | 2,96E-09 |
| Yars2 | 70120 | 19,274041 | 18,2635985 | 2,808501011 | 7,87E-11 | 3,48E-09 |
| Prps1 | 19139 | 32,31528 | 18,915306 | 2,909049527 | 7,91E-11 | 3,48E-09 |
| Zfp128 | 243833 | 11,441992 | -21,73047008 | 3,346211533 | 8,36E-11 | 3,66E-09 |

| | | | | | | |
|---|---|---|---|---|---|---|
| Ext1 | 14042 | 25,203081 | 17,05006622 | 2,625862365 | 8,41E-11 | 3,67E-09 |
| U2af1l4 | 233073 | 16,697619 | 17,41676886 | 2,682730195 | 8,46E-11 | 3,67E-09 |
| Slc30a5 | 69048 | 24,228361 | 17,94145966 | 2,780174682 | 1,09E-10 | 4,72E-09 |
| Gm38592 | 1,03E+08 | 58,693992 | 27,86427836 | 4,317983438 | 1,10E-10 | 4,72E-09 |
| Elovl2 | 54326 | 41160,054 | -12,22606041 | 1,896006064 | 1,13E-10 | 4,85E-09 |
| Cstf2 | 108062 | 70,305047 | 18,56295921 | 2,881652788 | 1,18E-10 | 5,04E-09 |
| Rcbtb2 | 105670 | 46,057274 | 17,47960035 | 2,726723605 | 1,45E-10 | 6,17E-09 |
| Sap25 | 751865 | 30,614106 | 19,1120392 | 2,994168499 | 1,74E-10 | 7,31E-09 |
| A230072C01Rik | 320742 | 15,032745 | 18,64728101 | 2,921068232 | 1,73E-10 | 7,31E-09 |
| Cfap300 | 234912 | 171,50214 | 27,50799471 | 4,317848554 | 1,88E-10 | 7,89E-09 |
| Gmppa | 69080 | 17,902616 | 17,56385948 | 2,758969888 | 1,94E-10 | 8,10E-09 |
| Rmdn3 | 67809 | 57,604022 | 18,69202235 | 2,93727449 | 1,97E-10 | 8,19E-09 |
| Rpp30 | 54364 | 36,165882 | 19,20993205 | 3,025203305 | 2,15E-10 | 8,92E-09 |
| H1f10 | 243529 | 16,05332 | 19,37189544 | 3,053672774 | 2,24E-10 | 9,23E-09 |
| Fam76a | 230789 | 26,495259 | 17,08574788 | 2,693416829 | 2,25E-10 | 9,23E-09 |
| Zfp934 | 77117 | 27,35263 | 19,35683123 | 3,051954652 | 2,26E-10 | 9,26E-09 |
| Psmg4 | 69666 | 22,726158 | 16,66788494 | 2,635777548 | 2,55E-10 | 1,04E-08 |
| Ccdc22 | 54638 | 21,915756 | 18,81201101 | 2,976381442 | 2,61E-10 | 1,06E-08 |
| Pter | 19212 | 13,667283 | 17,1115288 | 2,710438074 | 2,73E-10 | 1,10E-08 |
| Dym | 69190 | 15,821846 | 19,28873467 | 3,05649426 | 2,78E-10 | 1,12E-08 |
| Polr1e | 64424 | 14,839723 | 20,26661502 | 3,230243085 | 3,52E-10 | 1,41E-08 |
| Ncoa5 | 228869 | 17,912786 | 17,67703146 | 2,820002267 | 3,65E-10 | 1,46E-08 |
| Gm26788 | 1,08E+08 | 87,80307 | -26,69732024 | 4,277632326 | 4,34E-10 | 1,73E-08 |
| Glb1 | 12091 | 19,412324 | 17,37374648 | 2,784831949 | 4,41E-10 | 1,75E-08 |
| Mccc2 | 78038 | 21,204648 | 17,20622778 | 2,762555464 | 4,71E-10 | 1,86E-08 |
| Nxt2 | 237082 | 32970,852 | -15,57814518 | 2,50266566 | 4,83E-10 | 1,90E-08 |
| 2610044O15Rik8 | 72139 | 14,567329 | 20,20445151 | 3,247502981 | 4,92E-10 | 1,93E-08 |
| Phtf1 | 18685 | 174,78957 | 25,07894338 | 4,031416505 | 4,94E-10 | 1,93E-08 |
| Rps4l | 66184 | 14,597149 | 18,10174991 | 2,918025374 | 5,52E-10 | 2,15E-08 |
| Gsdmc2 | 331063 | 23,457806 | 18,08485691 | 2,9278419 | 6,54E-10 | 2,53E-08 |
| Tbp | 21374 | 62,378671 | 18,0763694 | 2,949972837 | 8,92E-10 | 3,44E-08 |
| Nupr1 | 56312 | 21,87295 | 18,39583664 | 3,013306328 | 1,03E-09 | 3,95E-08 |
| Dffa | 13347 | 18,733536 | 17,79003461 | 2,920193841 | 1,11E-09 | 4,26E-08 |

| | | | | | | |
|---|---|---|---|---|---|---|
| Fam50a | 108160 | 17,972017 | 17,33817188 | 2,866938022 | 1,47E-09 | 5,60E-08 |
| Slc25a1 | 13358 | 17,5766 | 17,57577212 | 2,9189762 | 1,73E-09 | 6,57E-08 |
| Rtl8a | 66158 | 11,810192 | 17,48110118 | 2,924737456 | 2,27E-09 | 8,60E-08 |
| Plekhj1 | 78670 | 13,90179 | 17,28900235 | 2,894619816 | 2,33E-09 | 8,79E-08 |
| Sec31a | 69162 | 15303,129 | -15,25234953 | 2,554566217 | 2,36E-09 | 8,87E-08 |
| Uvrag | 78610 | 11,763882 | 18,15143201 | 3,047611631 | 2,59E-09 | 9,67E-08 |
| Eif2b4 | 13667 | 11,795766 | 17,10371321 | 2,913166856 | 4,33E-09 | 1,61E-07 |
| Cdan1 | 68968 | 1742,1972 | -14,27766805 | 2,44504843 | 5,24E-09 | 1,94E-07 |
| 2810403D21Rik | 69964 | 16,077811 | 18,96324827 | 3,266208604 | 6,40E-09 | 2,37E-07 |
| Gm52051 | 1,15E+08 | 16,298584 | 17,94113247 | 3,100487691 | 7,18E-09 | 2,65E-07 |
| Lage3 | 66192 | 12029,934 | -13,34651156 | 2,308216776 | 7,37E-09 | 2,71E-07 |
| Ifitm2 | 80876 | 39,986414 | 24,92917043 | 4,318340781 | 7,79E-09 | 2,85E-07 |
| Nif3l1 | 65102 | 42,139681 | 24,10129243 | 4,188243853 | 8,69E-09 | 3,17E-07 |
| Il1r1 | 16177 | 209,32162 | 23,45075582 | 4,08856815 | 9,71E-09 | 3,52E-07 |
| Phf5a | 68479 | 21,104411 | 24,72853089 | 4,319192293 | 1,03E-08 | 3,73E-07 |
| Ltv1 | 353258 | 16,023423 | 19,96238677 | 3,505409957 | 1,24E-08 | 4,45E-07 |
| Gm5617 | 434402 | 22,164032 | 24,56009859 | 4,318480927 | 1,29E-08 | 4,64E-07 |
| Nr3c2 | 110784 | 6008,2944 | -15,64346119 | 2,753937045 | 1,34E-08 | 4,81E-07 |
| Lcmt1 | 30949 | 18,562507 | 16,13135232 | 2,858654355 | 1,67E-08 | 5,96E-07 |
| Zfp524 | 66056 | 13,365868 | 17,93079536 | 3,21245995 | 2,38E-08 | 8,46E-07 |
| Tmub1 | 64295 | 23,860732 | 14,38992373 | 2,579498232 | 2,42E-08 | 8,58E-07 |
| Prpf31 | 68988 | 12,030027 | 17,74633059 | 3,18157086 | 2,44E-08 | 8,59E-07 |
| Ddb2 | 107986 | 9954,9785 | -12,54192214 | 2,262229059 | 2,96E-08 | 1,04E-06 |
| Pwp2 | 110816 | 13,936665 | 18,26084391 | 3,311728434 | 3,51E-08 | 1,23E-06 |
| Cetn2 | 26370 | 712,27652 | 12,41553116 | 2,253952338 | 3,62E-08 | 1,26E-06 |
| Nab2 | 17937 | 4420,8638 | -12,51738844 | 2,273989846 | 3,70E-08 | 1,29E-06 |
| Hps5 | 246694 | 7444,5152 | -15,83817846 | 2,87911279 | 3,78E-08 | 1,31E-06 |
| Xrn1 | 24127 | 53331,96 | -11,40311238 | 2,079608095 | 4,17E-08 | 1,44E-06 |
| Ermp1 | 226090 | 11598,029 | -12,29278545 | 2,258537112 | 5,24E-08 | 1,80E-06 |
| Pnrc1 | 108767 | 264,77197 | 8,412214361 | 1,551780645 | 5,93E-08 | 2,03E-06 |
| Gtpbp8 | 66067 | 17,267522 | 18,6711288 | 3,448704069 | 6,16E-08 | 2,11E-06 |
| Lcat | 16816 | 994,1806 | -8,638533201 | 1,597699284 | 6,41E-08 | 2,18E-06 |
| Mgat1 | 17308 | 17221,294 | -12,42158978 | 2,314907406 | 8,05E-08 | 2,73E-06 |

| | | | | | | |
|---|---|---|---|---|---|---|
| Fam183b | 75429 | 46,15117 | 23,16864045 | 4,318309702 | 8,09E-08 | 2,74E-06 |
| Mospd2 | 76763 | 29,520181 | 14,79973984 | 2,767079303 | 8,87E-08 | 2,99E-06 |
| Pcyt2 | 68671 | 291,40374 | 9,543626907 | 1,790538226 | 9,82E-08 | 3,30E-06 |
| Tmsb10 | 19240 | 175,7209 | 12,34791661 | 2,317678666 | 9,95E-08 | 3,32E-06 |
| Smpd1 | 20597 | 336,27026 | -9,837545471 | 1,846407074 | 9,93E-08 | 3,32E-06 |
| Arl8b | 67166 | 13153,838 | -11,55068351 | 2,177304548 | 1,13E-07 | 3,75E-06 |
| Lmcd1 | 30937 | 4168,4154 | -12,04710246 | 2,277096539 | 1,22E-07 | 4,04E-06 |
| Pomt1 | 99011 | 48,490952 | 15,65044999 | 2,96162914 | 1,26E-07 | 4,17E-06 |
| Nek3 | 23954 | 14,381027 | 17,20406919 | 3,256783789 | 1,27E-07 | 4,20E-06 |
| Yaf2 | 67057 | 13580,357 | -11,7785056 | 2,231006588 | 1,30E-07 | 4,25E-06 |
| 1700088E04Rik | 27660 | 28,191202 | 22,3995246 | 4,261454476 | 1,47E-07 | 4,81E-06 |
| Mettl23 | 74319 | 14136,88 | -11,10118902 | 2,113585058 | 1,50E-07 | 4,90E-06 |
| Smpdl3a | 57319 | 5819,1137 | -8,996144276 | 1,717046165 | 1,61E-07 | 5,24E-06 |
| Gm32912 | 1,03E+08 | 1816,9952 | -12,96772688 | 2,48833836 | 1,87E-07 | 6,07E-06 |
| Decr1 | 67460 | 150,84582 | -9,510023982 | 1,828334899 | 1,98E-07 | 6,38E-06 |
| Ccp110 | 101565 | 202,71731 | 8,649221984 | 1,66416724 | 2,02E-07 | 6,51E-06 |
| Ik | 24010 | 286,54187 | 9,667426149 | 1,862983504 | 2,11E-07 | 6,77E-06 |
| Atp13a5 | 268878 | 283,13328 | 12,16515281 | 2,364643573 | 2,68E-07 | 8,57E-06 |
| 8030462N17Rik | 212163 | 2778,7759 | -8,846044481 | 1,721876931 | 2,79E-07 | 8,88E-06 |
| Agrn | 11603 | 327,60489 | 7,134997394 | 1,39108805 | 2,91E-07 | 9,25E-06 |
| Timm50 | 66525 | 17,678797 | 22,14424745 | 4,318387737 | 2,93E-07 | 9,28E-06 |
| Tbk1 | 56480 | 166,08485 | 10,65380967 | 2,083173679 | 3,15E-07 | 9,95E-06 |
| Klhdc7b | 546648 | 128,57396 | -8,223283605 | 1,610158874 | 3,27E-07 | 1,03E-05 |
| Eprs | 107508 | 438,89906 | 7,398447559 | 1,449253075 | 3,31E-07 | 1,04E-05 |
| Ccdc117 | 104479 | 17,317079 | 15,13388095 | 2,96657567 | 3,37E-07 | 1,05E-05 |
| Ndufa10 | 67273 | 23065,1 | -11,73030378 | 2,300771588 | 3,42E-07 | 1,07E-05 |
| Zdhhc17 | 320150 | 121,34954 | 10,7472002 | 2,114719843 | 3,73E-07 | 1,16E-05 |
| Ppp1r9a | 243725 | 263,47266 | 7,143250098 | 1,408490275 | 3,95E-07 | 1,22E-05 |
| Mrpl2 | 27398 | 38,98947 | 14,03048047 | 2,779179277 | 4,45E-07 | 1,38E-05 |
| Polh | 80905 | 21,811898 | 21,76781632 | 4,318030332 | 4,63E-07 | 1,42E-05 |
| Rpp14 | 67053 | 348,75092 | -12,53376401 | 2,48755487 | 4,69E-07 | 1,44E-05 |
| Tekt1 | 21689 | 191,65377 | 21,95671714 | 4,363578848 | 4,86E-07 | 1,49E-05 |
| Qpctl | 67369 | 70,272119 | 14,95903859 | 2,975243506 | 4,96E-07 | 1,51E-05 |

| | | | | | | |
|---|---|---|---|---|---|---|
| Ephx2 | 13850 | 31235,738 | -12,1400398 | 2,431590283 | 5,96E-07 | 1,81E-05 |
| Sgip1 | 73094 | 467,79954 | 8,973125126 | 1,808266145 | 6,97E-07 | 2,11E-05 |
| Glod4 | 67201 | 468,73674 | 10,83372154 | 2,186723883 | 7,26E-07 | 2,19E-05 |
| Map1b | 17755 | 378,56467 | 9,017129294 | 1,820679697 | 7,32E-07 | 2,21E-05 |
| Pcdhga6 | 93714 | 26,163484 | 14,85653423 | 3,003541017 | 7,56E-07 | 2,27E-05 |
| Morn5 | 75495 | 78,723104 | 21,4362332 | 4,337795554 | 7,74E-07 | 2,32E-05 |
| Meig1 | 104362 | 82,43991 | 21,23074672 | 4,318033733 | 8,80E-07 | 2,63E-05 |
| Endod1 | 71946 | 188,96269 | 9,768909199 | 1,990196868 | 9,18E-07 | 2,73E-05 |
| Tomm7 | 66169 | 167,54849 | 9,223585792 | 1,880343996 | 9,33E-07 | 2,77E-05 |
| Aass | 30956 | 19463,629 | -12,05208546 | 2,457756201 | 9,41E-07 | 2,78E-05 |
| Fbxo22 | 71999 | 527,76878 | -11,49648747 | 2,346745497 | 9,64E-07 | 2,84E-05 |
| Cbwd1 | 226043 | 197,5064 | 21,31633638 | 4,354885581 | 9,84E-07 | 2,90E-05 |
| Bpnt2 | 242291 | 384,46983 | 8,332132075 | 1,704151804 | 1,01E-06 | 2,97E-05 |
| Cpsf2 | 51786 | 417,42391 | -11,77937489 | 2,409687422 | 1,02E-06 | 2,97E-05 |
| Ip6k1 | 27399 | 205,22604 | -9,304152744 | 1,904863743 | 1,04E-06 | 3,03E-05 |
| Rbm27 | 225432 | 8590,924 | -9,486661169 | 1,950460101 | 1,15E-06 | 3,34E-05 |
| Rab1a | 19324 | 8765,4219 | -9,315151157 | 1,915130905 | 1,15E-06 | 3,34E-05 |
| Pls3 | 102866 | 33626,052 | -12,5762048 | 2,586829943 | 1,16E-06 | 3,37E-05 |
| 1810058I24Rik | 67705 | 41,912121 | -13,57532077 | 2,795861915 | 1,20E-06 | 3,46E-05 |
| Tmem29 | 382245 | 193,24341 | -10,22227422 | 2,110676892 | 1,28E-06 | 3,67E-05 |
| Kars | 85305 | 72,35526 | 20,24422198 | 4,181913594 | 1,29E-06 | 3,70E-05 |
| Gm6712 | 1,01E+08 | 283,65316 | -12,06679086 | 2,496583103 | 1,34E-06 | 3,84E-05 |
| Bop1 | 12181 | 66,196062 | 20,13582243 | 4,169414527 | 1,37E-06 | 3,90E-05 |
| Magohb | 66441 | 34,782418 | 20,79875349 | 4,309166661 | 1,39E-06 | 3,95E-05 |
| Tor1a | 30931 | 20019,78 | -13,36990937 | 2,770440528 | 1,39E-06 | 3,95E-05 |
| Crebrf | 77128 | 233,70329 | 8,584309709 | 1,77924091 | 1,40E-06 | 3,96E-05 |
| Adar | 56417 | 1128,1591 | -11,47626222 | 2,379982966 | 1,42E-06 | 4,00E-05 |
| Zfp57 | 22715 | 305,33732 | 8,094135021 | 1,678929949 | 1,43E-06 | 4,01E-05 |
| Serping1 | 12258 | 40,939568 | 20,8462618 | 4,325796524 | 1,44E-06 | 4,04E-05 |
| Rnf10 | 50849 | 161,84158 | 8,451495584 | 1,755405801 | 1,48E-06 | 4,11E-05 |
| Rcan3 | 53902 | 13,629184 | 15,17323216 | 3,151660332 | 1,48E-06 | 4,11E-05 |
| Fkbp7 | 14231 | 23,225989 | 20,57328931 | 4,284377343 | 1,57E-06 | 4,37E-05 |
| Zfp523 | 224656 | 133,15633 | 20,84772404 | 4,34889081 | 1,64E-06 | 4,52E-05 |

| | | | | | | |
|---|---|---|---|---|---|---|
| Tmem81 | 74626 | 137,18492 | 20,88771378 | 4,356822827 | 1,63E-06 | 4,52E-05 |
| Hspa1b | 15511 | 344,63225 | 8,255604538 | 1,724131633 | 1,68E-06 | 4,63E-05 |
| Alg1 | 208211 | 19,937005 | -20,57927894 | 4,301328253 | 1,71E-06 | 4,71E-05 |
| Ccl3 | 20302 | 48,606965 | 20,93736997 | 4,387034125 | 1,82E-06 | 4,98E-05 |
| Ankhd1 | 108857 | 384,22579 | 8,77313615 | 1,839738407 | 1,85E-06 | 5,07E-05 |
| Zfand5 | 22682 | 361,51077 | 8,645583529 | 1,816843553 | 1,95E-06 | 5,31E-05 |
| Uqcrh | 66576 | 515,99973 | 7,962644579 | 1,67687978 | 2,05E-06 | 5,57E-05 |
| Ifitm3 | 66141 | 241,5312 | 10,8398971 | 2,283511326 | 2,06E-06 | 5,59E-05 |
| Hnrnpa0 | 77134 | 20893,512 | -8,787195646 | 1,853168943 | 2,12E-06 | 5,73E-05 |
| Greb1 | 268527 | 241,00501 | -10,00309644 | 2,120843745 | 2,40E-06 | 6,46E-05 |
| Ier2 | 15936 | 316,67089 | 12,35022067 | 2,619064179 | 2,41E-06 | 6,48E-05 |
| Igtp | 16145 | 53,755062 | 20,28345662 | 4,317926068 | 2,63E-06 | 7,06E-05 |
| Ypel2 | 77864 | 362,50949 | 8,391080386 | 1,791743746 | 2,82E-06 | 7,55E-05 |
| Dctn6 | 22428 | 111,86666 | 19,39136699 | 4,146790119 | 2,92E-06 | 7,79E-05 |
| 2510002D24Rik | 72307 | 470,21798 | 12,49623377 | 2,674326069 | 2,97E-06 | 7,91E-05 |
| P2rx6 | 18440 | 54,401195 | 20,36221875 | 4,363551293 | 3,06E-06 | 8,13E-05 |
| Tm9sf1 | 74140 | 18824,837 | -10,12617202 | 2,170286752 | 3,07E-06 | 8,13E-05 |
| Nrf1 | 18181 | 7526,0367 | -10,65312562 | 2,287515054 | 3,21E-06 | 8,46E-05 |
| Zfp644 | 52397 | 444,90008 | 10,23806442 | 2,201333755 | 3,31E-06 | 8,70E-05 |
| Nbr1 | 17966 | 811,61575 | 11,04880025 | 2,379710033 | 3,44E-06 | 9,01E-05 |
| Tcf25 | 66855 | 17104,462 | -7,736000477 | 1,683893907 | 4,35E-06 | 0,000114 |
| Slc25a11 | 67863 | 25110,238 | -9,131715144 | 1,996194768 | 4,77E-06 | 0,000125 |
| Cfap74 | 544678 | 33632,536 | -11,15533298 | 2,438989313 | 4,79E-06 | 0,000125 |
| Zkscan17 | 268417 | 123,7946 | 10,63090989 | 2,327265434 | 4,92E-06 | 0,000128 |
| Rassf2 | 215653 | 226,13818 | 8,435476245 | 1,851190322 | 5,19E-06 | 0,000135 |
| Setd2 | 235626 | 218,40844 | 8,627278319 | 1,893967487 | 5,24E-06 | 0,000135 |
| Hint1 | 15254 | 187,28454 | 9,98291687 | 2,194464342 | 5,39E-06 | 0,000139 |
| Gsr | 14782 | 36,292705 | -10,94816363 | 2,407740066 | 5,44E-06 | 0,00014 |
| Med9 | 192191 | 72,557301 | 19,82874014 | 4,361858118 | 5,47E-06 | 0,00014 |
| Eif3i | 54709 | 198,4653 | 9,625907899 | 2,11832537 | 5,52E-06 | 0,000141 |
| Cyp2j12 | 242546 | 48,570737 | 19,59759638 | 4,321534339 | 5,76E-06 | 0,000147 |
| Pogk | 71592 | 159,95295 | 8,401129037 | 1,855642789 | 5,97E-06 | 0,000152 |
| Ttc4 | 72354 | 211,25423 | 8,092636511 | 1,787640125 | 5,98E-06 | 0,000152 |

| | | | | | | |
|---|---|---|---|---|---|---|
| Wasf3 | 245880 | 1947,9369 | -8,00779181 | 1,771421969 | 6,17E-06 | 0,000156 |
| Eno4 | 226265 | 752,72133 | 10,81340887 | 2,39261259 | 6,20E-06 | 0,000156 |
| Pigf | 18701 | 15,442186 | 13,85588393 | 3,067140982 | 6,26E-06 | 0,000157 |
| Smco3 | 654818 | 29,789233 | 19,81883486 | 4,386751413 | 6,25E-06 | 0,000157 |
| Iscu | 66383 | 241,51579 | 10,03301651 | 2,221178155 | 6,27E-06 | 0,000157 |
| Rbm28 | 68272 | 168,12427 | 8,052616166 | 1,783833879 | 6,36E-06 | 0,000159 |
| Lrrc4b | 272381 | 101,88072 | -6,979981662 | 1,548675405 | 6,57E-06 | 0,000164 |
| Ak4 | 11639 | 1939,0711 | -8,495320132 | 1,885667644 | 6,63E-06 | 0,000165 |
| Ociad1 | 68095 | 213,29373 | 7,914130585 | 1,757824721 | 6,72E-06 | 0,000167 |
| Gm38504 | 1,03E+08 | 32,351883 | 11,23314008 | 2,497574528 | 6,87E-06 | 0,00017 |
| Bcap29 | 12033 | 159,20578 | 8,508942432 | 1,894146899 | 7,05E-06 | 0,000174 |
| Asb8 | 78541 | 95,885142 | -8,261759651 | 1,845127877 | 7,55E-06 | 0,000186 |
| Cdkl4 | 381113 | 27,04635 | 19,511277 | 4,363735211 | 7,78E-06 | 0,000191 |
| Zer1 | 227693 | 742,89937 | -9,540690171 | 2,13762427 | 8,07E-06 | 0,000198 |
| Hspa1a | 193740 | 265,86313 | 9,412533787 | 2,110932105 | 8,24E-06 | 0,000201 |
| 5730408K05Rik | 67531 | 214,09471 | -9,319466136 | 2,092888146 | 8,47E-06 | 0,000206 |
| Zfp839 | 72805 | 508,55053 | -9,380690009 | 2,106644256 | 8,47E-06 | 0,000206 |
| Tsacc | 76927 | 49,570013 | 19,36728627 | 4,351457728 | 8,56E-06 | 0,000208 |
| Zkscan3 | 72739 | 393,74433 | 6,569878883 | 1,476840826 | 8,64E-06 | 0,000208 |
| Pih1d2 | 72614 | 23,590901 | 19,41428811 | 4,363784362 | 8,63E-06 | 0,000208 |
| Rgs10 | 67865 | 18,360286 | 19,51850838 | 4,387370601 | 8,64E-06 | 0,000208 |
| Tmem167 | 66074 | 24656,415 | -10,45534703 | 2,35108817 | 8,71E-06 | 0,000209 |
| Guf1 | 231279 | 2427,2399 | -11,19211584 | 2,517914075 | 8,79E-06 | 0,000211 |
| Xxylt1 | 268880 | 3574,0789 | -10,59943769 | 2,388032482 | 9,06E-06 | 0,000216 |
| Mrps21 | 66292 | 10098,747 | -11,46257967 | 2,585622431 | 9,28E-06 | 0,000221 |
| Parp14 | 547253 | 2884,1357 | -7,489465699 | 1,689956306 | 9,35E-06 | 0,000222 |
| 1700028K03Rik | 76421 | 47,168554 | 19,28322149 | 4,352845347 | 9,42E-06 | 0,000224 |
| Mras | 17532 | 1443,7703 | -9,501909916 | 2,146052417 | 9,53E-06 | 0,000226 |
| Klhl9 | 242521 | 29574,121 | -8,637264515 | 1,95266263 | 9,72E-06 | 0,00023 |
| Snrpe | 20643 | 89,155649 | -10,3400492 | 2,343199244 | 1,02E-05 | 0,00024 |
| Chchd2 | 14004 | 12283,971 | -6,647884063 | 1,507350915 | 1,03E-05 | 0,000243 |
| Gm52103 | 1,15E+08 | 125,67858 | -10,00793694 | 2,271961745 | 1,06E-05 | 0,000247 |
| Cipc | 217732 | 216,11596 | 8,393398822 | 1,90564402 | 1,06E-05 | 0,000247 |

| | | | | | | |
|---|---|---|---|---|---|---|
| Cep170 | 545389 | 545,32637 | 8,209188107 | 1,863556285 | 1,06E-05 | 0,000247 |
| Rps6ka1 | 20111 | 50,929144 | 19,13743036 | 4,345396795 | 1,06E-05 | 0,000247 |
| Ppp2cb | 19053 | 215,45954 | 9,635432441 | 2,191666411 | 1,10E-05 | 0,000256 |
| Rnf25 | 57751 | 54,690239 | -11,13477401 | 2,536273485 | 1,13E-05 | 0,000263 |
| Cdc40 | 71713 | 124,46219 | 8,820423972 | 2,012228665 | 1,17E-05 | 0,00027 |
| Tmem223 | 66836 | 402,13517 | 9,553082412 | 2,182978509 | 1,21E-05 | 0,000279 |
| Sobp | 109205 | 198,08855 | -9,188930374 | 2,102066128 | 1,23E-05 | 0,000284 |
| Supt6 | 20926 | 267,83031 | -8,281822128 | 1,895531966 | 1,25E-05 | 0,000287 |
| Afap1 | 70292 | 443,61317 | -7,380959154 | 1,692147761 | 1,29E-05 | 0,000295 |
| Tspan13 | 66109 | 90,092844 | 10,20582712 | 2,341372462 | 1,31E-05 | 0,000299 |
| Rhbdl2 | 230726 | 34,291344 | 18,96012847 | 4,352640714 | 1,32E-05 | 0,000302 |
| Pip4p2 | 72519 | 2836,1379 | -8,495966842 | 1,950880061 | 1,33E-05 | 0,000303 |
| Zmat1 | 215693 | 141,22199 | -10,30641731 | 2,370218847 | 1,37E-05 | 0,000312 |
| Suds3 | 71954 | 389,8066 | -8,946405477 | 2,061066541 | 1,42E-05 | 0,000322 |
| Ccdc32 | 269336 | 196,49869 | -9,303604409 | 2,146349421 | 1,46E-05 | 0,00033 |
| Aaas | 223921 | 19,949702 | 18,76484315 | 4,329367274 | 1,46E-05 | 0,00033 |
| Tmem237 | 381259 | 131,01737 | 9,933329929 | 2,294527712 | 1,50E-05 | 0,000337 |
| Taldo1 | 21351 | 423,68366 | 8,779016829 | 2,029379757 | 1,52E-05 | 0,000341 |
| Nfe2l2 | 18024 | 36808,273 | -8,223694759 | 1,90178213 | 1,53E-05 | 0,000343 |
| Clip1 | 56430 | 377,10872 | 7,850678707 | 1,816805771 | 1,55E-05 | 0,000346 |
| Vamp2 | 22318 | 3582,2695 | -8,601312372 | 1,990603182 | 1,55E-05 | 0,000346 |
| Tmem120b | 330189 | 21,015649 | 13,40075011 | 3,101305386 | 1,55E-05 | 0,000346 |
| Gm52092 | 1,15E+08 | 146,41531 | -8,621641372 | 1,996430001 | 1,57E-05 | 0,000349 |
| Pcx | 18563 | 111,60141 | -8,351118589 | 1,935567199 | 1,60E-05 | 0,000354 |
| Tmem251 | 320351 | 97,256665 | -9,224510158 | 2,141154061 | 1,65E-05 | 0,000364 |
| Rbm25 | 67039 | 37462,228 | -7,024732838 | 1,633877141 | 1,71E-05 | 0,000378 |
| Jmjd6 | 107817 | 114,52915 | 9,406230812 | 2,18917735 | 1,73E-05 | 0,000381 |
| Tubb5 | 22154 | 124,32058 | 9,150631812 | 2,130478747 | 1,75E-05 | 0,000382 |
| Frg1 | 14300 | 466,73445 | 8,532777085 | 1,986609476 | 1,75E-05 | 0,000382 |
| Dctn3 | 53598 | 13463,327 | -10,43900152 | 2,430314158 | 1,74E-05 | 0,000382 |
| Dnah12 | 110083 | 112,19223 | 9,344469759 | 2,175990675 | 1,75E-05 | 0,000382 |
| Nrxn2 | 18190 | 939,10479 | -7,453798779 | 1,738296722 | 1,80E-05 | 0,000392 |
| Vsnl1 | 26950 | 28,752465 | 18,5969271 | 4,338405047 | 1,81E-05 | 0,000394 |

| | | | | | | |
|---|---|---|---|---|---|---|
| Pecr | 111175 | 35,199774 | 18,6804603 | 4,360525795 | 1,84E-05 | 0,000398 |
| Eif4enif1 | 74203 | 55,893442 | -8,845372567 | 2,06606957 | 1,86E-05 | 0,000401 |
| Ndp | 17986 | 12204,257 | -10,12093064 | 2,364154737 | 1,86E-05 | 0,000401 |
| Tceal3 | 594844 | 99,259524 | 10,82885049 | 2,534879462 | 1,94E-05 | 0,000417 |
| Inpp5f | 101490 | 499,255 | 8,998434217 | 2,10961326 | 2,00E-05 | 0,000429 |
| Ep400 | 75560 | 141,44426 | 7,924112503 | 1,85959131 | 2,03E-05 | 0,000433 |
| Atp1b1 | 11931 | 246,01526 | 8,150183872 | 1,912680759 | 2,03E-05 | 0,000433 |
| Hax1 | 23897 | 15604,619 | -10,72944129 | 2,517462391 | 2,03E-05 | 0,000433 |
| Akip1 | 57373 | 47,019466 | 18,59722827 | 4,363526412 | 2,03E-05 | 0,000433 |
| Utp25 | 215193 | 2223,3634 | -9,243975862 | 2,170306451 | 2,05E-05 | 0,000436 |
| Tmem65 | 74868 | 10721,004 | -10,23506578 | 2,40528763 | 2,09E-05 | 0,000443 |
| Cfl1 | 12631 | 17824,379 | -7,105435051 | 1,67300209 | 2,17E-05 | 0,000458 |
| LOC118568253 | 1,19E+08 | 50,919489 | -10,1163211 | 2,383859688 | 2,20E-05 | 0,000464 |
| Atp6v0e2 | 76252 | 26332,233 | -9,346459485 | 2,202918067 | 2,21E-05 | 0,000465 |
| Apba2 | 11784 | 199,25271 | 7,242276944 | 1,708379465 | 2,24E-05 | 0,000471 |
| Col27a1 | 373864 | 79,679338 | 7,948484487 | 1,875543183 | 2,26E-05 | 0,000473 |
| Ctu2 | 66965 | 614,85052 | -10,95403483 | 2,585677998 | 2,27E-05 | 0,000475 |
| Atp1b2 | 11932 | 56307,903 | -6,897596445 | 1,630138364 | 2,32E-05 | 0,000485 |
| Mrpl11 | 66419 | 808,55028 | -10,36557791 | 2,450018777 | 2,33E-05 | 0,000485 |
| Arhgef40 | 268739 | 239,14824 | -7,735707606 | 1,830615723 | 2,38E-05 | 0,000495 |
| Ccdc12 | 72654 | 165,88032 | 9,537816072 | 2,258377402 | 2,41E-05 | 0,000499 |
| Uso1 | 56041 | 126,0064 | 10,09818452 | 2,390962257 | 2,41E-05 | 0,000499 |
| Rap1b | 215449 | 27,885119 | 18,24528929 | 4,322132098 | 2,43E-05 | 0,000502 |
| Ythdc1 | 231386 | 222,64155 | 6,474751294 | 1,5353172 | 2,47E-05 | 0,00051 |
| Hepacam | 72927 | 21251,691 | -6,399632381 | 1,519646099 | 2,54E-05 | 0,000523 |
| Zmym1 | 68310 | 64,93675 | 10,48818484 | 2,49106301 | 2,55E-05 | 0,000524 |
| B3galt1 | 26877 | 231,58031 | 6,605469072 | 1,569015512 | 2,55E-05 | 0,000524 |
| Dusp3 | 72349 | 195,6987 | 7,363621605 | 1,750970221 | 2,61E-05 | 0,000533 |
| Cox5a | 12858 | 10415,74 | -8,209955726 | 1,952540961 | 2,61E-05 | 0,000534 |
| Zmynd11 | 66505 | 203,09763 | 7,326283404 | 1,74345902 | 2,64E-05 | 0,000539 |
| Ube2d2a | 56550 | 159,41862 | 7,074348217 | 1,683728638 | 2,65E-05 | 0,000539 |
| 2310069G16Rik | 69659 | 12,918385 | 13,56732411 | 3,229903736 | 2,66E-05 | 0,00054 |
| Lrriq1 | 74978 | 191,40701 | 8,969720489 | 2,137118644 | 2,70E-05 | 0,000548 |

| | | | | | | |
|---|---|---|---|---|---|---|
| Fgfr1op2 | 67529 | 268,84168 | 9,922621851 | 2,364626754 | 2,71E-05 | 0,000548 |
| LOC118568733 | 1,19E+08 | 23,944125 | 18,24083416 | 4,355901978 | 2,82E-05 | 0,000569 |
| Ccnk | 12454 | 1071,084 | -8,340479753 | 1,993717132 | 2,87E-05 | 0,000578 |
| Pa2g4 | 18813 | 132,83502 | 7,542390068 | 1,804417805 | 2,92E-05 | 0,000585 |
| Zfp664 | 269704 | 131,21201 | 10,63776422 | 2,545290054 | 2,92E-05 | 0,000585 |
| Gm17484 | 1,01E+08 | 22,488418 | 18,24088221 | 4,364136666 | 2,92E-05 | 0,000585 |
| 1700113A16Rik | 76642 | 28,060682 | 18,10100319 | 4,34344989 | 3,08E-05 | 0,000615 |
| Gm51873 | 1,15E+08 | 795,12572 | -6,990937342 | 1,679980711 | 3,16E-05 | 0,000631 |
| Gnb2 | 14693 | 285,15202 | 6,585327899 | 1,583187114 | 3,19E-05 | 0,000633 |
| Rbbp5 | 213464 | 289,78778 | -9,701814374 | 2,332415208 | 3,19E-05 | 0,000633 |
| Gapdh | 14433 | 12449,621 | -6,221878334 | 1,497376717 | 3,25E-05 | 0,000644 |
| 0610010F05Rik | 71675 | 1973,8024 | -9,413767732 | 2,267088168 | 3,29E-05 | 0,000651 |
| Suz12 | 52615 | 238,67683 | 6,559638894 | 1,580992285 | 3,34E-05 | 0,000659 |
| Mtf2 | 17765 | 427,15302 | -8,435765133 | 2,03340451 | 3,35E-05 | 0,000659 |
| Antkmt | 214917 | 112,59536 | 8,79772401 | 2,122696132 | 3,40E-05 | 0,000668 |
| Kifbp | 72320 | 473,60501 | 7,701074335 | 1,858115654 | 3,40E-05 | 0,000668 |
| Gm49668 | 1,15E+08 | 2706,1781 | -9,433732623 | 2,276590815 | 3,42E-05 | 0,000669 |
| Stk17b | 98267 | 209,06281 | -9,993368346 | 2,413701184 | 3,47E-05 | 0,000678 |
| Sub1 | 20024 | 15609,658 | -7,629191132 | 1,844415806 | 3,53E-05 | 0,000688 |
| Spcs3 | 76687 | 133,61241 | 8,668162755 | 2,10365828 | 3,78E-05 | 0,000736 |
| Hnrnpab | 15384 | 65,740848 | 10,39234339 | 2,525144316 | 3,86E-05 | 0,00075 |
| Fcf1 | 73736 | 13251,73 | -8,598826949 | 2,091193069 | 3,92E-05 | 0,000759 |
| Xbp1 | 22433 | 209,93931 | 9,1167416 | 2,21708601 | 3,92E-05 | 0,000759 |
| Cyb5b | 66427 | 258,27905 | 7,417126277 | 1,804135121 | 3,94E-05 | 0,00076 |
| Zmym6 | 100177 | 110,2874 | 9,042888552 | 2,200624967 | 3,97E-05 | 0,000764 |
| Med1 | 19014 | 145,1474 | 8,40240669 | 2,044778153 | 3,97E-05 | 0,000764 |
| Fam117b | 72750 | 288,24117 | 7,838880322 | 1,908102541 | 3,99E-05 | 0,000766 |
| Rab8a | 17274 | 202,67051 | 9,282454619 | 2,260078142 | 4,01E-05 | 0,000768 |
| Ptk2 | 14083 | 422,0789 | 6,643392572 | 1,619245181 | 4,08E-05 | 0,000781 |
| Myo9a | 270163 | 6629,4584 | -8,703162326 | 2,123015999 | 4,14E-05 | 0,000788 |
| Caskin1 | 268932 | 4779,5059 | -8,076184254 | 1,969955965 | 4,14E-05 | 0,000788 |
| Ubl4a | 27643 | 11762,032 | -10,39392946 | 2,534964675 | 4,13E-05 | 0,000788 |
| Eef1b2 | 55949 | 190,00716 | 6,99438644 | 1,709239412 | 4,27E-05 | 0,000811 |

| | | | | | | |
|---|---|---|---|---|---|---|
| Fdps | 110196 | 671,05771 | -8,738958906 | 2,139946221 | 4,43E-05 | 0,00084 |
| Ttc14 | 67120 | 376,61497 | 6,97214516 | 1,708431363 | 4,48E-05 | 0,000848 |
| Aldoa | 11674 | 42924,012 | -7,862035375 | 1,931190939 | 4,68E-05 | 0,000883 |
| Gm11655 | 1,15E+08 | 273,6823 | -7,485988169 | 1,840223014 | 4,74E-05 | 0,000892 |
| Mrpl53 | 68499 | 10411,014 | -10,51711374 | 2,585483852 | 4,75E-05 | 0,000892 |
| Slc23a2 | 54338 | 158,14941 | 8,008498382 | 1,970857597 | 4,84E-05 | 0,000907 |
| Gm31328 | 1,03E+08 | 171,00253 | -11,11233169 | 2,735310082 | 4,85E-05 | 0,000909 |
| Tbc1d19 | 67249 | 41461,579 | -9,263343599 | 2,280547966 | 4,87E-05 | 0,00091 |
| Zfand2a | 100494 | 326,87602 | -10,66061917 | 2,62636507 | 4,93E-05 | 0,000919 |
| Dars | 226414 | 1134,5897 | -9,354534084 | 2,306211967 | 4,99E-05 | 0,000929 |
| Dock9 | 105445 | 22158,885 | -9,937869707 | 2,451084309 | 5,02E-05 | 0,000934 |
| Plekhb2 | 226971 | 301,83994 | 9,162262021 | 2,260660783 | 5,06E-05 | 0,000939 |
| Secisbp2 | 75420 | 155,02799 | 7,148353807 | 1,765093638 | 5,13E-05 | 0,000949 |
| Gm867 | 333670 | 156,55003 | 9,002513398 | 2,223387582 | 5,14E-05 | 0,000951 |
| Phf2 | 18676 | 149,78573 | 9,399513671 | 2,322298724 | 5,18E-05 | 0,000955 |
| Wbp2 | 22378 | 14583,093 | -8,044368604 | 1,991054697 | 5,34E-05 | 0,000983 |
| Tbrg1 | 21376 | 132,53324 | 10,42516286 | 2,58204018 | 5,40E-05 | 0,000993 |
| Gm2436 | 1E+08 | 1064,478 | -10,54095904 | 2,613312253 | 5,49E-05 | 0,001006 |
| Gm2446 | 1E+08 | 1064,478 | -10,54095904 | 2,613312253 | 5,49E-05 | 0,001006 |
| Rpl14 | 67115 | 258,44942 | 7,119772293 | 1,765778912 | 5,53E-05 | 0,001011 |
| Tmem138 | 72982 | 58,874063 | 8,708156759 | 2,162457658 | 5,65E-05 | 0,001031 |
| Nudc | 18221 | 325,0773 | 6,786230043 | 1,685362664 | 5,66E-05 | 0,001031 |
| Hipk2 | 15258 | 508,13017 | 5,786991833 | 1,438190686 | 5,73E-05 | 0,001041 |
| Prorsd1 | 67939 | 81,764096 | -9,208818854 | 2,290646927 | 5,82E-05 | 0,001055 |
| Dnah2 | 327954 | 65,437307 | 8,946262228 | 2,228534649 | 5,96E-05 | 0,00108 |
| Nudt19 | 110959 | 356,51541 | -9,733500303 | 2,425208 | 5,98E-05 | 0,001082 |
| Mospd1 | 70380 | 82,523131 | 9,482253225 | 2,363705392 | 6,03E-05 | 0,001088 |
| Srsf11 | 69207 | 7367,9799 | -5,560390366 | 1,386346639 | 6,05E-05 | 0,00109 |
| Vps53 | 68299 | 167,42127 | -10,95523478 | 2,731647638 | 6,06E-05 | 0,00109 |
| Tnip1 | 57783 | 45,988031 | 11,56194337 | 2,883741173 | 6,09E-05 | 0,001093 |
| Hmgcs1 | 208715 | 4459,8362 | -6,82918147 | 1,704388962 | 6,15E-05 | 0,001101 |
| Frmpd1 | 666060 | 107,83852 | -6,870621639 | 1,714774453 | 6,16E-05 | 0,001101 |
| Lrrc71 | 74485 | 1382,751 | -7,860743823 | 1,964133062 | 6,28E-05 | 0,001121 |

| | | | | | | |
|---|---|---|---|---|---|---|
| Tuba4a | 22145 | 2623,1091 | -8,116462072 | 2,02930251 | 6,34E-05 | 0,001131 |
| Hmgb2 | 97165 | 349,26249 | 10,80656875 | 2,70296262 | 6,39E-05 | 0,001136 |
| H2az2 | 77605 | 11315,492 | -7,014959083 | 1,755318215 | 6,43E-05 | 0,001142 |
| Tchp | 77832 | 169,55459 | 10,47517285 | 2,621813678 | 6,46E-05 | 0,001145 |
| Tceal1 | 237052 | 165,24369 | 9,211783804 | 2,306502662 | 6,50E-05 | 0,00115 |
| Mbnl2 | 105559 | 841,68739 | 5,304865584 | 1,32841837 | 6,51E-05 | 0,001151 |
| Blvrb | 233016 | 307,85651 | -9,711691991 | 2,432222396 | 6,53E-05 | 0,001151 |
| Rmnd5a | 68477 | 257,85666 | 9,718076405 | 2,435099357 | 6,58E-05 | 0,001159 |
| Dnah3 | 381917 | 53,780674 | 7,513807653 | 1,884993385 | 6,72E-05 | 0,00118 |
| Prr5 | 109270 | 252,62948 | -8,177476354 | 2,051997707 | 6,74E-05 | 0,001183 |
| Rxrb | 20182 | 327,4546 | -9,196943348 | 2,308277521 | 6,77E-05 | 0,001185 |
| Smim20 | 66278 | 16729,159 | -10,42325995 | 2,616364295 | 6,78E-05 | 0,001185 |
| Hint3 | 66847 | 233,90877 | 10,59759832 | 2,662405555 | 6,88E-05 | 0,0012 |
| Gm51971 | 1,15E+08 | 5001,0546 | -8,369832926 | 2,107804281 | 7,16E-05 | 0,001247 |
| Nop14 | 75416 | 205,7323 | 10,51793136 | 2,651068576 | 7,27E-05 | 0,001263 |
| S100a16 | 67860 | 15697,793 | -7,479155852 | 1,885982877 | 7,32E-05 | 0,001268 |
| Ptrh2 | 217057 | 141,20598 | -9,72335598 | 2,451847991 | 7,32E-05 | 0,001268 |
| Snx25 | 102141 | 482,00212 | -9,271395305 | 2,339136823 | 7,38E-05 | 0,001277 |
| Uqcr10 | 66152 | 5224,5178 | -6,069987299 | 1,531981131 | 7,43E-05 | 0,001282 |
| Timm10b | 14356 | 91,952906 | 7,978526621 | 2,014546473 | 7,48E-05 | 0,001289 |
| Clint1 | 216705 | 27247,46 | -10,14440303 | 2,564019231 | 7,61E-05 | 0,001309 |
| Gstm1 | 14862 | 40598,73 | -6,247096957 | 1,579700993 | 7,67E-05 | 0,001317 |
| Bin3 | 57784 | 105,19348 | -10,72457084 | 2,71505817 | 7,81E-05 | 0,001338 |
| Rpf2 | 67239 | 209,41765 | 9,383165997 | 2,375282732 | 7,80E-05 | 0,001338 |
| Noc4l | 100608 | 19274,976 | -16,37550708 | 4,148193677 | 7,89E-05 | 0,001349 |
| Vegfa | 22339 | 2289,5739 | -6,554991018 | 1,662306357 | 8,04E-05 | 0,001371 |
| Scg3 | 20255 | 36123,629 | -5,814216348 | 1,4754636 | 8,13E-05 | 0,001384 |
| Tspan15 | 70423 | 28464,96 | -8,32475783 | 2,114121772 | 8,23E-05 | 0,001399 |
| Prdx6 | 11758 | 19002,717 | -6,373215637 | 1,619000939 | 8,27E-05 | 0,001403 |
| Etfb | 110826 | 7694,1121 | -8,043390124 | 2,043795003 | 8,30E-05 | 0,001406 |
| Atf7ip | 54343 | 245,40032 | 6,315474267 | 1,605207321 | 8,34E-05 | 0,001411 |
| Tex264 | 21767 | 18671,482 | -9,189829358 | 2,337928781 | 8,47E-05 | 0,001429 |
| Mtmr6 | 219135 | 255,26809 | 9,348903367 | 2,378590194 | 8,48E-05 | 0,001429 |

| | | | | | | |
|---|---|---|---|---|---|---|
| Lancl1 | 14768 | 241,40963 | 9,72726346 | 2,477731834 | 8,64E-05 | 0,001454 |
| St7 | 64213 | 103,85742 | 8,696756446 | 2,218219406 | 8,83E-05 | 0,001484 |
| Cerox1 | 72834 | 133,1671 | 8,958224291 | 2,287745239 | 9,01E-05 | 0,001511 |
| Klf2 | 16598 | 164,12192 | 11,04435869 | 2,821429983 | 9,06E-05 | 0,001517 |
| Stard3nl | 76205 | 446,4901 | 8,61915138 | 2,203046998 | 9,14E-05 | 0,001528 |
| Eif2b3 | 108067 | 207,42046 | -10,12917057 | 2,590896703 | 9,25E-05 | 0,001543 |
| Pitpnm2 | 19679 | 102,62954 | 8,698082392 | 2,226970795 | 9,39E-05 | 0,001565 |
| Nova1 | 664883 | 171,5486 | 6,696723143 | 1,716885849 | 9,60E-05 | 0,001596 |
| Fermt2 | 218952 | 6339,4738 | -6,739507931 | 1,730869769 | 9,87E-05 | 0,001639 |
| Arrdc3 | 105171 | 576,54798 | 8,024572311 | 2,06364212 | 0,000101 | 0,001672 |
| Aph1a | 226548 | 89,913738 | 6,667520076 | 1,715091205 | 0,000101 | 0,001676 |
| 2310011J03Rik | 66374 | 161,21504 | 8,69478788 | 2,237386032 | 0,000102 | 0,00168 |
| Polr3b | 70428 | 93,023131 | -10,05968673 | 2,588405848 | 0,000102 | 0,00168 |
| Sfr1 | 67788 | 82,932849 | 7,723509178 | 1,989661762 | 0,000104 | 0,001707 |
| Kcnk2 | 16526 | 2399,9729 | -8,463167736 | 2,181478792 | 0,000105 | 0,00172 |
| Ccdc93 | 70829 | 158,47589 | 10,17014264 | 2,622408428 | 0,000105 | 0,001727 |
| Klhl11 | 217194 | 284,29667 | 7,498400533 | 1,934501598 | 0,000106 | 0,001739 |
| Enho | 69638 | 14880,949 | -7,712628142 | 1,99043644 | 0,000107 | 0,001745 |
| Bccip | 66165 | 151,45711 | 10,37241274 | 2,677345148 | 0,000107 | 0,001747 |
| Hdhd2 | 76987 | 130,84004 | 8,135509149 | 2,103401001 | 0,00011 | 0,001788 |
| Tmed4 | 103694 | 25716,596 | -8,438559041 | 2,181700747 | 0,00011 | 0,001788 |
| Pebp1 | 23980 | 233,89782 | 6,806439845 | 1,760689623 | 0,000111 | 0,001799 |
| Efnb3 | 13643 | 150,37292 | 7,935089322 | 2,052931048 | 0,000111 | 0,0018 |
| Vps29 | 56433 | 261,25592 | 7,149229643 | 1,849828968 | 0,000111 | 0,001801 |
| Snph | 241727 | 127,44037 | -8,039250325 | 2,0823521 | 0,000113 | 0,001829 |
| Acta2 | 11475 | 84,92942 | 8,946923261 | 2,318669795 | 0,000114 | 0,001841 |
| Zdhhc20 | 75965 | 300,21187 | 6,041729539 | 1,566789027 | 0,000115 | 0,001854 |
| Aldoc | 11676 | 52683,851 | -6,061782515 | 1,571862239 | 0,000115 | 0,001854 |
| Ccdc136 | 232664 | 112,21352 | 6,682707221 | 1,73615946 | 0,000119 | 0,001904 |
| Bace2 | 56175 | 165,7394 | 9,406531486 | 2,444065671 | 0,000119 | 0,001905 |
| Glul | 14645 | 29140,144 | -5,94399207 | 1,54465266 | 0,000119 | 0,001906 |
| Lsamp | 268890 | 31438,35 | -6,066993456 | 1,576922778 | 0,000119 | 0,001909 |
| Tmem256 | 69186 | 7784,5163 | -8,260110928 | 2,14720299 | 0,00012 | 0,001909 |

| | | | | | | |
|---|---|---|---|---|---|---|
| Dusp11 | 72102 | 136,69245 | 6,495650762 | 1,688802692 | 0,00012 | 0,001911 |
| Atf3 | 11910 | 455,90428 | 10,798917 | 2,808099953 | 0,00012 | 0,001913 |
| Aimp1 | 13722 | 101,75197 | 7,705770159 | 2,005127797 | 0,000122 | 0,001929 |
| Sp2 | 78912 | 41,723117 | -7,829277961 | 2,037756181 | 0,000122 | 0,001929 |
| Lgi3 | 213469 | 148,51228 | 8,143095855 | 2,119308051 | 0,000122 | 0,001929 |
| LOC118568500 | 1,19E+08 | 1532,1343 | -5,657752355 | 1,472352348 | 0,000122 | 0,001929 |
| Arfgef1 | 211673 | 122,03554 | 8,438107683 | 2,197134612 | 0,000123 | 0,001938 |
| Tenm3 | 23965 | 464,13505 | -6,603941951 | 1,720002083 | 0,000123 | 0,001943 |
| Copa | 12847 | 137,32755 | 7,968449744 | 2,077500019 | 0,000125 | 0,001971 |
| Tmcc2 | 68875 | 110,20555 | 8,806217292 | 2,296172085 | 0,000125 | 0,001971 |
| Ipo5 | 70572 | 139,82767 | 8,254069683 | 2,155209846 | 0,000128 | 0,002012 |
| Golim4 | 73124 | 54,057994 | 7,21317537 | 1,88367622 | 0,000129 | 0,002012 |
| Rftn2 | 74013 | 13247,812 | -7,787500773 | 2,03403539 | 0,000129 | 0,002012 |
| Grin3a | 242443 | 252,44878 | -9,218619633 | 2,407838332 | 0,000129 | 0,002012 |
| Man2b1 | 17159 | 245,41764 | -9,582510684 | 2,50310461 | 0,000129 | 0,002012 |
| Tubgcp2 | 74237 | 27,779783 | -8,539950151 | 2,231671035 | 0,00013 | 0,002021 |
| Csnk2a1 | 12995 | 414,98826 | 5,076477538 | 1,327983456 | 0,000132 | 0,002051 |
| Glyr1 | 74022 | 106,70077 | 6,961766152 | 1,822098869 | 0,000133 | 0,002064 |
| Nr4a1 | 15370 | 176,34996 | 8,13625077 | 2,129752985 | 0,000133 | 0,002065 |
| Vps41 | 218035 | 98,895007 | 8,774337192 | 2,298478172 | 0,000135 | 0,002085 |
| Nsmf | 56876 | 107,28167 | 6,758721078 | 1,772526907 | 0,000137 | 0,00212 |
| Scrg1 | 20284 | 10100,695 | -6,673961081 | 1,750595631 | 0,000138 | 0,002122 |
| Kremen1 | 84035 | 244,16294 | -7,885776164 | 2,068774359 | 0,000138 | 0,002124 |
| Cdr1 | 631990 | 100,51556 | 7,397524126 | 1,942161613 | 0,00014 | 0,002145 |
| Slc25a45 | 107375 | 58,280575 | -9,308178234 | 2,445863741 | 0,000141 | 0,00217 |
| Tmod1 | 21916 | 208,50606 | 10,74193303 | 2,825531658 | 0,000144 | 0,002202 |
| Sde2 | 208768 | 134,94706 | 8,177297732 | 2,153435277 | 0,000146 | 0,002238 |
| Git2 | 26431 | 133,02056 | 8,754866983 | 2,306925544 | 0,000148 | 0,002252 |
| Ibtk | 108837 | 387,19226 | 7,2028728 | 1,897871436 | 0,000148 | 0,002252 |
| Sod3 | 20657 | 159,74427 | 8,910005972 | 2,349836564 | 0,00015 | 0,002278 |
| Cenpc1 | 12617 | 76,305052 | 8,475203284 | 2,237248167 | 0,000152 | 0,002307 |
| Shtn1 | 71653 | 70,700854 | 9,678895379 | 2,556871895 | 0,000153 | 0,00233 |
| Vti1a | 53611 | 221,87822 | 6,668335041 | 1,762022174 | 0,000154 | 0,002335 |

| | | | | | | |
|---|---|---|---|---|---|---|
| Gpatch2l | 70373 | 68,851133 | 6,678270367 | 1,765287346 | 0,000155 | 0,002344 |
| Kndc1 | 76484 | 71,392854 | 7,053848933 | 1,865419519 | 0,000156 | 0,002354 |
| Pmm1 | 29858 | 15866,2 | -7,132960366 | 1,886320956 | 0,000156 | 0,002354 |
| Ruvbl2 | 20174 | 4137,8376 | -9,749868068 | 2,580903952 | 0,000158 | 0,002385 |
| Kctd1 | 106931 | 288,74608 | 7,878988508 | 2,08608255 | 0,000159 | 0,002389 |
| Ndufb7 | 66916 | 175,85639 | 8,782747 | 2,326440727 | 0,00016 | 0,0024 |
| Pam16 | 66449 | 6762,7064 | -9,377691092 | 2,484141365 | 0,00016 | 0,0024 |
| Gm4924 | 237412 | 405,77679 | 8,160391635 | 2,161916388 | 0,00016 | 0,0024 |
| Sat2 | 69215 | 185,76739 | -8,638688226 | 2,293693683 | 0,000166 | 0,002478 |
| Pggt1b | 225467 | 145,29085 | 7,044812001 | 1,870687634 | 0,000166 | 0,002478 |
| Msn | 17698 | 48,138906 | 8,788332615 | 2,334903804 | 0,000167 | 0,002494 |
| Abl1 | 11350 | 196,62773 | 7,64114041 | 2,031414275 | 0,000169 | 0,002515 |
| Ero1a | 50527 | 84,122269 | 7,809728345 | 2,077482346 | 0,00017 | 0,002534 |
| Spast | 50850 | 119,37674 | 8,394703111 | 2,233563467 | 0,000171 | 0,002538 |
| Fkbp8 | 14232 | 89,110208 | 7,106922145 | 1,891605878 | 0,000172 | 0,002541 |
| Utp14a | 72554 | 149,34959 | 9,076809118 | 2,41596346 | 0,000172 | 0,002541 |
| Wdr34 | 71820 | 154,54458 | -10,18483105 | 2,710711943 | 0,000172 | 0,002541 |
| Gm52313 | 1,15E+08 | 369,66227 | -8,084230236 | 2,152212084 | 0,000172 | 0,002545 |
| Ilrun | 224647 | 76,842265 | 8,493790148 | 2,263685674 | 0,000175 | 0,002583 |
| Cachd1 | 320508 | 399,1489 | 8,096750205 | 2,158716994 | 0,000176 | 0,002594 |
| 1810009A15Rik | 66276 | 106,52932 | 8,326850013 | 2,223868347 | 0,000181 | 0,002651 |
| Map2k7 | 26400 | 46,540771 | -7,577040148 | 2,023389025 | 0,000181 | 0,002651 |
| Stk4 | 58231 | 59,258563 | 8,183439429 | 2,185672245 | 0,000181 | 0,002651 |
| Rpl39 | 67248 | 77,456703 | 7,636671471 | 2,041841677 | 0,000184 | 0,00269 |
| Gm14399 | 1E+08 | 14115,295 | -8,176356511 | 2,18691908 | 0,000185 | 0,002701 |
| Synj1 | 104015 | 178,09439 | 6,864459929 | 1,837752388 | 0,000188 | 0,002735 |
| Cgrrf1 | 68755 | 84,702299 | 7,535849048 | 2,018400191 | 0,000189 | 0,002741 |
| Gtf3c2 | 71752 | 202,55266 | 7,652946599 | 2,04963408 | 0,000189 | 0,002741 |
| Coil | 12812 | 265,32635 | 8,389815126 | 2,246825465 | 0,000188 | 0,002741 |
| Hmgn3 | 94353 | 707,8255 | 5,297347642 | 1,41956264 | 0,00019 | 0,002757 |
| Txnl4a | 27366 | 150,98825 | 7,68464795 | 2,061405107 | 0,000193 | 0,002795 |
| Trip12 | 14897 | 112,99631 | 6,840705836 | 1,835194704 | 0,000193 | 0,002795 |
| Cav2 | 12390 | 15276,314 | -8,463404112 | 2,27107987 | 0,000194 | 0,002801 |

| | | | | | | |
|---|---|---|---|---|---|---|
| Gm41461 | 1,05E+08 | 795,9634 | -7,902918903 | 2,122274009 | 0,000196 | 0,002829 |
| Nono | 53610 | 161,58018 | 8,049249871 | 2,162231085 | 0,000197 | 0,002833 |
| Grk2 | 110355 | 63,440519 | -8,443925377 | 2,268137461 | 0,000197 | 0,002833 |
| Atp6v1f | 66144 | 21850,136 | -7,956706564 | 2,138512448 | 0,000199 | 0,002851 |
| Amer2 | 72125 | 277,5437 | -6,204427937 | 1,667837139 | 0,000199 | 0,002854 |
| Pdcd6 | 18570 | 20692,063 | -8,440489679 | 2,269489164 | 0,0002 | 0,002861 |
| Erich1 | 234086 | 101,78587 | 10,73196842 | 2,886371083 | 0,000201 | 0,002868 |
| Spin1 | 20729 | 490,39378 | 6,965381931 | 1,874086113 | 0,000202 | 0,00288 |
| Atp5b | 11947 | 20847,357 | -6,499765961 | 1,749499028 | 0,000203 | 0,002889 |
| Ndufb1 | 1,03E+08 | 4324,4242 | -8,556864967 | 2,303133022 | 0,000203 | 0,002889 |
| Csnk2a2 | 13000 | 219,02398 | 7,37447739 | 1,988799976 | 0,000209 | 0,002968 |
| Pcgf1 | 69837 | 35,444661 | 9,25080834 | 2,496265964 | 0,000211 | 0,002989 |
| Sel1l | 20338 | 217,24497 | 5,925319702 | 1,599098216 | 0,000211 | 0,00299 |
| Stk11 | 20869 | 75,738284 | 6,996069611 | 1,889377427 | 0,000213 | 0,003012 |
| Pip4k2b | 108083 | 286,7969 | -8,748204049 | 2,362533872 | 0,000213 | 0,003012 |
| Smc4 | 70099 | 538,00557 | 8,418313953 | 2,274828821 | 0,000215 | 0,003034 |
| Cdc73 | 214498 | 10477,746 | -8,015250105 | 2,166231514 | 0,000216 | 0,003036 |
| Hmbox1 | 219150 | 180,37577 | 8,101469263 | 2,190556761 | 0,000217 | 0,003052 |
| Slc25a36 | 192287 | 167,32992 | 6,467753217 | 1,749075683 | 0,000217 | 0,003055 |
| Rdh10 | 98711 | 101,43634 | 6,283412944 | 1,699434989 | 0,000218 | 0,003056 |
| Gm41559 | 1,05E+08 | 57,393259 | -7,488057098 | 2,026388307 | 0,00022 | 0,003075 |
| Mindy1 | 75007 | 120,14594 | 8,416266167 | 2,277691784 | 0,00022 | 0,003075 |
| Ndufaf7 | 73694 | 94,319306 | 7,892827752 | 2,136601668 | 0,000221 | 0,003082 |
| Dnajc8 | 68598 | 259,22304 | 5,431061168 | 1,470612574 | 0,000222 | 0,003091 |
| Ttll8 | 239591 | 72,522936 | 7,364661618 | 1,995571599 | 0,000224 | 0,003117 |
| Btf3l4 | 70533 | 64,048699 | -7,190721961 | 1,949282278 | 0,000225 | 0,003133 |
| Poli | 26447 | 84,829478 | -8,222498241 | 2,2299939 | 0,000227 | 0,003149 |
| Zfp846 | 244721 | 274,97742 | 7,538602595 | 2,046587283 | 0,00023 | 0,003191 |
| Chn2 | 69993 | 182,84316 | 7,944119336 | 2,157388641 | 0,000231 | 0,003202 |
| Nt5m | 103850 | 412,62601 | -9,98682802 | 2,712467624 | 0,000232 | 0,003203 |
| Pard3b | 72823 | 8556,8835 | -7,800626124 | 2,119295672 | 0,000233 | 0,003212 |
| Ube2d1 | 216080 | 147,24218 | 7,821518483 | 2,125409967 | 0,000233 | 0,003217 |
| U2af1 | 108121 | 1585,2247 | -7,27348614 | 1,976682386 | 0,000234 | 0,003217 |

| | | | | | | |
|---|---|---|---|---|---|---|
| Prkd3 | 75292 | 279,90537 | 8,92563033 | 2,42750519 | 0,000236 | 0,003248 |
| Tbca | 21371 | 257,74553 | -7,087551305 | 1,929477251 | 0,000239 | 0,003289 |
| Gnb5 | 14697 | 65,802951 | 8,052103934 | 2,193429489 | 0,000242 | 0,003314 |
| Appl1 | 72993 | 128,34683 | 7,26918725 | 1,980796652 | 0,000243 | 0,003325 |
| Gm26944 | 1,03E+08 | 541,73502 | -6,243695882 | 1,702255897 | 0,000245 | 0,003345 |
| Usp16 | 74112 | 208,46759 | 6,905591892 | 1,883030594 | 0,000245 | 0,003349 |
| Rnf13 | 24017 | 724,56191 | 5,3573639 | 1,46113216 | 0,000246 | 0,003353 |
| Med16 | 216154 | 21,974348 | -7,129234052 | 1,945997844 | 0,000249 | 0,003389 |
| Psmg1 | 56088 | 12,843311 | 11,16847623 | 3,050122807 | 0,000251 | 0,003409 |
| Twnk | 226153 | 71,266702 | 8,846850035 | 2,419601343 | 0,000256 | 0,003476 |
| Boc | 117606 | 42,301506 | 7,598127545 | 2,079072605 | 0,000258 | 0,00349 |
| Snrpd1 | 20641 | 150,6937 | 8,42451344 | 2,305138237 | 0,000258 | 0,00349 |
| Hdac3 | 15183 | 255,89726 | -6,001857134 | 1,643987399 | 0,000261 | 0,003537 |
| Nup85 | 445007 | 120,42807 | 9,082017087 | 2,488059298 | 0,000262 | 0,00354 |
| Nme2 | 18103 | 217,1242 | 5,763577829 | 1,58059427 | 0,000266 | 0,003583 |
| Nol9 | 74035 | 69,129121 | -6,826730419 | 1,87204385 | 0,000266 | 0,003583 |
| Gm35120 | 1,03E+08 | 64,674949 | -8,615076741 | 2,364128614 | 0,000268 | 0,003611 |
| Cabin1 | 104248 | 166,28781 | 5,964512041 | 1,637248966 | 0,000269 | 0,003621 |
| Lpin2 | 64898 | 169,50195 | 8,084973397 | 2,221679067 | 0,000274 | 0,00366 |
| Lrrcc1 | 71710 | 392,89808 | 7,787580646 | 2,140248533 | 0,000274 | 0,00366 |
| Fgf1 | 14164 | 392,95679 | 6,022433867 | 1,654755856 | 0,000273 | 0,00366 |
| Akr1a1 | 58810 | 221,34325 | 5,711462642 | 1,56971991 | 0,000274 | 0,00366 |
| Rev1 | 56210 | 489,95978 | -7,892757084 | 2,168735555 | 0,000273 | 0,00366 |
| Josd2 | 66124 | 10436,154 | -7,970727943 | 2,191541035 | 0,000276 | 0,003676 |
| Acad9 | 229211 | 60,074508 | -7,670610852 | 2,109194825 | 0,000276 | 0,003676 |
| Ssrp1 | 20833 | 145,05404 | 7,402950588 | 2,036423378 | 0,000278 | 0,003692 |
| Arl2bp | 107566 | 60,549632 | 7,529988959 | 2,072824052 | 0,00028 | 0,003724 |
| Usp47 | 74996 | 81,614703 | 7,070879549 | 1,948005443 | 0,000284 | 0,003761 |
| Srfbp1 | 67222 | 64,501443 | 7,887831294 | 2,174064699 | 0,000285 | 0,00378 |
| Esyt2 | 52635 | 105,51063 | 6,758501762 | 1,864865428 | 0,00029 | 0,003827 |
| Zswim8 | 268721 | 342,1724 | -7,592272217 | 2,095041264 | 0,00029 | 0,003827 |
| Cntnap2 | 66797 | 96,015818 | 8,594738566 | 2,371667102 | 0,00029 | 0,003827 |
| Hapln2 | 73940 | 248,59481 | 7,704564328 | 2,126479407 | 0,000291 | 0,003834 |

| | | | | | | |
|---|---|---|---|---|---|---|
| Zc3h15 | 69082 | 2082,3402 | -7,513091231 | 2,074280138 | 0,000292 | 0,003845 |
| Asf1a | 66403 | 58,013585 | -8,211486254 | 2,268332787 | 0,000295 | 0,00387 |
| Zfp148 | 22661 | 28731,779 | -7,256736377 | 2,005051331 | 0,000295 | 0,003877 |
| Npffr1 | 237362 | 1223,4982 | -6,106866412 | 1,687574641 | 0,000296 | 0,003879 |
| Gm33560 | 1,03E+08 | 80,966391 | -7,238357645 | 2,00112668 | 0,000298 | 0,003898 |
| Ghr | 14600 | 90,029957 | 6,784415314 | 1,876564724 | 0,0003 | 0,00392 |
| Lifr | 16880 | 222,44332 | -6,298344094 | 1,743622871 | 0,000304 | 0,003962 |
| Cse1l | 110750 | 223,28725 | 9,069220049 | 2,51197704 | 0,000306 | 0,00398 |
| Cyp7b1 | 13123 | 245,24618 | 6,625511222 | 1,835277587 | 0,000306 | 0,00398 |
| Zfp382 | 233060 | 91,893894 | -6,211729792 | 1,721014226 | 0,000307 | 0,00398 |
| Egfl6 | 54156 | 137,92813 | -8,528244147 | 2,362735533 | 0,000307 | 0,00398 |
| 1110032A03Rik | 68721 | 284,12319 | 8,643042977 | 2,394202708 | 0,000306 | 0,00398 |
| Ddr1 | 12305 | 130,61522 | 5,959225617 | 1,652149367 | 0,00031 | 0,004012 |
| Cst3 | 13010 | 232746,47 | -5,400784259 | 1,49818368 | 0,000312 | 0,004031 |
| Zfx | 22764 | 138,74176 | 7,736522165 | 2,146111624 | 0,000312 | 0,004031 |
| Bbs4 | 102774 | 243,75818 | 9,046763263 | 2,509729958 | 0,000313 | 0,004031 |
| Mir9-3hg | 101694 | 189,09509 | -7,179755254 | 1,992350362 | 0,000314 | 0,004042 |
| Tfdp2 | 211586 | 112,45838 | -6,304928706 | 1,751243752 | 0,000318 | 0,004085 |
| Olig2 | 50913 | 611,51183 | -6,876657473 | 1,910247954 | 0,000318 | 0,004085 |
| Arxes2 | 76976 | 30295,781 | -7,769819 | 2,158171193 | 0,000318 | 0,004085 |
| Rab7 | 19349 | 582,50106 | 6,135655156 | 1,706291252 | 0,000323 | 0,00414 |
| Skida1 | 72668 | 265,27604 | 8,450413641 | 2,350117066 | 0,000323 | 0,00414 |
| Slc9a7 | 236727 | 79,350896 | -7,853118167 | 2,185560611 | 0,000327 | 0,004175 |
| Rilpl2 | 80291 | 488,55849 | -8,741753991 | 2,433060676 | 0,000327 | 0,004175 |
| Prpsap2 | 212627 | 87,657031 | 8,822383633 | 2,457696847 | 0,000331 | 0,004221 |
| Tiam2 | 24001 | 359,84469 | -7,547429003 | 2,10296252 | 0,000332 | 0,004228 |
| Tmem19 | 67226 | 240,3185 | 6,559418987 | 1,828844755 | 0,000335 | 0,004251 |
| Rpl10a | 19896 | 234,42143 | 5,421450052 | 1,511652429 | 0,000335 | 0,004251 |
| Scnm1 | 69269 | 508,21699 | 7,370515896 | 2,055282101 | 0,000336 | 0,004251 |
| 4930570G19Rik | 329782 | 65,057702 | 8,707981549 | 2,428445286 | 0,000336 | 0,004251 |
| Zfp738 | 408068 | 11,563403 | 11,62954363 | 3,242973273 | 0,000336 | 0,004251 |
| Kctd2 | 70382 | 133,9467 | -8,459220712 | 2,361108523 | 0,00034 | 0,004297 |
| Htra1 | 56213 | 29254,41 | -5,247679808 | 1,464872991 | 0,000341 | 0,004298 |

| | | | | | | |
|---|---|---|---|---|---|---|
| Gria4 | 14802 | 117,52515 | -8,83364199 | 2,466488657 | 0,000342 | 0,004307 |
| P3h1 | 56401 | 66,546102 | 8,184524465 | 2,287372495 | 0,000346 | 0,004351 |
| Atraid | 381629 | 291,01779 | 6,723548468 | 1,878990642 | 0,000346 | 0,004351 |
| Rapgef2 | 76089 | 501,57222 | 8,298590863 | 2,319471996 | 0,000347 | 0,004351 |
| Sh3bp5l | 79566 | 120,21563 | 8,562137262 | 2,395034315 | 0,00035 | 0,004391 |
| Arrb1 | 109689 | 112,93716 | 5,737440032 | 1,604984719 | 0,000351 | 0,004391 |
| Srrm2 | 75956 | 13281,069 | -5,591587823 | 1,565339022 | 0,000354 | 0,00443 |
| BC005624 | 227707 | 147,25763 | 8,540886498 | 2,391424603 | 0,000355 | 0,004435 |
| Dpcd | 226162 | 177,19092 | 9,453703883 | 2,654105397 | 0,000368 | 0,004594 |
| Arpc5l | 74192 | 86,158243 | 6,983598992 | 1,960827527 | 0,000369 | 0,004594 |
| Tox2 | 269389 | 44,933376 | 7,449685412 | 2,092476932 | 0,000371 | 0,004612 |
| Chd4 | 107932 | 359,0558 | 4,698345269 | 1,320409957 | 0,000373 | 0,004641 |
| Vps37b | 330192 | 229,85667 | -6,405058873 | 1,801189186 | 0,000377 | 0,004672 |
| Mmgt2 | 216829 | 1924,5996 | -7,017035162 | 1,973387456 | 0,000377 | 0,004672 |
| Capza1 | 12340 | 168,2299 | 7,762233307 | 2,184085484 | 0,000379 | 0,004699 |
| Gm15478 | 1E+08 | 89,398831 | 6,227837479 | 1,753120053 | 0,000382 | 0,004721 |
| Gm39941 | 1,05E+08 | 166,68021 | 6,751468968 | 1,901684009 | 0,000385 | 0,004749 |
| Kdm7a | 338523 | 110,52867 | 7,951680624 | 2,23967777 | 0,000385 | 0,004749 |
| Prdx1 | 18477 | 19605,805 | -6,514480305 | 1,835474171 | 0,000386 | 0,004762 |
| A830082K12Rik | 320174 | 97,797467 | 8,231100156 | 2,319576903 | 0,000387 | 0,004768 |
| Fbxl20 | 72194 | 70,670716 | 6,919323137 | 1,951322151 | 0,000391 | 0,004809 |
| Echs1 | 93747 | 24347,347 | -8,270145976 | 2,33292257 | 0,000393 | 0,004821 |
| M6pr | 17113 | 153,06929 | 5,898008292 | 1,664140329 | 0,000394 | 0,00483 |
| Pnkp | 59047 | 145,49675 | 6,207464011 | 1,751643978 | 0,000394 | 0,004831 |
| Tpd52l2 | 66314 | 90,182916 | 7,578359875 | 2,140807893 | 0,0004 | 0,004895 |
| Hk1 | 15275 | 33,282345 | -6,527053818 | 1,845035341 | 0,000404 | 0,004914 |
| Dnajc16 | 214063 | 159,24221 | -7,004218024 | 1,979546208 | 0,000403 | 0,004914 |
| Evi2a | 14017 | 101,97949 | 7,907101761 | 2,234955821 | 0,000403 | 0,004914 |
| Arxes1 | 76219 | 339,55413 | -7,740083929 | 2,187676841 | 0,000403 | 0,004914 |
| Cox6b2 | 333182 | 50,833562 | 8,425424524 | 2,382589665 | 0,000406 | 0,004934 |
| Slc7a10 | 53896 | 25461,384 | -5,199727678 | 1,471435846 | 0,00041 | 0,004975 |
| Smim13 | 108934 | 50,957833 | -7,193033901 | 2,036012947 | 0,000411 | 0,004985 |
| Rspry1 | 67610 | 184,71635 | 8,34221449 | 2,362443383 | 0,000414 | 0,005011 |

| | | | | | | |
|---|---|---|---|---|---|---|
| Actr3 | 74117 | 147,60704 | 7,457085198 | 2,112478779 | 0,000416 | 0,005021 |
| Rgcc | 66214 | 474,05151 | 6,869974063 | 1,946036583 | 0,000415 | 0,005021 |
| 1700030L22Rik | 75576 | 170,64822 | -6,443020693 | 1,825589251 | 0,000417 | 0,005029 |
| Zfp703 | 353310 | 11271,293 | -8,221988387 | 2,330279578 | 0,000418 | 0,005041 |
| Gpr155 | 68526 | 179,00905 | 5,706401983 | 1,617767543 | 0,00042 | 0,005054 |
| Zmym5 | 219105 | 331,18099 | 5,820892116 | 1,650512003 | 0,000421 | 0,00506 |
| Topors | 106021 | 841,43449 | -7,617703651 | 2,161616762 | 0,000425 | 0,005104 |
| Slc7a6 | 330836 | 127,25531 | 7,404702355 | 2,101467854 | 0,000426 | 0,005107 |
| C230072F16Rik | 320784 | 104,15812 | 8,678569735 | 2,463431898 | 0,000427 | 0,005113 |
| Abhd10 | 213012 | 98,47277 | 6,126736611 | 1,739694847 | 0,000429 | 0,005131 |
| Rdx | 19684 | 366,06449 | 5,826451623 | 1,655633213 | 0,000433 | 0,005175 |
| Tsfm | 66399 | 37,255742 | 7,209937537 | 2,050865394 | 0,000439 | 0,005223 |
| Zfp287 | 170740 | 48,471057 | -6,488867904 | 1,845788379 | 0,000439 | 0,005223 |
| Cox4i1 | 12857 | 12123,436 | -6,887484978 | 1,959234207 | 0,000439 | 0,005223 |
| Elmod2 | 244548 | 83,453958 | 7,808442708 | 2,221162805 | 0,000439 | 0,005223 |
| Pcdhga11 | 93723 | 18,658604 | 15,33924164 | 4,364950902 | 0,000441 | 0,005241 |
| Tceal9 | 22381 | 154,76965 | 7,885205753 | 2,244052487 | 0,000442 | 0,005242 |
| Cox8a | 12868 | 7770,436 | -6,799924229 | 1,93597814 | 0,000444 | 0,005264 |
| Lrrc18 | 67580 | 29,799678 | -6,666353844 | 1,898596522 | 0,000446 | 0,005281 |
| Btbd8 | 1,01E+08 | 169,53943 | 6,71582542 | 1,913121747 | 0,000447 | 0,005291 |
| Pcyt1a | 13026 | 40,152525 | 9,076816765 | 2,585960383 | 0,000448 | 0,005292 |
| Pisd-ps1 | 236604 | 616,49418 | -6,153627099 | 1,75430459 | 0,000452 | 0,005332 |
| Phax | 56698 | 149,86855 | 7,521215832 | 2,145151164 | 0,000455 | 0,005351 |
| Cant1 | 76025 | 175,45076 | 7,792441054 | 2,222464667 | 0,000455 | 0,005351 |
| Nudt4 | 71207 | 176,80664 | 7,162204227 | 2,043718338 | 0,000457 | 0,005378 |
| Clu | 12759 | 70527,983 | -5,259169942 | 1,501321685 | 0,00046 | 0,005401 |
| Slc25a27 | 74011 | 155,1039 | 6,628989031 | 1,8932086 | 0,000463 | 0,005427 |
| Dnajb14 | 70604 | 318,63777 | 7,968016781 | 2,281487391 | 0,000479 | 0,005606 |
| Eif4g3 | 230861 | 496,8148 | 5,053679888 | 1,447215651 | 0,000479 | 0,005609 |
| Cox10 | 70383 | 1286,5903 | -7,621425141 | 2,183557702 | 0,000482 | 0,005637 |
| A930005G22Rik | 1,03E+08 | 44,67829 | 8,923089543 | 2,558763947 | 0,000488 | 0,005696 |
| Tmco3 | 234076 | 110,17783 | 6,616998165 | 1,899429877 | 0,000495 | 0,005766 |
| B4galt6 | 56386 | 98,393854 | 7,727509608 | 2,218697397 | 0,000496 | 0,005776 |

| | | | | | | |
|---|---|---|---|---|---|---|
| Strn4 | 97387 | 129,55899 | 6,066455973 | 1,742301727 | 0,000498 | 0,005792 |
| Ccnt2 | 72949 | 131,9681 | 7,439537589 | 2,138830361 | 0,000505 | 0,005852 |
| Calm2 | 12314 | 22183,09 | -6,604426418 | 1,898995562 | 0,000505 | 0,005852 |
| Psmb3 | 26446 | 206,52242 | 5,71066363 | 1,641917852 | 0,000505 | 0,005852 |
| Dhx57 | 106794 | 84,845385 | -7,347848908 | 2,112748763 | 0,000505 | 0,005852 |
| Snx3 | 54198 | 8428,6485 | -6,60814842 | 1,900394037 | 0,000507 | 0,005858 |
| Gtf2i | 14886 | 24536,135 | -7,533392236 | 2,166801202 | 0,000508 | 0,005863 |
| Ccdc24 | 381546 | 74,895908 | 8,431767053 | 2,426060759 | 0,00051 | 0,005876 |
| Gm4258 | 1E+08 | 492,92765 | 6,609077103 | 1,901590327 | 0,00051 | 0,005876 |
| Tmem80 | 71448 | 48,726168 | 8,250471446 | 2,377668457 | 0,00052 | 0,005991 |
| Pin1 | 23988 | 111,96447 | 8,858206376 | 2,554548256 | 0,000525 | 0,006037 |
| Mrpl37 | 56280 | 137,66543 | 8,313885274 | 2,39895126 | 0,000529 | 0,006075 |
| Mrgbp | 73247 | 107,62938 | -8,026984577 | 2,317587433 | 0,000533 | 0,006116 |
| Cog1 | 16834 | 35,280427 | 8,886313578 | 2,566054944 | 0,000534 | 0,00612 |
| Arsa | 11883 | 33,361488 | -8,342063588 | 2,411451611 | 0,000541 | 0,006197 |
| Abcb8 | 74610 | 65,316798 | 6,898160288 | 1,994558098 | 0,000543 | 0,006209 |
| Luzp1 | 269593 | 274,84539 | 7,386805455 | 2,136079716 | 0,000544 | 0,006211 |
| Bsn | 12217 | 337,68356 | -7,250414897 | 2,097617366 | 0,000547 | 0,006241 |
| Pax6 | 18508 | 112,76661 | 7,75939241 | 2,245347754 | 0,000549 | 0,006251 |
| Tmem135 | 72759 | 283,20349 | 6,332096829 | 1,832568576 | 0,00055 | 0,006254 |
| Ercc6l2 | 76251 | 166,86214 | -7,535677209 | 2,18141542 | 0,000551 | 0,006266 |
| Vps13b | 666173 | 76,079337 | -6,399742205 | 1,853128797 | 0,000553 | 0,006281 |
| Scaper | 244891 | 334,83751 | 8,532349079 | 2,470807073 | 0,000554 | 0,006281 |
| Zfp386 | 56220 | 87,55923 | 7,744719938 | 2,244630343 | 0,00056 | 0,006342 |
| Zfp410 | 52708 | 72,929002 | 7,128850099 | 2,067793967 | 0,000566 | 0,0064 |
| Gm40451 | 1,05E+08 | 257,04199 | -7,925506405 | 2,299459206 | 0,000568 | 0,006414 |
| Usp20 | 74270 | 75,876943 | -8,143845163 | 2,364025105 | 0,000571 | 0,006449 |
| Col9a3 | 12841 | 130,85514 | 8,468562157 | 2,459424567 | 0,000575 | 0,006479 |
| Ptn | 19242 | 54806,424 | -5,08805583 | 1,477830692 | 0,000575 | 0,006481 |
| Six5 | 20475 | 70,408471 | -8,099538557 | 2,352899358 | 0,000577 | 0,006487 |
| Calu | 12321 | 100,32762 | 6,490381593 | 1,885960782 | 0,000579 | 0,006503 |
| Car11 | 12348 | 118,2035 | -7,938875943 | 2,307406594 | 0,00058 | 0,006515 |
| S1pr5 | 94226 | 70,074556 | 7,759960401 | 2,256391335 | 0,000584 | 0,006529 |

| | | | | | | |
|---|---|---|---|---|---|---|
| Ramac | 67148 | 125,3085 | -7,55668921 | 2,197136282 | 0,000583 | 0,006529 |
| Zfp704 | 170753 | 54,276208 | 5,808247829 | 1,688802507 | 0,000583 | 0,006529 |
| Plekha3 | 83435 | 43,784413 | -6,822747009 | 1,98444856 | 0,000586 | 0,006546 |
| Uqcrc1 | 22273 | 9569,7915 | -6,63612852 | 1,931201027 | 0,00059 | 0,006583 |
| Vgll4 | 232334 | 136,64139 | 6,695610169 | 1,948834585 | 0,000591 | 0,006587 |
| Slc39a1 | 30791 | 192,12943 | -6,200358 | 1,804793883 | 0,000591 | 0,006587 |
| Carf | 241066 | 205,45553 | -9,116045784 | 2,654543969 | 0,000594 | 0,006613 |
| Fmnl2 | 71409 | 370,01102 | 7,079071618 | 2,061898344 | 0,000596 | 0,006626 |
| Tmem8b | 242409 | 166,31425 | 6,569634703 | 1,914787185 | 0,000601 | 0,006669 |
| Huwe1 | 59026 | 369,78062 | 5,729810798 | 1,670044616 | 0,000602 | 0,006669 |
| Tet2 | 214133 | 5379,3509 | -7,195216412 | 2,097618165 | 0,000603 | 0,00668 |
| Ptges2 | 96979 | 26,223605 | -9,161727787 | 2,671424704 | 0,000605 | 0,006683 |
| Nedd9 | 18003 | 89,35941 | 8,851479521 | 2,581028105 | 0,000605 | 0,006683 |
| Osbpl9 | 100273 | 254,7616 | 6,978511844 | 2,035289814 | 0,000606 | 0,006693 |
| Mrpl50 | 28028 | 58,056343 | 7,782394105 | 2,271863212 | 0,000614 | 0,00675 |
| Olfml1 | 244198 | 87,027567 | 8,012490549 | 2,338720292 | 0,000613 | 0,00675 |
| Crip2 | 68337 | 259,14301 | 8,486711493 | 2,47741568 | 0,000613 | 0,00675 |
| Gm12070 | 654472 | 74,180491 | -5,911246431 | 1,726357753 | 0,000617 | 0,006773 |
| S100a1 | 20193 | 11101,578 | -5,839679513 | 1,705514068 | 0,000617 | 0,006773 |
| Strbp | 20744 | 418,18684 | 4,107906785 | 1,200468731 | 0,000622 | 0,006818 |
| Cttn | 13043 | 103,02336 | 6,897342447 | 2,016197935 | 0,000624 | 0,006834 |
| Alg2 | 56737 | 63,130752 | -7,75981841 | 2,271312829 | 0,000634 | 0,006941 |
| Lrrc2 | 74249 | 22,304939 | -8,621648148 | 2,523804649 | 0,000635 | 0,006942 |
| Ctnnbip1 | 67087 | 261,19222 | 6,578585552 | 1,926130623 | 0,000637 | 0,006944 |
| Slc25a5 | 11740 | 10231,723 | -5,38766152 | 1,577376951 | 0,000636 | 0,006944 |
| 2310061I04Rik | 69662 | 39,329039 | 7,978018671 | 2,336240595 | 0,000638 | 0,006947 |
| Rars2 | 109093 | 73,33058 | 8,132355383 | 2,381562884 | 0,000638 | 0,006947 |
| Ttpa | 50500 | 91,782362 | 8,790920831 | 2,57490064 | 0,00064 | 0,006956 |
| Bag1 | 12017 | 89,378138 | 8,417045469 | 2,467072329 | 0,000645 | 0,007008 |
| Ufsp2 | 192169 | 97,469594 | 8,406578426 | 2,465196612 | 0,000649 | 0,007043 |
| Manf | 74840 | 149,88885 | 6,961215955 | 2,041629111 | 0,00065 | 0,007046 |
| Zfp948 | 381066 | 147,20317 | 6,924461963 | 2,031184447 | 0,000652 | 0,007046 |
| Gm39469 | 1,05E+08 | 77,767403 | 8,300466389 | 2,434645899 | 0,000651 | 0,007046 |

| | | | | | | |
|---|---|---|---|---|---|---|
| Chn1 | 108699 | 98,567817 | 7,181749629 | 2,108970596 | 0,000661 | 0,007136 |
| Bola2 | 66162 | 50,406028 | 7,210478244 | 2,118727805 | 0,000666 | 0,007184 |
| Lmna | 16905 | 189,95868 | 8,927749673 | 2,623655948 | 0,000667 | 0,007187 |
| Vwa5a | 67776 | 295,72391 | 7,854756356 | 2,31087289 | 0,000676 | 0,007278 |
| Zmiz2 | 52915 | 101,39899 | 8,182231802 | 2,40764104 | 0,000678 | 0,007282 |
| Gpalpp1 | 67467 | 485,0336 | 9,344147907 | 2,749636397 | 0,000678 | 0,007282 |
| D5Ertd579e | 320661 | 88,402289 | 7,271011135 | 2,139984888 | 0,00068 | 0,007287 |
| Ints7 | 77065 | 30,973852 | 8,458869245 | 2,489714835 | 0,00068 | 0,007287 |
| LOC118567753 | 1,19E+08 | 18,147573 | -7,40495528 | 2,179983751 | 0,000682 | 0,007299 |
| Tex2 | 21763 | 93,271654 | 7,011139924 | 2,064500237 | 0,000684 | 0,007311 |
| 1700047M11Rik | 67330 | 60,90063 | 8,029006555 | 2,365412692 | 0,000688 | 0,007349 |
| Eid1 | 58521 | 453,88033 | 7,621059734 | 2,245824728 | 0,00069 | 0,007357 |
| Grk6 | 26385 | 104,86621 | -8,226514913 | 2,424044181 | 0,00069 | 0,007357 |
| Fbxo7 | 69754 | 102,93601 | 6,568924553 | 1,936056579 | 0,000691 | 0,007363 |
| Mkrn1 | 54484 | 123,49255 | 7,737079146 | 2,280642039 | 0,000693 | 0,007367 |
| Clcn6 | 26372 | 184,1516 | -6,066518824 | 1,788596061 | 0,000694 | 0,007378 |
| Foxk2 | 68837 | 43,465848 | 7,361267222 | 2,170624167 | 0,000696 | 0,007383 |
| Dnajc1 | 13418 | 138,27643 | 6,396518237 | 1,888323608 | 0,000706 | 0,007467 |
| Bnip3l-ps | 1E+08 | 104,48102 | 7,562957236 | 2,232430822 | 0,000705 | 0,007467 |
| Mnt | 17428 | 243,09036 | -6,603240996 | 1,949391457 | 0,000706 | 0,007467 |
| 4930453N24Rik | 67609 | 238,1716 | 8,775428813 | 2,594610182 | 0,000719 | 0,0076 |
| Rad23b | 19359 | 68,262723 | 6,420492752 | 1,898706962 | 0,000721 | 0,007605 |
| Trim41 | 211007 | 155,6372 | 9,144480705 | 2,704333689 | 0,000721 | 0,007605 |
| Slc35a1 | 24060 | 93,480796 | 6,931437334 | 2,050466326 | 0,000724 | 0,007625 |
| D830031N03Rik | 442834 | 71,288592 | 7,846368649 | 2,321531026 | 0,000725 | 0,007633 |
| Rbbp7 | 245688 | 171,60944 | 6,815873328 | 2,017238663 | 0,000728 | 0,007638 |
| Ints6 | 18130 | 183,04564 | 6,778791936 | 2,006274629 | 0,000728 | 0,007638 |
| Cdk5rap2 | 214444 | 320,19244 | 6,303004849 | 1,865372664 | 0,000728 | 0,007638 |
| Dtnb | 13528 | 136,03007 | -7,345594025 | 2,175263743 | 0,000733 | 0,007675 |
| Sepsecs | 211006 | 72,552288 | 8,81098188 | 2,609089654 | 0,000733 | 0,007675 |
| Diablo | 66593 | 58,429356 | 8,246000242 | 2,444493289 | 0,000743 | 0,007767 |
| Ing1 | 26356 | 114,36541 | 8,038388933 | 2,385715191 | 0,000753 | 0,00787 |
| Gm30320 | 1,03E+08 | 40,647839 | -7,482748149 | 2,22105723 | 0,000754 | 0,007873 |

| | | | | | | |
|---|---|---|---|---|---|---|
| Lpar4 | 78134 | 451,3101 | 6,165638982 | 1,83122381 | 0,00076 | 0,007923 |
| Atp5j2 | 57423 | 5755,6226 | -6,360651785 | 1,889973604 | 0,000764 | 0,007957 |
| Twf1 | 19230 | 239,82136 | 5,132571714 | 1,525378476 | 0,000766 | 0,007969 |
| Baz1b | 22385 | 775,61854 | 5,869216972 | 1,745538012 | 0,000773 | 0,008021 |
| Timm23 | 53600 | 359,91608 | 6,14955436 | 1,828870242 | 0,000772 | 0,008021 |
| Cacna1d | 12289 | 64,675689 | 5,805938466 | 1,727507541 | 0,000777 | 0,008057 |
| Spata24 | 71242 | 29,418151 | 7,855711358 | 2,3392512 | 0,000784 | 0,008127 |
| Fam133b | 68152 | 5848,1717 | -7,199969635 | 2,14510807 | 0,000789 | 0,00817 |
| Kidins220 | 77480 | 572,78723 | -4,260888792 | 1,270065443 | 0,000794 | 0,008209 |
| Trim33 | 94093 | 181,95273 | 7,004492916 | 2,088395127 | 0,000797 | 0,008226 |
| Arl4a | 11861 | 171,89869 | 7,159838907 | 2,135840196 | 0,000802 | 0,008255 |
| Acot13 | 66834 | 13445,046 | -6,720603216 | 2,004889031 | 0,000802 | 0,008255 |
| Uqcrb | 67530 | 9116,245 | -6,686030525 | 1,994628248 | 0,000802 | 0,008255 |
| Cep83 | 77048 | 156,33539 | 7,925582894 | 2,364531033 | 0,000803 | 0,008255 |
| LOC118567737 | 1,19E+08 | 21,629284 | -7,552151961 | 2,253679638 | 0,000805 | 0,008272 |
| Actr1b | 226977 | 100,2542 | 8,959030033 | 2,674075737 | 0,000807 | 0,008284 |
| 2700046G09Rik | 67188 | 80,05786 | 9,029617001 | 2,696910016 | 0,000814 | 0,008341 |
| Cdkn2d | 12581 | 34,580679 | 8,207686778 | 2,453390666 | 0,000822 | 0,008414 |
| Glcci1 | 170772 | 65,773368 | 6,635801105 | 1,983840327 | 0,000823 | 0,008421 |
| Gm20773 | 665301 | 23,662459 | -6,680327005 | 1,999125613 | 0,000833 | 0,008504 |
| Gm20851 | 1E+08 | 23,662459 | -6,680327005 | 1,999125613 | 0,000833 | 0,008504 |
| Zfp157 | 72154 | 246,1714 | 7,126611436 | 2,133899059 | 0,000839 | 0,008554 |
| Nbas | 71169 | 90,00499 | 7,482708868 | 2,24106163 | 0,000841 | 0,00857 |
| Selenoi | 28042 | 245,44979 | 7,148304434 | 2,141825332 | 0,000845 | 0,008582 |
| Fndc3a | 319448 | 361,46361 | 6,991402012 | 2,094623033 | 0,000844 | 0,008582 |
| Pcgf6 | 71041 | 60,062951 | -7,269931748 | 2,177812772 | 0,000843 | 0,008582 |
| 9530068E07Rik | 213673 | 154,93274 | -7,998423878 | 2,396604015 | 0,000846 | 0,008582 |
| Gm52525 | 1,15E+08 | 39,220583 | -6,260628801 | 1,876882612 | 0,000851 | 0,008627 |
| Gramd1a | 52857 | 40,500315 | -6,986250099 | 2,095947333 | 0,000858 | 0,008694 |
| Ism1 | 319909 | 100,68605 | -5,51096735 | 1,655384122 | 0,000871 | 0,008814 |
| Tmed7 | 66676 | 596,85187 | -5,933701009 | 1,782759766 | 0,000874 | 0,008828 |
| Dhcr24 | 74754 | 263,43001 | -6,622178035 | 1,990048542 | 0,000876 | 0,008843 |
| Epc2 | 227867 | 47,09905 | 8,156060086 | 2,451856704 | 0,000879 | 0,008871 |

| | | | | | | |
|---|---|---|---|---|---|---|
| Pcnt | 18541 | 110,31626 | 6,924736382 | 2,083479369 | 0,000889 | 0,008953 |
| Zbed6 | 667118 | 49,898384 | 6,546462139 | 1,970336068 | 0,000892 | 0,00898 |
| Pou2f1 | 18986 | 174,78833 | 4,498626835 | 1,354704034 | 0,000898 | 0,009028 |
| Aldh9a1 | 56752 | 179,52698 | 6,101730274 | 1,837930608 | 0,000901 | 0,009046 |
| Lrig1 | 16206 | 150,86039 | 5,791820495 | 1,745237691 | 0,000905 | 0,009078 |
| Atp5md | 66477 | 3629,2299 | -6,324335663 | 1,905979854 | 0,000906 | 0,009084 |
| Gsk3b | 56637 | 501,02945 | 4,793784574 | 1,44602868 | 0,000916 | 0,009174 |
| Enpp5 | 83965 | 103,75932 | 6,424585802 | 1,939201757 | 0,000923 | 0,009235 |
| Mesd | 67943 | 57,999932 | 6,746757764 | 2,036626914 | 0,000924 | 0,009235 |
| Mlf2 | 30853 | 5137,7211 | -6,803364534 | 2,054565527 | 0,000929 | 0,009271 |
| Serinc5 | 218442 | 204,60138 | 6,906553524 | 2,08671363 | 0,000934 | 0,009314 |
| Mpzl1 | 68481 | 75,549805 | 6,788546409 | 2,052426216 | 0,000941 | 0,009378 |
| Gm39326 | 1,05E+08 | 118,94503 | 6,531191793 | 1,975737409 | 0,000947 | 0,009431 |
| Natd1 | 24083 | 134,06623 | 9,180561751 | 2,778026883 | 0,000951 | 0,009455 |
| Sec11c | 66286 | 12494,702 | -7,120289104 | 2,155172272 | 0,000954 | 0,009476 |
| Sec23ip | 207352 | 124,80237 | 6,328207804 | 1,915755828 | 0,000956 | 0,009486 |
| Cep295 | 319675 | 300,71099 | 5,52411867 | 1,672496196 | 0,000957 | 0,009487 |
| Cpeb2 | 231207 | 404,86517 | -7,146005496 | 2,1647804 | 0,000963 | 0,009542 |
| Rgs12 | 71729 | 85,246541 | 6,548785263 | 1,984746036 | 0,000968 | 0,009583 |
| Peli1 | 67245 | 226,90392 | -5,041827686 | 1,529147886 | 0,000977 | 0,009656 |
| Rarres2 | 71660 | 274,55138 | 6,092259391 | 1,848594773 | 0,000982 | 0,009689 |
| Micall1 | 27008 | 59,985655 | 7,71266451 | 2,340242177 | 0,000982 | 0,009689 |
| Psmd13 | 23997 | 93,072129 | 7,360336688 | 2,234799514 | 0,000989 | 0,009752 |
| Fam98a | 72722 | 32,115811 | 7,290451624 | 2,215310057 | 0,000999 | 0,009813 |
| 5730455P16Rik | 70591 | 101,57293 | 7,004928621 | 2,128498384 | 0,000998 | 0,009813 |
| Spcs1 | 69019 | 14253,965 | -6,576656741 | 1,998291784 | 0,000998 | 0,009813 |
| Mrps5 | 77721 | 46,695253 | 6,705994045 | 2,038941997 | 0,001006 | 0,009872 |
| Cct6a | 12466 | 87,719831 | 6,247351681 | 1,900978827 | 0,001015 | 0,009947 |
| Cpe | 12876 | 38698,801 | -4,792632583 | 1,458504324 | 0,001016 | 0,009947 |
| Ints14 | 69882 | 27,469635 | 9,210238384 | 2,802840477 | 0,001016 | 0,009947 |
| Yipf5 | 67180 | 264,90402 | 6,064564773 | 1,846065908 | 0,001019 | 0,009968 |
| Rufy2 | 70432 | 257,75352 | 7,11401378 | 2,165910504 | 0,001021 | 0,009979 |
| LOC105243553 | 1,05E+08 | 200,24949 | 5,250978107 | 1,599830785 | 0,00103 | 0,010042 |

| | | | | | | |
|---|---|---|---|---|---|---|
| Sec23a | 20334 | 66,428171 | 8,363365685 | 2,547912955 | 0,001029 | 0,010042 |
| B4gat1 | 108902 | 265,28186 | 5,442008142 | 1,659372126 | 0,00104 | 0,010117 |
| Orc2 | 18393 | 79,219414 | 7,823268486 | 2,385378606 | 0,001039 | 0,010117 |
| Cthrc1 | 68588 | 194,56014 | 8,59866756 | 2,622706557 | 0,001043 | 0,010144 |
| Nrd1 | 230598 | 303,35699 | 6,590864753 | 2,010809525 | 0,001047 | 0,010164 |
| Ndufc2 | 68197 | 10982,69 | -6,706163944 | 2,046703438 | 0,001051 | 0,010195 |
| Mrpl17 | 27397 | 84,941767 | 6,490455109 | 1,981402503 | 0,001054 | 0,010207 |
| Atp6v1c1 | 66335 | 188,18896 | 7,943490294 | 2,424819645 | 0,001053 | 0,010207 |
| Pdcd7 | 50996 | 144,09184 | 8,026689011 | 2,451849468 | 0,001061 | 0,010268 |
| Nkx6-2 | 14912 | 48,034445 | 7,810457615 | 2,386300308 | 0,001064 | 0,010279 |
| Cnot6l | 231464 | 98,456674 | 8,070809964 | 2,465968654 | 0,001065 | 0,010279 |
| Tle4 | 21888 | 583,60266 | -6,021299906 | 1,840318673 | 0,001068 | 0,010305 |
| Tm2d1 | 94043 | 176,66379 | 5,825821762 | 1,783603664 | 0,00109 | 0,010489 |
| Tmem185a | 236848 | 171,07971 | 8,599010446 | 2,632588973 | 0,001089 | 0,010489 |
| Adra2a | 11551 | 75,697873 | 6,369181999 | 1,950366358 | 0,001092 | 0,010505 |
| Prkar2a | 19087 | 59,044615 | 6,234577589 | 1,910418882 | 0,001101 | 0,010545 |
| Gm45913 | 545785 | 23,897992 | -6,917483623 | 2,119561248 | 0,0011 | 0,010545 |
| Stam2 | 56324 | 64,40845 | -7,24623075 | 2,220321738 | 0,0011 | 0,010545 |
| Catspere2 | 545391 | 79,18968 | 7,147917008 | 2,190265512 | 0,0011 | 0,010545 |
| Rtn1 | 104001 | 90,399344 | 7,815659023 | 2,395558185 | 0,001104 | 0,010568 |
| Tacc1 | 320165 | 122,26869 | 6,627518124 | 2,03166529 | 0,001106 | 0,010575 |
| Mapre3 | 100732 | 188,4623 | 5,464059808 | 1,678868942 | 0,001135 | 0,010848 |
| Tspan6 | 56496 | 109,52469 | 7,3140818 | 2,248655411 | 0,001143 | 0,010886 |
| Mllt3 | 70122 | 129,80856 | 6,771790338 | 2,082042313 | 0,001144 | 0,010886 |
| Aspscr1 | 68938 | 80,614193 | 5,972998173 | 1,836660435 | 0,001146 | 0,010886 |
| Rnf138 | 56515 | 373,9432 | -5,948925133 | 1,829221405 | 0,001145 | 0,010886 |
| Pak4 | 70584 | 52,789034 | 6,905177131 | 2,122567152 | 0,001141 | 0,010886 |
| Tmod3 | 50875 | 62,466314 | 8,28842418 | 2,548733837 | 0,001146 | 0,010886 |
| Ntsr2 | 18217 | 12948,387 | -5,922982304 | 1,821944243 | 0,00115 | 0,010916 |
| 3110053B16Rik | 382686 | 58,366341 | -7,21260388 | 2,220844989 | 0,001163 | 0,01103 |
| Tcf3 | 21423 | 80,224322 | 7,126266638 | 2,195504714 | 0,001171 | 0,011091 |
| B9d1 | 27078 | 111,94136 | 6,370641387 | 1,963159185 | 0,001174 | 0,01111 |
| Mcee | 73724 | 6063,9236 | -7,406312516 | 2,282714401 | 0,001176 | 0,011122 |

| | | | | | | |
|---|---|---|---|---|---|---|
| Pkn2 | 109333 | 151,51292 | 6,402912746 | 1,974220262 | 0,001182 | 0,01116 |
| Sfxn3 | 94280 | 75,777793 | 7,605017949 | 2,346232778 | 0,00119 | 0,011224 |
| Nav2 | 78286 | 421,19559 | 5,001827692 | 1,544422257 | 0,001201 | 0,011321 |
| Shoc2 | 56392 | 66,803688 | 6,370096561 | 1,969876087 | 0,001222 | 0,011495 |
| Ssx2ip | 99167 | 56,763317 | 7,692871835 | 2,378839203 | 0,001221 | 0,011495 |
| Lmo4 | 16911 | 352,91024 | -6,721158985 | 2,079669216 | 0,00123 | 0,011561 |
| Syap1 | 67043 | 45,610523 | -7,651736743 | 2,368770045 | 0,001237 | 0,011604 |
| Ddx3y | 26900 | 678,30121 | 4,964131678 | 1,536744084 | 0,001237 | 0,011604 |
| Cabcoco1 | 73287 | 144,36432 | 8,693807144 | 2,692251251 | 0,001241 | 0,011636 |
| Gm9234 | 668548 | 90,000239 | -5,664888064 | 1,754942491 | 0,001247 | 0,011675 |
| Scamp2 | 24044 | 205,38274 | 7,216123062 | 2,235944688 | 0,00125 | 0,01169 |
| Mfsd8 | 72175 | 99,748978 | -8,092031304 | 2,50791898 | 0,001253 | 0,011709 |
| Ccdc171 | 320226 | 232,72007 | 7,958269466 | 2,46674301 | 0,001254 | 0,011713 |
| 4931408D14Rik | 77059 | 64,422399 | -7,37997844 | 2,288839478 | 0,001263 | 0,01178 |
| Mbd6 | 110962 | 33,251647 | 6,856634213 | 2,126916051 | 0,001265 | 0,011793 |
| Spef2 | 320277 | 361,6154 | 7,093960707 | 2,200903626 | 0,001268 | 0,011804 |
| Zcchc4 | 78796 | 1762,0804 | -6,374039084 | 1,978107659 | 0,001272 | 0,011812 |
| Prr5l | 72446 | 114,21502 | 6,936931582 | 2,152842576 | 0,001272 | 0,011812 |
| Gramd1c | 207798 | 182,29911 | 7,793273383 | 2,418530963 | 0,001272 | 0,011812 |
| Adgrl1 | 330814 | 3457,6425 | -4,985564131 | 1,549131368 | 0,00129 | 0,011963 |
| Rer1 | 67830 | 198,89996 | 4,922303068 | 1,529622633 | 0,001291 | 0,011965 |
| Egfr | 13649 | 334,56315 | 5,673007389 | 1,763822316 | 0,001298 | 0,012024 |
| Ppp1r8 | 100336 | 65,748321 | 6,664232864 | 2,07413303 | 0,001313 | 0,012151 |
| Dnmt3a | 13435 | 356,59302 | 4,852790584 | 1,511053725 | 0,00132 | 0,012203 |
| Sumo3 | 20610 | 116,53654 | 7,551334934 | 2,352799286 | 0,00133 | 0,012278 |
| Atad2b | 320817 | 3429,3455 | -6,144176645 | 1,915763882 | 0,00134 | 0,012367 |
| Cspg5 | 29873 | 6385,4998 | -4,616196203 | 1,439685857 | 0,001344 | 0,012377 |
| Pxdn | 69675 | 55,953558 | 7,124237443 | 2,221722071 | 0,001343 | 0,012377 |
| Bzw1 | 66882 | 148,17691 | 5,422439481 | 1,692857952 | 0,001359 | 0,012507 |
| Atp13a1 | 170759 | 49,533886 | -6,788728319 | 2,122508497 | 0,001382 | 0,012692 |
| Immp1l | 66541 | 60,686524 | 7,700012197 | 2,407483738 | 0,001382 | 0,012692 |
| Tut4 | 230594 | 148,58469 | 5,626459548 | 1,759901639 | 0,001389 | 0,012725 |
| Rpgrip1 | 77945 | 62,678557 | 5,756437936 | 1,800829577 | 0,001391 | 0,012725 |

| | | | | | | |
|---|---|---|---|---|---|---|
| Zhx3 | 320799 | 72,488929 | -7,2107888 | 2,255643197 | 0,00139 | 0,012725 |
| Plp2 | 18824 | 23,812844 | -8,944260724 | 2,797655241 | 0,001388 | 0,012725 |
| Gm52702 | 1,15E+08 | 114,93757 | 6,559830825 | 2,053108908 | 0,001398 | 0,012778 |
| Mrs2 | 380836 | 192,27524 | -6,670076328 | 2,088368662 | 0,001404 | 0,012818 |
| Pak1ip1 | 68083 | 65,421998 | 6,55022403 | 2,051047177 | 0,001405 | 0,01282 |
| Map7d2 | 78283 | 91,631568 | 7,610104466 | 2,383992116 | 0,001412 | 0,012872 |
| Trmt5 | 76357 | 29,755976 | 7,863639445 | 2,463727089 | 0,001414 | 0,012878 |
| Mrpl10 | 107732 | 263,34114 | -6,572131264 | 2,059279017 | 0,001415 | 0,012879 |
| Fam120a | 218236 | 628,67314 | 5,834106493 | 1,830197696 | 0,001434 | 0,013037 |
| ND1 | 17716 | 40653,895 | -4,36164875 | 1,368418229 | 0,001436 | 0,01304 |
| Lmf2 | 105847 | 118,74449 | 6,377499791 | 2,002727158 | 0,001451 | 0,013155 |
| Proser1 | 212127 | 45,671208 | 7,309029705 | 2,295346003 | 0,001451 | 0,013155 |
| Myg1 | 60315 | 126,6345 | 7,110993208 | 2,233558668 | 0,001454 | 0,01317 |
| Hsdl1 | 72552 | 44,37099 | 7,593221156 | 2,38530752 | 0,001456 | 0,013175 |
| Bcr | 110279 | 68,66021 | 6,595752323 | 2,072441429 | 0,00146 | 0,013184 |
| Ak7 | 78801 | 56,425395 | 7,359966623 | 2,312490471 | 0,001459 | 0,013184 |
| Gm2423 | 1E+08 | 33,862015 | 7,544359657 | 2,371175147 | 0,001464 | 0,013213 |
| Slco3a1 | 108116 | 147,43703 | 7,6115828 | 2,393506459 | 0,001472 | 0,013274 |
| Bmpr2 | 12168 | 223,3036 | 6,076408303 | 1,911703753 | 0,00148 | 0,013323 |
| Coq8b | 76889 | 50,979988 | 7,984748848 | 2,512100868 | 0,00148 | 0,013323 |
| Letm1 | 56384 | 95,298821 | 5,775597088 | 1,817957506 | 0,001488 | 0,01337 |
| Dnajb4 | 67035 | 57,193428 | 6,86949441 | 2,162165077 | 0,001487 | 0,01337 |
| Cep290 | 216274 | 510,07141 | 5,872674677 | 1,8500431 | 0,001502 | 0,013479 |
| Creb3 | 12913 | 133,69928 | 6,779649954 | 2,137521165 | 0,001515 | 0,013574 |
| Raf1 | 110157 | 100,665 | 7,735720211 | 2,438922545 | 0,001515 | 0,013574 |
| 2610524H06Rik | 330173 | 37,784317 | 8,344521113 | 2,63107713 | 0,001516 | 0,013574 |
| Zfp709 | 236193 | 35,864781 | 6,087836236 | 1,920614105 | 0,001526 | 0,013639 |
| Magi1 | 14924 | 199,97217 | 4,803002361 | 1,515328845 | 0,001526 | 0,013639 |
| Tm7sf3 | 67623 | 117,66377 | 4,848670929 | 1,529984461 | 0,001529 | 0,013651 |
| Ift57 | 73916 | 86,481698 | 6,416039238 | 2,024876631 | 0,001532 | 0,013651 |
| Tmx4 | 52837 | 117,33843 | 5,612822806 | 1,771404844 | 0,001532 | 0,013651 |
| Mcl1 | 17210 | 113,74102 | 6,351753873 | 2,004829334 | 0,001534 | 0,013655 |
| Zfp687 | 78266 | 55,005393 | 5,70287122 | 1,800275673 | 0,001536 | 0,013664 |

| | | | | | | |
|---|---|---|---|---|---|---|
| Capn2 | 12334 | 70,582811 | -7,242138984 | 2,287701042 | 0,001547 | 0,01375 |
| Golga2 | 99412 | 135,7592 | -5,39199345 | 1,703454327 | 0,001549 | 0,013755 |
| Mief1 | 239555 | 67,87303 | 6,383315239 | 2,016922487 | 0,001551 | 0,013764 |
| St6galnac3 | 20447 | 142,26548 | 6,447638231 | 2,03758945 | 0,001554 | 0,013777 |
| Cops3 | 26572 | 80,332696 | -5,354956029 | 1,692448366 | 0,001556 | 0,01378 |
| Tmem50b | 77975 | 151,12361 | -5,676997753 | 1,794378184 | 0,001557 | 0,01378 |
| Eif1ax | 66235 | 90,642258 | 6,517290455 | 2,06202117 | 0,001574 | 0,013917 |
| Pbx2 | 18515 | 53,784958 | 7,464648861 | 2,363155668 | 0,001584 | 0,013991 |
| Zbed5 | 71970 | 78,213928 | -7,324840656 | 2,319049712 | 0,001586 | 0,013991 |
| Nr4a2 | 18227 | 204,98 | 8,38327491 | 2,654754515 | 0,001589 | 0,014013 |
| Atpaf2 | 246782 | 63,033198 | 8,320533037 | 2,637652968 | 0,001608 | 0,014161 |
| Ammecr1l | 225339 | 138,07494 | 6,538629444 | 2,072960556 | 0,001609 | 0,014162 |
| Haghl | 68977 | 64,68788 | 6,680734335 | 2,118820885 | 0,001616 | 0,014192 |
| Tmem132b | 208151 | 196,88634 | 6,62263953 | 2,100519333 | 0,001617 | 0,014192 |
| Apcdd1 | 494504 | 99,305831 | 7,264435336 | 2,303945237 | 0,001616 | 0,014192 |
| Rpl31 | 114641 | 7093,67 | -5,864297888 | 1,860253903 | 0,001619 | 0,014201 |
| Hip1r | 29816 | 72,33016 | 7,959751962 | 2,525284723 | 0,001621 | 0,014207 |
| Aebp1 | 11568 | 35,867041 | 6,509802207 | 2,065547903 | 0,001624 | 0,014214 |
| Cerk | 223753 | 81,688539 | 7,427103528 | 2,356789739 | 0,001625 | 0,014214 |
| Eml5 | 319670 | 128,18698 | 6,146747092 | 1,951233201 | 0,001632 | 0,014242 |
| Ss18l2 | 26901 | 60,595174 | 6,927323944 | 2,199110306 | 0,001632 | 0,014242 |
| Erbin | 59079 | 704,88155 | 6,426865155 | 2,040424764 | 0,001634 | 0,014242 |
| Gm45855 | 1,08E+08 | 34,61611 | -6,803070142 | 2,159779415 | 0,001633 | 0,014242 |
| Ablim1 | 226251 | 77,455194 | 5,945955178 | 1,888116173 | 0,001637 | 0,01426 |
| C030029H02Rik | 77383 | 63,126877 | 8,295495619 | 2,634763864 | 0,001641 | 0,01428 |
| Map1a | 17754 | 230,34555 | 6,016902834 | 1,911909052 | 0,001649 | 0,014331 |
| Gm10157 | 1,03E+08 | 76,543448 | -5,886308646 | 1,870493837 | 0,00165 | 0,014331 |
| Sec61b | 66212 | 103,93816 | 8,190564656 | 2,604989632 | 0,001666 | 0,014453 |
| Galnt2 | 108148 | 64,128228 | 7,394437052 | 2,352285456 | 0,001669 | 0,014474 |
| Nsun7 | 70918 | 56,341598 | -7,18298532 | 2,285541171 | 0,001673 | 0,014497 |
| CYTB | 17711 | 116931,91 | -4,449823642 | 1,416021749 | 0,001675 | 0,0145 |
| Ccnl1 | 56706 | 315,91212 | 5,002358565 | 1,592460699 | 0,001682 | 0,014534 |
| Cyp4v3 | 102294 | 168,23666 | 7,478993324 | 2,380857092 | 0,001682 | 0,014534 |

| | | | | | | |
|---|---|---|---|---|---|---|
| Ranbp3l | 223332 | 125,28789 | 6,397214276 | 2,037534487 | 0,001691 | 0,014601 |
| Tmem263 | 103266 | 83,033847 | 7,143764429 | 2,275600015 | 0,001694 | 0,014608 |
| Myl6 | 17904 | 7574,4695 | -6,505308133 | 2,073013441 | 0,001701 | 0,014655 |
| Gm3883 | 1E+08 | 114,06223 | -7,337177638 | 2,338635414 | 0,001705 | 0,014678 |
| Abcb10 | 56199 | 55,994783 | -7,271168757 | 2,318133453 | 0,001709 | 0,014702 |
| Caprin1 | 53872 | 191,74265 | 5,214590289 | 1,663105184 | 0,001716 | 0,014742 |
| Mob3b | 214944 | 260,88086 | 8,129365943 | 2,592819796 | 0,001717 | 0,014742 |
| Zfp612 | 234725 | 5203,1206 | -6,817861575 | 2,174936247 | 0,00172 | 0,014759 |
| Msl2 | 77853 | 50,745781 | 7,717079424 | 2,462803578 | 0,001728 | 0,014811 |
| Psme2 | 19188 | 80,236237 | -6,707010142 | 2,14073577 | 0,00173 | 0,01482 |
| Plaa | 18786 | 126,16406 | 7,098435668 | 2,266076281 | 0,001733 | 0,014835 |
| Cnot1 | 234594 | 75,519954 | 6,942710724 | 2,216583826 | 0,001735 | 0,014838 |
| Etfdh | 66841 | 68,650154 | 7,575473088 | 2,419008506 | 0,001738 | 0,014852 |
| Tns3 | 319939 | 85,400521 | 6,373977802 | 2,037830249 | 0,001761 | 0,015024 |
| H2az1 | 51788 | 387,1159 | 4,706713591 | 1,505064756 | 0,001764 | 0,015024 |
| Gabbr1 | 54393 | 1839,0722 | -4,962325676 | 1,586701803 | 0,001763 | 0,015024 |
| Sec22a | 317717 | 224,37337 | 7,971715368 | 2,548829516 | 0,001762 | 0,015024 |
| Rps2 | 16898 | 10204,733 | -6,362555328 | 2,035553465 | 0,001774 | 0,01509 |
| Lig4 | 319583 | 57,375054 | 7,680361119 | 2,457584463 | 0,001777 | 0,015105 |
| Pcbp2 | 18521 | 6359,6565 | -5,048293329 | 1,616018902 | 0,001785 | 0,015157 |
| Gm20716 | 1,06E+08 | 128,23187 | 8,822357829 | 2,824795101 | 0,001789 | 0,015181 |
| Brd7 | 26992 | 4283,3464 | -4,605071973 | 1,474641236 | 0,001791 | 0,015186 |
| Snrpg | 68011 | 5375,7097 | -6,482459958 | 2,077164742 | 0,001803 | 0,015277 |
| Dexi | 58239 | 42,184465 | -6,956922046 | 2,231121607 | 0,00182 | 0,015405 |
| Atg12 | 67526 | 93,562198 | 5,718564585 | 1,834302278 | 0,001823 | 0,015421 |
| Tomm6 | 66119 | 7491,6889 | -6,435814709 | 2,064558368 | 0,001825 | 0,015422 |
| Washc3 | 67282 | 142,10164 | -6,463096553 | 2,07393885 | 0,001831 | 0,015446 |
| Gm52667 | 1,15E+08 | 101,63633 | 6,096798524 | 1,956276 | 0,00183 | 0,015446 |
| Rhof | 23912 | 107,43476 | -6,101077517 | 1,958680841 | 0,00184 | 0,015509 |
| Schip1 | 30953 | 110,95614 | 5,301578712 | 1,703353967 | 0,001856 | 0,015601 |
| Nemf | 66244 | 555,70672 | -5,197025189 | 1,669787057 | 0,001856 | 0,015601 |
| Mrps12 | 24030 | 74,312913 | 7,489321155 | 2,406005619 | 0,001853 | 0,015601 |
| Actr6 | 67019 | 187,52541 | 7,890443403 | 2,535435524 | 0,001858 | 0,015605 |

| | | | | | | |
|---|---|---|---|---|---|---|
| Mpdu1 | 24070 | 140,25268 | 6,964554873 | 2,238365833 | 0,001862 | 0,015624 |
| Gm53055 | 1,18E+08 | 122,0444 | -7,515766259 | 2,415971825 | 0,001865 | 0,01563 |
| Pkig | 18769 | 73,234389 | 7,31173232 | 2,350402064 | 0,001866 | 0,01563 |
| Polr2j | 20022 | 53,323345 | -7,074733496 | 2,275019863 | 0,001872 | 0,015651 |
| 2210408I21Rik | 72371 | 213,69347 | 6,530827061 | 2,100141259 | 0,001873 | 0,015651 |
| Zfp871 | 208292 | 346,07861 | 6,332384811 | 2,036265747 | 0,001872 | 0,015651 |
| Anapc13 | 69010 | 6357,4421 | -6,02890919 | 1,939294805 | 0,001878 | 0,015672 |
| Hacl1 | 56794 | 98,881601 | 7,526179793 | 2,42077716 | 0,001877 | 0,015672 |
| Cables1 | 63955 | 505,59031 | -6,382113388 | 2,053095046 | 0,00188 | 0,015673 |
| Peg13 | 353342 | 36,335046 | 7,029490052 | 2,264203249 | 0,001905 | 0,015869 |
| Txndc9 | 98258 | 165,76278 | 6,652248213 | 2,144500069 | 0,001922 | 0,015996 |
| Prpf38a | 230596 | 202,1201 | 7,067977738 | 2,279037186 | 0,001927 | 0,016021 |
| 1700016K19Rik | 74230 | 120,80922 | 8,274677399 | 2,671150548 | 0,00195 | 0,016198 |
| Rplp1 | 56040 | 7285,557 | -5,642636191 | 1,821842003 | 0,001953 | 0,016217 |
| Fam102a | 98952 | 74,278135 | 6,511894 | 2,102966097 | 0,001958 | 0,016227 |
| Anks3 | 72615 | 121,57842 | 7,437050992 | 2,401611999 | 0,001957 | 0,016227 |
| Prnp | 19122 | 15390,466 | -4,204677705 | 1,358027757 | 0,00196 | 0,016233 |
| Slc38a6 | 625098 | 51,380525 | 7,655715478 | 2,475073049 | 0,001981 | 0,016387 |
| Car2 | 12349 | 26437,337 | -4,992653513 | 1,615014874 | 0,001992 | 0,016469 |
| Adcy5 | 224129 | 26,21166 | 7,152702011 | 2,31485956 | 0,002002 | 0,016538 |
| Itgb1bp1 | 16413 | 91,983781 | 5,907085621 | 1,913150923 | 0,002018 | 0,016652 |
| LOC118568744 | 1,19E+08 | 202,27596 | -6,051230483 | 1,960040694 | 0,00202 | 0,016656 |
| Vezt | 215008 | 610,86548 | -6,254161145 | 2,026461553 | 0,002027 | 0,016679 |
| Ttll5 | 320244 | 122,52536 | 5,54784684 | 1,797655565 | 0,002028 | 0,016679 |
| Htatsf1 | 72459 | 126,76223 | 6,770158285 | 2,193377087 | 0,002024 | 0,016679 |
| Ilf3 | 16201 | 83,766396 | 5,961511266 | 1,932083798 | 0,002032 | 0,0167 |
| Pyurf | 66459 | 76,486793 | 6,341523307 | 2,056794933 | 0,002048 | 0,016803 |
| Gatd3a | 28295 | 42,272774 | 7,060419448 | 2,289853649 | 0,002047 | 0,016803 |
| Psme3ip1 | 102122 | 291,49426 | 6,615314837 | 2,146268004 | 0,002054 | 0,016844 |
| Pfas | 237823 | 74,823062 | 6,364627703 | 2,066512792 | 0,002071 | 0,016964 |
| Ildr2 | 1E+08 | 217,56044 | -5,969084044 | 1,938313774 | 0,002073 | 0,01697 |
| Cldn11 | 18417 | 693,09383 | 5,868756885 | 1,905948121 | 0,002076 | 0,016972 |
| Smc1a | 24061 | 153,49456 | 7,438014992 | 2,415719576 | 0,002077 | 0,016972 |

| | | | | | | |
|---|---|---|---|---|---|---|
| Rnd2 | 11858 | 40,541936 | 6,459076919 | 2,098501115 | 0,002084 | 0,017005 |
| Dguok | 27369 | 55,350617 | 6,751909388 | 2,193522537 | 0,002083 | 0,017005 |
| Sppl3 | 74585 | 73,366717 | 7,115440115 | 2,312113514 | 0,002088 | 0,017018 |
| D330041H03Rik | 654822 | 53,437754 | 7,723577497 | 2,511492573 | 0,002103 | 0,017128 |
| Mt2 | 17750 | 6272,8703 | -4,56786089 | 1,48604796 | 0,002113 | 0,017199 |
| Spr | 20751 | 12,966855 | 8,484539419 | 2,760996525 | 0,002119 | 0,017233 |
| Abca3 | 27410 | 19,541939 | 6,261034599 | 2,039021585 | 0,002136 | 0,017357 |
| Otud7b | 229603 | 324,99235 | 5,624015613 | 1,832164114 | 0,002143 | 0,017401 |
| Arnt2 | 11864 | 205,09541 | -6,395910431 | 2,084877428 | 0,002157 | 0,017494 |
| Marcks | 17118 | 556,99019 | 4,972304785 | 1,621159818 | 0,002161 | 0,017517 |
| Rbbp4 | 19646 | 5558,881 | -5,792850722 | 1,889455011 | 0,00217 | 0,017576 |
| Vps35l | 71517 | 91,382306 | 6,657990829 | 2,172465076 | 0,002179 | 0,01763 |
| Aggf1 | 66549 | 114,01226 | 6,689481325 | 2,182949406 | 0,002181 | 0,017633 |
| Ddx47 | 67755 | 38,421056 | -8,206119971 | 2,679383567 | 0,002194 | 0,017721 |
| Ccnt1 | 12455 | 520,50662 | 6,202708194 | 2,025461728 | 0,002196 | 0,017726 |
| Dsc3 | 13507 | 150,91444 | -6,217568105 | 2,031334173 | 0,002207 | 0,017803 |
| Hmg20b | 15353 | 111,03996 | 6,599673522 | 2,157233474 | 0,002218 | 0,017878 |
| Wdr12 | 57750 | 42,829276 | 7,944645707 | 2,598718014 | 0,002235 | 0,017994 |
| Setd1b | 208043 | 86,885531 | 6,331307932 | 2,071378414 | 0,002239 | 0,018014 |
| Ergic1 | 67458 | 103,68667 | 6,40486634 | 2,095961567 | 0,002245 | 0,018045 |
| Csk | 12988 | 37,282219 | -5,948629397 | 1,946853542 | 0,002247 | 0,018049 |
| Zfp445 | 235682 | 201,90824 | 4,947021149 | 1,620279195 | 0,002264 | 0,018151 |
| Lclat1 | 225010 | 27,250559 | 7,972231886 | 2,611201645 | 0,002265 | 0,018151 |
| Slc66a2 | 66943 | 64,830296 | 8,350298558 | 2,734839081 | 0,002263 | 0,018151 |
| C1galt1c1 | 59048 | 162,1548 | 6,440725953 | 2,110325997 | 0,002273 | 0,018202 |
| Kdm4b | 193796 | 110,11174 | 5,674782461 | 1,859669074 | 0,002277 | 0,018218 |
| Arhgap1 | 228359 | 69,746693 | 7,627108742 | 2,500338653 | 0,002285 | 0,018268 |
| Marchf8 | 71779 | 159,05934 | 5,320563388 | 1,744488259 | 0,002289 | 0,018284 |
| Vmn2r12 | 627569 | 51,05364 | -6,977254315 | 2,288062442 | 0,002293 | 0,018301 |
| Mt1 | 17748 | 14690,467 | -4,396947927 | 1,442125592 | 0,002297 | 0,018315 |
| Dynll2 | 68097 | 177,02879 | 5,272973291 | 1,730540304 | 0,002311 | 0,018403 |
| Opa1 | 74143 | 105,40581 | 6,536376377 | 2,145079836 | 0,00231 | 0,018403 |
| Aopep | 72061 | 98,221146 | 7,037278107 | 2,309844304 | 0,002314 | 0,018411 |

| | | | | | | |
|---|---|---|---|---|---|---|
| Snhg1 | 83673 | 27,070533 | -6,692555514 | 2,198171622 | 0,00233 | 0,018522 |
| Nsun5 | 100609 | 57,377806 | -6,225772979 | 2,045541454 | 0,002338 | 0,018522 |
| Ptdss2 | 27388 | 142,14801 | -5,544073022 | 1,821550473 | 0,002338 | 0,018522 |
| Cxcl14 | 57266 | 4524,409 | -5,097384378 | 1,674386822 | 0,002332 | 0,018522 |
| Cfl2 | 12632 | 82,38445 | 5,535531675 | 1,818868038 | 0,002339 | 0,018522 |
| Depdc5 | 277854 | 137,49692 | 7,349043578 | 2,414269051 | 0,002335 | 0,018522 |
| Zranb1 | 360216 | 107,74083 | 5,782309879 | 1,900301952 | 0,002344 | 0,018542 |
| Cstf2t | 83410 | 86,623571 | 7,119517225 | 2,341271789 | 0,002359 | 0,018648 |
| Epb41l1 | 13821 | 43,532007 | 5,668297512 | 1,864190431 | 0,002361 | 0,018649 |
| Phactr2 | 215789 | 99,446118 | 5,170852221 | 1,701319272 | 0,002371 | 0,018711 |
| Slc1a3 | 20512 | 39827,958 | -4,342624804 | 1,428987194 | 0,002374 | 0,018711 |
| Abhd16a | 193742 | 46,604035 | -7,282963171 | 2,396578665 | 0,002374 | 0,018711 |
| Ndufaf3 | 66706 | 62,851944 | 6,969245867 | 2,293858931 | 0,00238 | 0,018739 |
| Ptprj | 19271 | 51,980155 | 7,52247624 | 2,478580727 | 0,002405 | 0,018925 |
| 4933434E20Rik | 99650 | 123,10167 | 5,213009292 | 1,718223116 | 0,002414 | 0,018976 |
| Togaram1 | 328108 | 94,803217 | 5,865601901 | 1,934370473 | 0,002427 | 0,019037 |
| Btg1 | 12226 | 346,25768 | 4,40575584 | 1,452788253 | 0,002424 | 0,019037 |
| Pds5b | 100710 | 143,53501 | 6,101284125 | 2,012115207 | 0,002427 | 0,019037 |
| Lonp2 | 66887 | 111,76796 | 6,386149307 | 2,107243147 | 0,002441 | 0,019129 |
| Orc6 | 56452 | 82,005987 | 7,598282615 | 2,50754722 | 0,002444 | 0,01914 |
| Pfdn4 | 109054 | 44,792551 | 6,819338249 | 2,250821515 | 0,002448 | 0,019154 |
| Mia | 12587 | 91,628556 | 7,8480939 | 2,590761202 | 0,002452 | 0,019167 |
| Sharpin | 106025 | 48,619638 | 7,337719524 | 2,42318293 | 0,002461 | 0,019224 |
| Mrpl51 | 66493 | 96,392387 | 7,333877044 | 2,423239931 | 0,002474 | 0,019314 |
| Zw10 | 26951 | 17,302846 | 7,889157365 | 2,608483764 | 0,002491 | 0,01943 |
| A930017M01Rik | 239410 | 43,527206 | 7,617160062 | 2,518792437 | 0,002493 | 0,019434 |
| Gtf2h3 | 209357 | 42,626992 | 7,565919724 | 2,502434454 | 0,002499 | 0,019464 |
| Mtmr2 | 77116 | 196,41805 | 5,629322843 | 1,86332914 | 0,002518 | 0,019598 |
| Oma1 | 67013 | 53,475293 | 6,907198516 | 2,286522696 | 0,002521 | 0,019601 |
| Pcdh7 | 54216 | 318,87908 | 4,866602314 | 1,611338044 | 0,002526 | 0,019625 |
| Acot2 | 171210 | 95,811308 | -5,621693212 | 1,861955962 | 0,002534 | 0,019673 |
| Slc8b1 | 170756 | 49,736611 | -6,261970804 | 2,074229692 | 0,002537 | 0,019678 |
| Synm | 233335 | 95,784417 | 6,574377227 | 2,17856685 | 0,002547 | 0,019726 |

| | | | | | | |
|---|---|---|---|---|---|---|
| Rrm1 | 20133 | 147,71385 | 8,991115753 | 2,979443998 | 0,002547 | 0,019726 |
| Stau2 | 29819 | 140,24166 | 7,290933747 | 2,416731574 | 0,002554 | 0,019767 |
| Inf2 | 70435 | 87,555489 | 6,352817889 | 2,106413531 | 0,002562 | 0,019811 |
| Sptlc2 | 20773 | 44,630141 | 6,145690451 | 2,038559288 | 0,002572 | 0,019876 |
| Kmt5b | 225888 | 126,5785 | 5,291235961 | 1,755473081 | 0,002577 | 0,019883 |
| Arf1 | 11840 | 1059,5851 | -6,377471416 | 2,115820813 | 0,002577 | 0,019883 |
| R74862 | 97423 | 77,161935 | 7,588242016 | 2,518354612 | 0,002585 | 0,019931 |
| Tsn | 22099 | 267,39056 | 5,434656887 | 1,80405281 | 0,002591 | 0,019962 |
| Pnpla8 | 67452 | 309,41771 | 5,859233555 | 1,945287518 | 0,002595 | 0,019976 |
| Cox19 | 68033 | 42,975777 | 7,31408013 | 2,428493034 | 0,002597 | 0,019976 |
| Rbm10 | 236732 | 49,743046 | 6,004384607 | 1,993822882 | 0,0026 | 0,01998 |
| Trim28 | 21849 | 110,22097 | 6,872531682 | 2,282664499 | 0,002606 | 0,020013 |
| Wdr70 | 545085 | 34,158987 | 5,97789751 | 1,986306225 | 0,002616 | 0,020051 |
| Flad1 | 319945 | 44,995891 | 7,292322856 | 2,423139791 | 0,002617 | 0,020051 |
| Ppp1r36 | 210762 | 180,82836 | 7,707350689 | 2,560926355 | 0,002616 | 0,020051 |
| Rps19bp1 | 66538 | 51,78328 | 8,036835232 | 2,670718745 | 0,002619 | 0,020051 |
| Glud1 | 14661 | 1336,2729 | -4,178291797 | 1,389281936 | 0,002634 | 0,020129 |
| Hecw2 | 329152 | 101,00497 | 6,584980443 | 2,189465202 | 0,002633 | 0,020129 |
| Sema4d | 20354 | 217,38378 | 6,343650938 | 2,109373344 | 0,002635 | 0,020129 |
| Arhgef25 | 52666 | 50,979932 | 6,675875706 | 2,221918486 | 0,00266 | 0,0203 |
| Lpgat1 | 226856 | 89,727453 | 6,319126757 | 2,103662536 | 0,002666 | 0,020304 |
| Gm13536 | 1,05E+08 | 1434,833 | -5,231609322 | 1,741666033 | 0,002666 | 0,020304 |
| Fzr1 | 56371 | 63,163869 | 8,174780743 | 2,721183122 | 0,002663 | 0,020304 |
| Spata33 | 320869 | 206,12014 | 8,095585471 | 2,695618207 | 0,002671 | 0,020325 |
| Mrps31 | 57312 | 44,196273 | 6,682007369 | 2,22513019 | 0,002674 | 0,020328 |
| Wdr11 | 207425 | 56,980149 | 7,160137292 | 2,384788231 | 0,002678 | 0,020342 |
| E030024N20Rik | 595139 | 220,60818 | -6,216761843 | 2,070673858 | 0,00268 | 0,020342 |
| Ddit4 | 74747 | 24,768871 | 5,665002855 | 1,88741487 | 0,002687 | 0,020367 |
| Ciao3 | 67563 | 78,736806 | 6,977624748 | 2,324708671 | 0,002686 | 0,020367 |
| Ppia | 268373 | 30404,3 | -5,966249502 | 1,988013208 | 0,00269 | 0,020374 |
| Zcchc14 | 142682 | 53,832868 | 6,052418086 | 2,017183574 | 0,002696 | 0,020405 |
| Creb5 | 231991 | 79,951608 | 5,989973712 | 1,998608915 | 0,002726 | 0,020583 |
| Cobll1 | 319876 | 85,998316 | 6,124275966 | 2,043570002 | 0,002728 | 0,020583 |

| | | | | | | |
|---|---|---|---|---|---|---|
| Mog | 17441 | 731,57209 | 6,317248738 | 2,107913469 | 0,002727 | 0,020583 |
| Fdxr | 14149 | 51,298338 | 8,315660615 | 2,774381588 | 0,002724 | 0,020583 |
| Slc25a40 | 319653 | 373,32613 | 6,16214291 | 2,056635673 | 0,002733 | 0,02061 |
| Astn1 | 11899 | 407,69026 | 4,208198076 | 1,404676889 | 0,002737 | 0,02062 |
| Slc35a5 | 74102 | 102,24351 | 5,625446592 | 1,877927244 | 0,002739 | 0,020623 |
| Smpd4 | 77626 | 65,809452 | 7,660950247 | 2,557682648 | 0,002742 | 0,020627 |
| BC005537 | 79555 | 34,534696 | 6,579828998 | 2,197152193 | 0,002747 | 0,02065 |
| Slc27a1 | 26457 | 5012,3364 | -4,272062978 | 1,427009956 | 0,002756 | 0,02067 |
| Gm30985 | 1,03E+08 | 26,134506 | -6,037970135 | 2,016664126 | 0,002753 | 0,02067 |
| Jam3 | 83964 | 154,9617 | 6,645984893 | 2,219881941 | 0,002755 | 0,02067 |
| Pisd | 320951 | 117,65691 | 6,229600982 | 2,081708084 | 0,002767 | 0,020734 |
| Cfap298 | 68001 | 55,078796 | 7,539406826 | 2,520678709 | 0,00278 | 0,020822 |
| Kctd3 | 226823 | 59,085318 | 6,485462669 | 2,168522375 | 0,002783 | 0,020826 |
| Dusp15 | 252864 | 51,917971 | 6,009395961 | 2,009568192 | 0,002786 | 0,020834 |
| Fam172a | 68675 | 32257,164 | -6,055829587 | 2,025390316 | 0,00279 | 0,020848 |
| Csgalnact2 | 78752 | 125,65299 | -6,662597566 | 2,229040217 | 0,002799 | 0,020897 |
| Kcna1 | 16485 | 290,89354 | 5,505587557 | 1,842679082 | 0,00281 | 0,020936 |
| Zfp407 | 240476 | 96,749867 | 4,820007526 | 1,613429586 | 0,002813 | 0,020936 |
| Ccdc148 | 227933 | 92,757708 | 5,410153284 | 1,810662939 | 0,002809 | 0,020936 |
| Lbr | 98386 | 95,232741 | 6,925497925 | 2,318500347 | 0,002817 | 0,020936 |
| Pyroxd1 | 232491 | 99,313102 | 6,972105249 | 2,334048078 | 0,002816 | 0,020936 |
| Skp2 | 27401 | 71,449556 | 8,278125836 | 2,77114648 | 0,002815 | 0,020936 |
| Tceal8 | 66684 | 638,46013 | 8,243129502 | 2,760115647 | 0,002822 | 0,020958 |
| Dclk2 | 70762 | 114,44172 | 6,336033079 | 2,122785697 | 0,002838 | 0,021062 |
| Srek1ip1 | 67288 | 39,680109 | 6,449450423 | 2,161252519 | 0,002844 | 0,021091 |
| Pcnx | 54604 | 138,98948 | 6,085970443 | 2,039708562 | 0,002847 | 0,021101 |
| Hspa14 | 50497 | 63,886065 | 8,518732087 | 2,855784483 | 0,002855 | 0,021139 |
| Ide | 15925 | 167,66565 | 5,44353721 | 1,825072336 | 0,002858 | 0,021146 |
| Rgs4 | 19736 | 158,36058 | 8,039164922 | 2,69691273 | 0,002874 | 0,021252 |
| Exosc3 | 66362 | 64,144415 | -6,294505824 | 2,112910938 | 0,002891 | 0,021362 |
| Gns | 75612 | 193,43158 | 6,502856335 | 2,184038665 | 0,002907 | 0,021439 |
| Syt10 | 54526 | 124,50739 | 7,169231557 | 2,407972178 | 0,002908 | 0,021439 |
| Spats1 | 71020 | 357,06379 | -6,677449374 | 2,242595385 | 0,002906 | 0,021439 |

| | | | | | | |
|---|---|---|---|---|---|---|
| Zfp532 | 328977 | 91,330262 | 6,555491722 | 2,202134863 | 0,002912 | 0,021452 |
| Iars | 105148 | 151,37 | -7,921181335 | 2,662237719 | 0,002926 | 0,02154 |
| Fryl | 72313 | 213,96618 | 5,405269066 | 1,816859502 | 0,002929 | 0,021547 |
| Foxn3 | 71375 | 524,63205 | 4,785755996 | 1,609426407 | 0,002943 | 0,02162 |
| Ndn | 17984 | 60,061842 | 6,82166876 | 2,2939269 | 0,002941 | 0,02162 |
| Ccdc47 | 67163 | 363,71701 | 5,744226939 | 1,932743229 | 0,002958 | 0,021711 |
| Zfp454 | 237758 | 107,45523 | 7,223537728 | 2,431312795 | 0,002968 | 0,021767 |
| Dynlt1c | 1E+08 | 54,855997 | 7,286891847 | 2,452980019 | 0,002972 | 0,021773 |
| Carnmt1 | 67383 | 209,90813 | -6,667314562 | 2,244509272 | 0,002973 | 0,021773 |
| Cul3 | 26554 | 215,65594 | 5,288849676 | 1,780661878 | 0,002976 | 0,021774 |
| Mrpl28 | 68611 | 391,47536 | 6,987786353 | 2,352761628 | 0,002978 | 0,021774 |
| Polr2h | 245841 | 34,070773 | -7,714440809 | 2,598737119 | 0,002992 | 0,021865 |
| Ddx50 | 94213 | 246,80418 | 6,981832071 | 2,352228851 | 0,002996 | 0,021874 |
| Plekha1 | 101476 | 83,464252 | 5,176888195 | 1,744779151 | 0,003006 | 0,021937 |
| Ndufaf2 | 75597 | 51,445815 | 5,916656681 | 1,994668695 | 0,003015 | 0,021964 |
| Zfp608 | 269023 | 326,06079 | -6,072712824 | 2,04720668 | 0,003014 | 0,021964 |
| Fgd5 | 232237 | 16,450699 | -6,675980155 | 2,251101478 | 0,00302 | 0,02199 |
| Arvcf | 11877 | 51,121823 | 6,160169083 | 2,077888206 | 0,00303 | 0,022047 |
| Sox7 | 20680 | 41,666556 | -5,737912258 | 1,93606441 | 0,00304 | 0,022097 |
| Bmyc | 107771 | 124,54089 | 6,894691469 | 2,326992186 | 0,003047 | 0,022137 |
| Adamts20 | 223838 | 71,810735 | 6,87277984 | 2,320345098 | 0,003057 | 0,022173 |
| Fcgrt | 14132 | 92,186652 | 5,172267627 | 1,746355651 | 0,003059 | 0,022173 |
| Jak2 | 16452 | 55,960543 | -5,693971636 | 1,922225171 | 0,003055 | 0,022173 |
| Flii | 14248 | 64,212806 | 6,748104851 | 2,279127578 | 0,003068 | 0,022223 |
| Ftsj3 | 56095 | 516,36704 | -7,988926137 | 2,698714935 | 0,003074 | 0,022247 |
| Zfp950 | 414758 | 450,83511 | 5,044670787 | 1,704339079 | 0,003077 | 0,022258 |
| Tead1 | 21676 | 134,63778 | 5,836857672 | 1,973879712 | 0,003106 | 0,022448 |
| Gtpbp6 | 107999 | 27,37361 | 8,165582399 | 2,762760132 | 0,003121 | 0,022539 |
| Lemd3 | 380664 | 45,918001 | 7,432676021 | 2,515467548 | 0,003129 | 0,022581 |
| Sfmbt2 | 353282 | 29,829213 | 7,028205059 | 2,379223211 | 0,003137 | 0,022623 |
| Rab2b | 76338 | 160,9087 | 6,620649313 | 2,242220789 | 0,00315 | 0,0227 |
| Tasp1 | 75812 | 42,18237 | 8,453098455 | 2,863650184 | 0,003159 | 0,022746 |
| Ifnar2 | 15976 | 135,32013 | -6,782229743 | 2,299451133 | 0,003183 | 0,022871 |

| | | | | | | |
|---|---|---|---|---|---|---|
| Zfp930 | 234358 | 16,006972 | -6,91794495 | 2,34538226 | 0,003182 | 0,022871 |
| Ccz1 | 231874 | 19,670469 | -8,530171346 | 2,891852823 | 0,003181 | 0,022871 |
| Pmvk | 68603 | 48,887449 | 6,434888901 | 2,182131822 | 0,003189 | 0,022899 |
| Ubxn6 | 66530 | 68,787146 | 5,966988916 | 2,023866154 | 0,003195 | 0,022926 |
| Cc2d1b | 319965 | 237,27443 | 6,349221973 | 2,15432269 | 0,003207 | 0,022992 |
| Inpp5a | 212111 | 210,33379 | 6,783894211 | 2,302234154 | 0,003212 | 0,023016 |
| F630111L10Rik | 320463 | 103,78921 | 7,789621096 | 2,644003808 | 0,003218 | 0,023037 |
| Gm52294 | 1,15E+08 | 393,85688 | 5,204179835 | 1,766816263 | 0,003224 | 0,023068 |
| Nadsyn1 | 78914 | 29,802776 | -7,109943969 | 2,414347187 | 0,003231 | 0,023099 |
| Tdrkh | 72634 | 101,97727 | -7,108318703 | 2,414262589 | 0,003237 | 0,023125 |
| Gm40989 | 1,05E+08 | 310,48681 | -4,966082248 | 1,687378719 | 0,00325 | 0,0232 |
| Jpt1 | 15374 | 82,622555 | 6,455114221 | 2,193638764 | 0,003254 | 0,023204 |
| Msrb1 | 27361 | 94,213658 | 8,09334195 | 2,75042191 | 0,003255 | 0,023204 |
| Ehmt1 | 77683 | 233,12297 | 5,656104163 | 1,922675125 | 0,003263 | 0,023247 |
| Dis3l2 | 208718 | 35,955604 | 8,154060031 | 2,773763008 | 0,003285 | 0,023387 |
| Fbxo4 | 106052 | 89,522381 | 7,241585011 | 2,464179645 | 0,003295 | 0,023439 |
| Usp8 | 84092 | 42,111532 | 7,613495914 | 2,590874885 | 0,003297 | 0,023439 |
| Zfp516 | 329003 | 89,913016 | 4,276686211 | 1,455638437 | 0,003303 | 0,023465 |
| Ptp4a3 | 19245 | 37,648066 | 6,074057564 | 2,068889078 | 0,003326 | 0,023558 |
| Upf2 | 326622 | 150,5329 | 5,520096555 | 1,880006602 | 0,003322 | 0,023558 |
| Gm16168 | 1,03E+08 | 95,350993 | 4,706335953 | 1,603010771 | 0,003325 | 0,023558 |
| Tm2d2 | 69742 | 186,95769 | 5,900292319 | 2,009364569 | 0,00332 | 0,023558 |
| Ankra2 | 68558 | 21,776006 | 6,404850701 | 2,18306954 | 0,003348 | 0,023679 |
| Exoc1 | 69940 | 92,617973 | -6,207870912 | 2,115936088 | 0,003348 | 0,023679 |
| Vps35 | 65114 | 140,68716 | 6,343157973 | 2,162741054 | 0,003358 | 0,023734 |
| Akt1 | 11651 | 38,895289 | 6,417597687 | 2,189287922 | 0,003375 | 0,023837 |
| Ssbp1 | 381760 | 213,30419 | 5,883966362 | 2,008482752 | 0,003394 | 0,023959 |
| Fzd4 | 14366 | 66,284397 | -5,794820319 | 1,978501621 | 0,003402 | 0,023993 |
| Pofut2 | 80294 | 27481,796 | -7,279550501 | 2,486415274 | 0,003414 | 0,024066 |
| Casd1 | 213819 | 275,37063 | -6,317382576 | 2,158050992 | 0,003419 | 0,024078 |
| Rpl28 | 19943 | 80,827844 | 4,799043909 | 1,640432067 | 0,003439 | 0,024198 |
| Hif3a | 53417 | 28,636938 | 7,088668096 | 2,423174067 | 0,003441 | 0,024198 |
| Gtf2f1 | 98053 | 44,698297 | 6,103982282 | 2,087461699 | 0,003454 | 0,024278 |

| | | | | | | |
|---|---|---|---|---|---|---|
| Sgms1 | 208449 | 49,411936 | 5,827489066 | 1,993518552 | 0,003464 | 0,024331 |
| Ecrg4 | 78896 | 16,661888 | 7,733265907 | 2,645686011 | 0,003467 | 0,024333 |
| Ppp3cb | 19056 | 58,571822 | 6,174640654 | 2,11310609 | 0,003477 | 0,024374 |
| Ran | 19384 | 224,13178 | 5,754454902 | 1,969499301 | 0,00348 | 0,024374 |
| Mzt2 | 72083 | 71,704333 | 6,638211831 | 2,271925786 | 0,00348 | 0,024374 |
| Tirap | 117149 | 27,635254 | -7,207922863 | 2,467210487 | 0,003484 | 0,024381 |
| 2410004P03Rik | 73667 | 138,2602 | 7,031078121 | 2,407309197 | 0,003492 | 0,024423 |
| Ankrd11 | 77087 | 602,05811 | 4,196727966 | 1,437465856 | 0,003506 | 0,0245 |
| Rnft1 | 76892 | 76,631246 | -6,142192804 | 2,104791716 | 0,003521 | 0,024588 |
| Raver2 | 242570 | 193,90272 | -6,45548827 | 2,213187638 | 0,003536 | 0,024678 |
| Pakap | 677884 | 36,477594 | 5,532040983 | 1,898084092 | 0,003562 | 0,024843 |
| Pja1 | 18744 | 49,953604 | 6,098007823 | 2,092732395 | 0,003569 | 0,024876 |
| Gm15927 | 1,03E+08 | 79,192144 | -6,38605573 | 2,192122814 | 0,003578 | 0,024916 |
| Jun | 16476 | 342,84004 | 4,78596613 | 1,644318132 | 0,003607 | 0,025078 |
| Cdk8 | 264064 | 623,28751 | -6,229144474 | 2,140004699 | 0,003605 | 0,025078 |
| Bud31 | 231889 | 102,23546 | 6,93938017 | 2,384256649 | 0,003609 | 0,025078 |
| Gm35857 | 1,03E+08 | 19,675111 | -6,315828121 | 2,171051039 | 0,003625 | 0,025172 |
| Dcaf17 | 75763 | 127,27447 | 4,816779229 | 1,656764724 | 0,003645 | 0,02528 |
| Ston2 | 108800 | 840,02751 | -5,403366767 | 1,858526572 | 0,003645 | 0,02528 |
| Wwtr1 | 97064 | 18023,623 | -7,263706439 | 2,498914388 | 0,003652 | 0,02531 |
| Zfp952 | 240067 | 40,145117 | 6,609809664 | 2,274250252 | 0,003657 | 0,025323 |
| Tbc1d23 | 67581 | 61,519586 | 5,485779257 | 1,888111138 | 0,003667 | 0,025381 |
| Naa50 | 72117 | 211,16265 | -6,188925815 | 2,131240924 | 0,003685 | 0,025488 |
| Car14 | 23831 | 268,7537 | 7,034705152 | 2,424326675 | 0,003711 | 0,025649 |
| Ywhaq | 22630 | 599,77807 | 4,534965549 | 1,563450126 | 0,003724 | 0,025721 |
| Dmxl2 | 235380 | 57,609465 | -5,613176823 | 1,936336981 | 0,003745 | 0,025819 |
| Phactr1 | 218194 | 200,00127 | 5,483003665 | 1,891225314 | 0,003741 | 0,025819 |
| Fads3 | 60527 | 54,770509 | 6,804668415 | 2,347434594 | 0,003746 | 0,025819 |
| Ttc30b | 72421 | 41,78631 | -8,439160702 | 2,912984098 | 0,003766 | 0,02594 |
| Slc22a23 | 73102 | 258,03495 | 6,476619582 | 2,235976753 | 0,003773 | 0,025966 |
| Zmym4 | 67785 | 54,855426 | 5,860867648 | 2,023663338 | 0,003778 | 0,02598 |
| Swt1 | 66875 | 172,66349 | 6,031694904 | 2,082992445 | 0,003783 | 0,026002 |
| Vdac3 | 22335 | 334,64685 | 4,644038499 | 1,60403898 | 0,003789 | 0,026023 |

| | | | | | | |
|---|---|---|---|---|---|---|
| Ubap2 | 68926 | 70,314771 | 6,006822621 | 2,074977733 | 0,003793 | 0,026032 |
| Lsm12 | 268490 | 63,314676 | 5,409698711 | 1,869530359 | 0,003808 | 0,026109 |
| Yes1 | 22612 | 13175,884 | -7,139224584 | 2,467300848 | 0,003809 | 0,026109 |
| Cbx3 | 12417 | 944,22693 | 4,400407781 | 1,521013671 | 0,003815 | 0,026129 |
| Stxbp4 | 20913 | 54,9477 | 5,852068007 | 2,023043151 | 0,003819 | 0,026142 |
| Iqcd | 75732 | 48,828889 | -7,076048085 | 2,44636219 | 0,003822 | 0,026143 |
| Git1 | 216963 | 55,957003 | 7,160397897 | 2,478674879 | 0,003867 | 0,026425 |
| Pphln1 | 223828 | 423,20804 | -6,106081846 | 2,113804627 | 0,003869 | 0,026425 |
| Invs | 16348 | 48,10424 | -5,366509185 | 1,858312207 | 0,003879 | 0,026477 |
| Rnf220 | 66743 | 109,21841 | 6,608953582 | 2,290026792 | 0,003902 | 0,026616 |
| Cdc42se2 | 72729 | 120,97524 | 5,89499895 | 2,043511893 | 0,003917 | 0,026702 |
| Kif3c | 16570 | 65,976351 | 6,544493304 | 2,271684696 | 0,003965 | 0,027011 |
| Ifrd1 | 15982 | 288,9675 | 4,683579844 | 1,625996823 | 0,003971 | 0,027014 |
| Sae1 | 56459 | 31,504928 | 7,577258214 | 2,630430348 | 0,003969 | 0,027014 |
| Ofd1 | 237222 | 23,395414 | 6,450005397 | 2,239548009 | 0,003976 | 0,027029 |
| Ric8b | 237422 | 76,253445 | 6,163269231 | 2,140241413 | 0,00398 | 0,027039 |
| Acvr2b | 11481 | 39,746387 | 5,634889583 | 1,957309922 | 0,003991 | 0,027091 |
| Rpl32 | 19951 | 9171,2956 | -5,432746294 | 1,887955506 | 0,004007 | 0,027166 |
| Ngef | 53972 | 570,9677 | -4,546876281 | 1,579988663 | 0,004005 | 0,027166 |
| Ankrd49 | 56503 | 42,784452 | 6,944608038 | 2,415237583 | 0,004036 | 0,027342 |
| Mark2 | 13728 | 72,435232 | 4,349859707 | 1,512939184 | 0,004039 | 0,027343 |
| Skiv2l | 108077 | 68,040185 | 5,201887403 | 1,81001623 | 0,004054 | 0,027425 |
| Fam8a1 | 97863 | 117,86255 | 5,054761869 | 1,759088705 | 0,004059 | 0,027444 |
| Kif1c | 16562 | 241,46191 | -5,685645789 | 1,978838657 | 0,004063 | 0,027451 |
| ND6 | 17722 | 16706,742 | -4,115290977 | 1,432453848 | 0,004067 | 0,02746 |
| Nfat5 | 54446 | 434,5846 | 4,336870575 | 1,509959075 | 0,004077 | 0,027504 |
| Coa6 | 67892 | 75,741246 | -6,85594889 | 2,387248646 | 0,00408 | 0,027509 |
| Dnajb2 | 56812 | 362,62798 | 4,650753031 | 1,619996303 | 0,004094 | 0,027583 |
| Kctd14 | 233529 | 37,310639 | 6,572143822 | 2,289649716 | 0,0041 | 0,027606 |
| Kat14 | 228714 | 23,119961 | 5,959383554 | 2,076416058 | 0,004104 | 0,027616 |
| Cbx1 | 12412 | 412,36812 | 4,650816765 | 1,620739175 | 0,00411 | 0,027639 |
| Prpf6 | 68879 | 51,654372 | 6,067462305 | 2,11509515 | 0,004122 | 0,0277 |
| Zfp428 | 232969 | 14,948885 | -6,741821355 | 2,350349678 | 0,004125 | 0,0277 |

| | | | | | | |
|---|---|---|---|---|---|---|
| Pef1 | 67898 | 68,059888 | 5,96059549 | 2,079198154 | 0,004147 | 0,027809 |
| Peak1 | 244895 | 22,88098 | 6,864620389 | 2,394903218 | 0,004152 | 0,027809 |
| Abhd5 | 67469 | 56,073929 | 7,903011325 | 2,757010841 | 0,00415 | 0,027809 |
| Tmem216 | 68642 | 17,480269 | 7,689192521 | 2,682467185 | 0,004151 | 0,027809 |
| Hmg20a | 66867 | 81,519411 | -5,671243901 | 1,979017383 | 0,004161 | 0,027847 |
| LOC118568277 | 1,19E+08 | 6180,0479 | -3,753152715 | 1,30987772 | 0,004167 | 0,027847 |
| Rpl36 | 54217 | 4760,5768 | -5,196928282 | 1,813738297 | 0,004166 | 0,027847 |
| Ccdc186 | 213993 | 52,990791 | 5,73138045 | 2,001186663 | 0,004183 | 0,027941 |
| Senp1 | 223870 | 93,072386 | 5,374986669 | 1,878603934 | 0,004221 | 0,028173 |
| Wipi1 | 52639 | 259,36582 | -5,914444955 | 2,067527149 | 0,004228 | 0,0282 |
| Nadk2 | 68646 | 339,26075 | 4,6339467 | 1,620121998 | 0,004233 | 0,028216 |
| Tspoap1 | 207777 | 41,458106 | 4,751450072 | 1,661532198 | 0,004241 | 0,028248 |
| Nae1 | 234664 | 27,400836 | 6,974955568 | 2,43940472 | 0,004246 | 0,028264 |
| Memo1 | 76890 | 6622,9594 | -5,131557346 | 1,7948455 | 0,004249 | 0,028266 |
| Tufm | 233870 | 59,758966 | 5,806851864 | 2,032246309 | 0,004272 | 0,028398 |
| Ripk1 | 19766 | 204,69265 | -5,882204977 | 2,058953157 | 0,004278 | 0,028421 |
| Purg | 75029 | 51,859789 | -6,147012184 | 2,151830099 | 0,004281 | 0,028424 |
| Mtcl1 | 68617 | 169,53756 | -5,667216909 | 1,984244172 | 0,004289 | 0,028453 |
| 1110051M20Rik | 228356 | 122,9861 | 6,14698892 | 2,15288042 | 0,0043 | 0,028512 |
| Rps17 | 20068 | 274,66541 | 4,13100705 | 1,447156852 | 0,00431 | 0,028515 |
| Cdon | 57810 | 72,573283 | -5,231615316 | 1,832528085 | 0,004306 | 0,028515 |
| Heph | 15203 | 23,099912 | -7,273746043 | 2,548073047 | 0,004309 | 0,028515 |
| Akt2 | 11652 | 234,26312 | 5,479699591 | 1,919806803 | 0,004313 | 0,028521 |
| Exd2 | 97827 | 94,750931 | 6,050240409 | 2,120650468 | 0,004331 | 0,028617 |
| Atf7 | 223922 | 47,406479 | -5,136515879 | 1,800961326 | 0,004343 | 0,028636 |
| Ttc13 | 234875 | 46,658992 | 6,397978642 | 2,24318082 | 0,004342 | 0,028636 |
| Commd2 | 52245 | 100,98292 | 7,634920841 | 2,677080596 | 0,004345 | 0,028636 |
| Stk19 | 54402 | 33,54409 | -7,045408425 | 2,470015814 | 0,004339 | 0,028636 |
| Gp1bb | 14724 | 107,51674 | -6,873025412 | 2,411896357 | 0,004377 | 0,028827 |
| Tmsb4x | 19241 | 13917,573 | -4,260787918 | 1,495420706 | 0,004383 | 0,028845 |
| Cend1 | 57754 | 32,219315 | 6,942001654 | 2,436701115 | 0,004387 | 0,028852 |
| Plpp3 | 67916 | 28463,311 | -3,556200574 | 1,248480977 | 0,004394 | 0,02888 |
| Cfap46 | 212124 | 87,913109 | 6,860269105 | 2,408959829 | 0,004402 | 0,028916 |

| | | | | | | |
|---|---|---|---|---|---|---|
| Tnc | 21923 | 131,03121 | 7,798400224 | 2,738962992 | 0,00441 | 0,028952 |
| Exosc2 | 227715 | 119,30507 | 5,06195378 | 1,778395801 | 0,004422 | 0,02901 |
| Oard1 | 106821 | 26,497579 | 7,350385652 | 2,5830067 | 0,004432 | 0,029054 |
| Cisd1 | 52637 | 209,27231 | 4,740097306 | 1,666477543 | 0,00445 | 0,029152 |
| Pias2 | 17344 | 117,43403 | 6,55534794 | 2,304944235 | 0,004455 | 0,029164 |
| Ttc12 | 235330 | 26,436725 | -6,401377046 | 2,251332768 | 0,004464 | 0,029191 |
| Vim | 22352 | 305,90555 | 4,791456198 | 1,685195973 | 0,004465 | 0,029191 |
| Sdhaf2 | 66072 | 41,469067 | -6,24745311 | 2,197390974 | 0,004467 | 0,029191 |
| Klf6 | 23849 | 196,13344 | 5,270620785 | 1,854029098 | 0,004472 | 0,029202 |
| Disc1 | 244667 | 34,672177 | -5,821403859 | 2,049053589 | 0,004497 | 0,029345 |
| Ctnnbl1 | 66642 | 38,168613 | 5,820788214 | 2,049572262 | 0,004511 | 0,029394 |
| Nmrk1 | 225994 | 42,935743 | 5,5445121 | 1,952139081 | 0,004508 | 0,029394 |
| Mbd4 | 17193 | 29,97247 | 6,678312973 | 2,351652215 | 0,004514 | 0,029394 |
| Lgals4 | 16855 | 449,32343 | -6,353259751 | 2,237338604 | 0,004516 | 0,029394 |
| Dnajc4 | 57431 | 110,8587 | 6,75516353 | 2,379530445 | 0,004527 | 0,029431 |
| Necap1 | 67602 | 54,557091 | 6,560185165 | 2,3108754 | 0,004528 | 0,029431 |
| Ttc5 | 219022 | 96,52526 | 4,852929313 | 1,70977914 | 0,004535 | 0,029458 |
| Slc25a23 | 66972 | 82,57212 | 5,702573993 | 2,009622927 | 0,004545 | 0,029503 |
| Ndufb11 | 104130 | 5555,4365 | -5,395240216 | 1,901759058 | 0,004554 | 0,029545 |
| Plcd4 | 18802 | 2453,0952 | -4,135476999 | 1,457972559 | 0,004562 | 0,029571 |
| Slc35b4 | 58246 | 46,121077 | 5,863452167 | 2,067302568 | 0,004564 | 0,029571 |
| Mrpl19 | 56284 | 34,481186 | 5,906852447 | 2,083054199 | 0,004573 | 0,029598 |
| A430027H14Rik | 1,15E+08 | 330,4476 | -6,056381514 | 2,135857137 | 0,004574 | 0,029598 |
| Vps13d | 230895 | 73,872617 | 5,656259619 | 1,9955883 | 0,004591 | 0,029685 |
| Zfp462 | 242466 | 89,557573 | 4,090917809 | 1,443545087 | 0,004598 | 0,029685 |
| Axl | 26362 | 359,38114 | -5,082972717 | 1,793454634 | 0,004594 | 0,029685 |
| Fam189a2 | 381217 | 59,028719 | 7,642759566 | 2,696996894 | 0,0046 | 0,029685 |
| Hsf2 | 15500 | 416,95708 | -6,208729549 | 2,191682285 | 0,004613 | 0,029753 |
| Pink1 | 68943 | 97,155758 | 5,562179041 | 1,963605401 | 0,004617 | 0,029755 |
| ND2 | 17717 | 63300,339 | -4,38873673 | 1,549864271 | 0,00463 | 0,02982 |
| Dnajc24 | 99349 | 67,206718 | 6,590531502 | 2,327728278 | 0,004636 | 0,02982 |
| Aph1c | 68318 | 52,229454 | 7,029764383 | 2,482798745 | 0,004635 | 0,02982 |
| Hnrnpl | 15388 | 101,71966 | 4,855660052 | 1,716294549 | 0,004667 | 0,030003 |

| | | | | | | |
|---|---|---|---|---|---|---|
| Dlc1 | 50768 | 141,51369 | 6,447201241 | 2,27984253 | 0,004685 | 0,0301 |
| Oscp1 | 230751 | 41,311473 | 5,729851329 | 2,027051926 | 0,004703 | 0,030157 |
| Adgrb3 | 210933 | 96,032261 | 6,003702176 | 2,123777476 | 0,0047 | 0,030157 |
| Parg | 26430 | 199,77057 | 7,568284957 | 2,677186071 | 0,004699 | 0,030157 |
| Ttc23 | 67009 | 64,534014 | -5,287438041 | 1,87095674 | 0,004712 | 0,030197 |
| Daam2 | 76441 | 4518,61 | -4,35715727 | 1,542134844 | 0,004722 | 0,030239 |
| Gm39673 | 1,05E+08 | 73,80597 | 5,740680752 | 2,032714983 | 0,004741 | 0,030339 |
| Otud6b | 72201 | 70,456869 | 4,526812579 | 1,603091731 | 0,004746 | 0,030352 |
| LOC115486538 | 1,15E+08 | 87,170875 | -5,23061072 | 1,85246876 | 0,004749 | 0,030352 |
| Cnpy4 | 66455 | 61,584721 | -6,389884195 | 2,263715418 | 0,004761 | 0,030378 |
| Atat1 | 73242 | 46,216309 | 5,600165413 | 1,98397482 | 0,004762 | 0,030378 |
| Gm34739 | 1,03E+08 | 13,528502 | -6,988415964 | 2,47561137 | 0,004759 | 0,030378 |
| Vkorc1l1 | 69568 | 50,392471 | -6,393893256 | 2,265460895 | 0,004768 | 0,030393 |
| Mplkip | 66308 | 36,447233 | 7,388472884 | 2,619335795 | 0,004791 | 0,030525 |
| Fam177a | 73385 | 517,52843 | -5,614953558 | 1,991619233 | 0,004813 | 0,030624 |
| Fam177a2 | 1E+08 | 517,52843 | -5,614953558 | 1,991619233 | 0,004813 | 0,030624 |
| Gm52861 | 1,15E+08 | 82,019779 | -5,208400391 | 1,847702507 | 0,00482 | 0,030644 |
| Taf1d | 75316 | 881,64657 | 6,826668768 | 2,421945614 | 0,004822 | 0,030644 |
| Clcc1 | 229725 | 226,15101 | 4,483987092 | 1,591022106 | 0,004828 | 0,030659 |
| Elavl2 | 15569 | 95,670384 | 6,257744319 | 2,221136797 | 0,004842 | 0,03073 |
| Nfib | 18028 | 3839,2419 | -4,5082451 | 1,600641223 | 0,004855 | 0,03079 |
| Ppp1cc | 19047 | 303,97573 | 4,52928224 | 1,608434746 | 0,004863 | 0,030825 |
| Pdp2 | 382051 | 145,52984 | 6,923230864 | 2,459179802 | 0,004874 | 0,030872 |
| Scyl1 | 78891 | 211,26363 | -6,319220038 | 2,245710887 | 0,004894 | 0,030983 |
| Naxd | 69225 | 108,61481 | -6,15426779 | 2,187253439 | 0,004898 | 0,030983 |
| Luc7l3 | 67684 | 12455,245 | -4,381463325 | 1,557967324 | 0,004919 | 0,031099 |
| Mtif2 | 76784 | 123,12151 | -4,657489706 | 1,65658835 | 0,004931 | 0,031143 |
| Glmp | 56700 | 89,621275 | 6,776948933 | 2,410495056 | 0,004932 | 0,031143 |
| Calb1 | 12307 | 22,198787 | -5,906609585 | 2,101277607 | 0,004939 | 0,031149 |
| Dnajc11 | 230935 | 124,30615 | 8,016084375 | 2,85172458 | 0,004939 | 0,031149 |
| Mark3 | 17169 | 1753,7258 | -5,422650484 | 1,930025443 | 0,00496 | 0,031259 |
| 3830406C13Rik | 218734 | 89,489319 | 5,018217784 | 1,786262308 | 0,004964 | 0,031267 |
| Senp3 | 80886 | 43,304934 | 6,201663839 | 2,208309034 | 0,00498 | 0,031345 |

| | | | | | | |
|---|---|---|---|---|---|---|
| Tm9sf3 | 107358 | 424,32947 | 5,305815214 | 1,889776636 | 0,004991 | 0,03139 |
| Tspan12 | 269831 | 291,0752 | 6,028208142 | 2,147524564 | 0,005 | 0,03139 |
| Fkbp1a | 14225 | 1629,2825 | -5,968220212 | 2,125899415 | 0,004995 | 0,03139 |
| Apmap | 71881 | 79,728995 | 6,275327476 | 2,235526326 | 0,004999 | 0,03139 |
| Insig1 | 231070 | 3986,437 | -4,294845676 | 1,5305215 | 0,005014 | 0,031432 |
| Chchd3 | 66075 | 148,86343 | 5,640503979 | 2,010206101 | 0,005017 | 0,031432 |
| Klf4 | 16600 | 252,75022 | 4,872217008 | 1,736275523 | 0,005014 | 0,031432 |
| Trmt1 | 212528 | 177,85548 | -5,913050799 | 2,107430289 | 0,005019 | 0,031432 |
| Col8a2 | 329941 | 15,671435 | 6,54368027 | 2,332799624 | 0,00503 | 0,031484 |
| Xpa | 22590 | 40,882331 | 7,234401111 | 2,580294073 | 0,005052 | 0,031598 |
| Phf21b | 271305 | 31,214572 | 5,482366195 | 1,956126792 | 0,005068 | 0,031666 |
| Nup54 | 269113 | 54,034297 | 6,556704174 | 2,33949618 | 0,005069 | 0,031666 |
| Frem2 | 242022 | 129,02236 | 5,392480102 | 1,925340175 | 0,005098 | 0,031805 |
| Lrrc4 | 192198 | 18,208275 | 6,895722148 | 2,461944691 | 0,005096 | 0,031805 |
| Zfyve9 | 230597 | 54,322978 | 5,59290819 | 1,997774199 | 0,005117 | 0,031906 |
| Micu2 | 68514 | 55,203913 | 7,450229874 | 2,661433345 | 0,005121 | 0,031909 |
| Mical2 | 320878 | 68,036701 | 5,860612319 | 2,09438924 | 0,005138 | 0,031997 |
| Celsr2 | 53883 | 305,34342 | -5,177036983 | 1,850470354 | 0,005147 | 0,032032 |
| Gstz1 | 14874 | 42,370588 | 5,751602904 | 2,057738743 | 0,005188 | 0,032267 |
| Frmd4b | 232288 | 91,79135 | 6,335936176 | 2,267112636 | 0,005195 | 0,032267 |
| Fundc1 | 72018 | 118,41456 | 6,863382304 | 2,455816664 | 0,005194 | 0,032267 |
| Vamp4 | 53330 | 105,21823 | 6,309388327 | 2,258359486 | 0,005209 | 0,032319 |
| Per2 | 18627 | 228,93344 | 7,105891264 | 2,543425971 | 0,005209 | 0,032319 |
| Gm7697 | 665578 | 21,917927 | 5,490763688 | 1,966025512 | 0,005225 | 0,032396 |
| Gm46429 | 1,08E+08 | 262,62133 | -4,918948841 | 1,761591548 | 0,005233 | 0,032425 |
| Mtfr1l | 76824 | 117,41587 | -6,979513871 | 2,50011261 | 0,005244 | 0,03247 |
| Mgat3 | 17309 | 193,02715 | 6,396237633 | 2,29235189 | 0,005267 | 0,032593 |
| Surf4 | 20932 | 239,45532 | 5,433600674 | 1,947970127 | 0,005281 | 0,032662 |
| Plaat3 | 225845 | 357,13493 | 4,041419571 | 1,449724862 | 0,005308 | 0,032788 |
| Copz1 | 56447 | 111,783 | 6,824578548 | 2,448052241 | 0,005307 | 0,032788 |
| Shroom2 | 110380 | 198,44339 | 5,789884023 | 2,077394942 | 0,005318 | 0,032831 |
| Fos | 14281 | 2039,7873 | 3,897490275 | 1,399048417 | 0,005339 | 0,03294 |
| Galnt11 | 231050 | 33,833275 | -6,383710663 | 2,292464413 | 0,005359 | 0,032977 |

| | | | | | | |
|---|---|---|---|---|---|---|
| ago-01 | 236511 | 43,947015 | -5,675573764 | 2,038035337 | 0,005356 | 0,032977 |
| LOC118568708 | 1,19E+08 | 40,436749 | 6,313521461 | 2,26684047 | 0,00535 | 0,032977 |
| Tmx1 | 72736 | 42363,565 | -7,441492203 | 2,672035612 | 0,005354 | 0,032977 |
| Dhdds | 67422 | 74,088165 | 6,921832956 | 2,486304492 | 0,00537 | 0,033024 |
| Cenpb | 12616 | 19,625012 | 6,172906191 | 2,217485103 | 0,005374 | 0,033029 |
| Hagh | 14651 | 86,418698 | 6,119675753 | 2,198597243 | 0,005378 | 0,033038 |
| Cdyl | 12593 | 79,189196 | 6,652780859 | 2,392589764 | 0,005426 | 0,033311 |
| Gm52026 | 1,15E+08 | 192,21815 | 5,714390609 | 2,055669047 | 0,005439 | 0,033368 |
| Exoc6b | 75914 | 156,35294 | 5,490016147 | 1,975507733 | 0,005452 | 0,033415 |
| Arih1 | 23806 | 417,33199 | 5,433903649 | 1,955371046 | 0,005453 | 0,033415 |
| Cisd2 | 67006 | 116,52059 | 5,16206245 | 1,859055037 | 0,005491 | 0,033627 |
| Ppp2r1a | 51792 | 154,13356 | 4,996087475 | 1,800148093 | 0,005514 | 0,033744 |
| LOC118567378 | 1,19E+08 | 244,61671 | -5,571999497 | 2,007808546 | 0,005517 | 0,033746 |
| H2ax | 15270 | 11,069594 | -8,270796303 | 2,980841705 | 0,005526 | 0,033778 |
| Rabl2 | 68708 | 20,054606 | 7,09734413 | 2,55872862 | 0,005541 | 0,033845 |
| Zc2hc1a | 67306 | 22,753632 | 6,685567468 | 2,410420666 | 0,005544 | 0,033845 |
| 1810013L24Rik | 69053 | 83,852385 | 5,645931234 | 2,036124957 | 0,005556 | 0,033879 |
| Bnip2 | 12175 | 174,93524 | 6,07723491 | 2,191567165 | 0,005554 | 0,033879 |
| Riok2 | 67045 | 59,581284 | -7,738303413 | 2,79095354 | 0,00556 | 0,033884 |
| Mrpl42 | 67270 | 250,75973 | 5,14357327 | 1,855479405 | 0,00557 | 0,033919 |
| Dcaf1 | 321006 | 28,006273 | 6,674378388 | 2,411548051 | 0,005646 | 0,034348 |
| 2310039H08Rik | 67101 | 51,787822 | 6,373761222 | 2,302986803 | 0,005647 | 0,034348 |
| Syde1 | 71709 | 2227,0465 | -5,837399746 | 2,109835231 | 0,005662 | 0,034417 |
| Tm4sf1 | 17112 | 63,936644 | 7,005416308 | 2,532282837 | 0,005667 | 0,034429 |
| Morf4l2 | 56397 | 74,025139 | 5,798865757 | 2,097646128 | 0,005702 | 0,034617 |
| Recql5 | 170472 | 13,104205 | 7,011155854 | 2,536496889 | 0,005708 | 0,034634 |
| Actr3b | 242894 | 47,424877 | 5,856738269 | 2,11963179 | 0,005726 | 0,034721 |
| Gm38667 | 1,04E+08 | 36,918262 | -7,589950796 | 2,747608176 | 0,005738 | 0,034775 |
| Lrch1 | 380916 | 74,910683 | 6,628635307 | 2,400021881 | 0,005746 | 0,034804 |
| Nek6 | 59126 | 38,626735 | -6,094761645 | 2,207257755 | 0,005758 | 0,034833 |
| Mir22hg | 1E+08 | 50,518501 | -6,025519379 | 2,182181514 | 0,005758 | 0,034833 |
| Pum1 | 80912 | 78,707614 | 5,132690491 | 1,859477564 | 0,005775 | 0,034914 |
| Zfp292 | 30046 | 151,88202 | 5,115587424 | 1,853718814 | 0,005787 | 0,034962 |

| | | | | | | |
|---|---|---|---|---|---|---|
| Dtd2 | 328092 | 36,574965 | 6,482127832 | 2,349512273 | 0,005799 | 0,035017 |
| Kcnd1 | 16506 | 26,736736 | 6,976285635 | 2,530962581 | 0,005845 | 0,03527 |
| Galnt6 | 207839 | 21,142137 | 6,308437328 | 2,288908989 | 0,00585 | 0,035279 |
| Noc2l | 57741 | 20,176998 | 7,512112945 | 2,726861828 | 0,005872 | 0,035369 |
| Grk5 | 14773 | 51,997992 | 8,421093772 | 3,056766967 | 0,005871 | 0,035369 |
| Sorbs1 | 20411 | 1033,7505 | 4,332434808 | 1,573016701 | 0,005883 | 0,035392 |
| Thap3 | 69876 | 15,197114 | 6,310534472 | 2,291266234 | 0,005884 | 0,035392 |
| Gm36245 | 1,03E+08 | 10,895466 | -7,250501898 | 2,632669445 | 0,005886 | 0,035392 |
| Lrrc51 | 69358 | 56,400534 | 6,785350297 | 2,46430184 | 0,005897 | 0,035414 |
| Lrrc8c | 100604 | 61,914176 | 6,20962352 | 2,255077378 | 0,005894 | 0,035414 |
| Ahnak | 66395 | 218,03188 | 5,591998086 | 2,032446839 | 0,005935 | 0,035619 |
| St3gal2 | 20444 | 82,888202 | 4,55691887 | 1,656521649 | 0,005943 | 0,035627 |
| Sec13 | 110379 | 117,40535 | 6,300888753 | 2,290433089 | 0,005942 | 0,035627 |
| Cct5 | 12465 | 186,40907 | 4,556966106 | 1,656778001 | 0,00595 | 0,035649 |
| Pdcd5 | 56330 | 9714,1746 | -6,354549458 | 2,310698657 | 0,005959 | 0,035676 |
| Gja1 | 14609 | 23262,669 | -3,99507115 | 1,45304855 | 0,00597 | 0,0357 |
| Fahd1 | 68636 | 28,496028 | 8,388266361 | 3,050895457 | 0,00597 | 0,0357 |
| Sf3b3 | 101943 | 96,41631 | 6,186457891 | 2,250884216 | 0,005988 | 0,035764 |
| 1700007K13Rik | 69327 | 65,131053 | 7,201157341 | 2,619991164 | 0,005986 | 0,035764 |
| Rgs7 | 24012 | 43,389724 | 6,162677128 | 2,242506547 | 0,005994 | 0,035779 |
| Cenpm | 66570 | 46,775266 | 6,903811221 | 2,512564668 | 0,006001 | 0,035802 |
| Knop1 | 66356 | 95,854774 | 5,612727321 | 2,045598849 | 0,006073 | 0,036209 |
| Hdac4 | 208727 | 50,547408 | 5,48136337 | 1,998029617 | 0,006081 | 0,036219 |
| Cadps | 27062 | 39,276834 | 8,218287208 | 2,995741422 | 0,006082 | 0,036219 |
| Samd4 | 74480 | 199,57269 | 5,644490113 | 2,058108943 | 0,006096 | 0,036271 |
| Gm19935 | 1,01E+08 | 193,5405 | 6,891038254 | 2,512716222 | 0,006098 | 0,036271 |
| Grid2 | 14804 | 104,56729 | 5,790043896 | 2,111578132 | 0,006106 | 0,036295 |
| Gnl3 | 30877 | 23,876663 | 6,83748937 | 2,494236083 | 0,006119 | 0,036354 |
| Cep120 | 225523 | 21,313851 | 6,768378426 | 2,469324206 | 0,006126 | 0,036369 |
| Coq10b | 67876 | 25,369046 | -6,276867719 | 2,29019404 | 0,00613 | 0,036373 |
| Dpm2 | 13481 | 48,23342 | -7,077457405 | 2,582729766 | 0,006138 | 0,036401 |
| Smarcb1 | 20587 | 197,47705 | 5,691012935 | 2,077112352 | 0,006146 | 0,036428 |
| Gcn1 | 231659 | 33,446241 | 6,040678475 | 2,205156324 | 0,006156 | 0,036464 |

| | | | | | | |
|---|---|---|---|---|---|---|
| Snx15 | 69024 | 39,132548 | 7,977108592 | 2,912761005 | 0,006169 | 0,036495 |
| Rabl6 | 227624 | 14591,824 | -7,644305954 | 2,791199467 | 0,006168 | 0,036495 |
| Vps16 | 80743 | 27,950017 | 5,958762035 | 2,176846019 | 0,006194 | 0,036622 |
| Irs2 | 384783 | 51,430339 | 5,227854257 | 1,910275364 | 0,006206 | 0,036671 |
| Caprin2 | 232560 | 55,964346 | 5,89683018 | 2,155540544 | 0,006225 | 0,036765 |
| Zup1 | 72580 | 260,07991 | 5,03705784 | 1,841574531 | 0,006234 | 0,036796 |
| Glmn | 170823 | 31,185681 | 6,872790705 | 2,512949796 | 0,006239 | 0,036801 |
| Teddm1b | 433365 | 99,103072 | -5,133535248 | 1,877882806 | 0,006263 | 0,036921 |
| Izumo4 | 71564 | 5453,9243 | -5,267000232 | 1,92693694 | 0,006269 | 0,036936 |
| Bet1l | 54399 | 67,646197 | 4,986162124 | 1,824795591 | 0,006287 | 0,036994 |
| Pld1 | 18805 | 49,565437 | 6,827114335 | 2,498405407 | 0,006284 | 0,036994 |
| Gabpa | 14390 | 64,927959 | -6,359265039 | 2,327680985 | 0,006295 | 0,03702 |
| Trpt1 | 107328 | 26,0594 | -6,652041465 | 2,435101078 | 0,0063 | 0,037031 |
| Wipf1 | 215280 | 29,439732 | -6,897865377 | 2,52590438 | 0,006317 | 0,037108 |
| Msantd2 | 235184 | 301,92319 | 5,187645327 | 1,900435698 | 0,006339 | 0,037192 |
| Tfb1m | 224481 | 64,900942 | -7,197607232 | 2,636758282 | 0,006339 | 0,037192 |
| Tmem106b | 71900 | 363,70389 | 4,456586949 | 1,632906601 | 0,006348 | 0,037215 |
| Slc1a2 | 20511 | 49961,217 | -3,826066324 | 1,401937946 | 0,00635 | 0,037215 |
| Rchy1 | 68098 | 157,18075 | 4,748689631 | 1,740410208 | 0,006363 | 0,037265 |
| Mpp7 | 75739 | 68,837643 | 6,077513618 | 2,228166978 | 0,00638 | 0,037301 |
| Mcrip1 | 192173 | 62,683392 | 6,037967162 | 2,21362274 | 0,006379 | 0,037301 |
| Zfp512 | 269639 | 78,067631 | 6,4594327 | 2,368080632 | 0,006378 | 0,037301 |
| Dync2h1 | 110350 | 63,391823 | 5,316035945 | 1,949272206 | 0,006388 | 0,037324 |
| Wnk2 | 75607 | 79,266129 | 6,848147211 | 2,511249293 | 0,006392 | 0,037325 |
| Gba2 | 230101 | 45,745463 | 6,725377196 | 2,466682603 | 0,006401 | 0,037359 |
| Calcoco1 | 67488 | 49,199477 | 6,461651049 | 2,37112126 | 0,006427 | 0,03748 |
| Coa7 | 69893 | 20,357008 | -6,885715971 | 2,526832032 | 0,006429 | 0,03748 |
| Arsb | 11881 | 485,83692 | 5,604897248 | 2,058008826 | 0,00646 | 0,037638 |
| Mga | 29808 | 279,29869 | 4,154547043 | 1,525933595 | 0,006477 | 0,03771 |
| Tmem185b | 226351 | 60,001609 | 5,049999199 | 1,855386943 | 0,006493 | 0,037782 |
| Cfap52 | 71860 | 31,064063 | 7,272040058 | 2,672731272 | 0,006512 | 0,037872 |
| Zc3h13 | 67302 | 423,30076 | 5,129080276 | 1,885252717 | 0,006516 | 0,037872 |
| Gm26618 | 1,03E+08 | 35,58315 | -7,149349054 | 2,628987462 | 0,006539 | 0,037988 |

| | | | | | | |
|---|---|---|---|---|---|---|
| Morf4l1 | 21761 | 366,265 | 4,225275071 | 1,553858372 | 0,006544 | 0,03799 |
| Rdm1 | 66599 | 74,888217 | 5,293473849 | 1,947065344 | 0,006554 | 0,038028 |
| Prox1 | 19130 | 31,852981 | 5,495467834 | 2,022060855 | 0,006573 | 0,038114 |
| Yipf3 | 28064 | 41,386461 | 6,477457159 | 2,383996699 | 0,006587 | 0,03815 |
| Celf4 | 108013 | 122,99367 | 7,336589338 | 2,700180025 | 0,006586 | 0,03815 |
| Sidt2 | 214597 | 64,949045 | 6,212706789 | 2,287206513 | 0,006602 | 0,038216 |
| Gm51666 | 1,15E+08 | 219,34495 | 5,121627271 | 1,885759358 | 0,006609 | 0,038233 |
| Rfk | 54391 | 41,67144 | 5,81162233 | 2,140362633 | 0,006623 | 0,038291 |
| Pdpr | 319518 | 20,938843 | 6,12590817 | 2,258188449 | 0,006673 | 0,038553 |
| Ermard | 381062 | 71,599192 | 6,290110973 | 2,318837167 | 0,006675 | 0,038553 |
| Dnajc15 | 66148 | 89,030804 | 6,176655265 | 2,278368025 | 0,006708 | 0,038718 |
| Dync1h1 | 13424 | 172,02155 | 4,062031353 | 1,499438018 | 0,006748 | 0,038926 |
| Socs3 | 12702 | 29,604991 | 5,976685828 | 2,207054397 | 0,006769 | 0,039026 |
| Npy | 109648 | 17,010744 | -8,497264003 | 3,139367682 | 0,006796 | 0,039158 |
| Ctdsp1 | 227292 | 55,231707 | 5,880714177 | 2,173247457 | 0,006811 | 0,03922 |
| Cul2 | 71745 | 88,051194 | -5,65211786 | 2,090011571 | 0,006844 | 0,039388 |
| Snx14 | 244962 | 122,71472 | 4,815477054 | 1,780781689 | 0,006848 | 0,03939 |
| Cdk5 | 12568 | 52,587178 | 5,231588565 | 1,935571818 | 0,006874 | 0,039518 |
| Gpbp1l1 | 77110 | 166,93741 | 5,472856214 | 2,025181769 | 0,006884 | 0,03955 |
| Timm8b | 30057 | 104,86995 | 6,300979569 | 2,332162028 | 0,006897 | 0,039602 |
| Ap1m1 | 11767 | 23,076689 | 6,59646933 | 2,442543866 | 0,00692 | 0,039713 |
| LOC118568133 | 1,19E+08 | 86,623569 | 4,413516123 | 1,63436667 | 0,006925 | 0,039715 |
| Lrrn3 | 16981 | 23,76634 | -6,746860534 | 2,498620286 | 0,006929 | 0,039717 |
| Ttc7b | 104718 | 28,492099 | -6,908633101 | 2,559404905 | 0,006948 | 0,039782 |
| Ogfod2 | 66627 | 42,920279 | 7,593820739 | 2,813236085 | 0,006948 | 0,039782 |
| Gm38809 | 1,05E+08 | 4897,6659 | -5,41454052 | 2,007449128 | 0,006992 | 0,039986 |
| Acbd6 | 72482 | 31,821028 | 5,660612487 | 2,098553007 | 0,006989 | 0,039986 |
| Tug1 | 544752 | 180,12228 | 4,489359187 | 1,665012086 | 0,007012 | 0,040075 |
| Arf5 | 11844 | 128,86794 | -5,145320198 | 1,908500602 | 0,007018 | 0,040087 |
| Hnrnph3 | 432467 | 447,15971 | -5,471854346 | 2,030308543 | 0,007037 | 0,040174 |
| Nploc4 | 217365 | 43,618804 | 6,21613365 | 2,306827137 | 0,007046 | 0,040178 |
| Zfand4 | 67492 | 24,538592 | 5,976723499 | 2,217880046 | 0,007043 | 0,040178 |
| Caskin2 | 140721 | 50,735575 | 6,339482237 | 2,353115821 | 0,007058 | 0,040226 |

| | | | | | | |
|---|---|---|---|---|---|---|
| Echdc1 | 52665 | 314,28466 | 5,246447141 | 1,947551802 | 0,007063 | 0,040229 |
| Ppm1k | 243382 | 233,62761 | 4,261036306 | 1,58218193 | 0,007078 | 0,040271 |
| Pdgfd | 71785 | 1017,2687 | -5,7532412 | 2,136214099 | 0,007077 | 0,040271 |
| Cahm | 1,01E+08 | 26,801433 | 7,20271339 | 2,674693267 | 0,007083 | 0,040275 |
| Aatf | 56321 | 77,681401 | 6,45338544 | 2,396827592 | 0,007093 | 0,040306 |
| Amotl1 | 75723 | 55,042609 | 5,672670987 | 2,107207539 | 0,007102 | 0,040335 |
| Lrrk2 | 66725 | 51,717989 | 6,231918065 | 2,31598704 | 0,007128 | 0,040459 |
| Rell1 | 100532 | 26,702381 | 6,780110933 | 2,52019407 | 0,007139 | 0,040498 |
| Itm2b | 16432 | 13306,942 | -4,497859422 | 1,67249677 | 0,00716 | 0,040597 |
| Vps13c | 320528 | 103,51346 | 4,651363809 | 1,729773984 | 0,007167 | 0,040611 |
| Zfp82 | 330502 | 26,400603 | -6,907373851 | 2,571603399 | 0,007231 | 0,040952 |
| Lrrc45 | 217366 | 18,610558 | 6,869568424 | 2,55853756 | 0,007254 | 0,041058 |
| Vps4a | 116733 | 42,688015 | -5,418781097 | 2,01858845 | 0,007265 | 0,041069 |
| Gm46558 | 1,08E+08 | 11,286235 | -4,998720434 | 1,862201941 | 0,007268 | 0,041069 |
| Dpy19l4 | 381510 | 250,03625 | 6,672097459 | 2,485311631 | 0,007261 | 0,041069 |
| Actr1a | 54130 | 57,0219 | 6,393740397 | 2,382521853 | 0,007283 | 0,041132 |
| Pprc1 | 226169 | 16,193219 | 8,109989095 | 3,022564802 | 0,007293 | 0,041164 |
| Ttbk2 | 140810 | 162,47765 | 6,039372274 | 2,252516182 | 0,007337 | 0,041385 |
| Slc7a2 | 11988 | 618,00979 | 4,761343044 | 1,776116169 | 0,007345 | 0,041412 |
| Ndufaf6 | 76947 | 54,525986 | 5,662996862 | 2,113202021 | 0,007366 | 0,041505 |
| Mto1 | 68291 | 16,590195 | -6,422127005 | 2,396845938 | 0,007375 | 0,041509 |
| Acvr2a | 11480 | 55,175265 | 6,452129666 | 2,407971631 | 0,007373 | 0,041509 |
| Zfyve27 | 319740 | 37,886925 | -5,84144412 | 2,180378536 | 0,007382 | 0,041524 |
| Scaf1 | 233208 | 60,481414 | 7,091998751 | 2,648279279 | 0,007407 | 0,041641 |
| Sh3glb2 | 227700 | 81,636804 | 4,8128919 | 1,797589276 | 0,007419 | 0,041686 |
| Lamtor3 | 56692 | 90,711094 | 6,007617762 | 2,245391464 | 0,007461 | 0,041874 |
| Ntn1 | 18208 | 84,272421 | 6,48306007 | 2,423056371 | 0,00746 | 0,041874 |
| Fcho2 | 218503 | 2314,2121 | -4,938716245 | 1,846425418 | 0,007479 | 0,041949 |
| Pigx | 72084 | 99,527576 | 6,969842506 | 2,606953206 | 0,007505 | 0,04205 |
| Ubxn11 | 67586 | 44,028004 | -6,023357847 | 2,252898782 | 0,007504 | 0,04205 |
| Pigyl | 66268 | 105,33436 | 5,230732192 | 1,957662745 | 0,007542 | 0,042231 |
| B230311B06Rik | 381914 | 29,617506 | -7,002317577 | 2,622098942 | 0,007574 | 0,042387 |
| Dnajb1 | 81489 | 271,80564 | 4,178401911 | 1,565253006 | 0,007597 | 0,042493 |

| | | | | | | |
|---|---|---|---|---|---|---|
| Phka2 | 110094 | 37,482237 | 5,769705373 | 2,161609684 | 0,007604 | 0,042507 |
| Rnf113a2 | 66381 | 20,322249 | -6,555207755 | 2,456309636 | 0,007614 | 0,04254 |
| Mettl16 | 67493 | 26,303485 | 5,27149786 | 1,975455102 | 0,007619 | 0,042544 |
| Ftl1 | 14325 | 15873,964 | -5,246103344 | 1,966072045 | 0,007623 | 0,042544 |
| Psmg2 | 107047 | 49,008871 | 7,593127347 | 2,846372968 | 0,007638 | 0,042604 |
| Amt | 434437 | 51,035551 | 4,581576513 | 1,718272199 | 0,007667 | 0,042741 |
| Cct8 | 12469 | 203,20786 | 4,607683343 | 1,728455585 | 0,007681 | 0,042794 |
| B3gat3 | 72727 | 82,643904 | 5,76558301 | 2,163378958 | 0,007697 | 0,042847 |
| Rbfa | 68731 | 33,073438 | 6,275975503 | 2,354975615 | 0,007699 | 0,042847 |
| 5430416N02Rik | 1,01E+08 | 41,998676 | 7,136522888 | 2,679370604 | 0,007733 | 0,043012 |
| Pgap2 | 233575 | 87,323634 | 4,86331952 | 1,826517317 | 0,007754 | 0,043102 |
| Arcn1 | 213827 | 19621,245 | -6,113410283 | 2,296630482 | 0,00777 | 0,043169 |
| Cnih1 | 12793 | 116,57523 | -4,631146374 | 1,740121648 | 0,007782 | 0,043211 |
| Itga7 | 16404 | 86,370521 | -5,735128069 | 2,158059735 | 0,007871 | 0,043683 |
| AU022252 | 230696 | 39,656214 | 5,885036916 | 2,214671997 | 0,007877 | 0,043691 |
| Alkbh1 | 211064 | 13,298304 | 6,85191256 | 2,578908017 | 0,007886 | 0,043717 |
| Cop1 | 26374 | 57,565262 | 4,53774009 | 1,708447755 | 0,007906 | 0,043802 |
| Hmgcs2 | 15360 | 22,059235 | -6,164909812 | 2,321455132 | 0,007916 | 0,043834 |
| Srsf7 | 225027 | 664,41086 | 4,383719311 | 1,651049799 | 0,007928 | 0,043877 |
| Tgfbr1 | 21812 | 59,914999 | -5,307344339 | 1,999058994 | 0,007933 | 0,043877 |
| Mpv17 | 17527 | 151,80047 | 4,672686622 | 1,760202821 | 0,00794 | 0,04389 |
| Tfdp1 | 21781 | 106,34777 | 5,242895591 | 1,975196704 | 0,007946 | 0,0439 |
| Prpf8 | 192159 | 370,81344 | -5,876865819 | 2,214394732 | 0,007956 | 0,043932 |
| Ncbp1 | 433702 | 59,401566 | 7,651438571 | 2,883713967 | 0,00797 | 0,043987 |
| Tmem248 | 71667 | 662,8081 | -5,013811517 | 1,889843215 | 0,007977 | 0,043994 |
| Ube2w | 66799 | 570,61 | 4,996668131 | 1,883475233 | 0,00798 | 0,043994 |
| Wdr60 | 217935 | 245,55289 | 4,878041716 | 1,839127845 | 0,007993 | 0,044026 |
| Fbxo2 | 230904 | 123,05477 | -5,648657434 | 2,129852897 | 0,007998 | 0,044026 |
| Uba1 | 22201 | 13234,738 | -6,043791875 | 2,278875404 | 0,007999 | 0,044026 |
| Rnpep | 215615 | 32,033681 | 5,497398054 | 2,073305802 | 0,008013 | 0,044077 |
| LOC118568151 | 1,19E+08 | 33,372972 | 6,020291611 | 2,270823218 | 0,008022 | 0,0441 |
| Taok2 | 381921 | 35,388884 | 5,106676048 | 1,92724311 | 0,008056 | 0,044237 |
| Psat1 | 107272 | 1521,0531 | -4,59643402 | 1,734598198 | 0,008053 | 0,044237 |

| | | | | | | |
|---|---|---|---|---|---|---|
| Ctnnd1 | 12388 | 132,10007 | -4,597580418 | 1,735631449 | 0,008075 | 0,044316 |
| Zbtb37 | 240869 | 145,50952 | 5,392013804 | 2,035776514 | 0,008082 | 0,044322 |
| Xpnpep3 | 321003 | 38,884387 | 5,912597435 | 2,232416201 | 0,008085 | 0,044322 |
| Smarcc2 | 68094 | 197,52402 | 4,146663269 | 1,565847639 | 0,008092 | 0,044341 |
| Nudcd2 | 52653 | 114,13587 | 5,095453461 | 1,924439567 | 0,008103 | 0,044349 |
| Hivep1 | 110521 | 100,02384 | 5,366249124 | 2,026675862 | 0,008102 | 0,044349 |
| Rps8 | 20116 | 273,83426 | 4,104657506 | 1,550663522 | 0,00812 | 0,044395 |
| Usp48 | 170707 | 138,40035 | 4,114532586 | 1,554323884 | 0,008117 | 0,044395 |
| Zfp318 | 57908 | 4012,782 | -4,865469229 | 1,838379839 | 0,00813 | 0,044427 |
| Trim24 | 21848 | 60,343622 | -5,813460462 | 2,197453396 | 0,008156 | 0,044542 |
| Aox1 | 11761 | 106,04145 | 5,727110384 | 2,165321957 | 0,008171 | 0,044599 |
| Cnot7 | 18983 | 447,45 | -5,721769937 | 2,163630009 | 0,008181 | 0,044628 |
| Retreg2 | 227298 | 104,18711 | 6,667907058 | 2,521806747 | 0,008191 | 0,044659 |
| Fbxw8 | 231672 | 131,75384 | -5,573189419 | 2,108587329 | 0,008215 | 0,044768 |
| Zdhhc8 | 27801 | 93,852379 | 5,791256833 | 2,191696389 | 0,008233 | 0,04484 |
| Stam | 20844 | 169,54373 | -5,949378113 | 2,251963639 | 0,008245 | 0,044881 |
| Mt3 | 17751 | 5482,6251 | -3,545247953 | 1,342278159 | 0,008261 | 0,044942 |
| Ppp1r3g | 76487 | 56,815029 | 5,692167514 | 2,155783192 | 0,00828 | 0,045009 |
| AI987944 | 233168 | 87,524166 | 6,367216495 | 2,411511179 | 0,008282 | 0,045009 |
| Tmem60 | 212090 | 151,80771 | 5,412052161 | 2,050196815 | 0,008296 | 0,04503 |
| Txndc16 | 70561 | 44,833082 | 5,851013954 | 2,216602531 | 0,0083 | 0,04503 |
| Snx33 | 235406 | 17,274196 | 7,164729957 | 2,71404033 | 0,008294 | 0,04503 |
| Srsf10 | 14105 | 271,26503 | 4,167601203 | 1,58013801 | 0,008352 | 0,04529 |
| Gm30992 | 1,03E+08 | 19,428254 | -5,37061129 | 2,036899845 | 0,008373 | 0,045327 |
| Slc25a42 | 73095 | 16,466819 | -6,446317806 | 2,44487184 | 0,008372 | 0,045327 |
| Cnot2 | 72068 | 94,248669 | 6,816220505 | 2,58490264 | 0,008366 | 0,045327 |
| Dhtkd1 | 209692 | 67,359479 | 5,427937005 | 2,058913464 | 0,008381 | 0,045349 |
| Pum2 | 80913 | 280,63791 | 4,672189882 | 1,772763331 | 0,0084 | 0,045378 |
| Tatdn3 | 68972 | 53,721054 | 6,497447312 | 2,465045193 | 0,008393 | 0,045378 |
| Tmem25 | 71687 | 71,928896 | 6,559494038 | 2,488721886 | 0,008397 | 0,045378 |
| Gm46967 | 1,08E+08 | 25,055628 | 6,613119821 | 2,509450415 | 0,008407 | 0,045388 |
| Nol4l | 329540 | 47,317956 | -5,863029582 | 2,225575338 | 0,008429 | 0,045483 |
| Dancr | 70036 | 30,550804 | 7,076007878 | 2,686572737 | 0,008442 | 0,045532 |

| | | | | | | |
|---|---|---|---|---|---|---|
| Tmem181c-ps | 1E+08 | 25,656104 | 5,1151504 | 1,942666203 | 0,008462 | 0,045545 |
| Nrbp2 | 223649 | 397,25053 | 4,363376188 | 1,657281289 | 0,008467 | 0,045545 |
| Ezr | 22350 | 648,23275 | 4,706766231 | 1,788110883 | 0,008482 | 0,045545 |
| Nap1l1 | 53605 | 409,87651 | -4,509244538 | 1,712388466 | 0,008456 | 0,045545 |
| Snapc5 | 330959 | 130,34983 | 4,93010854 | 1,872759314 | 0,008475 | 0,045545 |
| Ddrgk1 | 77006 | 318,62424 | 4,631965874 | 1,758942124 | 0,008454 | 0,045545 |
| Cfap69 | 207686 | 67,480724 | 5,934725904 | 2,254296864 | 0,008473 | 0,045545 |
| Maff | 17133 | 31,190372 | 6,319660386 | 2,400813559 | 0,008481 | 0,045545 |
| Pgd | 110208 | 37,639623 | 5,834258565 | 2,216589934 | 0,008486 | 0,045545 |
| Gem | 14579 | 54,263939 | 6,632378786 | 2,520168647 | 0,008495 | 0,04557 |
| Gpsm1 | 67839 | 71,621655 | 5,020138479 | 1,907942411 | 0,008509 | 0,045618 |
| Sgf29 | 75565 | 41,687158 | -6,885275972 | 2,617205134 | 0,008519 | 0,045648 |
| Tigd2 | 68140 | 54,693003 | 5,975199421 | 2,271474482 | 0,008525 | 0,045655 |
| Micu3 | 78506 | 231,10613 | 5,399184901 | 2,052788989 | 0,008534 | 0,045676 |
| Dll1 | 13388 | 19,748832 | 7,945926295 | 3,021246168 | 0,008538 | 0,045676 |
| Slc25a34 | 384071 | 34,361782 | 6,633631704 | 2,522778675 | 0,008551 | 0,045721 |
| Rundc1 | 217201 | 26,559382 | 6,590584773 | 2,506718751 | 0,008559 | 0,045741 |
| Ccdc91 | 67015 | 25,982097 | 6,790076812 | 2,582943923 | 0,008568 | 0,045764 |
| Brd8 | 78656 | 372,26005 | 4,44688705 | 1,692421146 | 0,008601 | 0,045863 |
| Smim11 | 68936 | 79,437783 | 5,617916632 | 2,138064589 | 0,0086 | 0,045863 |
| Nop58 | 55989 | 41,805885 | 5,78717175 | 2,202337507 | 0,008595 | 0,045863 |
| Wwc1 | 211652 | 1064,2256 | -4,6401655 | 1,766419949 | 0,008617 | 0,045888 |
| Ftl1-ps2 | 434624 | 10,891907 | -5,915294145 | 2,251978111 | 0,008621 | 0,045888 |
| Fktn | 246179 | 140,58961 | 5,876223419 | 2,237341751 | 0,008629 | 0,045888 |
| Hps3 | 12807 | 65,585215 | 6,808564432 | 2,592215821 | 0,008626 | 0,045888 |
| Ttc30a1 | 78802 | 13,563244 | 7,706861175 | 2,933912018 | 0,008619 | 0,045888 |
| Cebpg | 12611 | 39,115064 | 6,212301479 | 2,365600899 | 0,008637 | 0,045908 |
| Hhip | 15245 | 53,515207 | 6,380063571 | 2,430282265 | 0,008659 | 0,046 |
| Cds2 | 110911 | 225,94686 | 4,748224849 | 1,809700161 | 0,008696 | 0,046175 |
| Eif4e | 13684 | 130,55754 | 3,959712503 | 1,509820011 | 0,008725 | 0,046265 |
| Lman2 | 66890 | 333,46836 | -5,510458459 | 2,101178815 | 0,008727 | 0,046265 |
| Lgals1 | 16852 | 111,65841 | 6,264568737 | 2,388508817 | 0,008721 | 0,046265 |
| Timmdc1 | 76916 | 42,348116 | 4,802623478 | 1,832643571 | 0,008778 | 0,046507 |

| | | | | | | |
|---|---|---|---|---|---|---|
| Atp2b2 | 11941 | 53,497572 | 4,983937278 | 1,902314721 | 0,008795 | 0,046548 |
| Mrrf | 67871 | 197,5178 | 7,42132924 | 2,832611976 | 0,008794 | 0,046548 |
| Cox14 | 66379 | 174,30883 | 5,041636482 | 1,9251903 | 0,008825 | 0,046657 |
| Chkb | 12651 | 33,18188 | 6,537104212 | 2,496109669 | 0,008821 | 0,046657 |
| Apoe | 11816 | 39626,801 | -3,409209192 | 1,302376734 | 0,008853 | 0,046781 |
| Rnase4 | 58809 | 49,431864 | 6,17476626 | 2,359226008 | 0,008863 | 0,046811 |
| Eef2k | 13631 | 25,648503 | -6,059535681 | 2,316257684 | 0,008894 | 0,04695 |
| Pbx4 | 80720 | 21,88202 | 6,705558974 | 2,563842575 | 0,008911 | 0,047016 |
| Dcun1d2 | 102323 | 59,984626 | 5,506967939 | 2,106592342 | 0,008945 | 0,047166 |
| A830019P07Rik | 329056 | 37,880947 | -6,091062272 | 2,330895043 | 0,00897 | 0,047251 |
| Gcdh | 270076 | 260,95469 | 5,822991201 | 2,228193346 | 0,008967 | 0,047251 |
| Szrd1 | 213491 | 150,24316 | -5,179657669 | 1,983210688 | 0,009008 | 0,047424 |
| Sugct | 192136 | 11,733521 | -8,02614626 | 3,073372996 | 0,009014 | 0,047432 |
| Gm32790 | 1,03E+08 | 865,30576 | -5,252097112 | 2,011561296 | 0,009029 | 0,047484 |
| Nsun6 | 74455 | 130,89802 | 4,672098567 | 1,789569573 | 0,009035 | 0,047489 |
| Erg28 | 58520 | 109,8795 | -4,897108589 | 1,876525699 | 0,009063 | 0,047613 |
| Mfap3l | 71306 | 987,8914 | -5,039811731 | 1,931836244 | 0,009086 | 0,047706 |
| Lztr1 | 66863 | 67,387423 | 5,047021587 | 1,934731153 | 0,00909 | 0,047706 |
| Ndufs4 | 17993 | 151,64889 | -5,082814959 | 1,949100309 | 0,009113 | 0,047801 |
| Mef2a | 17258 | 76,628634 | 4,465467183 | 1,713237141 | 0,009149 | 0,047962 |
| Septin1 | 54204 | 35,643697 | 6,174635521 | 2,369528559 | 0,009165 | 0,04802 |
| Gnao1 | 14681 | 4974,3446 | -4,073685063 | 1,563743115 | 0,009185 | 0,048102 |
| Hikeshi | 67669 | 83,722278 | 6,384083549 | 2,451473767 | 0,009209 | 0,048204 |
| Tctn3 | 67590 | 129,679 | 6,842894496 | 2,627976489 | 0,009218 | 0,048223 |
| Arhgef2 | 16800 | 94,999631 | 5,47024839 | 2,101278305 | 0,009233 | 0,048261 |
| Ogfod1 | 270086 | 48,846902 | 7,006537845 | 2,691475912 | 0,009235 | 0,048261 |
| Epm2a | 13853 | 11,820425 | 5,411819977 | 2,079889966 | 0,009269 | 0,048388 |
| Gm51431 | 1,15E+08 | 56,197281 | -4,779648853 | 1,836917696 | 0,009268 | 0,048388 |
| Hspa2 | 15512 | 462,50884 | 3,921578541 | 1,508039071 | 0,00931 | 0,048578 |
| Vtn | 22370 | 552,32815 | 4,896855041 | 1,883673332 | 0,009332 | 0,048669 |
| Zfp420 | 233058 | 52,070051 | 6,697747141 | 2,576919028 | 0,009346 | 0,048714 |
| Cdk16 | 18555 | 15,622053 | 6,15381519 | 2,368005279 | 0,009357 | 0,048745 |
| 4430402I18Rik | 381218 | 143,61468 | 5,67556516 | 2,184278022 | 0,009367 | 0,048771 |

| | | | | | | |
|---|---|---|---|---|---|---|
| Ppil1 | 68816 | 37,461333 | 6,763482067 | 2,604386718 | 0,009405 | 0,048946 |
| Fut10 | 171167 | 149,5297 | 6,024842539 | 2,320586743 | 0,009425 | 0,048995 |
| Phf8 | 320595 | 43,407317 | -6,370119174 | 2,453514579 | 0,009423 | 0,048995 |
| Eif2b5 | 224045 | 11,08913 | 6,73940163 | 2,597013907 | 0,009457 | 0,04914 |
| Bcl2l13 | 94044 | 53,374665 | 5,270597704 | 2,031262366 | 0,009466 | 0,04916 |
| Lrif1 | 321000 | 643,01502 | -6,23181162 | 2,402209565 | 0,009481 | 0,049212 |
| Eif3k | 73830 | 83,857745 | 5,132334603 | 1,978771404 | 0,009495 | 0,049258 |
| Ric1 | 226089 | 124,49331 | 5,294255651 | 2,042032725 | 0,009524 | 0,049384 |
| Extl1 | 56219 | 37,458974 | 4,606755849 | 1,777048118 | 0,009532 | 0,049398 |
| Pstk | 214580 | 30,00254 | 6,843627304 | 2,640768948 | 0,009555 | 0,049492 |
| Pim1 | 18712 | 40,458047 | 5,42390743 | 2,093255526 | 0,009566 | 0,049523 |
| Add1 | 11518 | 2667,999 | -4,434277602 | 1,7116595 | 0,00958 | 0,04957 |
| Bfsp2 | 107993 | 39,078621 | -6,490483676 | 2,50564974 | 0,009588 | 0,049586 |
| Pelp1 | 75273 | 38,01786 | -6,081023184 | 2,347898594 | 0,009598 | 0,049611 |
| Gstp-ps | 1E+08 | 64,637459 | -4,339337111 | 1,67567892 | 0,009609 | 0,049615 |
| Zswim7 | 69747 | 35,908101 | -6,978087014 | 2,694485954 | 0,009604 | 0,049615 |
| Avl9 | 78937 | 38,411572 | 5,3161184 | 2,054247818 | 0,009657 | 0,049827 |
| Sec14l2 | 67815 | 24,38006 | 6,417091901 | 2,479768837 | 0,00966 | 0,049827 |
| Rexo4 | 227656 | 67,188266 | 6,264556255 | 2,421200286 | 0,009671 | 0,049853 |
| Vps54 | 245944 | 298,95101 | 5,85918748 | 2,264649531 | 0,009675 | 0,049853 |
| Rps13 | 68052 | 232,76001 | 3,972103782 | 1,535495378 | 0,009686 | 0,049857 |
| Rgs9 | 19739 | 27,843279 | 6,175319499 | 2,387109544 | 0,009683 | 0,049857 |
| Nabp2 | 69917 | 63,454825 | -4,37290942 | 1,691075752 | 0,009713 | 0,049947 |
| Bdh1 | 71911 | 74,203474 | 5,426545469 | 2,098447049 | 0,00971 | 0,049947 |
| Stub1 | 56424 | 72,258233 | 5,867067646 | 2,269115835 | 0,00972 | 0,049959 |

**Table S9.** Differentially expressed genes between Y-A$^+$/G$^+$ and O-A$^+$/G$^+$ astrocytes.

| Gene | Entrez_ID | baseMean | nonNorm_log2FoldChange | nonNorm_lfcSE | pvalue | padj |
|---|---|---|---|---|---|---|
| Coa6 | 67892 | 75,741246 | 24,69024859 | 2,453599702 | 8,06E-24 | 9,30E-20 |
| Cp | 12870 | 191,5301 | 22,66121578 | 2,487307719 | 8,18E-20 | 4,72E-16 |
| C1qa | 12259 | 17,401224 | -25,50539932 | 3,001595564 | 1,94E-17 | 7,47E-14 |
| Ccdc153 | 270150 | 187,77856 | 21,3639188 | 2,549897916 | 5,37E-17 | 1,55E-13 |
| Mlf1 | 17349 | 100,78536 | 21,12352185 | 2,573957685 | 2,27E-16 | 5,25E-13 |
| Skp2 | 27401 | 71,449556 | 22,80334447 | 2,795502161 | 3,43E-16 | 6,60E-13 |
| Fxyd6 | 59095 | 31,315546 | 20,28796338 | 2,627534328 | 1,15E-14 | 1,86E-11 |
| Pcdhgc3 | 93706 | 16,24265 | 23,75456031 | 3,082222474 | 1,29E-14 | 1,86E-11 |
| Vwa3a | 233813 | 51,507226 | 20,7065069 | 2,717443633 | 2,54E-14 | 3,26E-11 |
| 1700001L19Rik | 69315 | 67,110989 | 21,71127235 | 2,900163468 | 7,09E-14 | 7,57E-11 |
| Mcm3 | 17215 | 32,184892 | 19,3534343 | 2,589096219 | 7,72E-14 | 7,57E-11 |
| B230206L02Rik | 100039440 | 13,532208 | 21,29283718 | 2,84949001 | 7,87E-14 | 7,57E-11 |
| Zic1 | 22771 | 108,0344 | 18,32408857 | 2,485261556 | 1,67E-13 | 1,48E-10 |
| Zfp60 | 22718 | 73,698068 | 19,29324608 | 2,63600563 | 2,50E-13 | 2,06E-10 |
| Ap1s1 | 11769 | 13,406532 | 19,83162386 | 2,755299016 | 6,13E-13 | 4,71E-10 |
| Prpf31 | 68988 | 12,030027 | 21,99813717 | 3,151451871 | 2,95E-12 | 2,12E-09 |
| Gm16124 | 102636154 | 26,520751 | 18,75051725 | 2,760799226 | 1,11E-11 | 7,22E-09 |
| Spint2 | 20733 | 8,3005205 | 21,28353248 | 3,134861256 | 1,13E-11 | 7,22E-09 |
| Rnf128 | 66889 | 33,059008 | 18,44256831 | 2,74965778 | 1,98E-11 | 1,21E-08 |
| Spa17 | 20686 | 62,462287 | 19,67488576 | 2,981226922 | 4,12E-11 | 2,28E-08 |
| Cyb5r4 | 266690 | 7,4656695 | 21,39424405 | 3,242076079 | 4,14E-11 | 2,28E-08 |
| Cabcoco1 | 73287 | 144,36432 | 17,37018612 | 2,75326799 | 2,81E-10 | 1,47E-07 |
| Ccdc180 | 381522 | 200,07472 | 17,05340784 | 2,749188872 | 5,54E-10 | 2,78E-07 |
| Iqca | 74918 | 106,54843 | 18,1886445 | 2,936458477 | 5,86E-10 | 2,82E-07 |
| Pm20d1 | 212933 | 91,374552 | 25,96416178 | 4,320507472 | 1,86E-09 | 8,59E-07 |
| Fam166c | 75434 | 5,7988381 | 19,84765315 | 3,370872998 | 3,91E-09 | 1,74E-06 |
| 1700102H20Rik | 68230 | 9,9466143 | 20,23595433 | 3,443189649 | 4,18E-09 | 1,78E-06 |
| Gtf3a | 66596 | 9,7117852 | 20,48557631 | 3,533299527 | 6,72E-09 | 2,77E-06 |
| Pcdhga3 | 93711 | 20,955391 | 24,93477869 | 4,31804648 | 7,72E-09 | 3,07E-06 |

| | | | | | | |
|---|---|---|---|---|---|---|
| Npy | 109648 | 17,010744 | 19,11797611 | 3,348654715 | 1,14E-08 | 4,37E-06 |
| Cntf | 12803 | 9,6642923 | 19,84626475 | 3,565700206 | 2,61E-08 | 9,71E-06 |
| S100a11 | 20195 | 6,7092767 | 19,67136554 | 3,637512244 | 6,38E-08 | 2,30E-05 |
| 4930539J05Rik | 319587 | 6,8489051 | 17,48984238 | 3,386010144 | 2,40E-07 | 8,39E-05 |
| Pcdhga7 | 93715 | 84,084567 | 21,65785163 | 4,276593709 | 4,10E-07 | 0,0001392 |
| Rhbdl2 | 230726 | 34,291344 | 21,70438719 | 4,361480794 | 6,48E-07 | 0,0002136 |
| Tekt1 | 21689 | 191,65377 | 21,40353496 | 4,363914816 | 9,36E-07 | 0,0003001 |
| Fam183b | 75429 | 46,15117 | 21,09550628 | 4,338692475 | 1,16E-06 | 0,0003622 |
| Meig1 | 104362 | 82,43991 | 20,81105261 | 4,321370888 | 1,47E-06 | 0,0004452 |
| Ager | 11596 | 2,7921461 | 18,35298977 | 3,857568464 | 1,96E-06 | 0,0005796 |
| Sntn | 218739 | 263,68439 | 20,52425247 | 4,318705163 | 2,01E-06 | 0,00058 |
| Gm39468 | 105243583 | 19,65908 | -20,49997069 | 4,370531228 | 2,73E-06 | 0,0007672 |
| Laptm5 | 16792 | 42,567483 | -11,88832792 | 2,540927562 | 2,89E-06 | 0,0007932 |
| Csf1r | 12978 | 27,8148 | -12,61907439 | 2,807828057 | 6,98E-06 | 0,001874 |
| Ipo7 | 233726 | 55,148158 | 9,480274637 | 2,159481689 | 1,13E-05 | 0,0029724 |
| Morn5 | 75495 | 78,723104 | 18,94336815 | 4,365330983 | 1,43E-05 | 0,0036628 |
| Pcdhga8 | 93716 | 17,577333 | 18,65786269 | 4,324426514 | 1,60E-05 | 0,0040132 |
| Crygn | 214301 | 17,878726 | 18,340448 | 4,365517176 | 2,65E-05 | 0,0065199 |
| Fam227b | 75823 | 97,403026 | 18,28483519 | 4,365455278 | 2,81E-05 | 0,0067513 |
| Capsl | 75568 | 112,571 | 18,2085495 | 4,365593041 | 3,03E-05 | 0,0071459 |
| Ginm1 | 215751 | 239,88101 | 7,907012548 | 1,917613579 | 3,73E-05 | 0,0086192 |
| Serping1 | 12258 | 40,939568 | 17,55326761 | 4,369087667 | 5,88E-05 | 0,0133056 |
| Ctu2 | 66965 | 614,85052 | -10,45903932 | 2,680516949 | 9,55E-05 | 0,0211864 |
| Efcab1 | 66793 | 154,61346 | 10,89965281 | 2,799973579 | 9,91E-05 | 0,0215827 |
| Lrrc6 | 54562 | 43,953167 | 16,82725349 | 4,373110839 | 0,000119 | 0,0254649 |
| Gins4 | 109145 | 37,670464 | -8,962963775 | 2,350323924 | 0,000137 | 0,0287525 |
| Gm36908 | 102640972 | 21,900686 | 8,526764024 | 2,242809607 | 0,000144 | 0,0296043 |
| Trp53inp2 | 68728 | 319,17505 | 7,308721711 | 1,928410782 | 0,000151 | 0,0305019 |
| Ppp1r32 | 67752 | 296,85767 | 16,51317555 | 4,364607145 | 0,000155 | 0,030782 |
| Tfb1m | 224481 | 64,900942 | 10,23715001 | 2,719487609 | 0,000167 | 0,0326641 |
| Tmem212 | 208613 | 397,92475 | 9,639798345 | 2,564413679 | 0,000171 | 0,032808 |
| Fam120aos | 68128 | 174,95124 | 6,665477391 | 1,787541589 | 0,000192 | 0,0352405 |
| Tbc1d24 | 224617 | 19,515674 | 8,532421958 | 2,28613443 | 0,00019 | 0,0352405 |

| | | | | | | |
|---|---|---|---|---|---|---|
| Msl3l2 | 73390 | 54,571834 | 8,369863216 | 2,241662235 | 0,000189 | 0,0352405 |
| Adamts20 | 223838 | 71,810735 | 8,928700347 | 2,421678325 | 0,000227 | 0,0409242 |
| Nrep | 27528 | 84,043187 | 9,132312614 | 2,48055232 | 0,000232 | 0,0411619 |
| Dglucy | 217830 | 25,888668 | 8,398127257 | 2,305866966 | 0,00027 | 0,0472969 |
| Fars2 | 69955 | 71,361227 | 7,213790656 | 1,988860638 | 0,000287 | 0,0493766 |
